# Novel Alphaproteobacteria transcribe genes for nitric oxide transformation at high levels in a marine oxygen deficient zone

**DOI:** 10.1101/2023.11.21.568154

**Authors:** Claire E. Elbon, Frank J. Stewart, Jennifer B. Glass

## Abstract

Marine oxygen deficient zones (ODZs) are portions of the ocean where intense nitrogen loss occurs primarily via denitrification and anammox. Despite many decades of study, the identity of the microbes that catalyze nitrogen loss in ODZs are still being elucidated. Intriguingly, high transcription of genes in the same family as the nitric oxide dismutase (*nod*) gene from *Methylomirabilota* has been reported in the anoxic core of ODZs. Here, we show that the most abundantly transcribed *nod* genes in the Eastern Tropical North Pacific ODZ belong to a new order (UBA11136) of Alphaproteobacteria, rather than *Methylomirabilota* as previously assumed. Gammaproteobacteria and Planctomycetia also transcribe *nod*, but at lower relative abundance than UBA11136 in the upper ODZ. The *nod*-transcribing Alphaproteobacteria likely use formaldehyde and formate as a source of electrons for aerobic respiration, with additional electrons possibly from sulfide oxidation. They also transcribe multiheme cytochrome (here named *ptd*) genes for a putative porin-cytochrome protein complex of unknown function, potentially involved in extracellular electron transfer. Molecular oxygen for aerobic respiration may originate from nitric oxide dismutation via cryptic oxygen cycling. Our results implicate Alphaproteobacteria order UBA11136 as a significant player in marine nitrogen loss and highlight their potential in one-carbon, nitrogen, and sulfur metabolism in ODZs.

**Significance statement:** In marine oxygen deficient zones, microbes transform bioavailable nitrogen to gaseous nitrogen, with nitric oxide as a key intermediate. The Eastern Tropical North Pacific contains the world’s largest oxygen deficient zone, but the identity of the microbes transforming nitric oxide remain unknown. Here, we show that highly transcribed nitric oxide dismutase *(nod*) genes belong to Alphaproteobacteria of the novel order UBA11136, which lacks cultivated isolates. These Alphaproteobacteria show evidence for aerobic respiration, using oxygen potentially sourced from nitric oxide dismutase, and possess a novel porin-cytochrome protein complex with unknown function. Gammaproteobacteria and Planctomycetia transcribe *nod* at lower levels. Our results pinpoint the microbes mediating a key step in marine nitrogen loss and reveal an unexpected predicted metabolism for marine Alphaproteobacteria.

## Introduction

Marine oxygen deficient zones (ODZs) contribute up to half of the ocean’s nitrogen loss (DeVries et al., 2013) and are a major source of marine emissions of the potent greenhouse gas nitrous oxide (N_2_O) (Yang et al., 2020). The primary source of the N_2_O at the oxic-anoxic interface and in anoxic waters in ODZs is denitrification (Babbin et al., 2015; Frey et al., 2020). The microbial enzyme responsible for N_2_O production during denitrification is nitric oxide reductase (Nor), which uses electrons from cytochrome *c* (cNor) or quinol (qNor), to reduce nitric oxide (NO) to N_2_O (Wasser et al., 2002; Zumft, 2005; Kraft et al., 2011). In the qNor family, there are *bona fide* qNor enzymes and NO dismutase (NOD). NOD proteins lack the quinol-binding site, seemingly preventing the enzyme from taking up external electrons; instead, NOD is theorized to disproportionate NO into dinitrogen and O_2_ in methane-oxidizing *Methylomirabilota* bacteria (Ettwig et al., 2010; Ettwig et al., 2012) and in the alkane-oxidizing gammaproteobacterium HdN1 (Zedelius et al., 2011).

The Eastern Tropical North and South Pacific (ETNP and ETSP) ODZs are the world’s largest and second largest ODZs, and the subjects of extensive microbial ecology studies. Abundant NO reductase-like genes and transcripts in the ETNP and ETSP ODZ cluster in the same enzyme subfamily as NOD (Dalsgaard et al., 2014; Ganesh et al., 2014; Padilla et al., 2016; Fuchsman et al., 2017). Due to the similarity of ODZ Nod proteins to those of *Methylomirabilota* (NC10), it was initially presumed that ODZ bacteria also used Nod proteins to disproportionate NO into N_2_ and O_2_ for use in intra-aerobic methane oxidation (Dalsgaard et al., 2014; Padilla et al., 2016; Thamdrup et al., 2019). However, Fuchsman et al. (2017) found that the peak of *nod* gene abundance in the ETNP ODZ correlates with a peak of modeled N_2_O production (Babbin et al., 2015) and does not correlate with abundance of methane monooxygenase genes, suggesting that Nod proteins in the ETNP ODZ are potentially an important source of N_2_O, and are unlikely to be involved in methane oxidation. The plausibility that Nod proteins can reduce NO to N_2_O is supported by a study of a novel eukaryotic denitrification pathway in foraminifera (*Globobulimina* spp.) that produces N_2_O while expressing Nod (Woehle et al., 2018). Yet, the phylogenetic identity and metabolic context of marine Nod proteins, which are a key biological source of either N_2_O or O_2_+N_2_ in marine ODZs, remain unresolved.

In this study, we sought to determine the identity, predicted metabolism, and environmental niche of the ODZ organism responsible for the highly transcribed *nod* genes first discovered in Padilla et al. (2016). We found that the most abundantly transcribed *nod* genes in the ETNP ODZ belong to Alphaproteobacteria in the novel order UBA11136. Significant transcription of *nod* genes was limited to waters with <1 μM O_2_. These *nod*-transcribing alphaproteobacteria also transcribe genes involved in aerobic respiration, which was unexpected given that they inhabit anoxic waters, as well as genes involved in oxidation of formaldehyde, likely indicating methylotrophy. Genes encoding multi-heme cytochrome proteins potentially implicated in nitrogen or iron cycling were also transcribed.

## Results

### Transcribed nod sequences in the ETNP ODZ belong to Alphaproteobacteria, Gammaproteobacteria, and Planctomycetia

Our reanalysis of highly transcribed *nod* genes in the ETNP ODZ (Padilla et al., 2016) shows that these genes belong to Alphaproteobacteria rather than a member of *Methylomirabilota* as previously assumed. Querying the Nod amino acid sequences from Padilla et al. (2016) against ETNP ODZ metagenomes in the IMG-JGI database returned multiple 100% identity matches, including a *nod* gene (Ga0066848_100037855) on a scaffold with hypothetical genes with 100% identity to Alphaproteobacteria metagenome-assembled genomes (MAGs) from the ETNP ODZ (Uzun et al., 2020) **(Table S1)**. We binned previously sequenced ETNP ODZ metagenomes Ga0066848 (ETNP201310SV72) and Ga0066829 (ETNP201306SV43) (Ruiz-Perez et al., 2021) into MAGs. Contigs with the most highly transcribed *nod* genes were present in two Alphaproteobacteria MAGs (GTDB taxonomy: UBA11136 sp002686135; species representative: Rhodospirillaceae bacterium isolate ARS27) with 97% average nucleotide identity. Querying the Nod amino acid sequences from Padilla et al. (2016) against NCBI’s non-redundant protein database returned matches to other MAGs assigned to Alphaproteobacteria order UBA11136 from low-oxygen marine settings (ETNP, Saanich Inlet, and the Black Sea; 78-80% identity), the marine magnetotactic alphaproteobacterium *Magnetovibrio blakemorei* MV-1 (75% identity), Gammaproteobacterium HdN1 (66% identity), and *Methylomirabilota* spp. (66% identity; **Table S2**).

To glean additional insights into evolutionary relationships, we updated a previous Nod phylogeny (Hu et al., 2019) with additional amino acid sequences from marine MAGs (Tully et al., 2018; Cabello-Yeves et al., 2021; Lin et al., 2021) and ETNP ODZ metagenomes (Ruiz-Perez et al., 2021), subdivided in free-living cells (0.2-1.6 micron) and cells from the particle fraction (>1.6 micron; **Figure 1A; Table S3)**. The Nod topology was generally consistent with a previous phylogeny from Fuchsman et al. (2017), with additional taxonomic data from MAGs in the TARA oceans dataset further constraining Nod placement (Tully et al., 2018). As expected based on the binning and BLAST results, the Nod sequence from Padilla et al. (2016) (Ga0066848_100037855) clustered phylogenetically with marine Alphaproteobacteria (OTU III in Fuchsman et al. (2017), hereafter “Alpha-type Nod”); this clade contained three unique sequences, all of which were present in multiple metagenomes and all from the free-living fraction, and one of which was identical to that of Rhodospirillaceae NP1106 (MBV28360). Four unique ODZ Nod sequences clustered with marine Gammaproteobacteria (OTU II in Fuchsman et al. (2017), hereafter “Gamma-type Nod”); these sequences were monophyletic with a cluster of Gammaproteobacteria Nod cluster sequences from sewage sludge, including Gammaproteobacterium HdN1 (Ehrenreich et al., 2000) and other wastewater Gammaproteobacteria. Multiple ETNP ODZ metagenomes contained Gamma-type Nod sequences identical to those of Gammaproteobacteria NP964 (MBP20251). Gamma-type Nod had ∼70% identity to Alpha-type Nod. Several ODZ Nod sequences, all from the particle fraction (“PF”) clustered with marine Deltaproteobacteria in a clade of monophyletic *nod* genes from groundwater Methylomirabilis, Deltaproteobacteria, and Acidobacteria MAGs (∼65% identity to Alpha-type Nod). Six unique Nod ODZ protein sequences (two of which were present in multiple metagenomes) clustered with Planctomycetia (OTU I in Fuchsman et al. (2017), hereafter “Planctomycetia-type Nod”), and were primarily found in free-living cells (“FL”) and had ∼40% identity to Alpha-type Nod. Intriguingly, two ODZ sequences clustered in the eukaryotic *Globobulimina* clade (∼50% identity to Alpha-type Nod). Viral Nod sequences from Saanich Inlet (∼55% identity to Alpha-Nod) clustered with the viral Nod sequence previously reported by Gazitúa et al. (2021) from the Eastern Tropical South Pacific ODZ (St16 OMZ 317E – viral).

**Figure 1.**
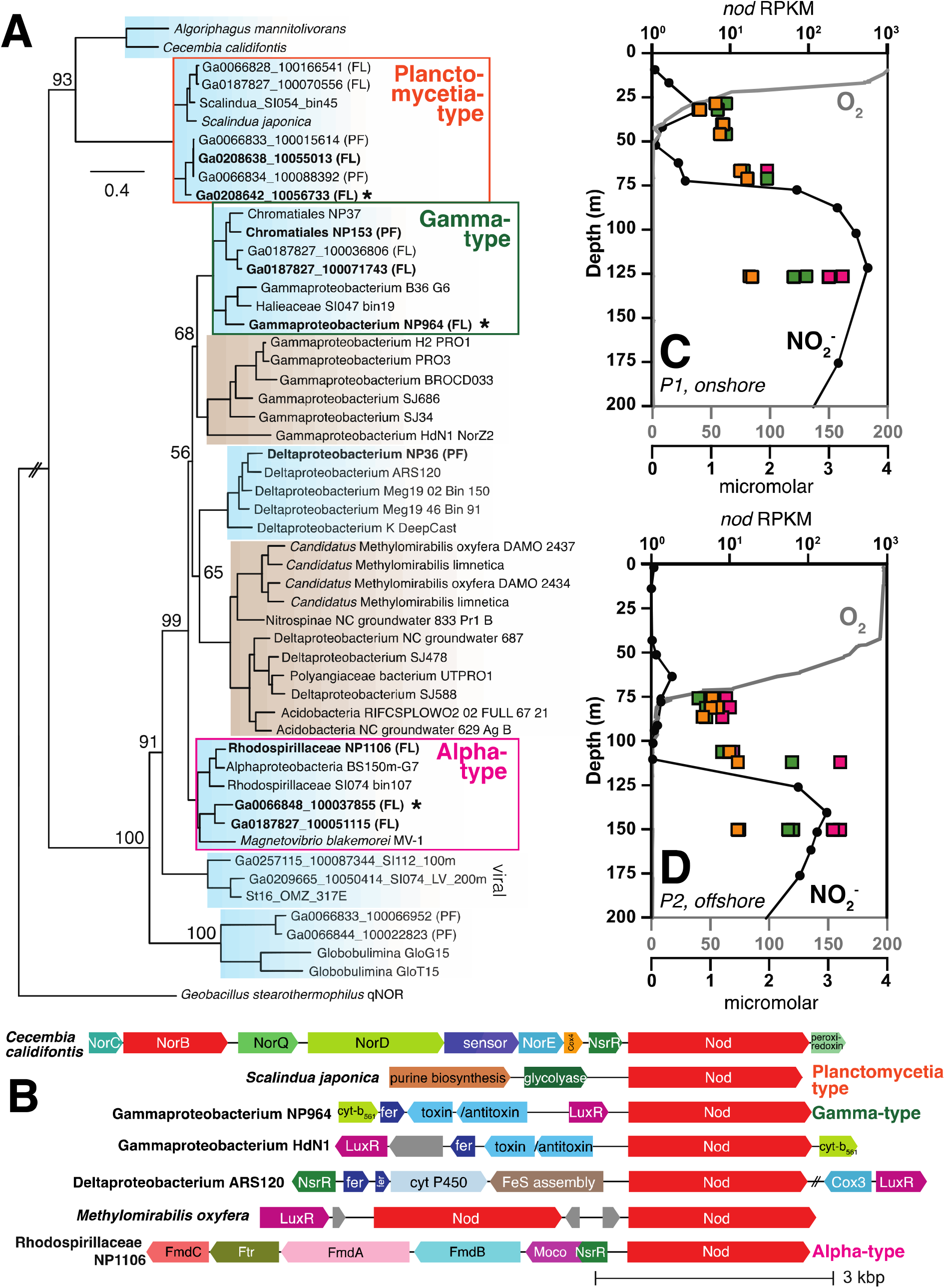
Marine Nod clades, gene neighborhoods, and depth profiles of transcription. (A) Maximum likelihood phylogeny of nitric oxide dismutase (Nod) amino acid sequences in marine (blue) and select terrestrial (brown) taxa, primarily from marine MAGs (Tully et al., 2018; Cabello-Yeves et al., 2021; Lin et al., 2021) and ETNP ODZ metagenomes (Ruiz-Perez et al., 2021). Branch support was evaluated using 1000 rapid bootstrap replicates, with bootstrap values shown for deep branches. The tree is drawn to scale, with branch lengths in number of substitutions per site. Bold sequences represent those present in multiple ETNP ODZ metagenomes (see **Table S3** for duplicate accession numbers). “PF” indicates genes from the particle fraction (>1.6 micron fraction) of filters. “FL” indicates genes from the free-living fraction (0.2-1.6 micron) collected on Sterivex filters. The most highly transcribed ETNP ODZ sequence is indicated with an asterisk. The qNor sequence *Geobacillus stearothermophilus* was used as the outgroup. (B) Gene neighborhoods surrounding *nod* genes in select taxa. GenBank contigs: *Cecembia calfifontis* SGXG01000001, *Scalindua japonica* BAOS01000045, Gammaproteobacteria NP964 PBRC01000062, Gammaproteobacterium HdN1 FP929140, Deltaproteobacteria NZCL01000067, *Candidatus* Methylomirabilis oxyfera FP565575, and *Rhodospirillaceae* NP1106 PCBZ01000014. Unlabeled gray genes are hypothetical. (C, D) Oxygen and nitrite concentrations (circles), and *nod* transcripts (squares, as reads per kilobase per million mapped reads (RKPM)) with depth in ETNP ODZ P1 (onshore) and P2 (offshore) sites (Mattes et al., 2022).

We investigated gene neighborhoods surrounding ODZ *nod* genes in the three main phylogenetic clusters of ODZ sequences: Planctomycetia-type Nod, Gamma-type Nod, and Alpha-type Nod. Whereas “unknown Nor-related” marine Bacteroidota sequences were located on an operon with other *nor* genes, there was no consistent gene neighborhood for *nod* sequences **(Figure 1B)**. Planctomycetia-type *nod* genes were not located in the vicinity of any genes with recognizable related function. Gamma-type *nod* gene neighborhoods contained ferredoxins and cytochrome *b*_561_ genes for electron transport. Upstream of the Alpha-type *nod* in *Rhodospirillaceae* NP1106 is a cluster of formylmethanofuran dehydrogenase genes (*fmd/fwd*) used in C1 metabolism via tetrahydromethanopterin/methanofuran-linked reactions.

Immediately upstream or downstream of *nod* genes, helix-turn-helix transcriptional regulators were common **(Figure 1B)**. Neighboring Gamma-type and *Methylomirabilis nod* genes, LuxR-type regulators were common; these regulators have diverse functions and their potential connection to Nod remains unclear. Neighboring Alpha-type and Bacteroidota (e.g. *Cecembia calidifontis*) *nod* genes, Rrf2-type regulators were present. The protein NsrR in the Rrf2 family regulates global cellular response to NO toxification by directly sensing NO with an iron-sulfur cluster (Bodenmiller and Spiro, 2006; Tucker et al., 2010). The presence of this NsrR-like regulator suggests that Nod in marine Alphaproteobacteria and Bacteroidota may be involved in nitrosative stress response and NO detoxification.

### Alphaproteobacterial nod is highly transcribed in anoxic waters

We assessed transcription of Alpha, Gamma*-*, and Planctomycetia-type *nod* genes from the oxycline to upper ODZ (secondary nitrite maximum) using ETNP ODZ metatranscriptomes from an onshore station with a shallower oxycline (P1; **Figure 1C**) and an offshore station with a deeper oxycline (P2; **Figure 1D**) (Mattes et al., 2022). In both oxyclines, transcription was low (4-10 reads per kilobase per million mapped reads (RPKM), n=8) for all three *nod* types (**Figure 1C, D)**. Below the oxyclines, *nod* transcripts began to rise and were highest at the secondary nitrite maxima, with Alpha-type (184-274 RPKM, n=4) > Gamma-type (55-95 RPKM, n=4) > Planctomycetia-type (13-19 RPKM, n=4; **Table S4)**.

### MAGs with highly transcribed nod gene represent a new order of Alphaproteobacteria

In order to assess the phylogeny of the *nod*-containing Alphaproteobacteria MAGs, we constructed an alphaproteobacterial phylogeny using the conserved protein NADH ubiquinone oxidoreductase subunit L (NuoL) as in Cevallos and Degli Esposti (2022), with additional representation of order UBA11136 including our MAG ETNP2013_S10_300m_22 (**Figure 2)**. MAG ETNP2013_S06_300m_15 was not included in the phylogeny because its *nuoL* gene was truncated. UBA11136 is situated in the phylogeny near other orders found in ODZs.

**Figure 2.**
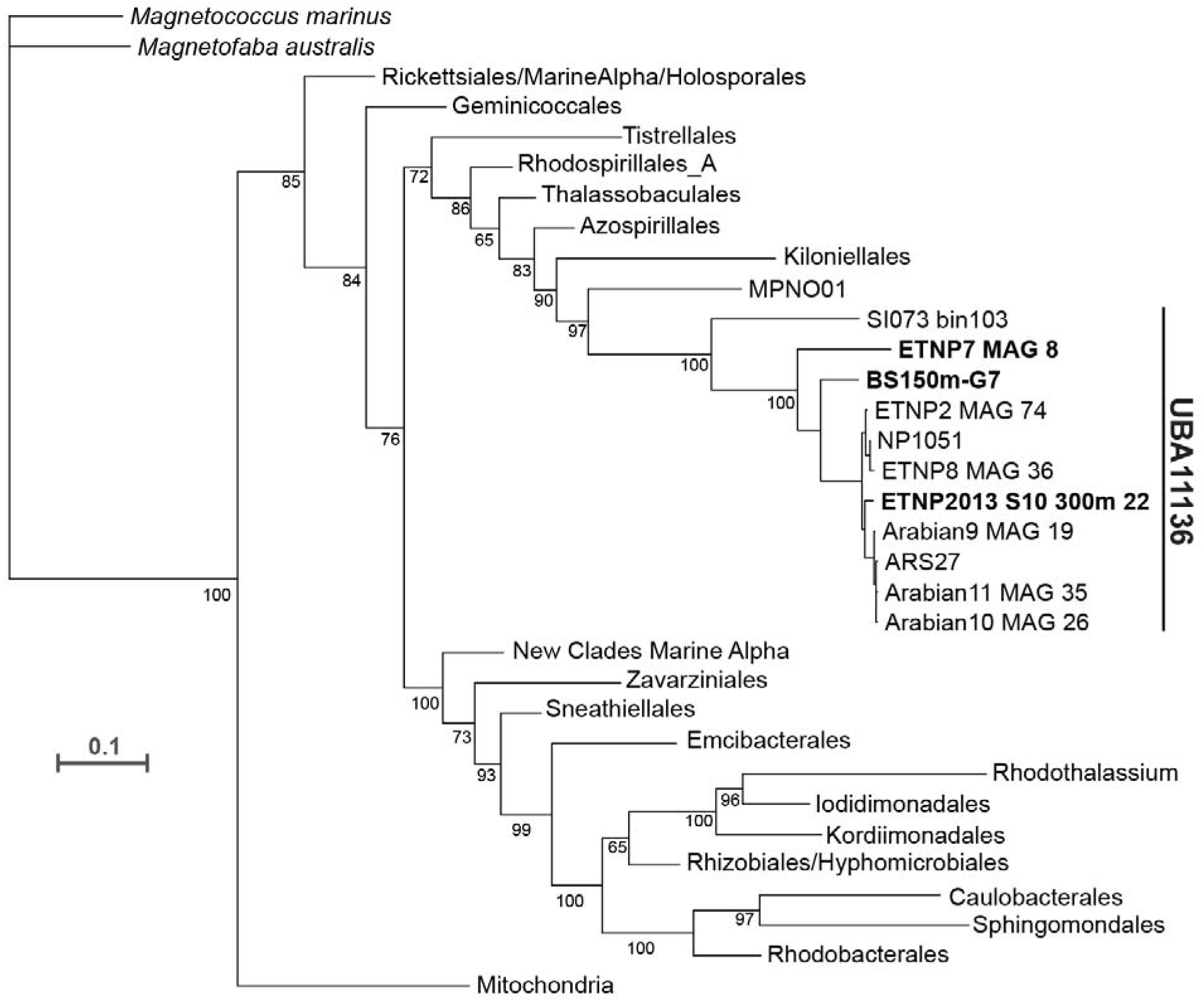
Phylogeny of Alphaproteobacteria showing order UBA11136, with *nod*-containing MAGs. The phylogeny was constructed using the Alphaproteobacterial phylogenetic marker NADH ubiquinone oxidoreductase subunit L as in Cevallos and Esposti (2022). Order UBA11136 is expanded and *nod*-containing MAGs are bolded. Taxonomic names are from Cevallos and Esposti (2022) and GTDB Release 08-RS214. The scale bar represents amino acid substitutions per site. The full phylogeny is shown in **Supplementary Figure S1**.

### Alphaproteobacteria transcribe genes for formate metabolism, aerobic respiration, and a multiheme cytochrome complex

To glean insight into potential roles for Nod in cellular context, we sought to reconstruct the electron transport chain of the Alphaproteobacteria with the most highly transcribed *nod* genes (Alphaproteobacterium MAG ETNP2013_S10_300m_22 and Alphaproteobacterium MAG ETNP2013_S06_300m_15, 73% and 69% estimated completeness, respectively) at the secondary nitrite maximum. Of total metagenomic reads, 0.38% map to ETNP2013_S10_300m_22 and 0.39% map to ETNP2013_S06_300m_15. In both MAGs, *nod* was in the top three most transcribed genes in the ETNP ODZ (∼44,000 FPKM; **Table S5)**, after a bacterial nucleoid DNA-binding protein and a potassium gated channel protein. In addition to *nod*, we found that genes for formaldehyde oxidation via tetrahydromethanopterin/methanofuran-linked reactions, including formylmethanofuran dehydrogenase (*fwd/fmd*) and formylmethanofuran--tetrahydromethanopterin N-formyltransferase (*ftr*), were transcribed in both MAGs **(Table S5)**. Both MAGs also transcribed NAD-dependent formate dehydrogenase **(Table S5)**. Thus, the alphaproteobacterium appears to be capable of conversion of formaldehyde to formate and use of formate as a source of electrons for NADH:ubiquinone oxidoreductase (Complex I; **Figure 3)**. The source of formaldehyde is likely methanol oxidation, as pyrroloquinoline quinone (PQQ)-dependent ethanol/methanol dehydrogenases were found in Alphaproteobacteria MAGs from low-oxygen marine settings **(Table S6)**. Methane monooxygenase genes were not found in the partial Alphaproteobacteria MAGs, precluding our ability to rule out the possibility of these genes in the missing portions of the genomes. The Alphaproteobacteria PQQ-dependent dehydrogenase genes contained the motif DYDG **(Table S6)**, which is characteristic of the lanthanide-containing form of the enzymes rather than calcium form (Keltjens et al., 2014).

**Figure 3.**
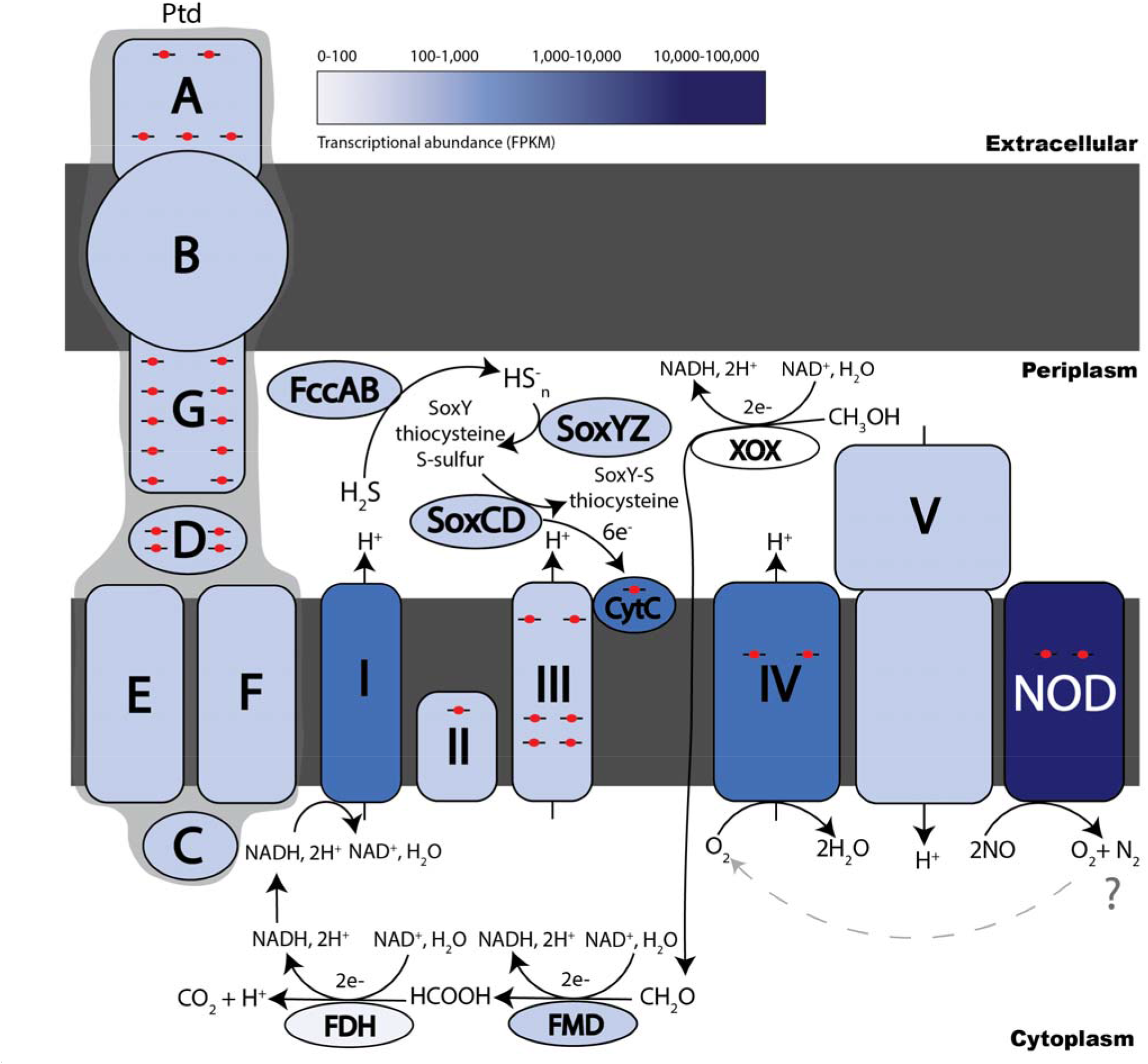
Schematic of the electron transport chain in *nod*-containing ODZ Alphaproteobacteria. Enzymes were included based on presence and transcriptional activity of metagenome-assembled genomes (MAGs) assigned to Alphaproteobacteria (GTDB taxonomy: UBA11136 sp002686135; see text). The color of each protein is chosen according to transcriptional activity and represented from 0-100, 100-1,000, 1,000-10,000, and 10,000-100,000 FPKM in gradient from lighter to darker blue (Table S5). Heme proteins are indicated by red circular hemes with the cartoon number corresponding to the number of actual hemes present on each protein. Hypothetical Ptd proteins are labelled A, B, C, D, E, F, and G, and location within the cell is determined using Psort bacterial localization prediction tool (Table S8). ETC complexes I-V found in Alphaproteobacteria MAGs are labelled with proposed interactions between formate oxidation and complex I NADH electron transfer. Highly transcribed NOD protein and predicted O_2_ generation is shown as feeding into A1 type CCO complex IV reduction. Additional electrons for CytC and the ETC are proposed to come from sulfur oxidation carried out by the flavocytochrome *c* sulfide dehydrogenase (FccAB, FCC), and sulfane-sulfur dehydrogenase (SoxCD) with the multi-enzyme carrier complex (SoxYZ).

A full aerobic electron transport chain (Complex I, II, III, and IV) and F0F1-type ATP synthase were transcribed in both bins **(Figure 3; Table S5)**. Complex IV (cytochrome c oxidase) was type A1 according to the Sousa et al. (2012) classification, and the *cox* operon in the GTDB species representative Rhodospirallaceae ARS27 was subtype b (COX2-COX1-CtaB-CtaG_Cox11-COX3-DUF983-SURF1-CtaA1-M32-Tsy-M16B) according to the Geiger et al. (2023) classification. Sulfur oxidation genes, including flavocytochrome c sulfide dehydrogenase (FccAB), sulfane hydrogenase (SoxCD), and carrier protein SoxYZ, were also transcribed, as were numerous transposes **(Figure 3; Table S5)**.

Genes for a multiheme cytochrome complex were transcribed in both bins. To our knowledge, this putative operon has not previously been described. Hereafter, we designate it the *ptdABCDEFG* operon for its sequence of <u>p</u>enta/tetra/<u>d</u>eca-heme proteins, interspersed with other conserved proteins. *ptdAB* genes are highly transcribed in our Alphaproteobacteria MAGs, but it is unclear if the rest of the operon is also highly transcribed, because it was truncated in our MAGs’ scaffolds. The *ptd* gene cluster consists of a penta-heme protein with a C-terminal beta-sandwich (PtdA), a porin (PtdB), a FAD/NAD(P)-binding oxidoreductase (PtdC), a periplasmic tetra-heme protein (PtdD), a cyclic nucleotide-binding domain protein with two 4Fe-4S clusters (PtdE), a cytoplasmic transmembrane ferric reductase-like protein (PtdF), and a periplasmic deca-heme protein (PtdG; **Figure 3; Tables S7, S8)**. The function of this complex is unknown, but the presence of genes encoding a porin and multiple multiheme proteins resembles porin-cytochrome protein complexes involved in extracellular reduction electron transfer during Fe(III) and Mn(IV) reduction (Richardson et al., 2012; Shi et al., 2014). PtdA has a homolog to a penta-heme cytochrome c_552_ protein of unknown function in a thermophilic purple sulfur gammaproteobacterium (Chen et al., 2019) and is in the same COG family (COG3303) as formate dependent nitrite reductase, NrfA. *ptdABCDEFG* genes were prevalent in Alphaproteobacteria, Gammaproteobacteria, Nitrospirales, and Planctomycetes MAGs from marine or high salinity environments **(Figure 4; Table S7)**.

**Figure 4.**
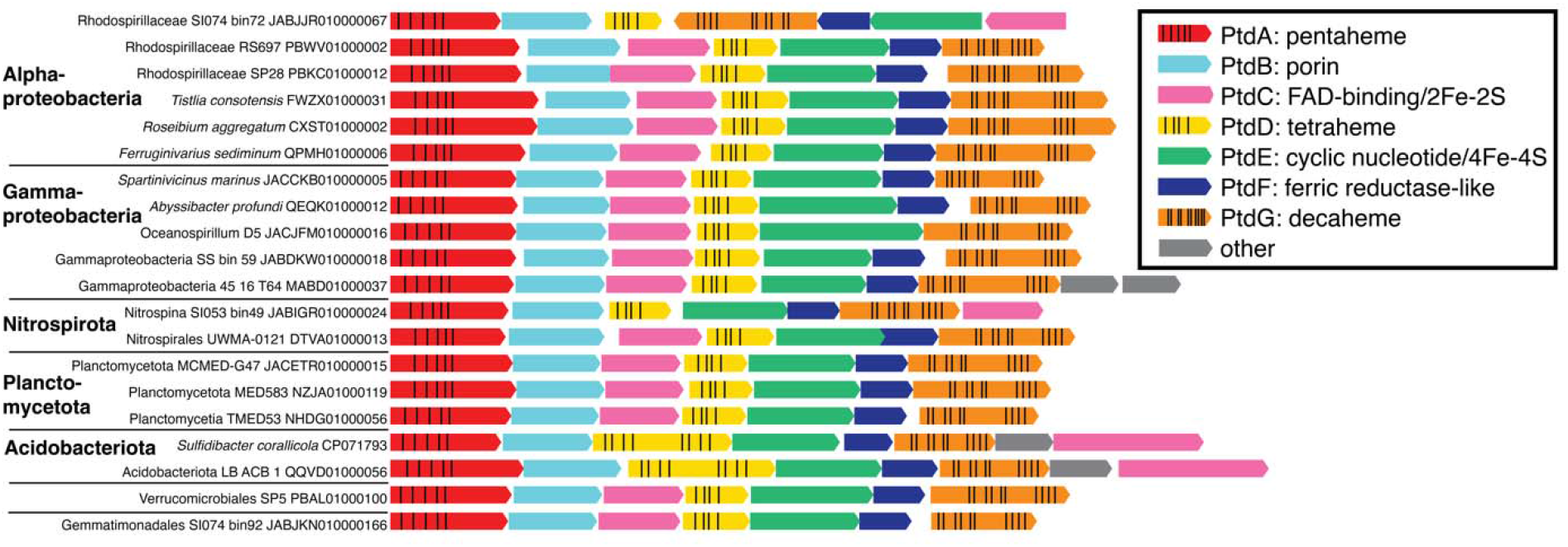
Gene neighborhoods of petaheme-tetraheme-decaheme genes from select organisms. Depicted heme spacing is approximate. All organisms are from saline environments (seawater, marine sediment, or saline spring).

## Discussion

This study predicts the previously ambiguous identity of the microorganisms that make the dominant nitric oxide-transforming protein (Nod) in the world’s largest ODZ, the Eastern Tropical North Pacific. Extensive horizontal gene transfer of *nod* genes between microbial genomes is evident from the lack of conservation of gene neighborhood and patchy phylogeny (Fuchsman et al., 2017), which may be mediated by viral infection (Gazitúa et al., 2021). We found that the most transcriptionally active *nod* genes in the ETNP upper ODZ belong to the novel Alphaproteobacteria order UBA11136. Alpha-type *nod* transcript abundances (∼200 RPKM) are similar to those of dissimilatory nitrate reductase (*narG*) in the ODZ (Tsementzi et al., 2016). The *nod*-transcribing Alphaproteobacteria are also transcribing genes for formaldehyde oxidation, likely as a source of electrons to the respiratory chain via NAD reduction by formate dehydrogenase. Sulfide may be used as a supplemental electron donor and/or may be concomitantly oxidized for detoxification (Callbeck et al., 2021; Schmitz et al., 2023).

Our discovery of a putative porin-cytochrome complex (*ptd* operon) in marine bacteria was unexpected. Porin-cytochrome complexes have been best studied for their role in extracellular electron transport, particularly for respiratory metal reduction and oxidation (Richardson et al., 2012; Shi et al., 2014). It is conceivable that the Ptd complex is involved in iron reduction in ODZs; there is iron reduction at the secondary nitrite maximum and it is hypothesized to be bacterially mediated, but the microbes involved have yet to be determined (Moffett et al., 2007; Glass et al., 2015). Alternatively, the presence of *ptdABCDEFG* genes in numerous nitrite-oxidizing bacteria (Nitrospirales) could imply the involvement of these genes in nitrogen cycling; PtdA was in the same COG family as formate-dependent nitrite reductase (Simon et al., 2000) and PtdC is similar to a flavohemoprotein with predicted nitric oxide dioxygenase activity, also annotated as hydroxylamine oxidoreductase-linked cytochrome. The function of PtdABCDEFG remains completely unknown and requires future biochemical characterization.

On the other end of the electron transport chain, high transcription of a heme/copper terminal oxidase suggests that O_2_ is being used as the terminal election acceptor in *nod*-transcribing Alphaproteobacteria MAGs. The transcribed heme/copper oxidase is A1-type (low O_2_ affinity), also present in mitochondria, and adapted for high O_2_ concentrations. Low O_2_ affinity A1-type heme/copper oxidases are transcribed in other anoxic environments (Berg et al., 2022). Because ODZs have extremely low concentrations of molecular oxygen below the oxycline, O_2_ for aerobic respiration may be generated *in situ* and rapidly consumed. Given that the function of Nod is proposed to be dismutation of two NO molecules into N_2_ and O_2_ (Ettwig et al., 2010), it is possible that the O_2_ source for aerobic respiration in the UBA11136 MAGs is NO dismutation, although other sources of O_2_ (e.g. *in situ* photosynthesis, mixing) in anoxic waters are also conceivable (Garcia-Robledo et al., 2017). The physiological uses of Gamma-type and Planctomycetia-type Nod may be different from Alpha-type Nod, although this remains to be investigated.

The source of NO, the presumed substrate for Nod, may be generated in the same organism using Nod, or generated by a different organism (or chemical pathway). Nitric oxide was positively correlated with nitrite in the Eastern Tropical South Pacific ODZ, and was only detectable when O_2_ was <1-2 μM (Lutterbeck et al., 2018). In the Eastern Tropical North Pacific, NO concentration and turnover rates were elevated at O_2_ <100 μM (Ward and Zafiriou, 1988). Both studies suggest that the NO in ODZs likely originates from nitrification or nitrifier denitrification, while genomic analyses indicate that the copper-containing nitrite reductase (*nirK*) in SAR11 bacteria (presumably performing denitrification) may be a key source of NO (Fuchsman et al., 2017). Because most ODZ denitrifiers specialize in only one of the three steps (NO_2_ ^-^ reduction, NO reduction, and N_2_O reduction) (Zhang et al., 2023), and known nitrite reductases were not identified in our MAGs, existing data indicate that the NO that is used as a substrate for alphaproteobacterial Nod is not generated *in vivo*. (Only 4 out of 32 *nod*-containing MAGs contained a nitrite reductase gene: two Gammaproteobacteria MAGs contained *nirK*, one Myxococcota MAG contained *nirS*, and one Scalindua MAG contained *nirS*). It is also possible that another uncharacterized enzyme produces NO.

This study suggests that marine Alphaproteobacteria from order UBA11136 are actively reducing NO under anoxia, as implied by their abundant transcription of *nod* genes. While there is strong evidence that the substrate for Nod in ODZs is NO based on its abundance, the products of this enzyme (N_2_O vs. N_2_+O_2_) remain uncertain. Nod is theorized to disproportionate NO into N_2_ and O_2_ in methane-oxidizing *Methylomirabilota* bacteria (Ettwig et al., 2010; Ettwig et al., 2012), but no biochemical characterizations of Nod have been published to date, and foraminifera expressing Nod produce N_2_O (Woehle et al., 2018). The apparent lack of other denitrification genes in *nod*-transcribing Alphaproteobacteria is consistent with the observation that denitrification in ODZs is largely divided into distinct microbial taxa (Dalsgaard et al., 2014; Fuchsman et al., 2017; Zhang et al., 2023). For example, although nitrate reductase (*narG*) genes are widely distributed amongst ODZ microbes (Zhang et al., 2023), SAR11 bacteria appear to dominate in *narG* transcriptional activity (Tsementzi et al., 2016). Our finding that the transcription of *nod* is catalyzed primarily by marine Alphaproteobacteria implies that this taxa contributes significantly to marine nitrogen loss.

## METHODS

### Nod phylogeny and gene neighborhood

Amino acid sequences of highly transcribed *nod* genes “ETNP 2014 Stn10 150m” and “ETNP 2013 Stn6 300m” were acquired from the authors of Padilla et al. (2016) (see Table S2 for sequences). These sequences were used for BLASTP searches of ODZ metagenomes in the IMG-JGI database and the non-redundant protein (nr) database in NCBI. Sequences (n=53, 731 gap-free sites) were aligned using the MAFFT online server with the L-INS-i method (Katoh et al., 2019). A phylogeny was generated with 1000 bootstraps using model LG+I+G4 using W-IQ-Tree (Trifinopoulos et al., 2016). The phylogeny was visualized using FigTree v.1.4.4, and the fasta file (Nod_alignment) is available as a supplemental dataset. Gene neighborhoods were generated using the EFI Gene Neighborhood Tool (Zallot et al., 2019) with single sequence BLAST of the UniProt database using the amino acid sequence Ga0066848_100037855 (JGI IMG) as the Nod query with an e-value cutoff of 10^-5^ and with 10 genes upstream and downstream the gene of interest.

### Transcription of nod genes in ETNP ODZ depth profiles

Magic Basic Local Alignment Search Tool (Boratyn et al., 2019) was used to search ETNP ODZ metatranscriptomes (PRJNA727903; Mattes et al. (2022)) using representative nucleotide sequences for Planctomycetia-like (Ga0066826_100064333 (JGI IMG)), Gamma-like (PBRC01000062.1:19833-22205 (NCBI)), and Alpha-like (Ga0066848_100037855 (JGI IMG)) *nod* genes. Default parameters were used except for the score threshold (18). Read hits were normalized to reads per kilobase million (RPKM).

### Metagenomic binning

Binning of metagenome-assembled genomes (MAGs) was performed using the KBase platform (Arkin et al., 2018). ETNP ODZ metagenomes were collected in 2013 and sequenced by Joint Genome Institute (JGI) using an Illumina HiSeq 2500 as described in Ruiz-Perez et al. (2021). Assemblies for the ETNP ODZ metagenomes (Ruiz-Perez et al., 2021) containing Alpha-type *nod* genes (ETNP201310SV72 (GOLD Analysis Project ID Ga0066848; stn10 300m) and ETNP201306SV43 (GOLD Analysis Project ID Ga0066829; stn6 300m) were imported from JGI IMG into Kbase. Metagenomic assemblies were binned into MAGs using MaxBin2 v2.2.4 (Wu et al., 2016). The two MAGs containing *nod* genes (MAG ETNP2013_S10_300m_22 from ETNP201310SV72 and ETNP2013_S06_300m_15 from ETNP201306SV43) were selected for further analysis. Average nucleotide identity was calculated using FastANI (Jain et al., 2018). MAG taxonomy and genome quality was evaluated by GTDB-Tk v2.3.2 (Chaumeil et al., 2022). MAGs were annotated with RASTtk v1.073 (Brettin et al., 2015). Metagenomic reads were mapped to MAGs using Bowtie2 (Langmead and Salzberg, 2012).

### Alphaproteobacterial NuoL phylogeny

Alphaproteobacterial NADH ubiquinone oxidoreductase subunit L (NuoL) and mitochondrial ND5 marker proteins (n=320) were aligned as in Cevallos and Degli Esposti (2022) with additional representation of UBA11136. A maximum likelihood phylogeny with 1000 bootstraps was constructed using IQ-tree (Nguyen et al., 2015) using the LG+F model with ultrafast bootstrap (Hoang et al., 2017). Taxonomic names and clades are from Cevallos and Esposti (2022) and GTDB Release 08-RS214. The fasta file (NuoL_alignment) is available as a supplemental dataset. Alphaproteobacteria MAGs containing *nod* genes (Table S2) were classified as belonging to order UBA11136 using GTDB-Tk v2.3.2 (Chaumeil et al., 2022).

### Mapping transcripts to metagenomic bins

Metatranscriptomic mapping to MAGs was performed using the KBase platform (Arkin et al., 2018). RNA-seq data (Mattes et al., 2022) were imported from the depth with highest *nod* transcription, the secondary nitrite maximum (126 meters, NCBI run SRR14460584) and aligned to MAGs using the Bowtie2 (Langmead and Salzberg, 2012) app in KBase. The Cufflinks v2.2.1 (Trapnell et al., 2012) app in Kbase was then used to assemble the aligned RNA-seq data into a set of transcripts and to calculate the relative abundances of the transcripts expressed in fragments per kilobase of exon per million fragments mapped (FPKM).

### Cellular localization and heme numbers

Cellular locations of Ptd proteins were predicted using PSORTb v3.0.3 analysis (Yu et al., 2010). Numbers of heme-binding motifs per protein were identified by counting CXXCH sequences. Ptd gene neighborhoods was generated using the EFI Gene Neighborhood Tool (Zallot et al., 2019) with single sequence BLAST of the UniProt database using the amino acid sequence Ga0066848_100031354 (JGI IMG) as the PtdA query, with an e-value cutoff of 10^-5^ and with 10 genes upstream and downstream the gene of interest.

## Supporting information

Nod alignment

NuoL alignment

## Data availability

The Kbase bioinformatic pipeline and MAGs are at https://narrative.kbase.us/narrative/106999. Original metagenomic reads are available at BioProject PRJNA375524 (ETNP201306SV43, SAMN06344130) and BioProject PRJNA375542 (ETNP201310SV72, SAMN06344148). MAG Alphaproteobacteria bacterium ETNP2013_S06_300m_15 was deposited into BioProject PRJNA375524 (BioSample SAMN38229257, WGS Accession JAZDBU000000000) and Alphaproteobacteria bacterium ETNP2013_S10_300m_22 was deposited into BioProject PRJNA375542 (BioSample SAMN38228782, WGS Accession JAZDCE000000000). All ETNP ODZ datasets used in this manuscript are listed in **Table S9**.

## Acknowledgements

We thank Laura Bristow for helpful discussions. We thank Mauro Degli Esposti for assistance with the NADH ubiquinone oxidoreductase subunit L phylogeny. We thank Cory Padilla, Anthony Bertagnolli, and Neha Sarode for sharing previous data. We acknowledge funding from an NSF Graduate Research Fellowship to CEE, the Simons Foundation, and NSF Awards 2022991 and 2054927 to FJS.

## Notes

https://narrative.kbase.us/narrative/106999

https://www.ncbi.nlm.nih.gov/bioproject/PRJNA375524/

https://www.ncbi.nlm.nih.gov/bioproject/PRJNA375542/

### Competing Interest Statement

The authors have declared no competing interest.

### Summary of Updates

Tempered our language to clarify that the functions are predictions only; changed the title as suggested by the reviewers; removed the old terminology Rhodospirillaceae and replaced it with the new GTDB order names; created a phylogeny (Figure 2, Figure S1) for the Alphaproteobacteria MAGs showing the new order they below to (UBA11136); mapped metagenomic reads to the Alphaproteobacteria MAGs and added the findings to the results; added additional citations for previously published metagenomes reanalyzed here; added an additional supplemental table (S9) with accession numbers, and latitude and longitude for all the datasets used in the manuscript; updated multiple sections of the methods with additional details; changed the coloring on Figure 4 (ptd gene synteny); additional numerous minor edits as suggested by the reviewers.

## References

Arkin, A.P., Cottingham, R.W., Henry, C.S., Harris, N.L., Stevens, R.L., Maslov, S. et al. (2018) KBase: The United States Department of Energy Systems Biology Knowledgebase. Nat Biotechnol 36: 566–569.

Babbin, A.R., Bianchi, D., Jayakumar, A., and Ward, B.B. (2015) Rapid nitrous oxide cycling in the suboxic ocean. Science 348: 1127–1129.

Berg, J.S., Ahmerkamp, S., Pjevac, P., Hausmann, B., Milucka, J., and Kuypers, M.M.M. (2022) How low can they go? Aerobic respiration by microorganisms under apparent anoxia. FEMS Microbiol Rev 46: 1–14.

Bodenmiller, D.M., and Spiro, S. (2006) The yjeB (nsrR) gene of Escherichia coli encodes a nitric oxide-sensitive transcriptional regulator. J Bacteriol 188: 874–881.

Boratyn, G.M., Thierry-Mieg, J., Thierry-Mieg, D., Busby, B., and Madden, T.L. (2019) Magic-BLAST, an accurate RNA-seq aligner for long and short reads. BMC Bioinf 20: 405.

Brettin, T., Davis, J.J., Disz, T., Edwards, R.A., Gerdes, S., Olsen, G.J. et al. (2015) RASTtk: a modular and extensible implementation of the RAST algorithm for building custom annotation pipelines and annotating batches of genomes. Sci Rep 5: 8365.

Cabello-Yeves, P.J., Callieri, C., Picazo, A., Mehrshad, M., Haro-Moreno, J.M., Roda-Garcia, J.J. et al. (2021) The microbiome of the Black Sea water column analyzed by shotgun and genome centric metagenomics. Environ Microbiome 16: 5.

Callbeck, C.M., Canfield, D.E., Kuypers, M.M.M., Yilmaz, P., Lavik, G., Thamdrup, B. et al. (2021) Sulfur cycling in oceanic oxygen minimum zones. Limnol Oceanogr 66: 2360–2392.

Cevallos, M.A., and Degli Esposti, M. (2022) New Alphaproteobacteria thrive in the depths of the ocean with oxygen gradient. Microorganisms 10: 455.

Chaumeil, P.A., Mussig, A.J., Hugenholtz, P., and Parks, D.H. (2022) GTDB-Tk v2: memory friendly classification with the genome taxonomy database. Bioinformatics 38: 5315–5316.

Chen, J.H., Yu, L.J., Boussac, A., Wang-Otomo, Z.Y., Kuang, T., and Shen, J.R. (2019) Properties and structure of a low-potential, penta-heme cytochrome c(552) from a thermophilic purple sulfur photosynthetic bacterium Thermochromatium tepidum. Photosynth Res 139: 281–293.

Dalsgaard, T., Stewart, F.J., Thamdrup, B., De Brabandere, L., Revsbech, N.P., Ulloa, O. et al. (2014) Oxygen at nanomolar levels reversibly suppresses process rates and gene expression in anammox and denitrification in the oxygen minimum zone off northern Chile. mBio 5: e01966.

DeVries, T., Deutsch, C., Rafter, P.A., and Primeau, F. (2013) Marine denitrification rates determined from a global 3-D inverse model. Biogeosciences 10: 2481–2496.

Ehrenreich, P., Behrends, A., Harder, J., and Widdel, F. (2000) Anaerobic oxidation of alkanes by newly isolated denitrifying bacteria. Arch Microbiol 173.

Ettwig, K.F., Speth, D.R., Reimann, J., Wu, M.L., Jetten, M.S., and Keltjens, J.T. (2012) Bacterial oxygen production in the dark. Front Microbiol 3: 273.

Ettwig, K.F., Butler, M.K., Le Paslier, D., Pelletier, E., Mangenot, S., Kuypers, M.M.M. et al. (2010) Nitrite-driven anaerobic methane oxidation by oxygenic bacteria. Nature 464: 543–548.

Frey, C., Bange, H.W., Achterberg, E.P., Jayakumar, A., Löscher, C.R., Arévalo-Martínez, D.L. et al. (2020) Regulation of nitrous oxide production in low-oxygen waters off the coast of Peru. Biogeosciences 17: 2263–2287.

Fuchsman, C.A., Devol, A.H., Saunders, J.K., McKay, C., and Rocap, G. (2017) Niche partitioning of the N cycling microbial community of an offshore oxygen deficient zone. Front Microbiol 8: 2384.

Ganesh, S., Parris, D.J., DeLong, E.F., and Stewart, F.J. (2014) Metagenomic analysis of size-fractionated picoplankton in a marine oxygen minimum zone. ISME J 8: 187–211.

Garcia-Robledo, E., Padilla, C.C., Aldunate, M., Stewart, F.J., Ulloa, O., Paulmier, A. et al. (2017) Cryptic oxygen cycling in anoxic marine zones. Proc Natl Acad Sci U S A 114: 8319–8324.

Gazitúa, M.C., Vik, D.R., Roux, S., Gregory, A.C., Bolduc, B., Widner, B. et al. (2021) Potential virus-mediated nitrogen cycling in oxygen-depleted oceanic waters. ISME J 15: 981–998.

Geiger, O., Sanchez-Flores, A., Padilla-Gomez, J., and Esposti, M.D. (2023) Multiple approaches of cellular metabolism define the bacterial ancestry of mitochondria. Sci Adv 9: eadh0066.

Glass, J.B., Kretz, C.B., Ganesh, S., Ranjan, P., Seston, S.L., Buck, K.N. et al. (2015) Meta-omic signatures of microbial metal and nitrogen cycling in marine oxygen minimum zones. Front Microbiol 6: 998.

Hoang, D.T., Chernomor, O., Haeseler, A.v., Minh, B.Q., and Vinh, L.S. (2017) UFBoot2: improving the ultrafast bootstrap approximation. Mol Biol Evol 35: 518–522

Hu, Q.-Q., Zhou, Z.-C., Liu, Y.-F., Zhou, L., Mbadinga, S.M., Liu, J.-F. et al. (2019) High microbial diversity of the nitric oxide dismutation reaction revealed by PCR amplification and analysis of the nod gene. Int Biodeterior Biodegradation 143: 104708.

Jain, C., Rodriguez, R.L., Phillippy, A.M., Konstantinidis, K.T., and Aluru, S. (2018) High throughput ANI analysis of 90K prokaryotic genomes reveals clear species boundaries. Nat Commun 9: 5114.

Katoh, K., Rozewicki, J., and Yamada, K.D. (2019) MAFFT online service: multiple sequence alignment, interactive sequence choice and visualization. Brief Bioinform 20: 1160–1166.

Keltjens, J.T., Pol, A., Reimann, J., and Op den Camp, H.J. (2014) PQQ-dependent methanol dehydrogenases: rare-earth elements make a difference. Appl Microbiol Biotechnol 98: 6163–6183.

Kraft, B., Strous, M., and Tegetmeyer, H.E. (2011) Microbial nitrate respiration–genes, enzymes and environmental distribution. J Biotech 155: 104–117.

Langmead, B., and Salzberg, S.L. (2012) Fast gapped-read alignment with Bowtie 2. Nat Methods 9: 357–359.

Lin, H., Ascher, D.B., Myung, Y., Lamborg, C.H., Hallam, S.J., Gionfriddo, C.M. et al. (2021) Mercury methylation by metabolically versatile and cosmopolitan marine bacteria. ISME J 15: 1810–1825.

Lutterbeck, H.E., Arévalo-Martínez, D.L., Löscher, C.R., and Bange, H.W. (2018) Nitric oxide (NO) in the oxygen minimum zone off Peru. Deep Sea Res Part II Top Stud Oceanogr 156: 148–154.

Mattes, T.E., Burke, S., Rocap, G., and Morris, R.M. (2022) Two metatranscriptomic profiles through low-dissolved-oxygen waters (DO, 0 to 33 μM) in the Eastern Tropical North Pacific Ocean. Microbiol Resour Announc 11: e01201–01221.

Moffett, J.W., Goepfert, T.J., and Naqvi, S.W.A. (2007) Reduced iron associated with secondary nitrite maxima in the Arabian Sea. Deep Sea Res Part I Oceanogr Res Pap 54: 1341–1349.

Nguyen, L.T., Schmidt, H.A., von Haeseler, A., and Minh, B.Q. (2015) IQ-TREE: a fast and effective stochastic algorithm for estimating maximum-likelihood phylogenies. Mol Biol Evol 32: 268–274.

Padilla, C.C., Bristow, L.A., Sarode, N., Garcia-Robledo, E., Ramírez, E.G., Benson, C.R. et al. (2016) NC10 bacteria in marine oxygen minimum zones. ISME J 10: 2067–2071.

Richardson, D.J., Butt, J.N., Fredrickson, J.K., Zachara, J.M., Shi, L., Edwards, M.J. et al. (2012) The ‘porin-cytochrome’ model for microbe-to-mineral electron transfer. Mol Microbiol 85: 201–212.

Ruiz-Perez, C.A., Bertagnolli, A.D., Tsementzi, D., Woyke, T., Stewart, F.J., and Konstantinidis, K.T. (2021) Description of Candidatus Mesopelagibacter carboxydoxydans and Candidatus Anoxipelagibacter denitrificans: Nitrate-reducing SAR11 genera that dominate mesopelagic and anoxic marine zones. Syst Appl Microbiol 44: 126185.

Schmitz, R.A., Peeters, S.H., Mohammadi, S.S., Berben, T., van Erven, T., Iosif, C.A. et al. (2023) Simultaneous sulfide and methane oxidation by an extremophile. Nat Commun 14: 2974.

Shi, L., Fredrickson, J.K., and Zachara, J.M. (2014) Genomic analyses of bacterial porin-cytochrome gene clusters. Front Microbiol 5: 657.

Simon, J., Gross, R., Einsle, O., Kroneck, P.M., Kroger, A., and Klimmek, O. (2000) A NapC/NirT-type cytochrome c (NrfH) is the mediator between the quinone pool and the cytochrome c nitrite reductase of Wolinella succinogenes. Mol Microbiol 35: 686–696.

Sousa, F.L., Alves, R.J., Ribeiro, M.A., Pereira-Leal, J.B., Teixeira, M., and Pereira, M.M. (2012) The superfamily of heme–copper oxygen reductases: Types and evolutionary considerations. Biochim Biophys Acta Bioenerg 1817: 629–637.

Thamdrup, B., Steinsdóttir, H.G.R., Bertagnolli, A.D., Padilla, C.C., Patin, N.V., Garcia-Robledo, E. et al. (2019) Anaerobic methane oxidation is an important sink for methane in the ocean’s largest oxygen minimum zone. Limnol Oceanogr 64: 2569–2585.

Trapnell, C., Roberts, A., Goff, L., Pertea, G., Kim, D., Kelley, D.R. et al. (2012) Differential gene and transcript expression analysis of RNA-seq experiments with TopHat and Cufflinks. Nat Protoc 7: 562–578.

Trifinopoulos, J., Nguyen, L.T., von Haeseler, A., and Minh, B.Q. (2016) W-IQ-TREE: a fast online phylogenetic tool for maximum likelihood analysis. Nucleic Acids Res 44: W232–235.

Tsementzi, D., Wu, J., Deutsch, S., Nath, S., Rodriguez, R.L., Burns, A.S. et al. (2016) SAR11 bacteria linked to ocean anoxia and nitrogen loss. Nature 536: 179–183.

Tucker, N.P., Le Brun, N.E., Dixon, R., and Hutchings, M.I. (2010) There’s NO stopping NsrR, a global regulator of the bacterial NO stress response. Trends Microbiol 18: 149–156.

Tully, B.J., Graham, E.D., and Heidelberg, J.F. (2018) The reconstruction of 2,631 draft metagenome-assembled genomes from the global oceans. Sci Data 5: 1–8.

Uzun, M., Alekseeva, L., Krutkina, M., Koziaeva, V., and Grouzdev, D. (2020) Unravelling the diversity of magnetotactic bacteria through analysis of open genomic databases. Sci Data 7: 252.

Ward, B.B., and Zafiriou, O.C. (1988) Nitrification and nitric oxide in the oxygen minimum of the eastern tropical North Pacific. Deep Sea Res Part A Oceanogr Res Pap 35: 1127–1142.

Wasser, I.M., De Vries, S., Moënne-Loccoz, P., Schröder, I., and Karlin, K.D. (2002) Nitric oxide in biological denitrification: Fe/Cu metalloenzyme and metal complex NO_x_ redox chemistry. Chem Rev 102: 1201–1234.

Woehle, C., Roy, A.S., Glock, N., Wein, T., Weissenbach, J., Rosenstiel, P. et al. (2018) A novel eukaryotic denitrification pathway in foraminifera. Curr Biol 28: 2536–2543 e2535.

Wu, Y.W., Simmons, B.A., and Singer, S.W. (2016) MaxBin 2.0: an automated binning algorithm to recover genomes from multiple metagenomic datasets. Bioinformatics 32: 605–607.

Yang, S., Chang, B.X., Warner, M.J., Weber, T.S., Bourbonnais, A.M., Santoro, A.E. et al. (2020) Global reconstruction reduces the uncertainty of oceanic nitrous oxide emissions and reveals a vigorous seasonal cycle. Proc Natl Acad Sci U S A 117: 11954–11960.

Yu, N.Y., Wagner, J.R., Laird, M.R., Melli, G., Rey, S., Lo, R. et al. (2010) PSORTb 3.0: improved protein subcellular localization prediction with refined localization subcategories and predictive capabilities for all prokaryotes. Bioinformatics 26: 1608–1615.

Zallot, R., Oberg, N., and Gerlt, J.A. (2019) The EFI web resource for genomic enzymology tools: leveraging protein, genome, and metagenome databases to discover novel enzymes and metabolic pathways. Biochemistry 58: 4169–4182.

Zedelius, J., Rabus, R., Grundmann, O., Werner, I., Brodkorb, D., Schreiber, F. et al. (2011) Alkane degradation under anoxic conditions by a nitratelJreducing bacterium with possible involvement of the electron acceptor in substrate activation. Environ Microbiol Rep 3: 125–135.

Zhang, I.H., Sun, X., Jayakumar, A., Fortin, S.G., Ward, B.B., and Babbin, A.R. (2023) Partitioning of the denitrification pathway and other nitrite metabolisms within global oxygen deficient zones. ISME Commun 3: 76.

Zumft, W.G. (2005) Nitric oxide reductases of prokaryotes with emphasis on the respiratory, heme–copper oxidase type. J Inorg Biochem 99: 194–215.

