## Supplemental Figure S1 for "Novel Alphaproteobacteria transcribe genes for nitric oxide transformation at high levels in a marine oxygen deficient zone"

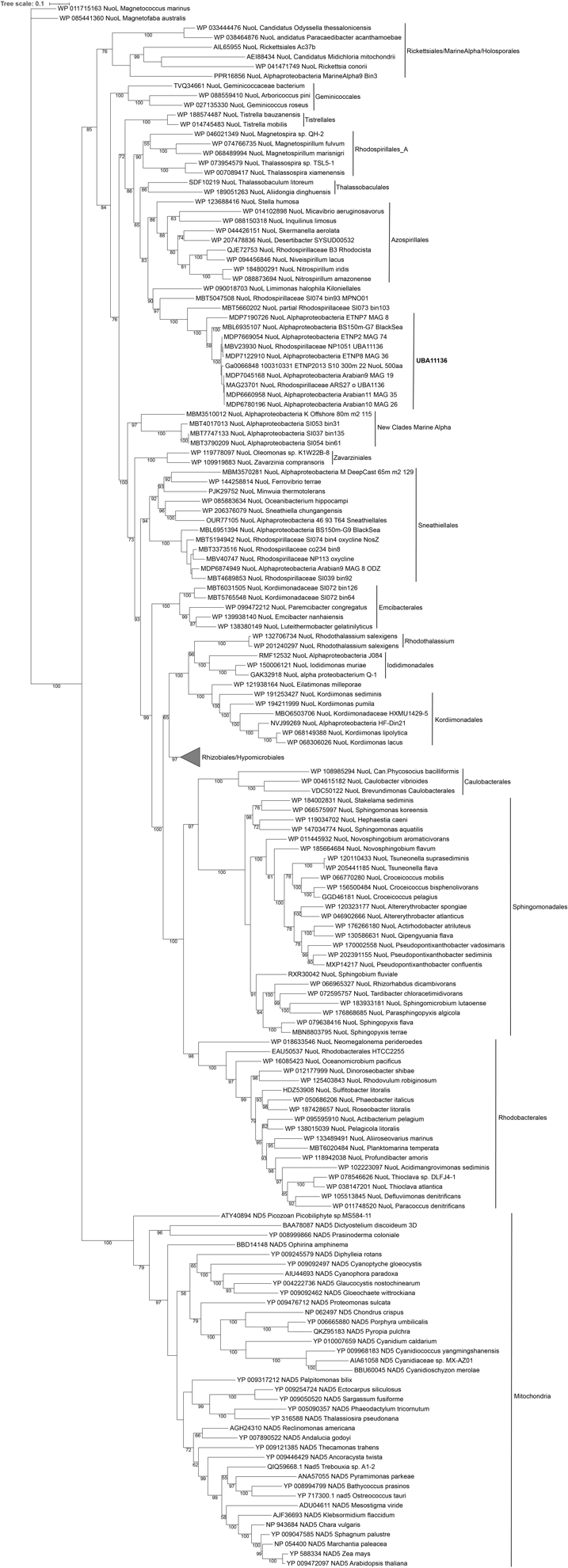

**Supplemental Figure 1. Expanded view (320 sequences) of the Alphaproteobacteria maximum likelihood phylogeny in Figure 2 of the main text.** All clades except Rhizobiales/Hypomicrobiales are expanded.
