## Supplementary material for "Novel Alphaproteobacteria transcribe genes for nitric oxide transformation at high levels in a marine oxygen deficient zone": Nod alignment

```
>Globobulimina_NorI_GloG15_CONTIG49717:890-3289
M-------------------------------------------------------ADVA
PKEGEWILRPAKTNWIATLSNKNYWAFYFLVIVAICVGGIGYF--GEVAHYEVPPLCDFV
--G-PDGSVVISAKSINHGEEVFHLRGLMAYGSFLGDGSERGPDYTAEALHELAIGMTKH
YESKLKHK------------------TQHDVEALKPRVRMELR----NNAYD-LEKDVIV
LNSAQISAWKRIKTYYRRIFAGADQG----------------TSTL---TPN---MFITN
PTDIADLAGFFFWGAWMSAAARPGEDYSYTHNWPHEPLAGNIPTPAVILWSAISVLILWF
GLAATLFIYGQLDHENEDEEDIKT----KGQPLTTQDLET---GVVRETQVDCYKFFVLS
MLAFAIQVIAGVCCAIDFV---RPL------GFS-ACYILGYNVLRSYHATFQIFWFFVA
WVGTTIWFLPRFQRRVPIGQKFLINLLFGGCCVVALGGAIGIPLGQTGYI-QGEMAYYFG
SQGWEFMELGRIFQDLLLCGFVLWILILFRGVYSYITWNT-IWSTPAWVFYGSLVMVWFL
FFSLKVTPKTNFIVSDFWRWMVVHMWVEVTFEVFTTVIVAFLYREMGLVSKKAAERATYI
AVMLFFLTATIGVGHNFYWIAKPTGVIAMGSAFSTTQVMPLILLTMDAWKFIQMRE----
KAELEQAKGNQKHIMRGVWLFMLGVNFWNVFGAGILGSFINLPVVNYYMHSTYLTGNHAH
AAMWGVKGNIAIAGVLFCVQHTVQEKYWSPKIVSVAFWSLNGGIAAQMFLSMFPAGLVHM
YTAMAEGLWAARTHDMYNSTIFQNLAKGRAFGGHIFLWGGLLPLVYFVVSRYFYLK-PVT
P-----NIDKNRKYDS--------F---WLDAAHTAPYSTKK------------------
--LV
>Globobulimina_NorII_GloT15_NODE_10819:151-2556
MST----------------------------------------------------DAKSY
PLIQDFVPSGTRNNWIATLSDKRWWTFYFAIIVAICVSGIGYF--GGVAHYQNPPNCDFV
--D-QNGKVVFPQSMIKHGEEIFHLRGLMSYGSFLGDGSEHGVDYTAEALHMMALSMGDF
YTAQLRQNTSR----------E---LAEHDMEAVKARVRLELR----ANTYE-EARDVVV
LNEAMVYAWKDLEKYHLRMHFEDGFG-----------------DLL----PN---ERFAK
KRDVSDMAAFFYWGGWISVAARPGEDYSYTHNWPHDPLVGNLPTAATILWSTLSVGVLFV
GLGITLFIWGQRKDEIEA--DDGT----TGKPLTTQDLEL---GVVHPTQLACYKFFVLA
MIAFAIQVLGGIACSVDVV---RPG------GLT-LCMWIPYTVLRSYHTTFQIYWFFVA
WVGTTIFFLPRFQKKPPAGQLFLINLLGVGCAIVAAGGLIGIPLGQGGYL-EGDMAYYFG
SQGWEFMELGRAFQDILLGGFVLWIFILFRGVYSYITWDT-IWSTPAWLFYGSLVMVWFL
FFSLKVTPDTDFVVSDFWRWMVVHMWVEVTFEVFTTVIVAYLYREMGLVSKKGAERATYI
AVMLFFLTATIGVGHNFYWIAKPTGVIALGSAFSTTQVMPLILLTLDAWKYIQVSE----
RAELERKKGNQKHVMKAAWLFLLGVNFWNVFGAGILGSFINMPIVNYYMHATYLTGNHAH
GAMFGVKGNVALAGVLFCAQHLVKEEAWSPKLLSLSFWCMNGGIAAQMFLSLFPTGMYHM
YINMMEGLYAARKFEVTQGYIFQTLAKGRAVGGHIFLWGGLLPLVYFILSRAAYLK-DAT
K------NAPGKRYQS--------F---WLDSAHTAGYKVK-------------------
--VY
>Ga0066833_100066952
MTLVV-------------------------------------------------------
-PESVIPKAAYDTNMAGKLANKNYWGVYGILVFAIIVLGTGYM--TGAAIFEVPPLVDFV
--S-ETGEVVLSTSKINHGEEIFHHRGLMSYGSMLGDGSERGPDFTAEALHIMAVSMGEF
YVKKHDGQTDAG---------S---NSTYDMEAIKARVITELH----LNTWNGDDKGVIV
LSEAKISAWQAVRKHYRRVYYREEASVSGSEDVSSADSAGSGRVGL---WSI---ERFAN
PGHADDLAAFFFWGAWCCVANRPSEDYSYTHNWPYDPLAGNTPTPEVMIWSVASLFVLFF
GLMITLYIYGQFPEEEEP---------LTQQPLTTQDLEA---DIVRPTQQATYRFFVLA
IAAFFIQVVSGIACAIDFI---RPG------GFS-VCQFLPFSVFRSYHTTFQIYWFFVA
WVGTTIFFLPRFSK-VPAAQHGLIDLLYIGCVVVALGGVIGIPLGQTGFL-EGPVAYYFG
SQGWEYMELGRFYQDVLLAGFVLWIVIMFRGVWPYLTLRR-AWSPPAWLFYGSIVMVAFL
FFSLKVTPKTNFIISDFWRWMVVHMWVEVTFEVFTTVIVSYLLVEMGLVTRKKAEKTTYI
```

```
AVMLFFLTATIGVGHNFYWIAKPTGVIALGSTFSTTQILPLILVTLDAWKIMQEKS----
RAEAEQKKGNQQYVMKEVFQFVVGINFWNIFGAGVLGSLVNLPIVNYYLHSTYLTGCHAH
GAMFGVKGNVALAGVLFCVQHLVEEKHWSGRLINISFWCLNGGMALMMFLSLFPTGLYQL
FMVIKYGFWYARSAEIVHGPVFQFLLKLRPIGGHIFFEGGLLPLLYFVFTRGFKLK-NET
D----PKSMGKDRYRS--------T---WHIDEADTKK----------------------
--EQ
>Ga0066844_100022823
MTLIA-------------------------------------------------------
-PESGIPKATYDTNMAGKLSNKKYWGVYAIIVIAIIVLGNGYM--TGAAISEVPPLVDFV
--S-ETGEVIMSTSKINHGEEIFHHRGLMSYGSVLGDGSERGPDFTAEALHIMAVSMGEF
YVEKHDGQTDAG---------A---GSAYDMEAIKARVIAELH----LNTWNGDDNGVIV
LNEAKIFAWQAVRKYYRRVYYREEASVSGSED-----------GGL---WPT---ERFAN
PEHADDLAAFFFWGAWCCVANRPSENYSYTHNWPYDPLAGNTPTPEVMIWSVASLFVLFF
GLMITLYIYGQFPEDDEP---------LTQQPLTTQDLEA---DIVRPTQRATYKFFVLA
IAAFFIQVISGIACAIDFI---RPG------GFS-VCEFIPFSVFRSYHTTFQIYWFFVA
WVGATIFFLPRFSK-VPTAQHGLIDLLYIGCIAVALGGVIGIPLGQTGYL-EGPMAYYFG
SQGWEYMELGRFYQDVLLAGFVLWIVIMFRGVWPFLTWKR-VWSPPAWLFYGSVIMVAFL
FFSLKVTPKTNFIISDFWRWMVVHMWVEVTFEVFTTVIVSYLLVEMGLVTRKMAEKTTYI
AVMLFFLTATIGVGHNFYWIAKPTGVIALGSTFSTTQILPLILITLDAWKIMQEKS----
RAEAEQKKGNQMYVMKEVFQFVVAINFWNIFGAGVLGSLVNLPIVNYYLHSTYITGCHAH
GAMFGVKGNVALAGVLFCVQHLVKEKHWSGRLLNISFWCLNGGMALMMFISLFPTGLYQL
FMVIKYGFWYARSAEIVQGPVFQFFLKLRPIGGHIFFEGGLLPLVYFIFSRWYTVK-NET
D----PKSKEENMYRS--------T---WHIDESDTKK----------------------
--VQ
>MBS93797_Chromatiales_NP37
M-----------------------------------------------------------
-------STSSQQNLALWLVNKKNWLTHFMIVAGISIAGLIYL--GGATYTGAPPLVDYV
--S-SSGDTVISQQQIKRGKELFHIRGLMSWGSFWGDGAERGPDFTADALHRTVMGMRAF
YGQDLEQRAG-----------S---LTQHDQDAISVRVKREVH----NNAYD-EEAGVIR
LNDAQVSALGDLNAHYTRMFNDPAYS-----------------EAY---EPS---GLISD
PDDLRDLTAFFFWGGWVSAADRPGETYSYTHNWPGDKDAGNEPTAATFFWSVFSVFALFV
GVMIVLYAYGEMKEQPVDVFDQSNESQSGGQSLTTYDLEN---EYVRPTQRATYKFFVLA
IIVFGCQILGGVIAATDFV---RPG------GVS-LNEIIPFNVARSYHTLLQIFWFFMC
WVGYTIFFLPRISK-VPPGQKSLINLLFFLCVVTGGGALVGIYMGQTGML-SDNMSYWFG
SQGWEFMELGRFFQFTLLAAFALWIWIIYRAVKPWLTRRN-VWSVPSWLLYGSGIMVAFL
FFGLLVQPEMNFAISDYWRWMVVHMWVEVTFEVFTTVIVGYMLVQMGLITRMMAERVIFL
AVMLFLVTATIGISHNFYWIAKPTGIIALGSVFSTLQVLPLLLLTLDAWQMRQEGL----
KADKNVVEGKQTIVMEGVWLFILGVNFWNIFGAGVFGSMINLPIVNYYEHATYLTGNHAH
AAMFGVKGNIALGGMLFCCQHLFHKSAWNPKLVRTSFWSMNIGLVLMMFLDLFPVGLYQT
WVAYTDGTWYARSSEIVTGPVFSFLTYCRTIGGAVFIWGGLLPMMWFILSRGTKLRRKEE
E----VEQGEWTVYEA--------D---WADQKDPALGG---------------------
----
>MBT76598_Chromatiales_NP153**
M-----------------------------------------------------------
-------SNANQQNLALWLVNKKNWLTHFLIVATICIAGLIYL--GGATYTGAPPLVDYV
--S-ATGETVISQQQIKRGQELFHIRGLMSYGSFWGDGAERGPDFTADALHRTVIGMRGF
YAQEKENSGV-----------V---LDKHDHGAIASRVKLEVH----ENTYD-ENAGIIH
INDAQIQAFNDLNDHYTQMFTNPDYTQCYRDCANPS------NEAF---LPS---GHISD
```

```
PDDLRALSAFFFWGGWVASTNRPGENYSYTHNWPGDKDAGNSPTAATFFWSVFSIFALFV
GVMVVLYAYGQMKEQPVDVFDSST--RNGGQSLTTYDLEN---EYVRPTQRATYKFFVLA
VIVFGVQIIGGVIAATDFI---RPG------GVS-LNDIIPFQVARSYHTLLQIFWFFMC
WVGYTIFFLPRISK-VPPGQKALINLLFVLCVVTGGGALVGIYMGQTGML-SDSMSYWFG
SQGWEFMELGRFFQFTLLAAFALWIGIIYRAVKPWLTRRN-LWSVPSWLLYGSGIMVMFL
FFGLLVQPEMNFAISDYWRWMVVHMWVEVTFEVFTTVIVGYMLVQMGLITRMMAERVIFL
AVMLFLITATIGISHNFYWIAKPTGIIALGSVFSTLQVPLLLLTLDAWQMRQEGL-----
TANKNVVEGKQTIVMEGVWLFILGVNFWNIFGAGVFGSVVNLPIVNYYEHATYLTGNHAH
AAMFGVKGNIALGGMLFCCQHLFHKSAWNPKLVRTSFWSLNIGLVLMMFLDLFPVGLYQT
WIAYTEGTWYARSSEIVTGPVFSFLTYCRTIGGAVFIWGGLLPLMWFILSRGFKLRRKEE
E----VEQGEWTVYES--------E---WAEQNNPSLGG---------------------
----
>Ga0187827_100036806
M-----------------------------------------------------------
-----STTSTSNQNIALWLVNKKNWLTHFMIVSAICIAGLIYL--GGATYMGSPPLEDYV
--SASTGETVITRTQIKRGQELFHIRGLMGYGSFWGDGAERGPDFSADALHRTVMGMRSF
YAQELADSGV-----------A---PTEYDAGAIAARVRGEVH----ENYYD-EVAGVIR
INDAQISAFSDVVDHYTAMFNDPTYS-----------------EAF---EPS---GFISD
PDDLEALSAFFFWGGWVSAANRPGETYSYTHNWPNDPDAGNSPTAATFFWSVFSIFALFV
GVMLVLYVYGQMREQDVDVFDTTSGNENNGQSLTTYDLEN---EYVRPTQRATYKFFVLA
MIVFLVQIIGGVLAATDFV---RPF------GVS-MNDIIPFTVARSYHTLLQIFWFFMC
WVGYTIFFLPRISK-VPPGQKTLINLLFVLCIVTGGGALAGIYMGQTGML-SDSAAYWFG
SQGWEFMELGRFFQFTLLAAFALWIFIIYRAVRPWLTKRN-LWSVPSWLLYGSGIMVMFL
FFGLMVQPDMNFAISDYWRWMVVHMWVEVTFEVFTTVIVGYMLVQMGLITRMMAERVIFL
AVMLFLITATIGISHNFYWIAKPTGIIALGSVFSTLQVPLLLLTLDAWQMRQEGH-----
TATQNMMEGKQTIVMEGVWLFILGVNFWNIFGAGVLGSVINLPIVNYYEHATYLTGNHAH
AAMFGVKGNIALGGMLFCCQHLFHKAAWNPKLVRTSFWSLNIGLALMMFLDLFPVGLYQT
WIAYTEGTWYARSSEIVTGPVFSFLTYCRTIGGMVFIWGGLVPMVWFILSRGTKLRLKEE
D----VAEGEWTVYEQ--------A---WSEQNDPSLGG---------------------
----
>Ga0187827_100071743*
M-----------------------------------------------------------
-------STTSNQNIALWLVNKKNWLTHFMIVSAICIAGLIYL--GGATYMGSPPLEDYV
--SASTGETVISRTQIKRGQELFHIRGLMGWGSFWGDGAERGPDFSADALHRTVLGMRSF
YAQELENSGV-----------S---PTEYDAGAIASRVRDEVH----VNNYD-EVAGVIR
INDAQISAFADLNEHYTAVFNDPTYS-----------------EAF---EPS---GFISD
PDDLTALTAFFFWGGWVSAANRPGENYSYTHNWPNDPDAGNSPTAATFFWSVFSIFALFV
GVMLVLYVYGQMREQDVDVFDTSAGNENNGQSLTTYDLEN---EYVRPTQRATYKFFVLA
MVVFLVQIIGGVVAATDFV---RPG------GVS-LNEIIPFTVARSYHTLLQIFWFFMC
WVGYTIFFLPRISK-VPPGQKSLINALFVLCVVTGGGALAGIYMGQTGML-SDGAAYWFG
SQGWEFVELGRFFQFTLLAAFALWIFIIYRAVKPWLTKRN-LWSVPSWLLYGSGIMVMFL
FFGLMVQPDMNFAISDYWRWMVVHMWVEVTFEVFTTVIVGYMLVQMGLITRMMAERVIFL
AVMLFLITATIGISHNFYWIAKPTGIIALGSVFSTLQVPLLLLTLDAWQMRQEGH-----
TATKNMIEGKQSIVMEGVWLFILGVNFWNIFGAGVLGSVINLPIVNYYEHATYLTGNHAH
AAMFGVKGNIALGGMLFCCQHLFHKAAWNPKLVRTSFWSLNIGLALMMFLDLFPVGLYQT
WVAYTDGTWYARSSEIVTGPVFTFLTYCRTIGGMVFIWGGLVPMVWFILSRGTKLRLKEE
D----VAEGEWTVYEN--------A---WSDQKDPSLGG---------------------
----
```

```
>RLA57063_Gammaproteobacterium_B36_G6
M-----------------------------------------------------------
----------SSQNLGTVLTSRKNWFGVFLLVAAISVAGLIYL--GGATYTGAPPLENYE
--S-STGDVIISRDQIHKGETVFHLRGLMSWGSFWGDGGERGPDFTADALHRTAMSMRRF
YELEVEANSGR----------P---ATQYDKDAISVRVVRELH----NNAYD-EDRGLIE
INDAQIFALKELNEHYTSMFMDPSYS-----------------EAF---DPA---GYISD
PTDIKNLTAFFFWGGWVSAANRPGENYSYTHNWPGDKAAGNAPTSATFMWSVFSVFALIL
GISAVLYVYGEMKDQDQDMFEAEE-----GTTLTTYDLEN---EYVRPTQKATYKFFVLA
IVVFAVQIIAGTIIATDFV---RPG------GMS-LNEIIPFPIARSYHTLLQIFWFFMC
WVGYTIFFLPRISK-VPTGQKLLIDVLFVLCLITGVGSIVGIYLGQSGVL-TGDAAYWLG
SQGWEFMEMGRFFQLTLLSSFALWIFIIYRAVKPWLTAKN-MWSVPSWLLYGSGIMVFFL
FFGLFIQPEMNFAIADYWRWMVVHMWVEVTFEVFTTVIIGYMLVQMGMITRVMAERVIFL
AVMLFLITATIGISHNFYWIAKPTGIIALGSVFSTLQVLPLLLLTLDAWQMREEGH----
WANKNRMAGKQNHVMEGVWLFILGVNFWNIFGAGVLGSVINLPIVNYYEHATYLTGNHAH
AAMFGVKGNIALGGMLFCCQHLFHKEAWNAKLVKTAFWSLNIGLAMMMFLDMFWVGLYQT
WVAFTDGTWYARSQAIATGNVFVYLTSARALGGLVFIGGGLLPLVWFILSRGSKLRLKEE
D----VEEGEWTVYEQ--------D---WSEQNDPALGG---------------------
----
>MBT4522754_Halieaceae_SI047_bin19
M-----------------------------------------------------------
----------SSQNLAITLMNRKNWFTIFLIVSAISIAGLIYL--GGATYTGAPPLENFE
--S-STGNVVISRDQIKKGEEVFHLRGLMSWGSFWGDGAERGPDFTADALHRTAMSMRRF
YEQEVADRTGQ----------A---ATQYDKDAISVQVIRELH----TNTYD-EDRGLIQ
INDAQIFALKELNEHYTSMFTDADYY-----------------EAF---DPA---GYISD
PDDIAKLTAFFFWGGWVAAANRPGESYSYTHNWPGDKDAGNFPTSATFMWSVFSIFALIL
GVCIVLYVYGQMKEEEQDMFEAEE-----GATLTTYDLEK---EYVRPTQKATYKFFMLA
ILVFGVQILAGIIIATDFI---RPG------GVS-LNEILPFPVARSLHTLLQIYWFFMC
WVGYTIFFLPRISK-VPNGQRALINILFTLCLITGVGAIVGIFLGQTGII-TGQMAYWFG
SQGWEFMELGRFFQFTLLASFALWILIIYRGVKPWLTMKN-MWSVPSWLLYGSGIMVFFL
FFGLLVQPDMNFAISDYWRWMVVHMWVEVTFEVFTTVIIGYMLVQMGMITRRMAERVIFL
AVMLFLITATIGISHNFYWIAKPTGIIALGSVFSTLQVLPLLLLTLDAWKMREEGR----
WADKNRIAGKQTHVMEGVWLFILGVNFWNIFGAGVFGSLINLPIVNYYEHATYLTGNHAH
AAMFGVKGNIALGGMLFCCQHLFQKPAWNAKLVRTAFWSLNIGLAMMMFLDLFWVGLYQT
WIAYTHGTWFARSQEIVTGDVFVYMTYARSLGGFVFIGGGLIPLIWFIMSRATKLRQGEE
D----VQEGEWTVYDK--------D---WGEQNNPALGG---------------------
----
>MBP20251_Gammaproteobacterium_NP964*********
M-----------------------------------------------------------
-----------SQNLAITLMNKKNWAALFALVSAISIAGLIYL--GGATYTGSPPLENFK
--A-ANGDIVISREAIKGGEEVFHLRGLMSWGSFWGDGGERGPDFTADALHRTAMSMREF
YAQELGGSTSQYD--------T---LPQYDKDAIAARVIRELH----ANTYD-EESGFIG
INDAEIYAFEELNKHYTRMFTDASYE-----------------EAF---SPA---GYISE
PDQLRDLTAFFFWGGWVSSSNRPGENYSYTHNWPGDKDAGNTPTDATFMWSVFSIFALIF
GIGAVLYVYGQMKEQPVDVFEGGDDNGNGQQSLTTYDLEN---EYVRPTQRATYKFFMLA
IIVFGVQILAGIIIATDFV---RPF------GMD-LNELIPFTVARSYHTLLQIYWFFMC
WVGYTIFFLPRISK-VPPGQKGLIDLLFALCVITGAGAIFGIYLGQTGVI-TGSMAYWFG
SQGWEFMELGRFFQFTLLTSFALWILIIYRGVKPWLTTKN-MWSVPSWLLYGSGIMVAFL
FFGLMIQPEMNFAISDYWRWMVVHMWVEVTFEVFTTVIIGYMLVQMGLITRLMAERVIYL
```

```
AVMLFLITATIGISHNFYWIAKPTGIIALGSVFSTLQVLPLLLLTLDAWQMRQEGN----
SAEKNRVEGKQTFVMEGVWLFILGVNFWNIFGAGVFGSLINLPIVNYYEHATYLTGNHAH
AAMFGVKGNIALGGMLFCCQHLFTRTAWNAKLVKTAFWSMNIGLAMMMFMDLFWVGLYQT
YVAFAEGTWMARSQEIVTGPVFVYLTSARALGGLVFIWGGLLPMMWFILSRGNKLRLKEE
D----VEEGEWTVYDK--------D---WSAQTDPSRGG---------------------
----
>MBV28360_Rhodospirillaceae_NP1106****
M-----------------------------------------------------------
--SKESNGNSIHQNLALILLNKNYWITHFMIVVVISVVGLIYL--GRVTYTGAPPLVDYV
--S-SAGDTVISREEIKRGEEVFHLRGLMSYGSFWGDGAERGPDFSADALHRTVVTMRSF
YENEAKEKG------------A---VSQYDQDAIAAQVKREVH----HNTWE-EDADLIR
INDAQIYAFEELNVHYTKMFSDPDYP-----------------EHF---M-T---GYITD
SEDIRALTAFFYWGGWVAAANRPGETYSYTHNWPYDPDAGNTPTSANYIWSIISILALFL
GIGVVLYVYGQMKSLGGEPFSGN------GTSLTTHDLEN---EYVRPTQRATYKFFALA
VILFGIQVLAGIISATDFV---RPY------GLY-LGDIIPFTVARSYHTLFQIFWFFMC
WVGYTIFFLPRLAK-VPNGQAFLINLLFALCIIVGGGALVGIYLGQTGVL-TGPAAYWFG
SQGWEFMELGRAFQIILLMAFALWIAIIYRGVKPWLTKKN-LWSVPAWLLYGSGIMVAFL
FFGLLVTPDTNFAIADYWRWMVVHMWVEVTFEVFTTVIVAYMLVQMGLVTRMMAERVIFL
AVILFLITATLGISHNFYWIAKPTGIIAIGSVFSTLQVLPLLLLTLDAWKMRQEGE----
RANEYVAQGKQKFLMDGVWMFILAVNFWNIFGAGVFGSLINLPIVNYFEHATYLTGNHAH
AAMFGVKGNIALGGMLFCCMHLFHKASWNPKLVMTSFWSLNIGLALMMFLDLFPVGVYQL
VAVLQEGLWYARSYEIVQGVVFQTLTYFRSLGGAVFVIGGLLPLIWFVLSRGSQLR-REV
E----VKDGEWSVYEK--------H---WATQEDSKLLG---------------------
----
>MBL6933931_Alphaproteobacteria_S150m-G7_BlackSea
M-----------------------------------------------------------
----KSNGNSIHTNLALILLNKNYWIWHFLVVVVIGVVGLIYL--GRATYTGAPPLVDYV
--S-SAGDTVISREEIKKGEEVFHLRGLMSYGSFWGDGAERGPDFTADALHRTVVAMKSF
YEDARSKETSR----------T---VSQHDKDAIAARVKREVR----KNTWD-EDANLIQ
INPAQAHAFEELNDHYTKMFSDPEYP-----------------EFF---M-S---GYITD
PEDIRNLTAFFYWGGWVAAANRPGETYSYTHNWPYDPDAGNTPTSATNIWSIISILALFL
GIGVILYVYGQMKTLQGDPFASN------GTSLTTHDLEN---EYVRPTQRATYKFFALA
VILFGLQVLAGIISATDFV---RPF------GLY-LGDIIPFTVARSYHTLFQIFWFFMC
WVGYTIFFLPRLAK-VPSGQAFLINLLFALCMVVGAGSLVGIYLGQTGIL-TGPAAYWFG
SQGWEFMELGRAFQIILLVAFALWILIIYRGVKPWLTKKN-LWSVPAWLLYGSGIMVAFL
FFGLLVTPETNFAIADYWRWMVVHMWVEVTFEVFTTVIVAYMLVQMGLVTRMMAERVIFL
AVMLFLITATLGISHNFYWIAKPTGIIAIGSVFSTLQVLPLLLLTLDAWQMRQEGE----
RANEYVAQGKQRFLMDGVWLFILAVNFWNIFGAGVFGSLINLPIVNYFEHGTYLTGNHAH
AAMFGVKGNIALGGMLFCCMHLFHRASWNPKLIKTAFWSLNIGVALMMFLDLFPVGVYQL
VAVLQEGYWYARSHEIVQGVVFQTLTYFRSIGGAVFVIGGLLPLIWFILSRGFQLR-REV
E----VKDGEWSVYEK--------N---WATQEDPKLLG---------------------
----
>MBT5374662_Rhodospirillaceae_SI074_bin107
M-----------------------------------------------------------
--STQSYKSSNRGNLALILLNKNYWIWHFLIVVVIGVVGLIYL--GRLTYTGAPPLVDFT
--S-STGETVISLQEIKHGQEVFHLRGLMSYGSFWGDGAERGPDFTADALHRTVVAMKSF
YENEAKEGG------------E---LSQYDEDAIAAQVKREVH----TNTWD-EDAGIIR
INAAQIYAIGELNDHYTRMFSDPDYP-----------------EFF---M-S---GYITD
```

```
PADIKGLTAFFYWGGWVAAANRPGEIYSYTHNWPYDPDAGNTPTSATYIWSIISILALFL
GIGVVLYVYGQMKTLQGDPFVSN------GTSLTTHDLEN---EYVRPTQRATYKFFALA
VILFGLQVLAGVISATDFI---RPF------GLY-MGDIIPFTVARSYHTLFQIFWFFMC
WVGYTIFFLPRLSK-VPRGQKFLINLLFMLCIIVGGGSLVGIYLGQTGIL-TGEAAYWFG
SQGWEFMELGRLFQIILLVAFTLWILIIYRGVKPWLTKKN-LWSVPAWLLYGSGIMVGFL
FFGLLVSPDTNFAVADYWRWMVVHMWVEVTFEVFTTVIVAYMLVQMGLVTRLMAERVIFL
AVMLFLLTATLGISHNFYWIAKPTGIIAIGSVFSTLQVLPLLLLTLDAWQMRQEGG----
RASEYQSQGKQTFVMEGVWLFILAVNFWNIFGAGVFGSLINLPIVNYFEHATYLTGNHAH
AAMFGVKGNIALGGMLFCCMHLFKESSWNPKLIKIAFWSLNIGVAMMMFLDLFPVGVYQL
MLVFQEGFWYARSQEVVQGTVFQTLTYFRSLGGAVFIIGGLLPIIWFILSRGRHLR-REV
N----VEDGEWSVYQK--------N---WAEQEDEKLPG---------------------
----
>Ga0066848_100037855*****
M-----------------------------------------------------------
---SASAETNGRGDYALVLINKKYWVGLFLIVAAISVSGLIYL--GRVTYTGAPPDANFV
--S-STGEVVLSRQLVARGEEVFHLRGLMGYGSFWGDGAERGPDFSAEALHRSVVGMRTF
YENEIKKDR------------A---VTQFDRDAIGARVKREVH----VNTWN-EEAGTVT
LNDAQVFAIQELNVHYTRMFMDPEYK-----------------HLF---Q-T---GYITD
PEDIRALTAFFYWGGWVAAANRPGEVYSYTHNWPYDPAAGNYPTSAVYIWSILSIFALFL
GIGAVLYVYGQMKDLGGEPFDGT------GTSLTTSDLEN---EYVRPTQRATYKFFALA
MILFGLQILAGIIGASDFI---RPF------GIF-LGDLVPFTVARSYHTLFQIFWFFMC
WVGYTIFFLPRLSK-VPPGQKFLINLLFVGCLIVGAGALVGIYLGQTGVL-TGSTAYWFG
SQGWEFMELGRAFQFILLGGFALWIFIIYRGVRTWLTKRN-MWSVPAWLLYGSGIMVGFL
FFGIFVSPETNFAIADYWRWMVVHMWVEVTFEVFTTVIVAYMLVQMGLVTRLMAERIIFL
AVMLFLITATIGISHNFYWIAKPTGIIALGSVFSTLQVLPLLLLTLDAWRMRQEGD----
RAQEYRSQGKQMFVMEGAWMFILAVNFWNIFGAGVFGSLVNLPIVNYFEHATYMTGNHAH
AAMFGVKGNIALAGMLFCCQHLFHKSSWNAKLVRTSFWSLNIGLGLMMFLDLFPVGVYQV
VIVLQEGLWYARSVDVVRGSVFVTLTYFRMIGGGVFVLGGLLPLIWFILSRGFSLV-REV
E----VEEGEWEVYEK--------D---WAAQEDPSLRHSRPGLAE--------------
----
>Ga0187827_100051115*
M-----------------------------------------------------------
---SVSAENNHGEDFALILINKKYWMGLFLIVAAISISGLIYL--GRATYTGAPPDANFI
--S-SSGKTVLSRQLVSRGEEVFHLRGLMSYGSFWGDGAERGPDFTADALHRTVVAMRTF
YENELKKER------------A---VTPYDKEAIAARVRLEVH----NNTWD-EKSGTIT
LNDAQIFAFDEVHKHYTSMFMDPNYP-----------------ELF---Q-T---GYITD
PENIRALSAFFFWGGWVSAANRPGENYSYTHNWPYDPDAGNHPTSATYIWSFLSIFALFL
GIGAVLYVYGQMKVLGGEPFVEN------GTSLTTSDLEN---EYVRPTQRATYKFFALA
MILFGLQVLAGIISATDFI---RPF------GIF-LGDIIPFTVARSYHALFQIFWFFMC
WVGYTIFFLPRLSK-VPSGQKFLINLLFLLCVITGGGALVGIYLGQTGALGTGEMAYWFG
SQGWEFMELGRAFQFVLLVAFALWIYIIYRGVKTWLTKRN-MWSVPAWLLYGSGIMVGFL
FFGILVSPETNFAIADYWRWMVVHMWVEVTFEVFTTVIVGYMLVQMGLVTRLMAERIIFL
AVMLFLITATIGISHNFYWIAKPTGIIALGSVFSTLQVLPLLLLTLDAWRMRQEGD----
RAEEHRAQGKQMFVMEGVWMFILAVNFWNIFGAGVFGSLINLPIVNYFEHATYMTGNHAH
AAMFGVKGNIALAGMLFCCQHLFHKNSWNPKLVRTSFWSLNIGIAMMMFLDLFPVGVYQM
VIVLQEGLWYARSSEIVMGTVWKTLTYFRMIGGGVFVLGGLLPLIWFILSRGMRLV-REV
E----VEEGEWTVYEK--------D---WAAQEDPTLRHSKPGLAE--------------
----
```

```
>OEJ69613_Magnetovibrio_blakemorei_MV-1
M-----------------------------------------------------------
--SSKLNDKLRFKNLSLILLNKKFWFTHFLIVTIISVAGLIYL--GQATYSGAPPLVHFK
--S-STGETVISRQLINAGERVFHLRGLMSYGSFWGDGAERGPDFTADALHRMTVSMRSF
YAGELKAQGTTQ---------D---LGQYDQDAIAARVRREVH----TNTWD-ESSDTIL
LNEAQIFALEELNRHYTRMFTDPHYP-----------------ELF---Q-I---GYITE
PKDIQALTAFFYWGAWVAAANRPGEDYSYTHNWPYDPDAGNTPTTATFIWSAMSILGLFL
GIGIVLYVYGQMKSMHYDPFNGNT-----GTSLTTHDLEN---NYIRPTQRGTYKFFALA
IVLFALQVLAGIIGATDFV---RPF------GLF-LGDIIPFTVARSYHTLFQIFWFFMC
WVGYTIFFLPRLSE-APRGQTFLINLLFTLCIIVGAGALVGIYLGQTGIL-TGDMAYWLG
SQGWEFMEFGRLFQIILLVAFTLWIFIIYRGVKPWLTRKN-MWSIPAWLLYGSGVMVFFL
FFGLLVTPETNFAVADYWRWMVVHMWVEVTFEVFTTVIVAYILVQMNLITRAMAERVVFL
AVMLFFITATVGIAHNFYWIAKPTGVIAFGSVFSTLQVPLLLLTLDAWRMRQEKT-----
RANTLRDQGKQTIVMDEVWLFILAVNFWNVFGAGVFGSVINLPIVNYYEHATYLTGNHAH
AAMFGVKGNIALAGLLFCCQHIFEKSSWNAKLIRRSFWSLNIGVAMMMFLDLFPVGVYQL
VDVLQNGLWHARSYEIVQGDVFKTLTYFRSLGGAVFVIGGVIPLVWFILSRGRRLV-REV
Q----IEPNSHTDYEK--------D---WAAQEDAP----------------------H-
----
>MBT38371_Deltaproteobacterium_NP36******
M-----------------------------------------------------------
--------SNSSRTIAQILQDKKTWWLHFLIVAAICVTGLVYL--GTETYTGAPPLEDYV
--S-ASGETVISREQIKHGQEVFHLRGLMSYGSFWGDGAERGPDFTADALHRTVMSMRAF
YEKEMGGA-----------------TTVYDKDAVAARVRREVH----TNTWD-ERAGHIV
ISPAQTQGYNDLIVHYTAMFNDPNYN-----------------EAF---EPA---GYISD
PEDLRDLSAFFFWGGWVSAANRPGEEYSYTHNWPYDPDAGNNATSQTWIWSFISIFGLFL
GILVVLYVYGQFKEQ-GDPFIGN------GTSLTTNDLEE---GRVRATQRATYKFFVFS
ILLFGAQVLAGIFASMDFV---RPF------GFS-LGDIIPFTVLRSYHTLFQIYWFFMA
WVGYTIFFLPRISK-VPNGQIGLINGLFALCIVTGVGALVGIYAGQTGMI-TGATAYWFG
SQGWEFMEIGRAFQYTLLGSFALWIYIIYRGVKPWLTTKN-IWSVPSWLLYGSGIMVAFL
FFGLMISPEQNFAVSDYWRWMVVHMWVEVTFEVFTTVIVGYILVQMGLISRMMCERVIFL
AVMMFLVTATLGISHNFYWIAKPTGIIAVGGVFSTLQVPLLLLTLDAWRGKSEAG-----
RARTHVTEGRQVFMMEGVWLFMLAVNFWNIFGAGVFGSLINLPIVNYYEHATYMTGNHAH
AAMFGVKGNIALGGMLFCIQHLVHKEAWSPKLVRTAFWSLQIGVALMMFLDLFPVGLYQI
MIVVQDGLWFARSQEILTGPVWTLLTYFRSLGGTLFLLGGVLPLIYFIVTRGSELK-EEA
D----SDEGEWTVYEK--------D---WAA-----------------------------
----
>MBW2395684_Deltaproteobacterium_Meg19_02_Bin_150
M-----------------------------------------------------------
--------SSSGRNLAQVLQDKNTWWVHFLIVTAICATGLIYL--GTQTYSGAPPLESYK
--S-ADGATVISREQIKHGQEVFHLRGLMSYGSFWGDGAERGPDFTADALHRTVVTMRTF
YANELGGV-----------------VSQYDEDAIAARVRRELH----TNTWD-EEAGHIV
INAAQIQAYEDLVVHYTAMFNDPGYG-----------------EAF---EPA---GYISD
PTDLRDLTSFFFWGAWVSAANRPGQVYSYTHNWPYDPDAGNTATSQTWIWSFISIFGLFL
GILAILYVYGQFKEQ-GDPFEGD------GTSLTTNDLEE---GRVRATQRATYKFFIFS
IVLFGCQVLAGVFASMDFV---RPF------GFS-LGDIIPFTVLRSYHTLFQIYWFFMA
WVGYTIFFLPRISK-VPNGQLFLINLLFAICCVTGAGALVGIYAGQTGMI-TGATAYWFG
SQGWEFMEIGRAFQYTLLTSFALWIYIIYRGVRPWLTAKN-IWSVPSWLLYGSGIMVGFL
FFGLMISPEQNFAVSDYWRWMVVHMWVEVTFEVFTTVIVGYVLVQMGLISRMMAERVIFL
```

```
AVMMFLVTATLGISHNFYWIAKPTGIIAVGGVFSTLQVLPLLLLTLDAWRMKSEAG----
RAQTNVVQGRQIFVMEGVWLFILAVNFWNIFGAGVFGSLINLPIVNYYEHATYMTGNHAH
AAMFGVKGNIALGGMLFCIQHLIHKEAWNEQLVRRAFWSLQGGLGLMMFLDLFPVGLYQC
MIVVQEGLWFARSQEILTGTVWKTLTYFRSIGGTLFIVGGVLPLIYFFITSARQLK-PEA
D----SDEGEWTVYEK--------D---WAA-----------------------------
----
>MBW2230945_Deltaproteobacterium_Meg19_46_Bin_91
M-----------------------------------------------------------
--------SNPSRNLAQVLQDKRTWWVHFLIVAAICATGLIYL--GTETYSGAPPLESFV
--SSSTGKVVFSREQIKKGQEVFHLRGLMGYGSFWGDGAERGPDFTADALHRTVVTMRDF
YADEIAATQ------------A---VTQYDTDAIAARVRREIH----TNTWD-EKAEQIL
INDAQVQAYRDLVVHYTRMFNDSEYG-----------------EAF---EPA---GYISD
PQDLENLTAFFYWGAWVSGANRPGQVYSYTHNWPYDPDAGNTATSQTWIWSFISILGLFL
GILAVLYVYGQFKEQ-GDPFEGN------GTSLTTNDLEE---GRVRATQRATYKFFVFS
IVLFGCQVLAGMFASMDFV---RPF------GFS-LGDIIPFTVLRSYHTLFQIYWFFMA
WVGYTIFFLPRISR-VPNGQLFLINALFAICMVTGAGALVGIYAGQTGMI-TGQTAYWFG
SQGWEFMEIGRFFQYTLLSSFALWIYIIFRAVRPWLTAKN-IWSVPSWLLYGSGIMVAFL
FFGLMISPEQNFAVSDYWRWMVVHMWVEVTFEVFTTVIVGYILVQMGLISRVMCERVIFL
AVMMFLVTATLGISHNFYWIAKPTGIIAVGGVFSTLQVLPLLLLTLDAWRMKSEAG----
RANTNLVAGRQIHVMEGVWLFILAVNFWNIFGAGVFGSLINLPIVNYYEHATYMTGNHAH
AAMFGVKGNIALGGMLFCVQHLISKEAWNEKLVRTSFWSLNLGMALMMFLDLFPVGLYQC
MIVVQEGLWYARSQEILTGTVWKTLTYFRSIGGTLFIVGGVIPLIWFFITSATRLE-AEA
D----SDEGEWTVYEK--------D---WAA-----------------------------
----
>MAG34007_Deltaproteobacterium_ARS120
M-----------------------------------------------------------
--------SDSNRNLAQILQDKKTWWVHFLIVAAICVSGLLYL--GQQTYSGAPPLESFV
--T-ADGQTVFSREQIKQGQEVFHLRGLMGYGSFWGDGAERGPDFTADSLHRTVLSMRAF
YEEELGANQS-----------E---LSLYDKDAVAARVRREVH----NNTWD-EQAGQIV
INDAQVRAYNDLVKHYTAMFNDPDYA-----------------EAF---EPT---GFISD
PEELRSLTAFFYWGGWVSAANRPGQVYSYTHNWPYDPDAGNTATSQTWIWSFISIFGLFL
GILAVLYIYGQFKEQ-GDPFVGN------GTSLTTNDLEE---GRVRATQRATYKYFIFS
IVLFGCQVVAGIFASMDFV---RPF------GFS-LGEIIPFSVLRSYHTLFQIYWFFMA
WVGYTIFFLPRISR-VPTGQIALINGLFALCMVTGVGALVGIYAGQTGMI-TGSTAYWFG
SQGWEFMEIGRFFQYTLLTSFALWIYIIFRAVKPWLTAKN-IWSVPAWLLYGSGIMVAFL
FFGLMISPEQNFAVSDYWRWMVVHMWVEVTFEVFTTVIVGYILVQMGLISRMMCERVIFL
AVMMFLVTATLGISHNFYWIAKPTGIIAVGGVFSTLQVLPLLLLTLDAWRGKSEGG----
RAEKNVLAGRQAFLMEGVWLFMLAVNFWNIFGAGVFGSLINLPIVNYYEHATYMTGNHAH
AAMFGVKGNIALGGMLFCIQHLIHKEAWNEQLVKRAFWSLQIGVALMMFLDLFPVGLYQC
MIVVQEGLWFARSQEILTGSVWKTLTYFRSIGGTLFLVGGVLPLIWFILKSATQMK-AEA
D----SDEGEWTVYEK--------D---WAA-----------------------------
----
>MBM4335308_Deltaproteobacterium_K_DeepCast_100m_m2_268
M-----------------------------------------------------------
--------SNTKRNLAQVLHDKKTWWVHFLIVAAICLSGLVYL--GTETYTGAPPIEDYV
--SATTGEVVIPLATIKKGQEVFHLRGLMLYGSFWGDGAERGPDFTADALHRTGVAMRSF
YEQQGGGN-----------------LAQYDKDAIDVRVIREIK----TNTWN-ESAKQIV
INDAQIHAFNELKAHYTRMFNDPEYE-----------------EAF---KPS---GAISA
```

```
PEDIEALTAFFFWGGWVSGAERPGQTYSYTHNWPYDPSTGNLPTNQTMVWSFISIFGLFL
GILVILYVYGQFKEE-GDPFTGN------GTSGTTNDLEQ---GRIRATQKATYKFFVFS
IILFGAQVLSGMLAATDFV---TPF------GVS-LASIIPFSVLRSYHTLFQIYWFFMA
WVGYTIFFLPRISK-VPNGQLFLINLLFAICMVTGVGALVGIYMGQTGMI-TGATAYWFG
SQGWEFMELGRAFQYTLLSSFALWIYIIYRSVRPWLTTKN-IWSVPSWLLYGSGIMVAFL
FFGLMISPDQNWAVADYWRWMVVHMWVEVTFEVFTTVIVGYMLVQMGLISRMMCERVIFL
AVMMFLVTATLGISHNFYWIAKPTGIIAVGGVFSTLQVLPLLLLTLDAWRMKSEAV----
RAKTHVAAGRQVFVMEGVWLFILAVNFWNIFGAGVFGSLINLPIVNYYEHATYMTGNHAH
AAMFGVKGNIALGGMLFCAQHLLTKADWSEKLVKNSFWSLQIGLLLMMTLDLFPVGLYQL
MIVFQEGLWYARSQEVITGPVWTFLTYMRSIGGTLFIVGGVLPLIWLFVSRVGKLR-KET
PEL--ESEGEWTVYEK--------D---WAA-----------------------------
----
>QOJ32248_Gammaproteobacterium_H2_PRO1
M-----------------------------------------------------------
-STKAANVSTGDSNLAMWLLNKKNWFTHFLIVAIISIAGLVYL--GRETYVGAPPLANFV
--T-SNGQVLMTTEQLARGKEIFHLRGLMLYGSFWGDGAERGPDFTADAMHRIGASMRSY
YEEETKRRTGAE---------S---LTGYEKDAIAQRVLREMH----NNTYN-AEAGTIV
LNDAQVFAFDELLQHYTRMFTDPTYP-----------------DRM---DPI---NQVSG
EQNLRDLTTFFYWGGWVSAANRPGEEYSYTHNWPYEPEVGNVPTTATFIWTFISIFALWI
GISVVLYVYGQMKEQPVDVFSQEG---ANGHSLTTSDLEN---GYVRETQRSTYKFFALA
MIVFGLQVLGGVVSAWDFI---RPF------GIN-LNELLPFTVSRSYHAILQIYWFFMC
WVGYTIFFLPRLTK-VPKGQKFLINLLFFLSCVVAVGAVGGIYAGQRGWI-DDTMSYWFG
SQGWEFIELGRFFQWVLLLGFSLWIYIIYRGVKPWISVKN-VWSVPAWLLWGSGVMVLFL
FFSVLMTPDSNFAISDYWRWMTVHMWVEVTFEVFTTVIVAYLLVQMGLVTRMMAERVIFL
AVMLFFVTAINGISHNFYWIAKPTGIIAVGSVFSTLQVLPLLLLTLDAWQMRQEGG----
RAHEFRVQGKQVVVMEGVWLFILGVNFWNIFGAGVFGSLITLPIVNYYEHATYMTGNHAH
AAMFGVKGNVALAGMLFCLQHLFQKTAWNEKLVRTVFWSFQIGLALMMFLDLFPVGLYQI
YVVLTEGLWYARSTEVIMGPVFATLTYLRTIGGAVFIFGGLLPIIYFVLSGASRLR-KEV
D----VTTDEWAQYHKEYGNEGGKE---WAAQEEPKVSR---------------------
----
>MBC6944429_Gammaproteobacterium_PRO3
M-----------------------------------------------------------
-STNAANVSTGDSNLAMWLLNKKNWFTHFLIVAIISIAGLVYL--GRETYVGSPPLANFV
--T-SSGQVVMTTEQLARGKEIFHLRGLMLYGSFWGDGAERGPDFTADSMHRIGASMRSY
YEEETKRRTGAQ---------S---LTEYEKDAIAQRVLREMH----GNTYS-AEAGTIT
LNDAQVFAYDELLTHYTRMFTDATYP-----------------DHM---DPI---NQVSG
EQNLRDLTAFFYWGGWVSAANRPGEEYSYTHNWPYEPEVGNVPTTATFIWTFISIFALWV
GISVVLYVYGQMKEQPVDVFSQEG---ANGHSLTTSDLEN---GYVRETQRSTYKFFALA
MIVFGLQVLGGVVSAWDFI---RPF------GIN-LNELLPFTVSRSYHAILQIYWFFMC
WVGYTIFFLPRLTK-VPKGQNFLINLLFVMSVIVALGCVFGIYAGQRGWI-DDKMAYLFG
SQGWEFIELGRVFQWILLAAFSLWIYIIYRGVKPWISVKN-VWSVPAWLLWGSGVMVLFL
FFSVLMTPDSNFAISDYWRWMTVHMWVEVTFEVFTTVIVAYLLVQMGLVTRMMAERVIFL
AVMLFFVTAINGISHNFYWIAKPTGIIAVGSVFSTLQVLPLLLLTLDAWQMRQEGG----
RAHEFRVQGKQVVVMEGVWLFILGVNFWNIFGAGVFGSLITLPIVNYYEHATYMTGNHAH
AAMFGVKGNVALAGMLFCLQHLFQKSAWNEKLVRTVFWSFQIGLALMMFLDLFPVGLYQI
YVVLTEGLWYARSTDVIMGPVFATLTYLRSIGGAVFIFGGLIPIIYFVLSGAGRLR-KEV
D----VTTDEWAQYHKEYGNEGGKE---WAAQEEPKVSR---------------------
----
```

```
>MBW7931464_Gammaproteobacterium_BROCD033
M-----------------------------------------------------------
-STNAANVSPGDSNLAMWLLNKKNWFTHFLIVAVISIAGLVYL--GQQTYSGAPPVADFV
--G-PNGEMIASEDMLQHGKHVFHIRGLMLYGSFWGDGAERGPDFTAEAMHRIGASMRSY
YEEEMKQRTGAS---------S---ITDYERDAIAQRVLKEMH----NNTYD-EATNKIT
LNDAQVFAFHELNEHYTRMFTDPTYP-----------------ERM---EG----GQIQG
EETIRDLTAFFFWGGWVSAANRPGQEYSYTHNWPYEPEVGNVPTTATFIWTFISIFALWI
GISVVLYVYGQMKEQPVDVFMGEG---GGGHSLTTSDLEN---GVVRETQRATYKFFVLA
MIVFGIQVFSGVIAAWDFI---KPF------GIN-LNDLIPFSVSRSYHAILQIFWFFMC
WVGYTIFFLPRLTK-VPKGQKSLINLLFLMSCIVAFGTVTGIYVGQRGWL-DETLGYWLG
SQGWEFIELGRLFQYILLAAFSLWIYIIYRGVKPWISMKN-IWSVPAWLLWGSGIMVLFL
FFSVLMTPSSNFAISDYWRWMTVHMWVEVTFEVFTTVIVAYLLVQMGLVTRVMAERVIFL
AVMLFLVTAINGISHNFYWIAKPTGIIAVGSVFSTLQVLPLLLLTLDAWQMRQEGG----
RAHEFRVQGKQVVVMEGVWLFILGVNFWNIFGAGVFGSLITLPLVNYYEHATYMTGNHAH
AAMFGVKGNVALAGMLFCMQHLVQKAAWSEKLVRMVFWSFQIGLALMMFLDLFPVGLYQI
YVVLTEGLWYARSNEVVMGPVFATMTYMRTIGGSVFIFGGLLPIIYFVLSRGGRMR-DEV
D----VSSNEWADYHKEYGHEAGQE---WAAQEEPKVKA---------------------
----
>MCC6201880_Gammaproteobacterium_SJ34
M-----------------------------------------------------------
-ATEAVAPSGSNSNLAMWLLNKKNWFTHFLIVAAISVVGLVYL--GQQTYSGAPPLVNFV
--T-KDGAVVFSKENIERGKELFHIRGLMAYGSFWGDGAERGPDFTAEALHRTVLSMREF
YANEMKARNGVA---------E---LPLYEQDAIAQRVIRELH----GNTYD-ETAQHVV
LNEAQIYAFEELNKHYTRMFTDPTYP-----------------DRM---DPV---NQLSG
ADNIRAVTAFFYWGGWVSAANRPGESYSYTHNWPPDPDAGNSATAATFIWTFASIFALWI
GISIVLYVYGQMKQQPIDLFEAQN--SENSHWLTTSDLEN---GYVRPTQRATYKFFAPA
MIVFGCQVLAGIIGATDFL---RPF------GIN-LNNLVPFTVARSYHTLLQIFWFFMC
WVGYTIFFLPRLAK-IPNGQKFLINLLFGIAVIVAVGALGGIYTGQRGWISDDKMAYWFG
SQGWEFIELGRFFQLLLLGGFTLWIYIIYRGVKPWLNRRN-VWSVPAWLLYGSGVMVLFL
FFGVLMVPDSNFAISDYWRWMVVHMWVEVTFEVFTTVIVAYLLVQMGLVTRLMAERVIFL
AVMLFFVTALNGISHNFYWIAKPTGIIAVGSVFSTLQVLPLLLLTLDAWQMRQEGN----
KADARRLQGKQTTVMNGVWLFILGVNFWNVFGAGVFGSLITLPIVNYYEHATYMTGNHAH
AAMFGVKGNIAIAGMLFCCQHLFPKSFWNDLLIRRIFWALNIGIALMMFLDLFPVGLYQI
FHVVTTGTWYARSTEIVMGPVFQTLTYLRMIGGAVFVLGGLLPLIWFVLSRGTRLR-QET
N----VETDEWSQYQQEYGKEHGKE---WAAQPEPRA-----------------------
----
>MCC7257357_Gammaproteobacterium_SJ686
M-----------------------------------------------------------
-STKAV-SNTGSSNLAMWMLNKKNWFAQFLIVAAISVVGLVYL--GQQTYSGAPPLVDFV
--S-STGEVVASEATIKRGKEVFHLRGLMGYGSFWGDGADRGPDFTADALHRTVVAMREF
YTEELKARNGTE---------T---LTAFETDAIGQRVIRELH----TNTYD-EAAGRIV
LTEGQIYALGELNKHYTRMFTDPTYK-----------------DRM---DPV---NQVSG
EDNLRAVSAFFFWGGWVAAANRPGEEYSYTHNWPYDPEAGNYATTATFVWTFISIFALWI
GIMVVLYVYGQMKLQPVDLFDTQS---GNGHSLTTSDLEN---GYVRPTQRSTYKFFALA
VIVFGLQVLAGIISATDFL---RPF------GIN-LNELVPFTVSRSYHTLLQIYWFFMC
WVGYTIFFLPRLTK-VPSGQKFLINLLFVVAAVVAVGAVGGIYTGQRGWFGDDELSYWFG
SQGWEFIELGRFFQLLLLGGFTLWIYIIYRGVKPWLTMKN-IWSVPAWLLWGSGVMVLFL
FFSVLMTPSDNFAISDYWRWMTVHMWVEVTFEVFTTVIVAYLLVQMGLVTRLMAERVIFL
```

```
AVMLFFVTAINGISHNFYWIAKPTGIIAVGSVFSTLQVLPLLLLTLDAWQMRQEGG----
RANELRVQGKQAHVMEGVWVFILGVNFWNVFGAGVFGSLITLPLVNYYEHATYMTLNHAH
AAMFGVKGNVALAGLLFCCQHLFQKSAWNDKLVSTSFWSLNIGVGLMMFLDLFPVGVYQI
WLVLTEGFWYARSTEVITGPVFVTLTYLRMIGGALFVLGGLLPLIWFVLSRGNRLQ-KEA
E----VGTDEWAQYQKEYGKEAGRE---WATQEEPRV-----------------------
----
>CBE69502_Candidatus_Methylomirabilis_oxyfera_DAMO_2437
M--------------------------------------------------------SPN
-PSGTAAKGKNERTFAQVLLIKKYWWLHALIVTAISTIGLIAL--GVWTYAGAPPLVNFV
--S-KSGDVVIAEHSMNRGKQVFHLKGLMLYGSFWGDGAERGPDFTAEALHRTFVSMGKY
YEMQIEKEQGR----------P---ATQDEKDGIAGKVKREIH----QNGYD-AAAGVIR
LNDAQIFAYNELVDHYTKMFTDPTYE-----------------EAFQKGRIQ---SYVSN
PEDIKGLAGYFFWGGWVAGANRPGEIYSYTHNWPYDPDAGNLPTYATYIWSFLSILVLFA
GTMLVLYVYGEMKSLPGEPFNGRD------WSLTTVDLENKGDAYVRPTQRATYKFFAFA
VILFLVQVLAGILGAEDFV---GGGPGEAI-LGA-FGLVIPFSVVRSYHAIVQIYWFFMA
WVGYTLFFLPRISK-VPNGQRFLINLLFALCVLVGAGALFGIYAGHTGML-TDDMAYWFG
SQGWEFLELGRFWHILMLASFCLWVYIIFRAVKPWITSQN-LWSVPAWLFYGSGIMVLFL
FFGMFMTPSQNFAISDYWRWMNIHMWVEVTFEVFTTCIVGYMLVQMGLVNRAMAERVIFL
AVMMFLVTALIGISHNFYWIAKPTGIIALGSVFSTMQVLPLLLITLDAWKMRTERT----
KAHEHLSEGKQRFVMDGVWTFILAVNFWNIFGAGVMGSLINLPIVNYYEHGTYLTNNHAH
GAMFGVKGNIAIAGMLFACQHLFQRSAWNEKLIKTVFWSLQVGIVMMMLMDMFPVGLYQL
AHIFQYGFWYGRQQSFVTNEVWHTLTWLRSIGGVVFLFGGVLPLCWFILSRAGRMV-REA
AV---VEEGEWTIYDREKAK----EREAWAASDEAF------------------------
----
>WP_107560883_Candidatus_Methylomirabilis_limnetica
M--------------------------------------------------------SPN
-PNEAVSKAKMHRTFAQALLIKKYWWLHALIVTAISVIGLIAL--GVWTYAGAPPLVNFV
--SSTTGEVVIPEWEMNRGKQVFFLKGLMTYGSFWGDGAERGPDFTAEALHRTSLSMGTH
YRNEIAKTR------------A---VTQDDEDMIKTRVAREIH----TNLYD-SAAGVIR
LNDAQIFAYHELNTHYTRMFTDRTYV-----------------EAFQNGRIE---SYIKN
PADIKALTGFFFWGGWVSGANRPGEVYSYTHNWPYDPAVGNFPTYATYIWSFLSIFVLFL
GTMLVLYVYGEMKTLPGEPFNGRD------WSLTTVDLENKGDAYVRPTQRATYKFFAFA
VILFLVQILAGIISAEDFI---GGGPVDAM-LGM-VGLTIPFSVARGWHLIVQIYWFFMA
WVGYTIFFLPRISK-VPNGQRFLINLLFTLCVIVGAGALFGIYLGHTGFFANDEMAYWFG
SQGWEFLELGRFWHVLMLISFCLWVFIIFRAVKPWLTSQN-MWSVPAWLFYGSAIMVLFL
FFGMFMTPQQNFAISDYWRWMNIHMWVEVTFEVFTTCIVGYMLVQMGLVNRAMAERVIFL
AVMMFLVTALIGISHNFYWIAKPTGIIALGSVFSTMQVLPLLLITLDAWKMRTERV----
KAHEAVAEGKQRFVMDGVWTFILAVNFWNIFGAGVMGSMINLPIINYYEHGTYVTSNHAH
GAMFGVKGNIAIGGMLFACQHLFQRSAWNEKLIKTVFWSLQIGLVMMMMMDMFPVGLYQV
AAIIQHGFWYGRQQSFITNEVWHTLTLLRAIGGAVFLFGGVLPLTWFILSRGTRMV-REA
AT---VEEGEWTIYDKEKAK----EREAWAASDEAF------------------------
----
>CBE69496_Candidatus_Methylomirabilis_oxyfera__DAMO_2434
MR----------------------------------------------------------
-SSSSSGMGKTNRTFGQALLIKKYWWLHALIVTVISVIGLVAL--GVWTYTSAPPLTNYV
-LS-STGETVIPEWQIQRGKQVFHLKGLMTYGSFWGDGGERGPDFTAEALHHTYVSMSKF
YENEIAKER------------P---VTQADRDMISVRVKREIH----ENGYD-AAANIIR
INPAQVFAYQELITHYTRMFTDATYE-----------------EAFMKGRIE---NHISS
```

```
PEDLKALAGYFFWGGWVSGANRPGFDYTYTHNWPPDPLVGNTPTFETYLWSFISIFVLFC
GTMLVLYVYGEMKVLPGEPFNGRD------WSLTTVDLENKGDAYVRPTQRATYKFFAFA
VILFLVQVLAGILSAEDFV---GGGPGSAIATTV-LGFTIPFTVTRGWHTIVQIYWFFMA
WVGYTLFFLPRISK-VPNGQRFLINLLFTLCLIVGAGALFGIYLGHTGYM-TDDMAYWFG
SQGWEFLELGRFWHILMLASFCLWVYIIFRAVKPWITSQN-LWSVPAWLFYGSGIMVLFL
FFGMFMTPSQNFAIADYWRWMNIHMWVEVTFEVFTTCIVGYMLVQMGLVNRAMAERVIFL
AVMMFLVTALIGISHNFYWIAKPTGIIALGSVFSTMQVLPLLLITLDAWKMRTERT----
KAHENIAEGKQRFVMDGVWTFILAVNFWNIFGAGVFGSLINLPIVNYYEHGTYLTGNHAH
AAMFGVKGNIAIAGMLFACQHLFQRSAWNEKLIKGIFWSLQVGLVLMMMLDLFPVGLYQV
ATVFKEGLWAARAQAHVTDSVWITLTWMRTIGGAVFLFGGVLPLVYFILSRAGRMV-REA
SV---VEEGEWTIYDREKAK----EREAWAAGDEAF------------------------
----
>WP_107560884_Candidatus_Methylomirabilis_limnetica
M--------------------------------------------------------SPH
-PSTPVSKAKVYRTFAQALLIKKYWWLHALIVGLISVVGLIAL--GVWTYASAPPLVNFV
AVS-NSGTVVIPEWEIQRGKQVFFLKGLMTYGSFWGDGGERGPDYTAEALHHTSVSMRKY
YSAEIVKSR------------P---LTQDDLGMIDSRVKREIH----TNLYD-EKAGVIA
LNDAQIFAYQELITHYTRTFTDATYE-----------------EAFMKGRIE---NHISN
PADLKALASFFFWGGWVSGANRPGEDYSYTHNWPPDQSVNNYPTFPTYLWSFISIFVLFA
GTMLVLYIYGEMKTLPGEPFNGRD------WSLTTVDLENKGDAYVRPTQRATYKFFAFA
VILFLVQILAGILSAEDFV---GGGPGSFLATSV-LGFTIPFTVTRGWHLIVQIYWFFMA
WVGYTLFFLPRISK-VPNGQRFLINLLFTLCVIVGAGALFGIYFGHTGWFANDEMAYWFG
SQGWEFLELGRFWHILMLSSFCLWVFIIFRAVRPWLTSQN-MWSVPAWLFYGSGMMVLFL
FFGLFMTPTQNFAIADYWRWMNIHMWVEVTFEVFTTCIIGYMLVQMGLVNRAMAERVIFL
AVMMFLVTALIGISHNFYWIAKPTGIIALGSVFSTMQVLPLLLITLDAWKMRTERI----
KAHEAVAEGKQRFVMDGVWTFILAVNFWNIFGAGIMGSMINLPIVNYYEHGTYLTGSHAH
AAMFGVKGNIAIAGMLFACQHLFQRSAWNEKLIKTIFWSLQIGLSLMIGLDLFPIGLYQV
ATVFKEGLWAARAQSHVTDSVWVTLTWLRTIGGAIFLFGGVLPLAWFILSRGTRMV-REA
AT---VEEGEWTIYDKEKAK----ERESWAASDEAF------------------------
----
>MBI3325094_Nitrospinae_NC_groundwater_833_Pr1_B
MS------------------------------------------------------SSTS
-RASRSHNGSRSRTFAQILLVKRYWWLHLLIVAAISTVGLVAL--GTWTYTGSPPLADFV
--SSTTGRTVIPLDQIQRGKEVFHLKGLMDYGSFWGDGATRGPDFTADALHRTVVYMRSF
YENEMKDR-------------P---VTQFERDGIAARVQRELH----ANAWD-ETAGVIR
INDAQIHAYRELIGHYTRMFTDSTYP-----------------EAFQLGRIT---GYISD
PADLKALAAFFFWGGWVSAADRPGESYSYTHNWPYDPAAGNTPTFATMFWSVASILGLFL
GIMLVLYVYGQMKTLPGDPFNGAN-----GGTLTTIDLENKGDAYVRPTHRSTYKFFAFA
VILFLVQVLAGILSAEDYV---RGGPGTAA-LNL-LGMTIPFTVTRSWHLILQIYWFFMC
WVGYTIFFLPRLSR-VPNGQRFLINLLFAVCVLVGAGALFGIYLGQTGRL-SDRVAYWFG
SQGWEFLELGRFWHILMLSAFVLWVGIIFRAVKPWITSQN-LWSVPAWLFYGSGIMVAFL
FFGMLMTPGQNFAISDYWRWMNIHMWVEVTFEVFTTCIIGYMLVQMGLVSRLVAERIIFL
AVMMFLVTALIGISHNFYWIAKPSGIIALGSVFSTMQVLPLVLITLDAWRMRAEKF----
RANEYLVQGKQQFVMDGVWMFILAVNFWNIVGAGVFGSLINLPIVNYFEHSTYLTANHAH
AAMFGVKGNVALGGMLFCCQHLFQRASWNAKLVKTVFWALNGGLVLMMFFDLFPVGVYQL
SVVFTHGFWYARSQEVVTGPVFTTLTWLRTLGGVVFLFGGVLPLVWFILSRARRLV-REV
E----VQEGEWTVYDKEKGK----D---WAAQEEEPIPSA--------------------
----
```

```
>MBI2962797_Deltaproteobacterium_NC_groundwater_687_Ag
M-------------------------------------------------------QKSN
-VNRRNGSSKSRRNFGTWLLAKPNWFIQFAIVSGISIVGLIAL--GSWTYSGAPPRVAMV
--SAASGEPVVPIEQIRRGQELFHIRGLMSWGSFWGDGAERGPDFTADALHRTVVGMRSF
YERQMEKER------------P---LTQSDKDAITVRVQREIK----QNGYD-AAAGVIR
INDAQIHAYEELQTHYKRVFTDPTYP-----------------AKF---RLD---NYITD
PEDLRALTGYFFWGGWVAGAARPGETYSYTHNWPYDPEAGNNPTMPTVLWSFLSILALFA
GAMLVLYVYGEMKALPGDPFNGAN-----GGTLTTIELEK-GYDFVRPTQRATYKFFAFA
VILFLVQVLAGILSAEDFV---GGGPGEAI-VQV-FGISLPFTVVRAYHTILQIYWFFMC
WVGYTIFFLPRLSK-VPNGQRFLINLLFTLCVIVGAGALFGIYFGQMGYL-SDTAAYWFG
SQGWEFLELGRFWHILMLASFVLWITIIFRGVRPWITKQN-MWSVPAWLFYGSGIMVMFL
FFGLGATTTSNFAIADYWRWMTVHMWVEVTFEVFTTCIVGYMLVQMGLLNRAMAERVIFL
AVMMFLITATVGISHNFYWIAKPTGIIALGSVFSTLQVLPLLLITLDAWRMRNEKI----
RAGEHLVEGKQKFVMEGVWLFVLAVNFWNIVGAGVFGSLINLPIVNYFEHGTYLTGNHAH
AAMFGVKGNVALAGLLFCCQHLFPRLAWNEALLRRTFWSLQIGIVLMMTLDLFPVGLYQL
AAVLTHGYWYARTNEFVTGPVFATLTWMRVIGGVVFLFGGVLPLVWFVLSRGPKMV-REL
E----VEEGEWTVYDK--------D---WAAHEEEILRALK-------------------
----
>MCC6766257_Deltaproteobacterium_SJ478
M--------------------------------------------------------HTD
-VKARTRNSRTRQNLGKWMLAKPNWLTQFAIVTAVSLLGLIAL--GTWTYGSAPPRVPFV
--SASTGKEVIPLSDILRGQELFHIRGLMSWGSFWGDGAERGPDFTAEALHHTVVSMRSF
YEAQAASGG------------T---SSQTDRDAIAVRVQREIK----ENGYD-EAAGVIR
INDAQIRAYQDLQTHYARVFTDPTYP-----------------AKF---RLA---NYITD
PNDLRALSAYFFWGGWVSGAKRPGEPYSYTHNWPYDPDAGNVPTTPTVMWSFLSILVLFA
GAMLVLYVYGQMKELPGDPFNGAN-----GGTLTTAELER-GYEFVRPTQRATYKFFAFA
VILFLAQVLAGILSAEDFV---SGGPGTAI-VKV-LGVPFSFTVTRAWHTILQIYWFFMC
WVGYTIFFLPRLSR-VPNGQRFLINLLFALCVIVGAGALFGIYFGHMGYM-SDTASYWLG
SQGWEFMELGRFWHILMLASFVLWIAIIYRGVRPWITKQN-MWSVPAWLFYGSGIMVLFL
FFGLGATPSGNFAITDYWRWMTVHMWVEVTFEVFTTCIVAYLLVQMGLLNRSMAERVIFL
AVMMFLITAIVGISHNFYWIAKPTGIIALGSVFSTMQVLPLLLMTLDAWRMRNEKL----
RAGEHRAQGKQKFVMEGVWLFILACNFWNIVGAGVFGSLINLPIVNYFEHGTYLTHNHAH
AAMFGVKGNIALAGVLFCCQHLFPRAAWNEKLLRTSFWSLQAGLVLMMVLDLFPVGLYQM
AAVVTHGFWYARTNDFVTGPVWVKLTWLRTIGGVIFLFGGVLPLLWFILSRGRVLV-REL
E----LEEGEWTVYEK--------D---WAAHEEEIEHALTS------------------
----
>OQY65997_Polyangiaceae_bacterium_UTPRO1
M--------------------------------------------------------HTD
VVKARGRTSRSRQNIGKWMLAKPNWLTQFAIVTAVSVTGLVAL--GMWTYGSAPPRVPFV
--SASTGKEVIPLADIVRGQEIFHIRGLMSWGSFWGDGAERGPDFTAEALHHTVVSMRSF
YESAASKDR------------P---LAQADRDAIAMRVQREIK----DNGYD-EAAGVIR
INDAQIRAYEELQTHYTRVFTDPTYP-----------------AKF---RLA---NYITD
PNDLRAITAYFFWGGWVSGAKRPGETYSYTHNWPYDPEAGNVPTMPAVLWSFLSILVLFV
GTMLVLYVYGQMRDLPGSPFNGAH-----GGTLTTSELER-GYEFVRPTQSATYKFFAFA
VILFLIQVLAGILSAEDFI---GGGPGEAI-VKV-FGLALPFTVVRAWHTILQIYWFFMC
WVGYTIFFLPRLSH-VPKGQRFLINLLFAVCVIVGAGALFGIYFGHMGYL-SDTAAYWLG
SQGWEFVELGRFWHILMLSAFVLWIAIIFRGVRPWITKQN-MWSVPAWLFYGSGIMVLFL
FFGLGATPTGNFAITDYWRWMTVHMWVEVTFEVFTTCIVAYLLVQMGLMNRAMAERVIFL
```

```
AVMMFLVTALVGISHNFYWIAKPTGIIALGSVFSTMQVLPLLLMTLDAWRMRNEKL----
RAAEHREAGKQQFVMEGVWLFILACNFWNIVGAGVFGSLINLPIVNYFEHGTYVTGNHAH
AAMFGVKGNIALAGVLFCCQHLFPRAAWNEKLIKTSFWSFQIGLVLMMTLDLFPVGLYQM
AAVVQHGFWFARTNEFVTGPVFVTLTYLRVIGGMVFLFGGLLPLLWFILSRGPRLV-REL
D----IEEGEWTVYGK--------D---WAAHEEEILSALK-------------------
----
>MCC6850618_Deltaproteobacterium_SJ588
M--------------------------------------------------------NAN
-VRTRTWNSRTRQNLGKWLLAKPNWLTQFAIVTSVSVLGLVAL--GTWTYGSAPPRVPFV
--SATTGKEVIPLADLARGQELFHLRGLMSWGSFWGDGAERGPDFTAEALHRTVVSMRSF
YEAESAKNQ------------PSPEAAQADRDAITVRVQREIK----HNGYD-EAAGVIR
INDAQIQAYQDLQAHYTRVFTDPTYP-----------------ARF---RLA---NYITD
PNELRALTAFFFWGGWVSGAKRPGETYSYTHNWPYDPDAGNFPTTPAVVWSFLSILVLFA
GAMLVLYVYGQMKDLPGDPFNGAK-----GGTLTTYELER-GYEFVRPTQRATYKFFAFA
LILFLVQVLAGILSAEDFV---SGGPGEAI-VKV-LGLSLPFTVVRAWHTILQIYWFFMC
WVGYTIFFLPRLSR-VPKGQRFLINLLFALSVTVGAGVLFGIYFGHMGYL-TDSAAYWLG
SQGWEFMELGRLWHIMMLGAFVLWIGIIFRGVRPWITKAN-MWSVPAWLFYGSGIMVLFL
FFGLGATPYQNFAITDYWRWMTVHMWVEVTFEVFTTCIVAYLLVQMGLMNRAMAERVIFL
AVMMFVVTAVVGISHNFYWIAKPTGIIALGSVFSTMQVLPLLLMTLDAWRMRNEKL----
RAAEHQAAGRQTFVMEGVWLFVLACNFWNIVGAGVFGSLINLPIVNYFEHGTYVTGNHAH
AAMFGVKGNIALAGVLFCCQHLFPRASWNEKLIKTSFWSFQIGLVLMMTLDLFPVGLYQM
AAVVTHGFWYARTNEFVTGPVWVILTWLRTIGGVIFLFGGLLPMLWFILSRGLKLV-REL
E----LEEGEWTVYDK--------D---WAAHEEEIVHALK-------------------
----
>OFW12443_Acidobacteria_RIFCSPLOWO2_02_FULL_67_21
MV------------------------------------------------------NHRN
RPNGTGNGSKTQQNFGKWLLAKRNWWVQFTVVAGISLVGLLAL--GVWTYAGAPPLTRFV
--SSATGETVVPLDGIQRGREVFHLRGLMAWGSFWGDGAERGPDFTADALHRTNVAMKAY
YEQGIAKER------------P---VTQTDRDAIAVRVQREIK----ANGYD-EAADVIR
VNDAQIYALRELEAHYTRMFTDPTYS-----------------EAF---TLD---GYITD
PDDLKALAAFFYWGGWVAGAARPGETYSYTHNWPYDPEVGNLPTMPTVVWSFLSILVLFA
GAMLVLYVYGQMKDLPGDPFNGAN-----GGTLTTIELEK-GYDFVRPTQRATYKFFAFA
MVLFLVQVLAGIISAEDFV---QGGPGQAI-ISL-FGISIPFTVARSWHTILQIYWFFMC
WVGYTIFFLPRLSR-VPKGQRFLINLLFAMSVIAGAGVLFGIYFGQMGYM-SDTASYWFG
SQGWEFLEMGRFWHIAILAAFVLWIAIIFRGVRPWITRQN-LWSVPAWLFYGSGIMVAFL
FMSLGATTSGNFAIEDYWRWMNVHMWVEVTFEVFTTCIVGYMLVQMGLLNRAMAERVIFL
AVMMFLVTAIVGISHNFYWIAKPTGIIALGSVFSTLQVMPLLLITLDAWRMRSEKI----
RAGEHQQEGRQKFVMEGVWLFILAVNFWNIVGAGVFGSLINLPIVNYFEHGTYLTGNHAH
AAMFGVKGNIALGGMLFCCQHLFPRAAWNEKLIKNSFWSLQIGIVLMMTLDLFPVGMYQL
AAVLTHGFWYARTNEFVTGPVFTTLTWMRVIGGMVFLFGGLLPLMWFILSRGRVLV-REL
E----VEEDEWTVYEK--------D---WAAHEEEILRALK-------------------
----
>MBI2828937_Acidobacteria_NC_groundwater_629_Ag_B
M-----------------------------------------------------------
-NNNRHNNSRTRQNFGKWLLAKPNWWVQFSVVAGISLAGLLAL--GTWTYMGAPPYTSFV
--SASTGETVIPYATVARGKEVFHLRGLMSWGSFWGDGAERGPDFTADALHRTVVSMRSY
YENELSRDR------------P---VTQADKDAISVRVQREIHENTRENGYD-EQADVIR
INDAQIQAYRDLIVHYTRMFTDPSYP-----------------EKF---NLD---GFISD
```

```
PDDLRALTAFFYWGGWVAGANRPGETYSYTHNWPYDPAAGNNPTMATVLWSFLSILALFA
GAMLVLYVYGQMKELPGDPFNGAN-----GGTLTTFELEK-GYDFVRPTQRATYKFFAFA
VILFVVQVLAGILSAEDFV---SGGPGMAI-TSV-LGISLPFSVVRAWHTLLQIYWFFMA
WVGYTIFFLPRLSR-VPKGQRFLINLLFTLCVVAGAGVLFGIYFGQMGYM-SDTMSYWFG
NQGWEFLEMGRFWHIAILGAFALWIFIIFRGVRPWLTKQN-MWSVPAWLFYGSGIMVLFL
FMSLGATMGGNFAITDYWRWMNVHMWVEVTFEVFTTCIVGYMLVQMGLLNRAMAERVIFL
AVMMFLVTAIVGISHNFYWIAKPSGIIALGSVFSTLQVLPLLLITLDAWRMRNEKV----
RAGEHLQEGKQKFVMEGVWLFILAVNFWNIVGAGVFGSLINLPIVNYFEHSTYLTGNHAH
AAMFGVKGNVALAGLLFCCQHLFPRAQWNEKLIRTSFWSLQSGIVLMMTLDLFPIGLYQL
AAVLTHGFWYARTNDFVTGPVWVTLTWLRTIGGVVFLFGGVLPLTWFVLSRGRSLL-REV
E----IEEGEWTVYDR--------D---WASHEEEILQVLKS------------------
--RE
>CBL43845_Gammaproteobacterium_HdN1_NorZ2
MST---------------------------------------------------------
-NSLLGGSIRAKKNIALWLVSKKNWILQFLIVAAICTGGLLYL--GAQTYISAPPIVNFV
--N-DKGDTVFSEQQIMDGKHVWSIRGLMAYGSFWGDGAERGPDFTADSLHRSVVAMREF
YAKEKKAELGVD---------A---LPVYETDAIAARVIRELH----NNAYD-EEAGIVR
LNPAQEYAFAEVREHYRKMFNDATYS-----------------EAF---HPQ---GLISD
PEQITSLTSFFFWGGWASAANRPGETYSYTHNWPHDPGAGNAPTPATWIWSFLSILALFL
CIVVVLYVYGQMRELPIDVFGTST---ENPFALTTHDLEN---GYVRPTQKATYKFFALA
MILFGVQIFAGIAAAWDFV---KPF------GIS-LNDFLPFTASRSFHAIIQIVWFFVC
WVGYTIFFLPRLSK-LPSSQRTMINSLFAMIVIVGLGTLIGVYLSTMGFL-DGWVAYWFG
TMGWEFMEMGRFFQLFLLVTFSFWIYIIYRGVKNWLTRKN-IWSVPAWLLYGSGIMVLFL
FFGVLILEDQNFAIADYWRWMVVHMWVEVTFEVFTTVIVGYLLVQMGLVTRLMAERTIFL
AVMLFLITAVIGISHNFYWIAKPTGIIALGSIFSTLQVLPMLLLTLDAWKMRQELG----
RAHDLRQQGRQVHVMEGVWLFILGVNFWNIVGAGVLGSLINLPIVNYYEHATYLTPNHAH
AAMFGVKGNIAIAGMLFCLQHMFHKSAWSETRIKGIFWSLQIGLVLMMVLDMFPVGLYQV
WLTLQDGLWHARSSEVILGPVFASFTYLRVIGIAVFVLGGALPLIWFVLSRGRTLQ-QET
E----VAEGEWTAYED--------D-EPWVAVKK--------------------------
----
>Ga0257115_100087344_SI112_100m_VIRAL
MTHLL--SLIVLIIALMGFLYLISWVFKIINKIIDIFKIKEEKMTTEFKNGEYIITTKDD
-SETVRSVLSFKDSVSALLLNKKYWFAHFWIVSIISILGLVYM--GGATYMGAPPVPTYV
--A-SNGEVIITEAEIMKGQEIFHLRGLMNYGSFWGDGAERGPDFTAETLHTMAVAMKGY
YAEQLKPGSVTQDDKLRKWKFD---ISEYDEGAIETRVKKELH----NNTYT-EERNRVV
LNDAQIYGIEYVRWYYEQMFTNADHP-----------------EAF---HPI---GYITD
KTDLNNLAAFFYWGSWTSAADRPGDDFSYTSNWPYDPQAGNEPPPSLMMWSFLSIFILWI
GIMLVLYVYGQMRTMPGGPFDMGA----RGPTLTTADLES---GYVRPTQLATYKFFALS
IVMFGLQVLAGILGAMDFA---NPF------AHK-LANILPFNILRSYHTLLQIYWFFVA
WVGYTIFFLPRLSE-VPKHQEFLINLLFGICVTVGVGGIVGVYLGQAGYM-TGWAAYWFG
SQGWEFMELGRFFQMLLLGGFSLWIYIIYRGVKPWITKKT-FWSVPAWLLWGSGIMVFFL
FFGLFATIDSNWAIADFWRWMVVHMWVEVTFEVFTTVIVGYILVQMGLVTKVMAERVIYL
AVILFLITATIGVAHNFYWIAKPESVIALGSVFSTLQILPLLLLTLDAWQMRKEGI----
GAYQSRDEGKQTFIMEGVWLFVIAVNFWNIVGAGIFGSLINLPIVNYFEHATYLTGNHAH
AAMFGVKGNVALAGILFVCQHLFTTESWNDKIIKYAFWSLNIGVVLMMFLDLFPAGLYQL
SIVLEDGFWLARAQETINGTVFQTLTYFRSIGGAVFVT-GVVALIWFILSRSMHLK-SET
GKA--ASTQDWTTAEN--------D---WQK-----------------------------
----
```

```
>Ga0209665_10050414_SI074_LV_200m
M---------------------------------------------------------DE
-STSNGTKKEFKGTLAHMLLSKKYWFIHFWIVSVISIMGLVYM--GAATYMGAPPVPDFK
--N-ANGELVISEADIMKGQEIFHLRGLMNYGSFWGDGAERGPDFTAEALHAITVSMKGY
YANQIKPGSVTQDDRLREWRFR---ISDYDEGAIEARIAKELH----VNTFN-EDTNTVI
LNDAQTDSFEYINWYYTQMFTNADHP-----------------EAF---YPT---NYISD
KQQLKELSAFFFWGAWTSAADRPGDAFSYTSNWPYDPAAGNTPPPGLMMWSFLSIGILLI
GIMIVLYVYGQMRELPGSPFKLNP----TGKILTTPDLEA---GYVRPTQKSTYKFFAVA
IVAFGLQVLAGILGAMDFT---NPF------AHE-LAGVLPFNILRSYHTLLQIYWFFLM
WVGYTIFFLPRLSK-VPKGQNTLINILFGICVAVGLGGIFGVYLGQAGYL-TGWAAYWFG
SQGWEFMELGRVWQTLLLGGFTLWIFIIYRGVKPWITKKT-FWSVPAWLLWGSGIMVFFL
FFGLFATIDQNWAIADFWRWMVVHMWVEVTFEVFTTVIVGYILVQLGLVSRAMAEKVIYL
AVILFLITATIGVAHNFYWIAKPEGVIAMGSVFSTLQILPLLLLTLDAWQMRKEGH----
GAYTNMANGTQNHAMDGVWLFILAVNFWNIFGAGVLGSLINLPIVNYFEHATYLTGNHAH
AAMFGVKGNIALAGMLFVLQHLVKPEFWNPKLIKLSFWSLNLGIVFMMFFDLFPAGLYQF
SIVLEDGLWLARAQETITGTVFQTLTYFRSIGGLVFVV-GVVSLIWFVLTRGTKLK-DET
GKTVQIQTQEWNDLDN--------D---WVDEEHAVERGVEEEIKEINNFQVQRSAWKYK
K---
>St16_OMZ_317E_VIRAL
MVNIIILMGLMMIFMVALAFVIFLSLV-ATKNIIINFF-----------------NTNTN
-RNNMKQDKEFKGTVAHMLLSKKYWFAHFWVVSVISIMGLLYM--GGATYMGAPPVPDYK
--N-ANGEMVFSSDEIMKGQELFHLRGLMNYGSFWGDGAERGPDFTAEALHATTVAMKGY
YAEQLKPGSVTQDDKLRKWRFD---IEEYDKGAIEAQVAKELH----INTYD-EDTNTVV
LNDAQVFSFEYINWYYTQMFTNADHP-----------------EAF---HPL---NYITD
ETQLRELSAFFFWGAWTSAADRPGDDFSYTSNWPYDPAAGNEPPPSLMMWSFLSIGILMA
GIMIVLYVYGQMRELPGSPFKLNA----TGQLLTTPDLEA---GYVRPTQKSTYKFFAIA
IVAFGLQVVAGILGAMDFA---NPF------AHQ-LADVLPFNILRSYHTLLQIYWFFLM
WVGYTIFFLPRLSK-VPKGQGTLINILFTICVIVGLGGVFGVYLGQAGYM-TGWAAYWFG
SQGWEFMELGRFWQMLLLGGFTLWIFIIYRGVKPWITKKT-FWSVPAWLLWGSGIMVFFL
FFGLFATIDQNWAIADFWRWMVVHMWVEVTFEVFTTVIVGYILVQMGLVSRSMAEKVIYL
AVILFLITATVGVAHNFYWIAKPEGVIAMGSVFSTLQILPLLLLTLDAWQMRKEGH----
GAFKNLANGTQNHVMDGVWLFILAVNFWNIMGAGVLGSLINLPIVNYFEHATYLTGNHAH
AAMFGVKGNIALAGMLFVLQHLVKPEFWNPKLIKTAFWSLNLGIVLMMFFDLFPAGLYQF
MIVLEDGFWLARAQETITGTVFQTLTYFRSIGGLVFVV-GVVSLIWFVLTRGTKLK-DET
GKS--IQTQNWNDLDN--------D---WIDEQHAVANGVEEEVNELSTFE--QALIKHN
KKEQ
>WP_026951429_Algoriphagus_mannitolivorans
M-----------------------------------------------------------
---------NTEPSFLTYLMKPKNWWLPFLLIFTVSISGLIFI--GYQTYNEAPPIPDYV
--D-SEGNMIISQAEILAGQEVFHRYALMEYGSMFGDGALRGPDFTAQTLHTLAESIKEF
YRVQ--SPL------------D---PKPFELAGIEQMVKNEIK----KNRYQ-TRSNQTE
LTPAQVFAVSEINTFYQRVFLDPEFH-----------------EAF---KPA---GYLTD
PKEIEDLSKFFYWGAWVCSVARPGETYSYTHNWPFDPEAGNTPTASTVLWSMLGLFGLVL
GLGAVMYYYGQFEQLTEEYYAKGA-----SEMVTEEKVRS---FRPTPTQRATFKFFFVA
ILLFFIQVLAGVLTVHDFVGFTSFF------GLD-LQALLPVTISRSWHVQISLYWISAC
WVGISFFILPILAKGEPKGQLKLINLLFAIFFVLVAGSLVGIYLGPMNML--GSLSRWLG
HQGWEFVELGRLYQYMLLAVFALWAVIVYRGLKGVMAKAK-PWELPNWLVYAVVCILILL
VSGFVASPETNFVVADFWRWMVVHMWVEAFFEVFTTIIVGYMMVMMGLVNRQAVTKVVYL
```

```
AALLFLGSGLLGISHNFYWNAKPVGTLALGSVFSTLQVVPLILLTLEAWRMQRMPQL-L-
AKKNNESGRSSLFAMPGVFLYLLGVNFWNFFGAGVFGLIINLPIVNYYEHGTYLTVNHGH
AALMGVYGNLSIAGILFALRYLVQPTSWNPQLVKLSFWSINLGLVFMVVLDLFPAGIHQL
FTVMDQGYWFARSQDFIQSTPFQTMTWLRIVGGSLFLLGGVIPLTWMVIKSSRQLK-YEQ
P------FVLNRFHQERKEM----E-----------------------------------
--LH
>WP_130276701_Cecembia_calidifontis
M-----------------------------------------------------------
---------KPDTKFLNYIMKQKNWWLPFMVIFVISIVGLLFI--AYQTYEEAPPIPHYV
--D-EKGEEIITQEQILRGQEVFHRYALMEYGSMFGDGALRGPDFTAQALNERAKSMFRF
YQESLNENG------------S---RSEFESEGILRKVQKEIK----ENTYD-ENNNQVN
ISFAQVAAIRDLEGFYQEMFLNPDFH-----------------EAF---KPT---GYISD
PTEIKDLSAFFFWGSWVCSVERPGEKYSYTHNWPYDEFAGNVATPSTLIWSIVGLLGLVL
GLGIVLYYYGQFEQLSEEYYAKGA-----SEMVTEEKLQQ---YKPTPTQRATFKFFYVA
VILFLIQVLAGVLTVHDFVGFTKFF------GWD-IQEVLPVTISRSWHVQLSLFWISAC
WVGISFFILPLLAKSEPKGQLFLINLLFGIFFVMVGGSFVGIFMGPMGML--GDYSRWLG
HQGWEFVELGRLYQYFLLAIFALWAIIVYRGVKNTLRPGM-PWGLPNWLVYSIVCILLLL
LSGFVARPETNFVIADFWRWMVVHMWVEAFFEVFTTIIVGYMMVMMGLVNRQAVVKVVYI
AALLFLGSGLLGISHNFYWNAKPVGTLALGSVFSTLQVVPLILLTLEAWRMQRMPYL-L-
STKQKERHRSSLFAMPGVFLFIIGVTFWNFFGAGVFGLIINLPIANYYEHGTYLTVNHGH
AALMGVYGNLSIAAILFALRFLLKPEAWNTQLVKTAFWSINIGLLLMVTLDLFPAGIHQL
VAVMEEGYWFARSQEFIQSVPFQAMTWLRIVGGAFFCIGGLFPLTWFVLKSAKHLK-PAY
E----MEKIDHHLEEVEEEI----E-----------------------------------
--TV
>Ga0066828_100166541
------------------------------------------------------------
-------------------MKSTNWWKYLLAVLVVGASGVIFM--GISTYKDAPPKPDYI
--S-PSGVEIVQQASVERGQLVFQRYALMEYGSMFGDGAARGPDFTAEALHQVAVEMNTF
YGQQVASGSTD----------E---LSQIEKDGISVRVKRELK----ANRYD-RERNIVV
LTEGQAYAAGRLAEYYNSKF-KGDHK-----------------EAF---KPA---GYITD
DAELKDLSAFFFWGAWVCAVERPGGESSYTHNWPFDEYAGNTPTPSVILWSVIGMLFLIF
GLGAVLCTYSYYSKASQLQ--------VKEDPVDNKSVDA---SAPTASQRATYKFFVVA
VSLFFIQILAGVLTIHDFVGFTTFF------GYN-ISELLQITITRSWHVQLSILWIATC
WIAGSIFILPGIYRQEPKRQVLLINVLFGLLVSVVIGMFVGCFLGPKNLL--GDHWRLLG
NQGWEFVELGKLWQVVLFAALVMWSVIVYRGVKPALKGQS-AFSLPYWILYSVIAITILF
LSSFVGGKNTNFVIADFWRWCVIHMWAECFFEVFTTMVIACYMVLMGLVSRQGATRVIYL
ATLLFLGSGLLGISHNFYWNAKPAALMAMGSVFSTLQVIPLVLLTLEVWKFVRVPGYSF-
GPSTNSAVTGSSFGFSEAFLFLIAVNFWNFFGAGVFGLIINLPIMNYYEHGTYLTVNHGH
AALMGVYGNLSIAAVLFCSRHVITSRRWNVRLLRSTFWAINVGLLLMVVMDTFPAGVLQF
RSVVENGYWVARSQEFILGSAFQVLTWMRAVGGVLF-FAGVIPLLYFMISRLNSLK-AAS
T----VYSSEESVLIKEELT----K------------CDA--------------------
----
>Ga0187827_100070556
------------------------------------------------------------
-------------------MKSTNWWKYLLAVLVVGASGVIFM--GISTYKDAPPKPDYI
--S-PSGVEIVQQASVERGQLVFQRYALMEYGSMFGDGAARGPDFTAEALHRVAVEMNAF
YGQQVASGSTD----------E---LSQIEKDGISVRVKRELK----ANRYD-RERNIVV
LTEGQAYAAGRLAEYYNSKF-KGDHK-----------------EAF---KPA---GYITD
```

```
DAELKDLSAFFFWGAWVCAVERPGGESSYTHNWPFDEYAGNTPTPSVILWSVIGMLFLIF
GLGAVLCTYSYYSKASQLQ--------VKEDPVDNKSVDA---SAPTASQRATYKFFVVA
VSLFFIQILAGVLTIHDFVGFTTFF------GYN-ISELLQITITRSWHVQLSILWIATC
WIAGSIFILPGIYRQEPKRQVLLINVLFGLLVSVVIGMFVGCFLGPKNLL--GDHWRLLG
NQGWEFVELGKLWQVVLFAALVMWSVIVYRGVKPALKGQS-AFSLPYWILYSVIAITILF
LSSFVGGKNTNFVIADFWRWCVIHMWAECFFEVFTTMVIACYMVLMGLVSRQGATRVIYL
ATLLFLGSGLLGISHNFYWNAKPAALMAMGSVFSTLQVIPLVLLTLEVWKFVRVPGYSF-
GPSTNSAVTGSSFGFSEAFLFLIAVNFWNFFGAGVFGLIINLPIMNYYEHGTYLTVNHGH
AALMGVYGNLSIAAVLFCSRHVITSRRWNVRLLRSTFWAMNVGLLLMVTMDTFPAGVLQF
KSVVENGYWVARSQEFILGGAFQMLTWMRAVGGVLF-FAGVIPLLYFMISRLNSLK-VAA
T----AHSSEELVLTKEELP----E------------CDVQ------------------
--VA
>MBT3880006_Scalindua_SI054_bin45
------------------------------------------------------------
-------------------MKSTNWWKYLLAVLVVGASGVIFM--GFSTYKDAPPKPDYI
--S-PSGVEIVQRAAVERGQLVFQKYALMEYGSMFGDGAARGPDFTAEALHRIAVEMNDY
YGRQVTNNNLD----------E---LSQIEKDGISIRVKRELK----ANRYD-GERNIVV
LTEGQAYAAERLVEYYSSKF-KGDHK-----------------EAF---KPA---GYITD
DSELKDLTAFFFWGAWVCAVERPGGESSYTHNWPFDEYAGNTPTPSVILWSVIGMLFLIF
GLGAVLCTYSYYSKTSQLQ--------VKENPVNNKSVDA---SAPTASQRATYKFFVVA
VALFFIQIVAGVLTIHDFVGFTTFF------GYN-ISEFLQITITRSWHVQLSVLWIATC
WIAGSIFILPGIYRQEPKRQVLLINVLFGLLVSVVVGMLVGCFLGPKNLL--GDHWRLLG
NQGWEFVELGKLWQVVLFASLVMWSVIIYRGVKPALKGQS-AFSLPYWILYSVIAITILF
LSSFVGGKNTNFVIADFWRWCVIHMWAECFFEVFTTMIIACYMVFMGLVSRQGAIRVIYL
ATLLFLGSGLLGISHNFYWNAKPAALMAMGSVFSTLQVIPLVLLTLEVWKFVRVPEYSL-
RRSANSAVAGSSFGFSEAFLFLIAVNFWNFFGAGVFGLIINLPIINYYEHGTYLTVNHGH
AALMGVYGNLSIAAVLFCSRHVIIARRWNVRLLRSTFWAINVGLLLMVVMDTFPAGVLQF
RSVVENGYWVARSQEFILGGTFQVLTWLRAVGGVLF-FAGVVPLLYFMISRLNSLK-AAA
T----EHSSEEPILIKEEIP----E------------CDVR------------------
--VA
>WP_096896255_Scalindua_japonica
------------------------------------------------------------
-------------------MKSTNWWKYLLAVLVVGASGVIFM--GFSTYKDAPPKPDYI
--S-PEGVEIVQKSAVERGQLVFQKYALMEYGSMFGDGAARGPDFTAEALHRIAVDMNNY
YGRQVTDGNLS----------E---LSQIEKDGISVQVRRELK----ENRYD-MERNIVV
LTEGYAYAAEQLVDYYRLKF-KGNHK-----------------EAF---KPA---GYITD
DSELNDLSAFFFWGAWVCAAERPGGESSYTHNWPFDEYAGNVPTPSVILWSVIGMLFLIF
GLGAVLCTYSYYSKASQLQ--------VKENPVNNKSVEA---SIPTATQRAAYKFFVVA
VALLFIQIVAGVLTIHDFVGFTTFF------GYN-ISEFLQITITRSWHVQLSILWIATC
WIAGSIFLLPGIHRQEPNRQVLLINVLFGLLVSVVVGMLVGCFLGPKNLL--GDNWRLFG
NQGWEFVELGKVWQIVLFAALIMWAVIIYRGVKPALKGQS-AFSLPYWILYSVVAITILF
LSSFVGGKNTNFVIADFWRWCVIHMWAECFFEVFTTIVIACYMVFMGLVSRQGATRVIYL
ATLLFLGSGLLGISHNFYWNAKPAALMAIGSVFSTLQVIPLVLLTLEVWKFVRIPGYSI-
RRSEDSAVTGSSFGFSEAFLFLIAVNFWNFFGAGVFGLIINLPIINYYEHGTYLTVNHGH
AALMGVYGNLSIAAVLFCSRHVITSRRWNVRLLRSTFWAINVGLLLMVLMDTFPAGVLQF
RSVVENGYWVARSQEFILGGAFQVLTWMRAVGGVLF-FAGVIPLLYFMVSRLNSLK-AAA
T----TQKSEEPVLITEELP----E------------RDVQ------------------
--VA
```

```
>Ga0066833_100015614
------------------------------------------------------------
-------------------MQRNNWWKYLLVVLVAGTAGVLFM--GIKTYKDAPPKPDYV
--S-SSGEPVFMRASVERGQQVFQKYALMEYGSMFGDGAARGPDFTAEALHQIAVEMNDY
YGKQFTNGRTE----------E---LTQIEKDGISVRVKRELK----TNYYD-GERDIVV
LTEGQVYAAGQLVEYYRLKF-KGDHR-----------------ESF---KPA---GYITD
DSELEDLAAFFFWGSWVCAAERPGGESSYTHNWPYDEYAGNIPTPSVILWSVIGMLFFIF
GLGAVLCIYGYYAKSSGLQPP------AKEKPVSNESISS---SVPTAGQRATYKFFAVA
VAMFFLQIVAGVLTIHDFVGFTTFF------GYD-IAEYLQITITRSWHVQLSILWISVC
WIAGSIFILPSIHRQESKHQVTLINVLFWLLITVVVGMLVGCFMGPKNML--GDHWRLLG
NQGWEFVELGKLWQIFLFAALIMWAVIIYRGVKPALKGQS-AFSLPYWILYSVVAITVLF
TSGFVSGPNTNFVIADFWRWCVIHMWAECFFEVFTTIIVACYLVFMGLITRQAATRVIYL
ATLLFLGSGLLGISHNFYWNAKPAALMAMGSVFSTLQVVPLILLTIEVWKFARVPDYSLG
KRSENEAVKGTSFGFSEAFLFLIAVNFWNFFGAGVLGLIINLPIINYFEHGTYLTVNHGH
AALMGVYGNVSLAAVLFCSRHLIVSRRWNVRLLRSSFWAINVGLLLMVALDTFPAGVLQF
KLVVESGYWIARSQEFILGGPFQVLTWMRAVGGVVFFFGGVIPLFYFMMTRLNSLK-AAA
P----GHVPADPVRIEEPLP----E------------LNVQ-------------------
--TA
>Ga0066834_100088392
------------------------------------------------------------
-------------------MQRNNWWKYLLVVLVAGTAGVLFM--GIKTYKDAPPKPDYV
--S-SSGEPVFMRASVERGQQVFQKYALMEYGSMFGDGAARGPDFTAEALHQIAVEMNDY
YGKQFTNGRTE----------E---LTQIEKDGISVRVKRELK----TNYYD-RERDIVV
LTEGQVYAAGQLVEYYRLKF-KGDHR-----------------ESF---KPA---GYITD
DSELEDLAAFFFWGSWVCAAERPGGESSYTHNWPYDEYAGNIPTPSVILWSVIGMLFFIF
GLGAVLCIYGYYAKSSGLQPP------AKEKPVSNESISS---SVPTAGQRATYKFFAVA
VAMFFLQIVAGVLTIHDFVGFTTFF------GYD-IAEYLQITITRSWHVQLSILWISVC
WIAGSIFILPSIHRQESKHQVTLINVLFWLLITVVVGMLVGCFMGPKNML--GDHWRLLG
NQGWEFVELGKLWQIFLFAALIMWAVIIYRGVKPALKGQS-AFSLPYWILYSVVAITVLF
TSGFVSGPNTNFVIADFWRWCVIHMWAECFFEVFTTIIVACYLVFMGLITRQAATRVIYL
ATLLFLGSGLLGISHNFYWNAKPAALMAMGSVFSTLQVIPLILLTIEVWKFARVPDYSLG
KRSENEAVKGTSFGFSEAFLFLIAVNFWNFFGAGVLGLIINLPIINYFEHGTYLTVNHGH
AALMGVYGNVSLAAVLFCSRHLIVSRRWNVRLLRSSFWAINVGLLLMVALDTFPAGVLQF
KLVVESGYWIARSQEFILGGPFQVLTWMRAVGGVVFFFGGVIPLFYFMMTRLNSLK-AAA
P----GHVPADPVRIEEPLP----E------------LNVQ-------------------
--TA
>Ga0208638_10055013********
------------------------------------------------------------
-------------------MQRNNWWKYLLVVLVAGTAGVLFM--GIKTYKDAPPKPDYV
--S-SSGEPVFMRASVERGQQVFQKYALMEYGSMFGDGAARGPDFTAEALHQIAVEMNDY
YGKQFTNGRTE----------E---LTQIEKDGISVRVKRELK----TNYYD-RERDIVV
LTEGQVYAAGQLVEYYRLKF-KGDHR-----------------ESF---KPA---GYITD
DSELEDLAAFFFWGSWVCAAERPGGESSYTHNWPYDEYAGNIPTPSVILWSVIGMLFFIF
GLGAVLCIYGYYAKSSGLQPP------AKEKPVSNESISS---SVPTAGQRATYKFFAVA
VAMFFLQIVAGVLTIHDFVGFTTFF------GYD-IAEYLQITITRSWHVQLSILWISVC
WIAGSIFILPSIHRQESKHQVTLINVLFWLLITVVVGMLVGCFMGPKNML--GDHWRLLG
NQGWEFVELGKLWQIFLFAALIMWAVIIYRGVKPALKGQS-AFSLPYWILYSVVAITVLF
TSGFVSGPNTNFVIADFWRWCVIHMWAECFFEVFTTIIVACYLVFMGLITRQAATRVIYL
```

```
ATLLFLGSGLLGISHNFYWNAKPAALMAMGSVFSTLQVVPLILLTIEVWKFARVPDYSLG
KRSENEAVKGTSFGFSEAFLFLIAVNFWNFFGAGVLGLIINLPIINYFEHGTYLTVNHGH
AALMGVYGNVSLAAVLFCSRHLIVSRRWNVRLLRSSFWAINVGLLLMVALDTFPAGVLQF
KLVVESGYWIARSQEFILGGPFQVLTWMRAVGGVVFFFGGVIPLFYFMMTRLNSLK-AAA
P----GHVPADPVRIEEPLP----E------------LNVQ-------------------
--TA
>Ga0208642_10056733*********
------------------------------------------------------------
-------------------MQRNNWWIYLLVVLVTGAAGVLFM--GIKTYKDAPPKPDYV
--S-SSGEEVFTRASIERGQLVFQKYALMEYGSMFGDGAVRGPDFTAEALHQIAVQMNDY
YGRQITNDSSA----------E---LTQIEKDGISVRVKRELK----TNYYD-RERGIVV
LTEGQVYAAERLVEYYRSKF-KGDHR-----------------ESF---KPA---GYITD
DSELEDLAAFFFWGSWVCSAERPGGESSYTHNWPYDEYAGNIPTPSVILWSVIGLLFLIF
GLGAVLCIYGYYAKSAGLQSP------DEENPVNNENVSS---SVPTAGQRATYKFFMVA
VAMFFLQIVAGVLTIHDFVGFTTFF------GYN-ISEYLQITITRSWHVQLSILWISVC
WIAGSIFILPSIHRQESKHQVTLINVLFWLLVTVVVGMLAGCFMGPKNML--GDHWRLLG
NQGWEFVELGKLWQVFLFAALIMWAVIIYRGVKPALKGQS-AFSLPYWILYSVVAIAVLF
TSGFVSGPNTNFVIADFWRWCVIHMWAECFFEVFTTIIVACYLVFMGLITRQAATRVIYL
ATLLFLGSGLLGISHNFYWNAKPAALMAMGSVFSTLQVIPLILLTIEVWKFARVPGYSLG
KRSENAAVKGTSFGFSEAFLFLIAVNFWNFFGAGVFGLIINLPIINYFEHGTYLTVNHGH
AALMGVYGNVSIAAILFCSRQLIVSRRWNVRLLRSSFWAVNVGLLLMVTMDTFPAGVLQF
KSVVENGYWAARSQEFILGGPFQVLTWMRAVGGSVFFFGGVIPLFYFMITRLNSLK-AAA
P----GHVSEDPVRIEEPLP----E------------FNVQ-------------------
--TA
>BAG66139_Geobacillus_stearothermophilus
MEV---------------------------------------------------------
-------NRTVSPNIQTG-RKTTNSFLKSILIFTILISSTVLLVGGYWIFKEMAPRPKEV
-RS-ESGEVLMTKETIIGGQAVFQKYGLMDYGTVLGHGSYMGPDYTAEALKVYTEGMQDY
KAKERYNKPFA----------D---LTDDEKSIIREQVIKEMR----KNRYN-PVTDVLV
LTDAQVYGLEKVRDYYRDVFTNGDGWGLKKGLIK---------ESD---MPKANRAWVAD
SDQIQQIADFFFWTAWLSSTLRIGDEITYTNNWPYYEDAGNTMSFSAVWWSGASVTILIL
FIGIILYVFYRYQLSMQEAYA--------EGKFPVIDLRR---QPLTPSQVKAGKYFVVV
SALFFVQTMFGALLAHYYTEPDSFF------GINWIYDILPFNIAKGYHLQLAIFWIATA
WLGMGIFIAPLVGGQEPKKQGLLVDLLFWALVVLVGGSMIGQWLGVNGYL--GNEWFLLG
HQGWEYIELGRIWQIILVVGMLLWLFIVFRGVKRGLKRESDKGGLIHLLFYSAIAVPFFY
IFAFFIQPDTNFTMADFWRWWIIHLWVEGIFEVFAVVVIGFLLVQLRLVTKKSTVRALYF
QFTILLGSGVIGIGHHYYYNGSPEVWIALGAVFSALEVIPLTLLILEAYEQYKM------
-----MRDGGANFPYKATFWFLISTAIWNLVGAGVFGFLINLPAVSYFEHGQFLTPAHGH
AAMMGVYGMFAIAVLLYSLRNIVKPEAWNDKWLKFSCWMLNIGLAGMVVITLLPVGILQM
KEAFIHGYWASRSPSFLQQDVVQNLLLVRAVPDTIFLI-GVVALLVFAIKALFHLR-KPT
H-----GEGEELPVAN--------H---WMKDRLKNS-----------------------
----
```
