## Supplementary material for "Novel Alphaproteobacteria transcribe genes for nitric oxide transformation at high levels in a marine oxygen deficient zone": NuoL alignment

```
>WP_085441360_NuoL_Magnetofaba_australis
MST-----YKLIVLLPLMGSIIAGLFG----RRLGNQMSQIVTIGGIGLALLLSIKAFFEIAL--GDAP-
AVHETFFTWIPSGDFVVTLGVLVDRLTAIMLIVVTGVSTLVHIYSVNYMEEDPD-------
VPRFFSYLSLFSFAMLSLVTAPNFLQLFFGWEGVGLASYLLIGFWFKKESACNAAIKAFLVNRVGDFGFALGVF
GVFMVFGTLDFLGDKGVFMALTNYQAT---
MMFLGHEFNTMTLICLLLFMGAMGKSAQFFLHTWLPDAMEGPTPVSALIHAATMVTAGVFLVCRASPLFEQSET
ALMVVTVIGAVTAIFAASVGLVQNDIKRVIAYSTCSQLGYMFFAAGVSAYAASMFHLMTHAFFKALLFLGAGAV
IHAMHHEQDMRKMGGLWKKIPLTYALMMIGTLALT--GFPY---------
LAGWWSKDAILESAWAAHTGVGTFAWVIGLIAAFMTTFYSFRLVFMTFHGKPH---DEHHYDHA----------
---------------------------------------------------
HEAPWFMRAPNLLLAVGALLAGYLGHGIIEAG--------WF--
KDAIFLAEGHDALAHAHHAPAHVKWLPFVMFLGGLFLALLMYIWV-----------------------------
------------------------------
PTLPKKVSELCPRGYQFLLNAWYFDKLYDAIFIKPAKAIGKGLWQT-GDATIIDGYGVNGTA-
NLMVRMGAVLKRMQSGYVYHYAFAMLAGVLVLITFYA------------
>WP_011715163_NuoL_Magnetococcus_marinus
MST-----YKLIVLLPLLGSLIAGLLG----RTIGTRMSQMVTIGGIGLSLLLSIQAFWQIAV--NDGE-
VVREIFWSWVISGDFQVTLGVLVDRLTAVMLIVVTGVSTLVHIYSVDYMHEDPD-------
NPRFFSYLSLFSFAMLMLVTSPNFLQLFFGWEGVGLASYLLIGFWFKKESACNAAIKAFLVNRVGDFGFALGVL
AIFMVFGTLDY--SAVFGAARGEFNQT---
MHFLGYEFTTITLICLLLFLGAMGKSAQLFLHTWLPDAMEGPTPVSALIHAATMVTAGVFLVCRASPLFELSET
ALAVVTIIGALTAFFAATVGMVQNDIKRVVAYSTCSQLGYMFFAAGVSAYAASMFHLMTHAFFKALLFLGAGAV
IHAMHHEQDMRKMGGLVKKIPLTYGLMMIGTLALT--GFPG---------
LAGFFSKDAILESAYAAHSATGTFAFWLGIMAAGMTTFYSFRLVFMTFHGKPK---DHHAYDHA----------
---------------------------------------------------
HEAPWFMRGPNIALAIGALFAGYLGVPLIEAG--------WF--
KEAIVLAAGHNALEHAHHVPAWVKWLPFVMFVLGLSVAVVLYVLA-----------------------------
------------------------------
PTLPAKIAQMCPRGYNFLLNAWYFDKLYDKLFVKGAQCLGQGLWKT-ADETIVDGYGVNGWA-
RTLGRWGASLRRMQSGYVYHYAFAMFVGVLALATFYS------------
>WP_068489994_NuoL_Magnetospirillum_marisnigri
MI-------TATVFLPLLGAAVAGLLG----RFIGDRGAQILTCGLLLISAVLAGLIFQTVAL--GHQP-
QTVVIA-DWITSGTFNVAWTLKVDTLTAVMLIVVTWVSAAVHVYSVGYMSHDTS-------
IARFQSYLSLFTFAMLMLVTSDNLVQMFFGWEGVGVASYLLIGFWYEKPSASAAAIKAFVVNRVGDFGFALGIF
GVYFMFGTVGF--DAIFAAAPDKAAAT---
VTFFGVEWHAITICCLLLFVGAMGKSAQLGLHTWLPDAMEGPTPVSALIHAATMVTAGVFMVARMSPLFEFSDT
ALTVVTVVGAATAIFAATVGCVQNDIKRVIAYSTCSQLGYMFFAIGVSAYQAAIFHLMTHAFFKALLFLGAGSV
IHAMSDEQDIRRMGGIWKHVKVTWVLMWIGSLALA--GIPV---------
FAGYYSKDTILEAAWASHSIAGQGAYWLGCTAAFLTAFYSWRLLLLTFHGRPR--ADEHVMAHV----------
---------------------------------------------------
HESPAVMLIPLLFLALGAVFAGWLGYDLFVGH-HNAE---FW--
GKSILVTAAHPSLENAHHVPSWAALLPTVVAVSGIALAYLLYSFA-----------------------------
------------------------------
PGIPGALAKAFQPIYLFLLNKWYFDELYDKIFVKPAFKLGKGLWQG-GDGALIDGVGPDGVA-
AGTLAVARKVGRMQSGYLYHYAYAMLIGVAVFVTWYVFNAR--------
>WP_014745483_NuoL_Tistrella_mobilis
MY-------VALILLPLAAAIVAGFFG----RVIGDRGSQLVTTGAVGLAALLSVVAFFDVAV--GGNA-
TTVNLF-TWIRSGELAFSWALRIDSLTAVMLIVVNGVSFMVHIYSIGYMSHDPH-------
```

```
KPRFMAYLSLFTFAMLTLVTADNLVQMFFGWEGVGLASYLLIGFWYKRPSANKAAMKAFVVNRVGDFGFSLGIF
ALFFVTGSVGF--DAIFADLGTVAGTS---
IDFLGMTLPTIEVIAVLLFIGAMGKSAQLGLHTWLPDAMEGPTPVSALIHAATMVTAGVFMVARLSPVFELTSF
TSDMITVIGASTAFFAATVGLTQTDIKRVIAYSTCSQLGYMFFAAGVGAYPAAIFHLMTHAFFKALLFLGAGSV
IHAMSDEQDMRKMGGIWKLIPVTYTFMWIGSLALA--GLPP---------
FAGFFSKDMILETAYAAHTGVGAYAFWLGIAAAFMTAFYSWRLLFMTFHGKPR--ADKHTMDHV----------
---------------------------------------------------
HESPLVMTAPLFPLAVGAIVAGYIGLPMVDPE-LH-----FW--
NGAITMLGEHNILEEAHHVPGWVKLLPLVMSVGGIALAWFLYIRR-----------------------------
------------------------------
PDLPGKIARDFSGLHKFLLNKWYFDELYDAIFVRPSLWIGRVLWQA-GDRKIIDGLGPDGIA-
AVSVDLARRAGRMQSGYVYHYAFAMLIGLALIVSYF-FGLEA-------
>WP_188574487_NuoL_Tistrella_bauzanensis
MY-------VALILLPLAAAIVAGFFG----RVLGDRGSQFVTTGAVGLAALLSVVAFFDVAV--YGNA-
TTIELF-TWIRSGGLDFAWALRIDALTAVMLIVVNGVSFMVHLYSIGYMSHDPH-------
KPRFMAYLSLFTFAMLTLVTADNLVQMFFGWEGVGLASYLLIGFWYKKPSANKAAMKAFVVNRVGDFGFSLGIF
GVFLVTGSVTF--DAVFGELGNVAGTT---
IDFLGMSLPTIEVLGVLLFIGAMGKSAQLGLHTWLPDAMEGPTPVSALIHAATMVTAGVFMVARLSPLFELTVF
TSDMITVIGASTAFFAATIGLTQTDIKRVIAYSTCSQLGYMFFAAGVGAYPAAIFHLMTHAFFKALLFLGAGSV
IHAMSDEQDMRRMGGIWKLIPVTYMMMWIGSLALA--GIPP---------
FAGFFSKDMILETAYAAHSTVGAYAFWMGIAAAFMTAFYSWRLLFMTFHGKPR--ADKHTMEHV----------
---------------------------------------------------
HESPLIMTVPLFLLAVGAITAGYIGQPMVDAG-LA-----FW--
NGAITMLGEHNILEEAHHVPGWVKLLPLVMSAAGIGLAWLFYIRR-----------------------------
------------------------------
PDLPGKVASDFSGLYKFLLNKWYFDELYDAIFVRPALWIGRVLWQG-GDRRIIDGFGPDGIA-
AVSLGLARRAGRMQSGYVYHYAFAMLIGLALIVSYFFFGLEAGW-----
>WP_074766735_NuoL_Magnetospirillum_fulvum
MTV------IATVFLPLLGAVAAGLFG----RVLGARGSQIVTTGLLMMAAVSAAMVFDRVAL--GGQT-
ETVILA-DWIKSGGLELSWALKVDTLTAVMLVVVTWVSAAVHLYSIGYMAHDPS-------
IPRFQAYLSLFTFAMLMLVTADNLVQLFFGWEGVGLASYLLIGFWYEKPSASAAAIKAFVVNRVGDFGFLLGIF
GLFALFGTIGL--DAIFAAP-AKAGAT---
LTVLGLKINAITVCCLLLFVGAMGKSAQLGLHTWLPDAMEGPTPVSALIHAATMVTAGVFLVARMSPLFELSDA
ALAVVTVVGAATAIFAATIGCVQNDIKRIIAYSTCSQLGYMFFALGVSAYQAAIFHLMTHAFFKALLFLAAGSV
IHAMSDEQDIRRMGGIWKHVKVTWALMWVGSLALA--GIPF---------
FAGYYSKDIILEAAWAAGTPVGQIAYWVGLLAAVLTAFYSWRLLLLTFHGRPR--ADEQVMAHV----------
---------------------------------------------------
HESPAVMIVPLLFLGAGAMFAGWIGYDLFVGP-ASHD---FW--
GHALTVADIHPALDNAHHVPEWVALMPTIAAAGGIATAYLLYVFA-----------------------------
------------------------------
PSLPGMLAGAFPGLHRFLLNKWYFDELYDAILVRPAFRLGRGLWKG-GDGALIDGLGPDGVA-
ATSIRMGRKVAAAETGYLFHYAFAMLIGVAALVTWYMAFKN--------
>WP_046021349_NuoL_Magnetospira_sp._QH-2
MY-------AAIVFLPFLAFLIAGILAFVTPKHRIDRNAQWVTSGSLVISWILSIFAFIDVAL--GGNP-
VTIELF-RWIQSGSMDVSWALKVDTLTAVMLIVVTTVSAMVHIYSIGYMHHDPD-------
VPRFMAYLSFFTFCMLMLVTADNLLQMFFGWEGVGVASYLLIGFWYDKPSANAAAIKAFLVNRVGDFGFALGIF
GTFLLFDTISL--DAIFAAVPGKADTM---
LEVFGMEVHALTALCLLLFVGAMGKSAQLGLHTWLPDAMEGPTPVSALIHAATMVTAGVFMVARLSPMFEFSET
```

```
ALMVVTYVGAFTAFFAATVGLVQNDIKRVIAYSTCSQLGYMFFACGVSAYSAGIFHLMTHASFKALLFLGAGSV
IHAMSDEQDMRRMGGIWKMVPMTYILMWIGSLALA--GVPP---------
FAGFFSKDMILESAWGAHSSVGNLAFWLGVAAAFMTAFYSMRLLMMTFHGKPR--ADETTMAHV----------
---------------------------------------------------
HESPKVMILPLLVLAAGAIGAGYMGYEYFVGH-DAAA---FW--
GNAIYVAEHHQALENAHHAPGWVKLLPLVMGVSGLGLGFLFYGPM-----------------------------
------------------------------
SSMPGWLVTNFRSVYDLLLNKWWFDELYDFLFVRPAFKLGRGFWKG-GDGAVIDGCGPDGIA-
SGVLRWVRRATSMQTGFVYHYAFSMLIGVAGLVTWYLFTLG--------
>WP_007089417_NuoL_Thalassospira_xiamenensis
MY-------PLIVFLPLIAATIAGIIDHHHGVSTSDRISQLITVGGVVTAFIISIFAFVDVAL--GGNP-
VTVQLF-TWISSGNFTAEWALRFDTLTCVMLIVVTGVSSMVHIYSIGYMSHDPD-------
KPRFMAYLSLFTFAMLMLVTADNLIQLFFGWEGVGLASYLLIGFWYSKPSASAAAIKAFVVNRVGDFGFALGIF
AVYVLFDSVQF--DVIFANAEEVAGTT---
LLFLGHEFSALEITCFLLFIGAMGKSAQLGLHTWLPDAMEGPTPVSALIHAATMVTAGVFMVARLSPIFEYAET
TLAIITVVGASTAFFAATIGCVQNDIKRVIAYSTCSQLGYMFFACGVSAYSAGVFHLMTHGFFKALLFLGAGSV
IHAMSDEQDMRKMGGIYKMVPLTFAVMMVGTLAIT--GFPG---------
LAGFYSKDLVIESAFAAHTAVGTYAFWAGILAALMTSFYSWRLIFMTFFGTPR--ADERTMAHV----------
---------------------------------------------------
HESPAVMTLPLLFLTIGAIFAGWFAKDWFGGG-DYEEMLAFW--
NGAIFMAEGHQALEHAHHVPGWVKLAPFVAMVSGFVIALVMYKLV-----------------------------
------------------------------
PSLPRQLANTFNGLYRFALNKWYFDELYDKIFVKPAFYLGYGFWKS-GDGALIDGVGPDGVA-
AACRNIARRVSALQSGFVYHYAFAMLIGIAALVSYTIWKMG--------
>WP_073954579_NuoL_Thalassospira_sp._TSL5-1
MY-------ALIVFLPLIAAIIAGLIDHHHGVSTTDRLSQLVTVAGVVIAAALSVAAFIEVAL--NGNP-
VTLDLF-TWISSGDFTANWSLRFDTLTCVMLIVVTGVSSMVHIYSIGYMSHDPD-------
KPRFMAYLSLFTFAMLMLVTADNLIQLFFGWEGVGLASYLLIGFWYSKPSASSAAMKAFIVNRVGDFGFALGIF
AVYVLFGTVQF--DTIFANAEEISGST---
LVFLGHDFSALNIATFLLFIGAMGKSAQLGLHTWLPDAMEGPTPVSALIHAATMVTAGVFMVARLSPIFEYAET
TLAIVTVVGATTAFFAATIGCVQNDIKRVIAYSTCSQLGYMFFACGVSAYSAGVFHLMTHGFFKALLFLSAGSV
IHAMSDEQDMRKMGGIWKMVPLTFGVMIIGTLAIT--GVPG---------
FAGYYSKDMIIESAFAAHTSVGLYAFWMGVIAAFMTSFYSWRLIFMTFFGAPR--ADERTMAHV----------
---------------------------------------------------
HESPAVMTVPLLILAVGAVFAGWFAVNWFGGN--YEEMLSFW--
NGAIFVAENHQALEHSHHVPEWVKMAPFVAMMLGLALAVLMYRLV-----------------------------
------------------------------
PSLPRMLANTFNGIYRFLLNKWYFDELYDAIFVKPAFALGRGFWKA-GDGALIDGVGPDGVA-
AACRNIARRISTLQSGFVYHYAFAMLIGIAALVSYTIWKMG--------
>WP_189051263_NuoL_Aliidongia_dinghuensis
MA-------TAIVFLPLIAAIIVGLLG----RTLGDRVSQLITCGCVLVAAALSVLVFKDVAL--DHHT-
QTISLL-NWIDTGPLSVDWAIRLDTLTAVMLIVVTGVSSMVHVYSIGYMSHDHS-------
IPRFMAYLSLFTFAMLMLVTADNLVQMFFGWEGVGLASYLLIGFWYDRPSANAAAIKAFVVNRIGDFGFSLGIF
AAFFLFGTLQF--DQIFAAAPGAVGKT---
FHFLSWDVDALTCTSILLFIGAMGKSAQLGLHTWLPDAMEGPTPVSALIHAATMVTAGVFMVCRLSPLLEYSND
ALHFIAIIGAATAIFAASVGMAQNDIKRVIAYSTCSQLGYMVFAAGVSAYPAAMFHLATHASFKALLFLSAGSV
IHAMSGEQDMRKMGGLWKLIPVTYVMMWIGNLALA--GIPP---------
FAGYWSKDTIIEAAYAAGTPYGHFAYWLGIAAAFLTSFYSWRLLFMTFHGKSR--ADHHTLEHV----------
```

```
---------------------------------------------------
HESPMVMVVPLIVLAIGAVFAGWLGNADFVGE-GLKE---FW--
GKSILILDSHPGLAAREEIPPLMSYMALIVGVAGIVLAWVFYVAV-----------------------------
------------------------------
PSLPGAFTSAAKTLYLACLNKWYFDEIYDFLVVKPAMWLGWGLWKG-GDGMIIDGLGPDGVA-
AVTRGLSRRASKLQTGYVYHYAFAMLIGVLCLVSWYLLSQR--------
>WP_088150318_NuoL_Inquilinus_limosus
MD-------IAAIFLPLAGFLIAGLFG----RLIGDRASMAVTTLAVCVAALISWWLFFDVAI--DGNV-
RTHSLL-LWINSGTFVADWALKFDTLTAVMLIVVNTVSAMVHIYSIGYMSHDHS-------
KPRFFAYLSLFTFAMLMLVTADNLIQLFFGWEGVGLASYLLIGFWYEKESARAAAVKAFLVNRVGDFGFALGIF
AVYKIFGTVRY--EEIFAVP-HMVDAR---
INFLGMDLPALTTACLLLFVGAMGKSAQLGLHTWLPDAMEGPTPVSALIHAATMVTAGVFMLARMSPLFEYAPD
ALAVVCVVGALTAFVAATIGLTQFDIKRVIAYSTMSQLGYMFFAIGVSAYQAAIFHLMTHAFFKALLFLGAGSV
IHAMGGEQDMRKMGGIWRLIPTTYALMWIGSLALAGIGIPGV------
FGFAGFYSKDMILEAAWAAHSGVGNLAYWLGVIAALMTAFYSGRLLFLTFHGKPR--ADHHTMEHA--------
-----------------------------------------------------
HESPPVMLVPLIVLAAGAVLAGVIGFGIFVGD-GREG---FW--
REAILVLHGQDSIEAAHHAPEWVSLLPLLAGLVGLGASWLFYIQS-----------------------------
------------------------------
PGLPGAVVRAFKPIHQLFFRKWYFDEIYDALLVKPAFVLGRVFWKA-GDGAIIDRLGPDGVA-
ATSIGIARRASRLESGYVYHYAFVMVIGVVALVSWYVFGGRI-------
>WP_088873694_NuoL_Nitrospirillum_amazonense
MAV------IAAIFLPLLAFIAGFFG----RRIGDLGSQLVTCGAMVAALVLSIILFIDVAV--NGHA--
TVVPLANWVASGDFHVDWALRVDTLTAVMLIVVNGVSTMVHIYSVGYMSHDPD-------
KPRFMSYLSLFTFMMLMLVTSDNLVQMFFGWEGVGVASYLLINFWFEKKSANDASMKAFLVNRVGDFGFALGIF
SIFIIFGSVEF--GTIFPAAAGKAHAT---
FNFLGWHVSALDCACLLLFVGAMGKSAQLGLHTWLPDAMEGPTPVSALIHAATMVTAGVFMLARMSPVFEYAPT
ALAVVTVLGALTAFVAATIGMTQFDIKRVIAYSTMSQLGYMFFAIGVSAYQAAVFHLMTHAFFKALLFLGAGSV
IHAMSGEQDMRKMGGIARLIPFTYVVMWIGNLALAGVGIPGV------
YGFAGFYSKDIVLEAAYGSGTAVGHFAYWLGIAAAFMTAFYSWRLLIMTFHGKPR--ASKEVMDHV--------
-----------------------------------------------------
HESPLVMTGPLTVLAIGALVAGIGGYEWFVGHAARAE---FW--
RGAIQVLHAHDSIHAAHEGPEWVGLLPLLVGLAGIALAYVCYMFV-----------------------------
------------------------------
PGIPATFTKIVKPLHVLVFNKYFFDQIYDVLFVKPAQVLGFGLWKA-GDGAIIDGLGPDGVA-
KTTRGVAAGVSRLQSGYVYHYAFAMVIGVVALVGWFYFKP---------
>WP_184800291_NuoL_Nitrospirillum_iridis
MAV------IAAIFLPLLASFIAGFFG----RRIGDRGSQFVTCAAMLVALGLSIHLFIDVAI--NGHA-
QIVTLA-NWVASGAFHVDWALRVDTLTAVMLIVVNGVSTMVHIYSVGYMSHDPD-------
KPRFMSYLSLFTFMMLMLVTSDNLLQMFFGWEGVGVASYLLINFWFEKKSANDASMKAFLVNRVGDFGFALGIF
TVFVIFGTVEF--KPIFAAA-SQAHAT---
FNFLGWNVSALDCACLLLFVGAMGKSAQLGLHTWLPDAMEGPTPVSALIHAATMVTAGVFMLARMSPVFEYAPT
ALAVVTVVGALTAFVAATIGMTQFDIKRVIAYSTMSQLGYMFFAIGVSAYQAAIFHLMTHAFFKALLFLGAGSV
IHAMSGEQDMRKMGGIWKLIPFTYAVMWIGNLALAGVGIPGV------
YGFAGFYSKDIILEAAYGSGTAVGHFAYWLGIAAAFMTAFYSWRLLILTFHGKPR--ATKDVMDHV--------
-----------------------------------------------------
HESPVVMTGPLTVLALGALVAGIGGYEWFVGHEARAEG--FW-RGAI-
QVLHAHDSIHAAHEGPEWVGVLPLLVGLTGIGLAYVCYMFV---------------------------------
```

```
--------------------------PGIPATFTRVVKPLHTLVFNKYFFDQIYNALFVKPAQLLGYGLWKG-
GDGAIIDGLGPDGVA-KTTRGIAGGVSRLQSGYVVYHYAFAMVIGVVALVGWFYFKP--------
>WP_207478836_NuoL_Desertibacter_SYSUD00532
ME-------VLAIFLPLLGAFIAGFFG----RWIGDRGAQIVTCAGLIVSAMISIWLFNDIAL--GGNP-
RTVELF-TWMDSGALEVSWSLRLDSLTAVMLIVVTVVSSCVHVYSVGYMSHDHS-------
IPRFMAYLSLFTFFMLMLVTADNLLQMFFGWEGVGLASYLLINFWYEKPSANAAAMKAFIVNRVGDFGFALGIM
ATFAIFGTIQF--DAIFGAVPQAANQD---
MNFLGFQAHALTVTCLLLFMGAMGKSAQLGLHTWLPDAMEGPTPVSALIHAATMVTAGVFMVCRLSPMFEYAPA
ALAVVTIVGALTAFVAATIGFTQFDIKRVIAYSTMSQLGYMFFAAGVSAYGAAMFHLMTHAFFKALLFLGAGSV
IHAMSDEQDMRRMGGIWKMIPFTYAMMWVGSLALA--GIPV---------
FAGYYSKDMILEAAYADHSAVGTLAFWLGIAAAFLTAFYSWRLIIMAFHGKPR--ADEKTMAHV----------
---------------------------------------------------
HESPLVMTLPLLVLAIGAVFSGMVAYGWFVGE-GRAE---FW-
KGRAILVLHENDTVEAAHHVPGWVPLAPLVVALAGIATAYVFYMFR----------------------------
-------------------------------
PDLPGALAQRFAGLHRFLYNKWYFDELYDRIFVRPAKALGFGLWQA-GDRAVIDGVGPDGVA-
AATRGMAARAARLQSGYVVYHYAFAMVVGVVVLVGWFYLRG--------
>WP_094456846_NuoL_Niveispirillum_lacus
ME-------IAAIFLPLAGAFIAGFFG----RLIGDRGAQIVTCAGVLIAAVLSVILFIKVGF--
GGPEAARTVELFTWIDSGTFEVSWSLRIDTLTAVMLVVVNVVSSMVHVYSVGYMSHDPH-------
RARFMAYLSLFTFMMLMLVTADNLLQMFFGWEGVGVASYLLINFWYEKKSANDASMKAFLVNRVGDFGFALGIM
AIYMIFGTVEF--DGIFAQAASKADAT---
FNFLGWEVHALTCACLLLFIGAMGKSAQLGLHTWLPDAMEGPTPVSALIHAATMVTAGVFMLARMSPVFEYAPT
ALEVVTYLGAATAFVAATIGLTQFDIKRVIAYSTMSQLGYMFFAIGVGAYPAAIFHLMTHAFFKALLFLGAGSV
IHAMSDEQDMRNMGGIWKLIPGTYTMMWIGNLALA--GIPF---------
FAGFYSKDMILEVAYAKHSAAGSFAFWMGIAAAFMTAFYSWRLLIMTFHGKPR--ASEKVMAHV----------
---------------------------------------------------
HESPAVMLVPLVVLSIGALFAGFVAYPWFVGH-DMNA---
FKAGSIVMLTGEHGSIIEAAHHVPGWVPYAPLVAGVSGIVLAYVFYMFS-------------------------
----------------------------------
PGLPGAFVNTFKALHAFVFRKYMFDELYDALFVRPAMKLGFGLWKG-VDIGVIDGLGPDGAA-
AVTRRLSLASSRVQSGYVVYHYAFVMVIGVVALTGWFILAR--------
>QJE72753_NuoL_Rhodospirillaceae_B3_Rhodocista
ME-------IAAIFLPLLGAFIAGFFG----RQIGDRGAQIVTCAGVLVAAVLSCILFAQIGF--
GGPEAARTVELFTWIDSGSFEVSWALRMDTLTAVMLVVVNVVSAMVHVYSVGYMSHDPG-------
KARFMAYLSLFTFMMLMLVTADNLLQMFFGWEGVGLASYLLINFWYEKTSANNASMKAFLVNRVGDFGFALGIM
AVFVIFGTVQF--DEIFAQAASKADAS---
FNFLGFQVHALTCACLLLFIGAMGKSAQLGLHTWLPDAMEGPTPVSALIHAATMVTAGVFMLSRMSPVFEYAPD
ALAVVTVVGGLTAFLAATIGMTQFDIKRVIAFSTMSQLGYMFFAIGVGAYPAAIFHLMTHAFFKALLFLGAGSV
IHAMSDEQDMRNMGGIWRMIPFTYAMMWIGNLALA--GIIP---------
FAGYYSKDMILEVAYADHSWYGTLAFWLGMAAALMTAFYSWRLLLLTFHGQPR--ANEKVMAHV----------
---------------------------------------------------
HESPLVMTAPLAVLAVGAVFSGLVAYEWFVGH-DMAH---
FWSGSITMLAGEHGTIIDAAHHVPGWVPLAPLAMGILGIALAYVFYMFK-------------------------
----------------------------------
PDLPGKFVNSFRLLHGIVFRKYYFDELYDALFVRPSMKLGYGLWKG-GDQAVIDGFGPDGAA-
SVTRGLSAVTSRLQTGYVVYHYAFVMVIGVVALVGWFYLR---------
>WP_014102898_NuoL_Micavibrio_aeruginosavorus
```

```
ME-------LIAVFAPLLGFVIAFLFG----KQIGDKGAQFVTCTGLIMAAVASCSLFYSVII--QSQP-
HVVSLE-NWITVGDFTVDWALRFDQLSVVMMCVINIVSACVHVYSVGYMSHDPC-------
KARFMSYLSLFTFAMLMLVSADNLVQLFFGWEGVGLASYLLIGFWNHKHSANAAAVKAFVVNRVGDFGLALGIF
TIFMVFGTVQF--QGIFDQAAQHKDTM---
FMFLGYEVHALTLACLLLFMGAVGKSAQLGLHTWLPDAMEGPTPVSALIHAATMVTAGVFLVSRFSPVFEYAPT
ALMVVCIFGAVTALVAATIGLTQFDIKRVIAYSTMSQLGYMFFALGVSAYPAAMFHLTTHAFFKALLFLGAGSV
IHAMSDEQDMRKMGGIWHLIPSTYILMWVGSLALA--GMPF---------
FAGYYSKDMILETAFADHTWFGTMAYWFGIVAALLTAFYSWRLILMTFHGKPR--ADEKVMAHV----------
---------------------------------------------------
HESPLIMLLPLTVLAAGAILAGSVLYGGFVGSPNAAWDKHFW--
GGSIFVLPENDTVEAAHHVPTWVKVLPIFVASAGIILAYIFYMFA-----------------------------
------------------------------
PNMPSVMVKVFKPIHALFYNKWWFDELYDRVFTRNAFRLGRFFWQK-GDKAVIDGYGPDGVA-
SMYARFAGVLSRFQTGYVFQYAFVMMIGLIGLVSWFVFRATMGQ-----
>WP_044426151_NuoL_Skermanella_aerolata
ME-------IAAIFLPLLAFIAGFFG----RLIGDRGSQYITSGAVLLSALLSIWLFVDVAL--HGNA-
RTIELF-TWIDSGTFEVSWALKFDTLTAVMLIVVNVVSSMVHVYSIGYMSHDHH-------
KPRFMAYLSLFTFFMLMLVTADNFVQMYFGWEGVGVASYLLINFWYEKPSANNASMKAFIVNRVGDFGFALGIM
AIFYLFGSVHF--DTVFAAAASKAGQT---
MNFLGFNLDALTVACLLLFMGAMGKSAQLGLHTWLPDAMEGPTPVSALIHAATMVTAGVFMVARLSPIFEYAPI
ALNVIAIVGASTAFVAATIGLTQYDIKRVIAYSTMSQLGYMFFALGVSAYSAAMFHLMTHAFFKALLFLGAGSV
IHAMSDEQDIRKMGGVWRLIPITYAMMWIGSLALA--GIPF---------
FAGYYSKDMILEAAYAAHSGVGAYAFWLGIAAALMTAFYSWRLIILTFHGRPR--ANDRVMAHV----------
---------------------------------------------------
HESPAIMTVPLAVLAVGALFSGMFAYNWFVGE-GREA---FW--
GEAIFVLHEHDTVEAAHHVPSWVPLAPFIAGVIGIALAYLFYMFV-----------------------------
------------------------------
PQLPGIVTNALGGVYRFVYRKWMFDELYDRLFVQPARYLGYGLWKS-GDGAIIDGVGPDGIA-
AATRDVAFRAARLQSGYVYHYAFVMVIGVVLLVGW-FYF-FG-------
>WP_123688416_NuoL_Stella_humosa
MF-------AAAIFLPLLGAIVAGFFG----RWIGDRGAQIVTCAGLLISMLLGWYILYLVGF--RGET-
RTVELF-TWISSGTFEVAWALRFDTLTAVMVFVVTTVSAMVHVYSVGYMSHDHS-------
IPRFMAYLSIFTFFMLMLVTADNFVQMFFGWEGVGVASYLLIGFWFDRQSANAAAMKAFIVNRVGDFGFALGIM
AIFYLFGTVQF--DAVFQAAPQMAGRN---
FEFLGMQVDALTTACILLFIGAMGKSAQLGLHTWLPDAMEGPTPVSALIHAATMVTAGVFMLCRLSPVFEYAPT
ALAIVTLVGATTAIFAATVGLTQFDIKRVIAYSTCSQLGYMFFAIGVSAYQAAIFHLMTHAFFKALLFLGAGSV
IHAMSDEQDMRRMGGIWRLIPTTYTLMWIGSLALA--GLPF---------
FAGFYSKDIILEAAFGAHSGVGQYAFWLGILAALLTAFYSWRLLFLTFHGKPR--ANEKVMAHV----------
---------------------------------------------------
HESPPVMIVPLVVLALGALFSGWLAYEAFVGH-DQKA---FW--
GASIFVASHHDPLEAAHHVPGWVKLLPIFVAVAGIALAWIAYIAR-----------------------------
------------------------------
PDLPRLAAEALRPVHRFFFNKWYFDELYDRIFVRPSFYLGRGLWKT-GDGALIDGVGPDGVA-
AATQAMSIRASRLQTGYVYHYAFAMLIGVVVLVSWYLYVRI--------
>MBT5660202_NuoL_partial_Rhodospirillaceae_SI073_bin103_UBA11136
--------------------------------------------------------------------------
------------------------------------VHVYSIGYMAKDDA-------
IPRFFAYLNLFTFFMLMLVTADNMVQLFFGWEGVGLASYLLIGFWYKRPAANAAAVKAFVVNRVGDFGALGIF
```

AIFLLFGSVNF--GEIFAAAPHHAEAT---
MSFLGFDFHAYTIICLLLFMGAMGKSAQIGLHIWLPDAMEGPTPVSALIHAATMVTAGVFMVVRLSPLFEYAPG
VLNVIAVVGGTTAFFAATIAITQNDIKKVIAYSTCSQLGYMFCAIGVSAYGAAIFHLMTHAFFKSLLFLCAGSV
IHAMSDEQDMRKMGGIWRKVPVTYTMMWIGSLALA--GVPF---------
FAGYYSKDMILEAAWGAHSLPGGYAFLMGTIAAFLTAFYIFRLIFLTFHGEPR--ADEKVMAHI----------
---------------------------------------------------
HESPKSMTVPLVVLAVGAIFAGYLGYDSFVGE-GEKA---FW--
GTSILVLDSHTALIDGHHAPFGIKILPVILGALGILLAYITYFMF-----------------------------
------------------------------
PGLASHAAERFPGLYAFLYNKWYFDELYDRLIVKPALVLGRGLWKS-GDGEVIDGVGPDGIA-
ASVIALGRRIAGIQSGYVYHYAFAMIIGIAGILTCFLYASVG-------
>MBL6935107_NuoL_Alphaproteobacteria_BS150m-G7_UBA11136
LY-------VATIFLPLIGAIAAGLFG----RWVGDRGAQLIACGMMVLSGLFSIIIFRDVAL--DGNE-
RVIELF-TWIRSGGFEASWALKFDTLSAVMVFVVSVVSALIHIYSIGYMAKDPS-------
IPRFFSYLSLFTFFMLMLVSADNLVQLFFGWEGVGLCSYLLIGFWYDRDSANAAAIKAFVLNRIGDFGFALGIF
AVFYLFGSVSF--ETIFAAAPAQAETS---
FEFLGLEVHALTLICLLLFIGAMGKSAQIGLHVWLPDAMEGPTPVSALIHAATMVTAGVFMVVRMSPLFELAPS
VLALVAVIGALTAFMAGTIALAQNDIKRVIAYSTCSQLGYMFFAIGVSAYGAAIFHLMTHAFFKSLLFLGSGSV
IHAMSDEQDMRKMGGIWRKIPLTYAVMWVGSLALV--GIPF---------
FAGYYSKDIIIESAWGAHSLVGNFAYVIGVVVAFLTAFYSCRLLFMTFHGEPR--ADEKVMAHV----------
---------------------------------------------------
HESPKVMTVPLVLLAVGAIFSGSLGYDLFVGD-GGEG---FW--
RASILVLPGHDALAEAHHAPWYIKLTPLVLAVLGMALAFLVYFMF-----------------------------
------------------------------
PGLAGHAAERFPRVHAFLYNKWYFDEIYDRLLVRPALILGRGLWKS-GDGELIDGIGPDGIA-
ASVMRMGRRAAALQSGYVYHYAFAMIIGVVGFISWYLYGLMG-------
>MAG23701_NuoL_Rhodospirillaceae_ARS27_o_UBA1136
LY-------VATIFLPLIGAVVAGLCG----RWTGDRGAQLVTCGLMVLSGLFSIVIFREVAL--DGNE-
RAIELF-TWIRSGGLEAAWALKFDTLTAVMVFVVSVVSALIHIYSIGYMAKDPS-------
IPRFFSYLSLFTFFMLMLVSADNLVQLFFGWEGVGLCSYLLIGFWYDRDSANAAAIKAFVLNRIGDFGFVLGIF
AVFYLFGSVNF--ETIFTAAPDQAETT---
FEFLGLEVHALTLICLLLFVGAMGKSAQLGLHVWLPDAMEGPTPVSALIHAATMVTAGVFMVVRMSPLFELAPL
VLALVAVVGALTAFMTATIALAQNDIKRVIAYSTCSQLGYMFFAIGVSAYGAAIFHLMTHAFFKSLLFLGSGSV
IHAMSDEQDMRKMGGLWRRIPLTYTVMWVGSLALA--GIPF---------
FAGYYSKDIIIESAWAAHGVVGNFTYILSVVVAFLTAFYSYRLLFMTFHGESR--ADERVMAHV----------
---------------------------------------------------
HESPKVMTVPLVLLSVGAIFSGYLGYDLFVGD-GGEE---FW--
RASILVLPGHDALAEAHHVHWYIKIVPLVMAGLGILLAFVVYFMF-----------------------------
------------------------------
PGLAGHAAERFPRLHTFLYNKWYFDEVYDRLLVRPALAFGRALWKT-GDGELIDGIGPDGIA-
ASVMRMGRRAAALQSGYVYHYAFAMIIGVVGFISWYLYELAG-------
>MDP7045168_NuoL_Alphaproteobacteria_Arabian9_MAG_19
LY-------VATIFLPLIGAVVAGLCG----RWTGDRGAQLVTCGLMVLSGLFSIVIFREVAL--DGNE-
RAIELF-TWIRSGGLEAAWALKFDTLTAVMVFVVSVVSALIHIYSIGYMAKDPS-------
IPRFFSYLSLFTFFMLMLVSADNLVQLFFGWEGVGLCSYLLIGFWYDRDSANAAAIKAFVLNRIGDFGFVLGIF
AVFYLFGSVNF--ETIFTAAPDQAETT---
FEFLGLEVHALTLICLLLFVGAMGKSAQLGLHVWLPDAMEGPTPVSALIHAATMVTAGVFMVVRMSPLFELAPL
VLALVAVVGALTAFMTATIALAQNDIKRVIAYSTCSQLGYMFFAIGVSAYGAAIFHLMTHAFFKSLLFLGSGSV

IHAMSDEQDMRKMGGLWRRIPLTYTVMWVGSLALA--GIPF---------
FAGYYSKDIIIESAWAAHGVVGNFTYILSVVVAFLTAFYSRLLFMTFHGESR--ADERVMAHV-----------
---------------------------------------------------
HEAPKVMTVPLVLLSVGAIFSGYLGYDLFVGD-GGEE---FW--
RASILVLPGHDALAEAHHVPRYIKIVPLVMAGLGILLAFVVYFMF-----------------------------
------------------------------
PGLAGHAAERFPRLHTFLYNKWYFDEVYDRLLVRPALAFGRALWKT-GDGELIDGIGPDGIA-
ASVMRMGRRAAALQSGYVYHYAFAMIIGVVGFISWYLYELAG-------
>MDP7122910_NuoL_Alphaproteobacteria_ETNP8_MAG_36
LY-------VATIFLPLIGAVVAGLCG----RWTGDRGAQLITCGLMVLSGLFSIVIFREVAL--DGNE-
RVIELF-TWIRSGGLEAAWALKFDTLTAVMVFVVSVVSALIHIYSIGYMAKDPS-------
IPRFFSYLSLFTFFMLMLVSADNLVQLFFGWEGVGLCSYLLIGFWHDRDSANAAAIKAFVLNRIGDFGFVLGIF
AVFYLFGSVNF--ETIFTAAPDQAETT---
FEFLGLEVHALTLICLLLFVGAMGKSAQLGLHVWLPDAMEGPTPVSALIHAATMVTAGVFMVVRMSPLFELAPL
VLALVAVVGALTAFMTATIALAQNDIKRVIAYSTCSQLGYMFFAIGVSAYGAAIFHLMTHAFFKSLLFLGSGSV
IHAMSDEQDMRKMGGLWRRIPLTYTVMWVGSLALA--GIPF---------
FAGYYSKDIIIESAWAAHGVVGNFTYILGVVVAFLTAFYSCRLLFMTFHGESR--ADERVMAHV----------
---------------------------------------------------
HESPKVMTVPLVLLSVGAIFSGYLGYDLFVGD-GGEE---FW--
RASILVLPGHDALAEVRYAPQYIKIAPLAMAGLGILLAFVVYFMF-----------------------------
------------------------------
PGLAGHAAERFPRLHTFLYNKWYFDEVYDRLLVRPALAFGRALWKS-GDGELIDGIGPDGIA-
ASVMSMGRRAAALQSGYVYHYAFAMIIGVVGFISWYLYELAG-------
>MBV23930_NuoL_Rhodospirillaceae_NP1051_UBA11136
LY-------VATIFLPLIGAVVAGLCG----RWTGDRGAQLITCGLMVLSGLFSIVIFREVAL--DGNE-
RAIELF-TWIRSGGLEAAWALKFDTLTAVMVFVVSVVSALIHIYSIGYMAKDPS-------
IPRFFSYLSLFTFFMLMLVSADNLVQLFFGWEGVGLCSYLLIGFWHDRDSANAAAIKAFVLNRIGDFGFVLGIF
AVFYLFGSVNF--ETIFTAAPDQAETT---
FEFLGLEVHALTLICLLLFIGAMGKSAQLGLHVWLPDAMEGPTPVSALIHAATMVTAGVFMVVRMSPLFELAPL
VLALVAVVGALTAFMTATIALAQNDIKRVIAYSTCSQLGYMFFAIGVSAYGAAIFHLMTHAFFKSLLFLGSGSV
IHAMSDEQDMRKMGGLWRRIPLTYXVMWVGSLALA--GIPF---------
FAGYYSKDIIIESAWAAHGVVGNFTYILGVVVAFLTAFYSCRLLFMTFHGESR--ADERVMAHV----------
---------------------------------------------------
HESPKVMTVPLVLLSVGAIFSGYLGYDLFVGD-GGEE---FW--
RASILVLPGHDALAEVRYAPQYIKIAPLAMAGLGILLAFVVYFMF-----------------------------
------------------------------
PGLAGHAAERFPRLHTFLYNKWYFDEVYDRLLVRPALAFGRALWKS-GDGELIDGIGPDGIA-
ASVMRMGRRAAALQSGYVYHYAFAMIIGVVGFISWYLYELAG-------
>Ga0066848_100310331_ETNP2013_S10_300m_22_NuoL_500aa
--------------------------------------------------------------------------
----------------------------------ALIHIYSIGYMAKDPS-------
IPRFFSYLSLFTFFMLMLVSADNLVQLFFGWEGVGLCSYLLIGFWHDRDSANAAAIKAFVLNRIGDFGFVLGIF
AVFYLFGSVNF--ETIFAVAPDQAETT---
FEFLGFEVHALTLICLLLFIGAMGKSAQLGLHVWLPDAMEGPTPVSALIHAATMVTAGVFMVVRMSPLFELAPL
VLALVAVVGALTAFMTATIALAQNDIKRVIAYSTCSQLGYMFFAIGVSAYGAAIFHLMTHAFFKSLLFLGSGSV
IHAMSDEQDMRKMGGLWRRIPLTYTVMWVGSLALA--GIPF---------
FAGYYSKDIIIESAWAAHGVVGNFTYILGVVVAFLTAFYSCRLLFMTFHGESR--ADERVMAHV----------
---------------------------------------------------

```
HESPKVMTVPLVLLSVGAIFSGYLGYDLFVGD-GGEE---FW--
RASILVLPGHDALAEAHHVPRYIKIVPLVMAGLGVLLAFVVYFMF-----------------------------
------------------------------
PGLAGHAAERFPRLHTFLYNKWYFDEVYDRLLVRPALAFGRALWKS-GDGELID--------------------
--------------------------------------
>MDP6780196_NuoL_Alphaproteobacteria_Arabian10_MAG_26
LY-------VATIFLPLIGAVVAGLCG----RWTGDRGAQLVTCGLMVLSGLFSIVIFREVAL--DGNE-
RAIELF-TWIRSGGLEAAWALKFDTLTAVMVFVVSVVSALIHIYSIGYMAKDPS-------
IPRFFSYLSLFTFFMLMLVSADNLVQLFFGWEGVGLCSYLLIGFWYDRDSANAAAIKAFVLNRIGDFGFVLGIF
AVFYLFGSVNF--ETIFTAAPDQAETT---
FEFLGLEVHALTLICLLLFVGAMGKSAQLGLHVWLPDAMEGPTPVSALIHAATMVTAGVFMVVRMSPLFELAPL
VLALVAVVGALTAFMTATIALAQNDIKRVIAYSTCSQLGYMFFAIGVSAYGAAIFHLMTHAFFKSLLFLGSGSV
IHAMSDEQDMRKMGGLWRRIPLTYTVMWVGSLALA--GIPF---------
FAGYYSKDIIIESAWAAHGVVGNFTYILGVVVAFLTAFYSRLLFMTFHGESR--ADERVMAHV-----------
---------------------------------------------------
HEAPKVMTVPLVLLSVGAIFSGYLGYDLFVGD-GGEE---FW--
RASILVLPGHDALAEAHHVHWYIKIVPLVMAGLGILLAFVVYFMF-----------------------------
------------------------------
PGLAGHAAERFPRLHTFLYNKWYFDEVYDRLLVRPALAFGRALWKT-GDGELIDGIGPDGIA-
ASVMRMGRRAAALQSGYVYHYAFAMIIGVVGFISWYLYELAG-------
>MDP6660958_NuoL_Alphaproteobacteria_Arabian11_MAG_35
LY-------VATIFLPLIGAVVAGLCG----RWTGDRGAQLVTCGLMVLSGLFSIVIFREVAL--DGNE-
RAIELF-TWIRSGGLEAAWALKFDTLTAVMVFVVSVVSALIHIYSIGYMAKDPS-------
IPRFFSYLSLFTFFMLMLVSADNLVQLFFGWEGVGLCSYLLIGFWYDRDSANAAAIKAFVLNRIGDFGFVLGIF
AVFYLFGSVNF--ETIFTAAPDQAETT---
FEFLGLEVHALTLICLLLFVGAMGKSAQLGLHVWLPDAMEGPTPVSALIHAATMVTAGVFMVVRMSPLFELAPL
VLALVAVVGALTAFMTATIALAQNDIKRVIAYSTCSQLGYMFFAIGVSAYGAAIFHLMTHAFFKSLLFLGSGSV
IHAMSDEQDMRKMGGLWRRIPLTYTVMWVGSLALA--GIPF---------
FAGYYSKDIIIESAWAAHGVVGNFTYILSVVAAFLTAFYSRLLFMTFHGESR--ADERVMAHV-----------
---------------------------------------------------
HEAPKVMTVPLVLLSVGAIFSGYLGYDLFVGD-GGEE---FW--
RASILVLPGHDALAEAHHVHWYIKIVPLVMAGLGILLAFVVYFMF-----------------------------
------------------------------
PGLAGHAAERFPRLHTFLYNKWYFDEVYDRLLVRPALAFGRALWKT-GDGELIDGIGPDGIA-
ASVMRMGRRAAALQSGYVYHYAFAMIIGVVGFISWYLYELAG-------
>MDP7669054_NuoL_Alphaproteobacteria_ETNP2_MAG_74
LY-------VATIFLPLIGAVVAGLCG----RWTGDRGAQLITCGLMVLSGLFSIVIFREVAL--DGNE-
RAIELF-TWIRSGGLEAAWALKFDTLTAVMVFVVSVVSALIHIYSIGYMAKDPS-------
IPRFFSYLSLFTFFMLMLVSADNLVQLFFGWEGVGLCSYLLIGFWHDRDSANAAAIKAFVLNRIGDFGFVLGIF
AVFYLFGSVNF--ETIFTAAPDQAETT---
FEFLGLEVHALTLICLLLFIGAMGKSAQLGLHVWLPDAMEGPTPVSALIHAATMVTAGVFMVVRMSPLFELAPL
VLALVAVVGALTAFMTATIALAQNDIKRVIAYSTCSQLGYMFFAIGVSAYGAAIFHLMTHAFFKSLLFLGSGSV
IHAMSDEQDMRKMGGLWRRIPLTYTVMWVGSLALA--GIPF---------
FAGYYSKDIIIESAWAAHGVVGNFTYILGVVVAFLTAFYSCRLLFMTFHGESR--ADERVMAHV----------
---------------------------------------------------
HESPKVMTVPLVLLSVGAIFSGYLGYDLFVGD-GGEE---FW--
RASILVLPGHDALAEVRHAPWYIKIAPLVMAGLGILLAFVVYFMF-----------------------------
------------------------------
```

```
PGLAGHAAERFPRLHTFLYNKWYFDEVYDRLLVRPALAFGRALWKS-GDGELIDGIGPDGIA-
ASVMSMGRRAAALQSGYVYHYAFAMIIGVVGFISWYLYELAG-------
>MDP7190726_NuoL_Alphaproteobacteria_ETNP7_MAG_8
LY-------VATIFLPLIGAVAVGLLG----RWTGDRGAQLITCGLMVLSGLFSLVIFREVAL--DGNA-
RTIELF-TWIHSGEFEVSWALKFDTLTAVMVFVVSVVSALIHIYSIGYMAKDPS-------
IPRFFSYLNLFTFFMLILVTANNLIQLFFGWEGVGLCSYLLIGFWYDRASANTAALKAFVVNRIGDFGFALGIL
AVFYLFSSVAF--ETIFTAAPAQAGTT---
FEFLGLEVHALTLICLLLLIGAMGKSAQLGLHVWLPDAMEGPTPVSALIHAATMVTAGVFMVVRLSPLFELAPS
VLAAAAIIGALTAFITATIALVQNDIKRVIAYSTCSQLGYMFFAVGVSAYGAAIFHLMTHAFFKSLLFLGSGSV
IHAMSGERDMRQMGGIWRRIPLTYVAMCVGSLSLI--GIPF---------
FAGYYSKDIILESAWEAHGFAGNFAYILGVVVAFLTAFYSYRMLFMTFHGDSR--AGEKVMAHV----------
---------------------------------------------------
HESPKVMTVPLIILSVGAIFSGYLGYDLFVGD-GGEG---FW--
RASILALPSHDALADTHYASWYIQITPYIQITLGILLAFITYSKF-----------------------------
------------------------------
PGLADSAAERFPRLHAFLYNKWYFDEVYDRLFVRPTLVLARGLWKS-GDGRLIDGIGPDGIA-
ASVLRVGRRAAALQSGYVYHYAFAMIIGVVGFISWHLYGSMG-------
>MBT5194942_NuoL_Rhodospirillaceae_SI074_bin4_oxycline_NosZ
MY-------HAIVFLPLIAFLIVGTLG----RVLGDRPSQWITCGAVIISAVLSLVAFWQVAL--GGQP-
VKVEVA-SWIVSGTFDVSWSLRIDQLTAVMLVVVNGVSALVHVYSVGYMSHDPH-------
KPRFMAYLSLFTFAMLMLITSDNFLQMFFGWEGVGLASYLLIGFWYHKPAANAAAIKAFLVNRIGDFGFGLGIM
ATFMVFGSVDF--DTVFAASPDQVGKT---
FHFLSWEWDVMTTICILLFIGAMGKSAQVPLHTWLPDAMEGPTPVSALIHAATMVTAGVFLVARCSPMFEFSPT
ALVVVTIVGGTTAFFAATVGLVQNDIKRVIAYSTCSQLGYMFFALGVGAYPVAIFHLFTHAFFKALLFLGSGSV
IHAMSDEQDMRRMGGIYKYIPQTYIMMWIGSLALA--GLPI---------
FAGFYSKDAILESAFAAHTGAGQYAFWLGLAAAVMTAFYSWRLLFMTFHGQPR--ASQEVMSHV----------
---------------------------------------------------
HESPWSMLVPLYFLAAGSILSGILFVHYFVGD-GAGA---FW--
GNSIFMLPDNNILEAMHHVPTWVKVAPIVAGLFGILVAWQFYIRR-----------------------------
------------------------------
TDIPARLAKVHHEAYLFLLNKWYFDELYDLTIVNPAKWLGRLLWKG-GDGRIIDGFGPDGIA-
ATVMRVARRASALQSGYLYHYAFAMLIGVAVLVTLFYTSTGGGH-----
>WP_108680428_NuoL_Methyloceanibacter_sp._wino2
MY-------HAILFLPLIGALFAGLFG----RFVGYRACEIVTISLMFVAAFLSWIAFYQVAI--QGED-
VRISLI-GFMHSGAFETYWSIRIDTLTAVMLVVVNTVSSVVHLYSVGYMDEDPH-------
KERFFAYLSLFTFAMLALVTSDDLVQLFFGWEGVGLASYLLIGFWYKKPSANAAAIKAFVVNRVGDFGFLLGIF
TIFYVFGAVQY--DTIFAAAPDQVGKT---
ITFLSWEVPALELMCFLLFMGAMGKSAQFLLHTWLPDAMEGPTPVSALIHAATMVTAGVFMVARMSPLFDLAPY
ALAFVAFIGATSAFFAATAAVVQTDIKRVIAYSTCSQLGYMFAAEGVGAYGAGIFHLFTHACFKALLFLCAGAV
IHTLADNQDMRRMGGLRKIIPFTWVMMLIGSLALT--AFPL---------
TSGFVSKDFIIETTWFDHTAVGQYTYWMVLIAAFLTSLYTWRLMFMTFEGNFR--GPKDVKAHA----------
---------------------------------------------------
HEPPWTMGVPLMVLAAGALFAGLGFRHLFIGA-DQAS---FW---NRDSIVLPH---
GGHHDVPFWLVLSPTIAMTLGFLIAWYCYMWR------------------------------------------
-----------------PLVPFGLLKRFPATNQFFFHAWYFDALYDRIFVRPAKWLGNFLWKV-
GDGKIIDGLGPNGVA-ARVLDITRGVVKLQTGYIYHYAFIMLIGVALLITYFIVTGGAKP-----
>PJK29752_NuoL_Minwuia_thermotolerans
MY-------SLIVFAPLLGAILAGFFG----RMIGDRGSMLVTSGLMTLAAVLSWIVFLDVTF--GHAP-
```

THVELL-TWITAGEFRVNWALQVDQLTAVMLIVVNTVACLVHWYSIGYMAHDPH-------
KARFFSYLSLFTFAMLMLVTSDNFVQLYFGWEGVGLASYLLIGFWYHKPSANAAAMKAFIVNRVGDFGFALGIA
GIFLVFEDVTF--AGVFAATPEVAGQT---
FQFLGYEVDILTTICLLLFMGAMGKSAQLPLHTWLPDAMEGPTPVSALIHAATMVTAGVFLVVRCSHMFEYSPD
ALAVVTVVGASTAFFAATIGLVQNDIKRVIAYSTCSQLGYMFFAAGLSAYPVAIFHLFTHAFFKALLFLGAGSV
IHAVADEQDMRKMGGLARHIKKTYIMMWIGSLALAGIGIPG------
FAYVFGFYSKDLIVESAFGAGSGVGQFAYIMGVGAAIMTAFYSWRLLFMTFHGKPR--APRETMAHI-------
------------------------------------------------------
HDSPPVMMIPLYILAVGAVFSGLLFIGPLTGH-HWHD---FW--
GDSILILPQHGAMEAAHEVPLWVKLSPLVASLVGILIAWVMYVRK-----------------------------
------------------------------
TDLPGNFANTHRPLYLFLLNKWYFDELYDWIFVRPAKALGRILWKG-GDGRIIDGLGPDGIS-
ASVLRATRRITMLQTGYVVYHYAFAMLIGIAVLVTFYFYAIGV-------
>WP_090071583_NuoL_Cohaesibacter_marisflavi
MY-------SAIVFLPLIGFLIAGLFG----RTLGHKASEIITSTLLVIAAFLSWVAFLSVGI--
GHGESQTVNIL-TWLSSGDLEINWSIRVDTLTVVMLVVVNTISALVHIYSIGYMHGDPH-------
RARFFAYLSLFTFAMLSLVTADNLLQMFFGWEGVGLASYLLIGFWYQKPSANAAAMKAFIVNRVGDFGFLLGLC
GIYVLFGSISF--DTIFANASAMQEQT---
IHFLGQDFNALTTICLLLFMGAMGKSAQFLLHTWLPDAMEGPTPVSALIHAATMVTAGVFMVARLSPLFELSPT
ALEVVTFIGATTAFFAATVGLVQNDIKRVIAYSTCSQLGYMFVALGIGAYGAAVFHLFTHAFFKALLFLGAGSV
IHAVSDEQDMRRMGGLRKHIKLTYLMMLVGTFALTGVGIPGT-----
ILGFAGFNSKDAIIESAFAAQNGMANYAFAMTVIAALFTSFYSWRLIFLTFHGRER--MSADVKAHI-------
------------------------------------------------------
HESPAVMTVPLIVLALGAVLAGMVFAGYFYGH-HYEE---FW--
KGALFTGAENHVMHESHDVPMMVKLAPFVMMAVGFLVAFGFYILS-----------------------------
------------------------------
PSTPKKLAERHNWLYKFLLNKWYFDELYNFLFIRGAKGLGSLLWKG-GDEGVIDRFGPNGIA-
ARVVALTSRINRLQTGYVVYHYAFAMMIGVAILITYSMLS-GGAH-----
>SDF10219_NuoL_Thalassobaculum_litoreum
MY-------AAIVFLPLIGALIAG-FGN---KKLGDRGAQIVTCGAMLTSAVLGIVAFWEVAL--QGNA-
QTVELF-TWIDSGSFEASWALRWDTLTAVMVIVVTVVSSCVHVYSVGYMSHDPS-------
IPRFMAYLSFFTFAMLMLVTSDNLIQMFFGWEGVGVASYLLIGFWYDRPSANAAAIKAFVVNRVGDFGFALGIF
GCFLLFDAVSF--DAIFAAAPEMAGTT---
FGFLWWEEVDALTVAILLFIGAMGKSAQLGLHTWLPDAMEGPTPVSALIHAATMVTAGVFLVARMSPLFEYAPT
ALVVVTVVGAATAFFAATVGTTQNDIKRVIAYSTCSQLGYMFFALGVSAYPAAIFHLMTHAFFKALLFLSAGSV
IHALHDEQDMRNMGGIWRKIPWTYAMFWIGSLALA--GIPF---------
FAGYYSKDMILESAFAAHTGAGQFAFWAGIVAALLTAFYSWRLLFMTFHGKPR--MDQHTFDHA----------
---------------------------------------------------
HESPPVMLVPLIVLAIGAVFSGFIGYEYFVGH-EMDA---FW--
GSAIKVLEENNVIEAAHHSPTWVKLLPLVVGVIGIGAAYVAYILN-----------------------------
------------------------------
PGIPAKVTSAIRPVYLFLLNKWYFDELYDWLFVKRSWQAGLMFWKT-GDGTIINGFGPDGVS-
AMTRWCAGQVARLQTGYLYHYAFAMLIGVVALVSWYLAF-GG-------
>WP_090018703_NuoL_Limimonas_halophila
ME-------VVAVFLPLIGAALAGLFG----GVLGDRGSQLVTCGLLIVSAVLSVIVFVDVAL--YDNA-
RTTELF-TWFASGDLELSWAIRMDTLSAVMLATVTVISALIHVYSIGYMEHDAS-------
IQRFFSYISLFTFFMLMLVTADNFLQLFFGWEGVGLASYLLIGFWYQKPSANAAAIKAFLVNRVGDIGFALGIA
ATFFVFQTTSF--DQVFAAAPEMADTG---

```
FSFLGIEAPALTIISILLFIGAMGKSAQLGLHTWLPDAMEGPTPVSALIHAATMVTAGVFMLARVSPILEQAPT
ALAVITIIGALTAFFAASVGMVQNDLKRVIAYSTCSQLGYMMFAVGVSAYGAAIFHLMTHAFFKALLFMGAGSV
LHAMDEEHDMRKMGGIWRMLPITYAVMWIGSLALA--GIPP---------
FAGFFSKDMILEAAYAAHSGTGQLAFWLGVIAAGMTAFYSWRLLFMTFHGKPR--ASAETMKHV----------
---------------------------------------------------
HESPKVMTLPLMALAVGAVLAGWLGYSAFVGH-GFEH---FW--SGSIL---
GHEAIKAAHHVPLWVKLLPLVVSVGGIAVAYVMYIAR-------------------------------------
----------------------PELPAALASRLRPVHAFLYNKWYFDELYDLVFVRPAWALGNGLWRA-
GDIAVIDGLGPNGVS-SVARGLARRASQLQTGYVYHYAFAMLIGVVALVTWYMMTQIG-------
>WP_027135330_NuoL_Geminicoccus_roseus
MY-------TILVFAPLVGALIAGLFG----RLIGDRASQVVTCGFMAISAICAISSFFLNI---YGEP-
FKVVLF-DWIVVGSFDTQWALRIDGLSATMMLVVSFISFLIHVYSVGYMSHDAS-------
IPRFMSYLSLFTFAMLMLVTSDNLLQLFFGWEGVGVASYLLIGFWYEKPSACAAAMKAFIVNRVGDFGLILGLA
GCYLVFDSIQY--DVIFPQVAEFADSS---
IVLFSYEFNTLTIIGVLLFIGAMGKSAQLGLHTWLPDAMEGPTPVSALIHAATMVTAGVFLVARFSPLYEYAPT
ALAMVTLVGASTAFFAATIGTVQNDIKRVIAYSTCSQLGYMFFACGVSAYSAGVFHLFTHAFFKALLFLSAGSV
IHAMSDEQDMRKMGGIWRKIPYTYAMMWIGSLALM--GVPF---------
FAGYYSKDAILEAAYASHAPFAGYAFWLGLAAAFLTAFYSGRLLWMTFHGKPR--ADHHTMEHV----------
---------------------------------------------------
HESPWVMLVPLFCLAAGAVFAGWLAKGALDPD-AA-----WW--
GGAIFMAHEPNIMEELHHVPALVKFAPLIVGIAGFGLSWVFYLMM-----------------------------
------------------------------
PGIPGRLASAARPVYLFLLNKWYFDELYDRIFVRPSKMIGSALWRG-GDVFVIDGFGPDGIA-
ATTAQLAKQTARLQTGYVYHYAFAMLIGVAALVTWYLFFVS--------
>WP_088559410_NuoL_Arboricoccus_pini
MLH------TILVLCPALGALIVGLFG----RYLGDRASQIITCGLMAITALCAWIGVFTYM---GEPA-
FKVHWF-TWISVGGLEADWSLRIDSLSTIMMLVVGFISFLIHVYSVGYMSHDPS-------
KPRFMAYLSLFTFAMLMLVTSDNIIQLFFGWEGVGVASYLLIGFWYTRPSACAAAMKAFIVNRVGDFALMLGIG
ALFLVFDSVSY--DVIFNKAASLADTQ---
IVLFGGTFQTLNIITLLLFIGAMGKSAQIGLHTWLPDAMEGPTPVSALIHAATMVTAGVFLVARFSPVFEFAPS
TLAFVTFIGATTCIFAATVGMTQFDIKRVIAYSTCSQLGYMFFACGVGAYQAGVFHLMTHAFFKALLFLGAGSV
IHAMSDEQDMRKMGGIWKKIPLTYAMMWIGTLALIGFGIPGIG------
GFAGFYSKDAILEAAYASNGSFHLYAFWMGVLAAVLTAFYSGRLIFLSFHGEPR--ADHHTMEHV---------
----------------------------------------------------
HESPPIMTVPLMVLGVGAVIAGFVAFGMVEPE-GN-----WW--
GGAIFTGPENHVLHAMHEVPLWVSLAPFFAMAFGLGLSWVFYIAM-----------------------------
------------------------------
PGTAPKIAAAVRPLDNFLRNKWYFDELYDRIFVRPALALGRGLWHG-GDQGIIDRFGPDGVA-
GTTWGTALRIARLQTGYVYHYAFVMLIGVAVFVSWYLFLQR--------
>TVQ34661_NuoL_Geminicoccaceae_bacterium
MY-------TAIVFLPLLGAIVAGLFG----RIIGDRAAQIVTCSLLCISALLSGIALLTVPF--GEP--
FKVPLA-TWIVSGSFEVDWALRIDALTVVMLVVVTWVSALVHIYSIGYMSHDAS-------
IARFMSYLSLFTFAMLMLVTADNFLQLFFGWEGVGVASYLLIGFWYEKASANNAAMKAFIVNRVGDFALILGIA
AIFLVFDSVDF--DTVFAAAPAFAEHQ---
IALFGVTWSTLDLICILLFIGAMGKSAQLGLHTWLPDAMEGPTPVSALIHAATMVTAGVFLIARVSPIIEYAPV
ALAVIVLIGATTAFFAATVALTQNDIKRVIAYSTCSQLGYMFFALGVGAYAAAIFHLFTHAFFKALLFLAAGSV
IHGMSNEQDMRRMGGIWRHMKFTYAMMWIGSLALV--GVPF---------FAGFYSKDLILEAAFAA-
GGVGFYAFALGIAAAVMTAFYSGRLLFMTFHGKPR--ASEEVMHHV----------------------------
```

```
---------------------------------HESPLVMTIPLGLLAVGAVFAGYLGLPMVADD-HA-----
FW--GGSIYLEMERNIIYLAHYVPTWVVIAPIVAGFVGLGLAYVIYIQR-------------------------
----------------------------------
TELADRLAERFRPLYQFSLNKWYFDELYDALFVRPARQAGTFLWRR-VDAGIIDDYGPNGVA-
ATALEGTKRVVKLQTGYIYHYAFVMLIGVAALVSWFLFVARG-------
>WP_138380149_NuoL_Luteithermobacter_gelatinilyticus
MY-------QAIVFLPLLGAIIAGFFG----KQIGDRGAQIVTCSFLVISALLSWIAFGQVAL--GHQE-
AYVTVL-NWISSGDLQLDWAFRIDTLTAVMLVVVTTVSALVHIYSVGYMSHDPH-------
IPRFMSYLSLFTFAMLMLVTADNFLQLFFGWEGVGLASYLLIGFWYKKPSANAAAIKAFVVNRVGDFGFALGIV
AIFMAFGSLDF--TTVFSSVPDYTDNV---
IYFLGMELDLITTISILLFIGAMGKSAQLGLHTWLPDAMEGPTPVSALIHAATMVTAGVFMVARCSPIFEYSPD
ALAFVTFIGASTAIFAATIGVAQNDIKRVIAYSTCSQLGYMFFALGVSAYGAAIFHLFTHAFFKALLFLGSGSV
IHAMSDEQDMRKMGGLWKKIPITFAMMTIGTLALT--GFPF---------
FAGYYSKDMIIESAFASSASMSDYAFAMGVLAALFTSFYSWRLVFMTFFGESR--ASKKVQDHV----------
---------------------------------------------------
HESPQVMLIPLYLLAVGAIASGFVFSEYFTGH-HYED---FW--GTSLKV-
LGDNVMEAAHHVPAWVVWSPFIAMVTGFAVAFYSYILA------------------------------------
-----------------------PETPAAAARTFKYLYNLFLNKWYVDEIYDFLFVRPAKRLGAFLWKK-
GDLGTIDRYGPDGVS-ATVVALARKFREMQTGYLYHYAFAMLIGVAAFTTWFIV--GGN------
>WP_139938140_NuoL_Emcibacter_nanhaiensis
MY-------QTIVFLPLVMSFIAGMFG----NRIGDRGAQIITCGGLIVSAILSWVAFFDIAL--GHNV-
AHVEVL-SWIVSGDFNLSWAFKVDTLTSVMLVVVTTVSCVVHIYSVGYMSHDPD-------
IPRFVSYLSLFTFAMLMLVTADNFLQMFFGWEGVGLASYLLIGFWYKKPSACAAAIKAFVVNRVGDFGFALGIF
AIFMTFGTLDF--ATVFASVPDVAGQK---
IHFLNWELDLITTICILLFIGAMGKSAQLGLHTWLPDAMEGPTPVSALIHAATMVTAGVFMVARCSPIFEYSPD
ALAVVCVIGASTAIFAATIGVAQNDIKRVIAYSTCSQLGYMFFAIGVSAYGAAIFHLFTHAFFKALLFLGSGSV
IHAMSDEQDMRKMGGLWKKIPVTFGLMTIGTLALT--GFPF---------
FAGYYSKDMIIESAFASHSPVGPYAFAMGVVAALFTSFYSWRLVFMTFFGESR--ASKKVQDHV----------
---------------------------------------------------
HESPQVMLVPLYILAVGAVFAGWVFHEYFTGH-HYEE---FW--GHALKL-
VGDNVMHEAHHVPGWVVWSPFVAMVTGFAVAWISYIYA------------------------------------
-----------------------PWIPGAAASTFKYLYNLFLNKWYVDEIYDFLFVRPAKRLGVFLWKR-
GDEQTIDRYGPDGVS-ASVAALARKFKEMQTGYLYHYAFVMLIGVAAITTWFIV--GGN------
>WP_099472212_NuoL_Paremcibacter_congregatus
MY-------QIIVFGPLLAAIIAGLFG----RKLGDKGAQAITCGALILSAILSWVAFFQIGL--GHTV-
EHVTVM-NWVTSGDLSFNWAFKVDTLTAVMLVVVNTVSCLVHIYSVGYMSHDPH-------
KPRFMAYLSLFTFAMLMLVTADNFVQMFFGWEGVGLASYLLIGFWYKKPSACAAAIKAFLVNRVGDFGFALGIL
LVFMATGSVDF--ETVFAKIPELQGQQ---
IHFLWMELDLITTMCILLFIGAMGKSAQLGLHTWLPDAMEGPTPVSALIHAATMVTAGVFMVARCSPIFEFSPD
ALAFVTLVGASTAIFAATIGTTQNDIKRVIAYSTCSQLGYMFFALGVSAYGAAIFHLFTHAFFKALLFLGSGSV
IHAMSDEQDMRKMGGLWNKIPITFAMMTIGTLALT--GFPF---------
LAGFYSKDMIIESAFAANNPWSDYAFFMGVFAALLTSFYSWRLVFMTFFGKSR--ASQSVQDHV----------
---------------------------------------------------
HESTNWMLVPLYVLAAGALAAGFVFKENFVGH-HYED---FW--
GQALFLDAAANVMEAAHHVPGWVVAAPFVVMVLGFLGAIYAYIWK-----------------------------
------------------------------
PETPAAFAGSFRYLYNLFYNKWYIDEIYDFLFVEPAKKIGVFLWKK-GDENVIDRFGPDGAA-
ASVMTLARRFKALQSGYVYHYAFAMLMGVAAFVTWFI---GGAS-----
```

```
>WP_085883634_NuoL_Oceanibacterium_hippocampi
MY-------VLIVFLPLAAALIAGFFG----RQLGDRGAQVVTSGAVLTSALLSWVALFQVAF--GHAP-
VTITLF-DWVVSGDFAASWAIRVDALTAVMLVVVNTVSSLVHVYSIGYMSHDKA-------
KGRFMAYLSLFTFAMLMLVTADNLLQLYFGWEGVGLASYLLIGFWYHKASANAAAIKAFVVNRVGDFGFALGVV
GIFYIFGSIEF--DQIFAAAPEQVGKS---
FVFMGMEVDVLTTLCLLLFLGAMGKSAQLFLHTWLPDAMEGPTPVSALIHAATMVTAGVFMVARLSPLFEYAPD
ALTVVTVVGGSTAFFAATVGLVQNDIKRVIAYSTCSQLGYMFFAAGVSAYPVAIFHLFTHAFFKALLFLCAGSV
IHAVSDEQDMRRMGGLWKHIKATYVLMWIGSLALA--GIWP---------
FAGYFSKDMVLEAAYAAGTTVGSYAFWLGIAAAIMTAFYSWRLLFMTFHGAPR--ASKEVMDHV----------
---------------------------------------------------
HESPKVMLIPLYVLALGSICAGFIFAPYMVGH-HYQE---FW--
GNSILVLEEHVAMAEAHDVAAWVKFLPLVAGVVGILIAFQLYIRR-----------------------------
------------------------------
PEMPAELARTHQALYQFLLNKWYFDELYDWIFVRPATWLGRVLWKG-GDGRIIDGLGPDGIA-
MTVVRVAARAKQIQTGYIYHYAFAILIGIAAFVTLFFY--GVD------
>MBT5765548_NuoL_Kordiimonadaceae_SI072_bin64
MY-------QTIVFLPLLASIIAGLFG----NRIGVRGAQGVTCGALIVSAVLSWVAFFQVAMAPGDHA-
THVQVL-SWIVSGDFNVNWAFQIDTLTAVMLVVVNTVSCLVHIYSVGYMSHDPH-------
IQRFMSYLSLFTFAMLMLITSDNFVQMFFGWEGVGLASYLLIGFWYKKPSACAAAIKAFVVNRVGDFGFALGIF
AIFMTFGSADF--AVVFASVGDHVDKT---
IHFLGMELDLLTTICLLLFIGAMGKSAQLGLHTWLPDAMEGPTPVSALIHAATMVTAGVFMVARCSPLFEYSPD
ALAVVAVIGASTAFFAATVGMSQFDIKRVIAYSTCSQLGYMFFALGVSAYGAAVFHLFTHAFFKALLFLGSGSV
IHASSDEQDMRNMGGLRKQIPVTFWMMTIGTLALT--GFPF---------
LAGYYSKDMIIESAFAAHSGVGTYAFGLGVAAALMTSFYSWRLVFMTFFGESR--ASKEVQDHV----------
---------------------------------------------------
HESPQVMLIPLYVLALGALLSGFVFKEYFVGH-NYEE---FW--AGALFL-
IGDNVMEAAHHVPTWVIASPFIAMVVGFATAWYFYVKE------------------------------------
-----------------------PSMPGKFTSTFRFAHALSFNKWYFDELYDIIFVKPAMKLGMIFWKR-
GDENIIDRYGPDGVS-AAVVRVAQKFKQFQTGYLYHYAFVMLIGLSALVTFFV--MGGL------
>MBT6031505_NuoL_Kordiimonadaceae_SI072_bin126
MY-------QTIVFLPLLASLIAGLFG----NRIGDRGAQIVTCGAMIVSAVLSWVAFYQVALTPGDNS-
LHVQVL-SWIVSGDFNVNWAFQIDSLTAVMLVVVNTVSCLVHIYSVGYMSHDPH-------
IPRFMSYLSLFTFAMLMLITSDNFVQMFFGWEGVGLASYLLIGFWYKKPSACAAAIKAFVVNRVGDFGFALGIF
AIFMTFGSADF--AVVFADVGSYTDKS---
IHFLGMELDLLTTICILLFIGAMGKSAQLGLHTWLPDAMEGPTPVSALIHAATMVTAGVFMVARCSPLFEYAPA
ALEVVCIVGASTAFFAATVGMSQFDIKRVIAYSTCSQLGYMFFALGVGAYGAAIFHLFTHAFFKALLFLGSGSV
IHASSDEQDMRNMGGLRKKIPLTFWMMTIGTLALT--GFPF---------
LAGYYSKDMIIESAFAAHSSVGSYAFAMGVAAALMTSFYSWRLVFMTFFGESR--ASKEVQDHV----------
---------------------------------------------------
HESPQVMLVPLYVLAFGALFSGFIFHEYFVGA-HYED---FW--AGALFL-
VGDNVMEAAHHVPMWVVEAPFVAMTVGLATAYYFYIRR------------------------------------
-----------------------PEMPGKFVETFRFLHAISFNKWYFDELYDILFVRPAMKLGYILWKR-
GDENIIDRYGPDGVS-AAVVRVAEKFRSLQTGYLYHYAFAMLIGLSAFVTWFV--MGGQ------
>MBT3790209_NuoL_Alphaproteobacteria_SI054_bin61
MY-------TALVFLPLLGAVLAGLFG----RILGDRGSQAVTLIGITTSALLSVYVFFDVT---
GNNNPVVVDLF-TWIESDSFEVSWALRVDQLTAVMLMVVCVVSAVVHWYSVGYMSHDKA-------
IPRFMSYLSLFTFAMLMLVTADNFLQMFFGWEGVGLCSYLLIGFWYDRPSANAAAIKAFLVNRVGDFGFALGIM
AIFFTFNSVEF--DVVFAAAPEMAGKT---
```

```
92… FMFLGQEWDILTTITMLLFVGAMGKSAQLGLHTWLPDAMEGPTPVSALIHAATMVTAGVFMLARCSYLFEYAPF
…   TLEVVTVIGATTAFVAATIGLVQNDIKRVIAYSTMSQLGYMFFAIGVSAYPAAIFHLLTHAFFKALLFLGAGSV
…   IHAMSDEQDMRNMGGIYKLIPGTYLMMWIGSLALA--GVPL---------FAGYYSKDMIIEVAFAG-
…   SGVGTYAFTMGILAALMTAFYSWRLLFMTFHGPAR--ADEKVMAHV----------------------------
…   ---------------------------------HESPKVMMLPLLVLALGAIFAGWFWAGDMIGD-GRVA---
…   FW-GGESIFIQPANHIFENAHHVSAWVKYSPTALGAIGILLAWWFYIKN-------------------------
…   ----------------------------------
…   PNIPKAIATMHREAHLFLLNKWYFDELYDLIFVNPAKRLGHFLWKF-GDGKIIDGLGPDGIS-
…   RLVLNITRRAQTLQSGYVYHYAFAMMLGVAGFVSWFVFGLGG-------
93  >MBT4017013_NuoL_Alphaproteobacteria_SI053_bin31
94  MY-------TALVFLPLLGAVIAGLFG----RVLGDRGSQAVTLIGITTSALLSLYVFFDVT---
…   GNNNPVVIELF-TWMESDSFEVSWALRIDQLTAVMLMVVTVVSAVVHWYSVGYMSHDKA-------
…   IPRFMAYLSLFTFAMLMLVTADNFMQMFFGWEGVGLCSYLLIGFWYDRPSANAAAIKAFLVNRVGDFGFALGIM
…   AIFFTFNSVEF--DVVFAAAPDVAGKT---
…   FVFLGQEWDILTTITMLLFVGAMGKSAQLGLHTWLPDAMEGPTPVSALIHAATMVTAGVFMLARCSPIFEFAPF
…   TLEVVTIIGATTAFVAATIGLVQNDIKRVIAYSTMSQLGYMFFAIGVSAYPAAIFHLMTHAFFKALLFLGAGSV
…   IHAMSDEQDMRKMGGIYKLIPGTYLMMWIGSLALA--GVPL---------
…   FAGYYSKDMIIEVAFASNSGVGMYAFTMGILAALMTAFYSWRLIFMTFHGPAR--ADEKVMAHV----------
…   ---------------------------------------------------
…   HESPKVMMLPLLVLAFGAIFAGWFWFDDMVGD-GRAA---FW--
…   GQSIFIQPANQIFENAHHVSAWVKYSPTALGAFGILLAWWFYIKN-----------------------------
…   ------------------------------
…   PGIPKTIATAHREVHSFLLNKWYFDELYDLIFVRPAKRIGYFLWKI-GDGKIIDGFGPDGIS-
…   RLVLNITRRAQALQTGYVYHYAFAMMLGVAGFVSWFVFGLGG-------
95  >MBT7747133_NuoL_Alphaproteobacteria_SI037_bin135
96  MY-------TALVFLPLLGAVLAGLFG----RILGDRGSQAVTLIGITTSALLSVYVFFDVT---
…   GNNNPVVVDLF-TWIESDSFEVSWALRVDQLTAVMLMVVCVVSAVVHWYSVGYMSHDKA-------
…   IPRFMSYLSLFTFAMLMLVTADNFLQMFFGWEGVGLCSYLLIGFWYDRPSANAAAIKAFLVNRVGDFGFALGIM
…   AIFFTFNSVEF--DVVFAAAPEMAGKT---
…   FMFLGQEWDILTTITMLLFVGAMGKSAQLGLHTWLPDAMEGPTPVSALIHAATMVTAGVFMLARCSYLFEYAPF
…   TLEVVTVIGATTAFVAATIGLVQNDIKRVIAYSTMSQLGYMFFAIGVSAYPAAIFHLLTHAFFKALLFLGAGSV
…   IHAMSDEQDMRNMGGIYKLIPGTYLMMWIGSLALA--GVPL---------
…   FAGYYSKDMIIEVAFASGSGVGTYAFTMGILAALMTAFYSWRLLFMTFHGPAR--ADEKVMAHV----------
…   ---------------------------------------------------
…   HESPKVMMLPLLVLALGAIFAGWFWAGDMIGD-GRVA---FW--
…   GESIFIQPANHIFENAHHVSAWVKYSPTALGAIGILLAWWFYIKN-----------------------------
…   ------------------------------
…   PNIPKAIATMHREAHLFLLNKWYFDELYDLIFVNPAKRLGHFLWKF-GDGKIIDGLGPDGIS-
…   RLVLNITRRAQTLQSGYVYHYAFAMMLGVAGFVSWFVFGLGG-------
97  >MBT5047508_NuoL_Rhodospirillaceae_SI074_bin93_MPNO01
98  MLY------VLCIFLPLIGAAIAGLLG----PWIKARGAMIVTCGALLISALISPFILIEVGL--EGKT-
…   TTIQIF-NWISSGAFEVDWALRFDTLTAVMLVVVTVVSCAVHFYSIGYMSHDPS-------
…   IPRFFSYLSLFTFFMLMLVTADNFLQLFFGWEGVGLASYLLIGFWYEKPSANAAAIKAFLVNRVGDFGFALGIM
…   GVFLLYGTVSF--DAVFDATPGKAAAT---
…   IEFLGGQYHALTVICLLLFVGAMGKSAQLGLHTWLPDAMEGPTPVSALIHAATMVTAGVFMVARLSPMFEYSPS
…   ALAVVTVVGASTAIFAATVGVCQNDIKRVIAYSTCSQLGYMFFACGVSAYAAGIFHLFTHAFFKALLFLGAGSV
…   IHAMSDEQDMRKMGGIWRMIPVTYAVMWIGSLALA--GIPP---------
…   FAGFYSKDIVLEAAYASNTGIGEYAFWMGIAAFLTAFYSWRLLLMTFHGEAR--ADERVMAHV-----------
```

```
---------------------------------------------------
HESPKVMLIPLILLAVGAIFAGYIGVSYFVGE-GAVE---FW--
GNAILVLPSHPALEAAHHVPFWVKALPLVVGVSGIALAYILYVLA-----------------------------
------------------------------
PGLPAAIAGRFRFIHQFLLNKWYFDELYDFLFVRPAFILGRGFWKS-GDGALIDGIGPDGIA-
AATLNLARRAAALQSGYLYHYAFAMLTGVVVLVTWYLFHQVG-------
>MBT4689853_NuoL_Rhodospirillaceae_SI039_bin92
MY-------HAIVFLPLIAFLIVGPFG----RVLGDKPSQWITCGAVITSAVLSIVAFWQIAL--
GGGDTVKVEIA-SWIVSGTFDVSWALRIDALTAVMLVVVNGVSALVHVYSVGYMSHDPH-------
KPRFMAYLSLFTFAMLMLITSDNFLQMFFGWEGVGLASYLLIGFWYHKPSANAAAIKAFLVNRVGDFGFGLGIM
AIFMVFGSLDF--DTVFAAVPGQVGNT---
FHFLNWEFDIITTICILLFIGAMGKSAQVPLHTWLPDAMEGPTPVSALIHAATMVTAGVFLVARCSPMFEFSPT
ALVVVTIVGGFTAFFAATVGLVQNDIKRVIAYSTCSQLGYMFFALGVGAYPVAIFHLFTHAFFKALLFLGSGSV
IHAMSDEQDMRRMGGIFKYVPQTAIMMWIGSLALA--GLPI---------
FAGFYSKDAILESAFAAHTGAGMFAFYAGMAAAVMTAFYSWRLLFMTFNGDPR--ASQEVMSHV----------
---------------------------------------------------
HESPPSMLIPLYFLAAGSILSGMLFVHYFVGD-GHQA---FW--
GDAIFMGPENHILEAMHHVPTWVKIGPIAAGIIGIFVAWLFYIKR-----------------------------
------------------------------
KDIPVNLAKVHHEVYLFLLNKWYFDELYDLIFVRPAKWLGRILWRG-GDGRIIDGYGPDGIA-
AMVMHLARRASALQSGYLYHYAFAMLIGVAVLVTWYY-TTGGGH-----
>MBV40747_NuoL_Rhodospirillaceae_NP113_oxycline
MY-------HAIVFLPLLAFLIVGTFG----RVFGDKPSQWITCGAVIASAILSLVAFWQVAL--GGNP-
VKVEIA-SWIVSGTLDVSWALRIDALTAVMLVVVNGVSALVHVYSIGYMSHDPH-------
KPRFMAYLSLFTFAMLMLITSDNFLQMFFGWEGVGLASYLLIGFWYHKPSANAAAIKAFLVNRVGDFGFGLGIM
ATFMVFGSIDF--NTVFAAAPDQVGKT---
FHFLSWEWDVMTTICILLFIGAMGKSAQVPLHTWLPDAMEGPTPVSALIHAATMVTAGVFLVARCSPMFEFSPT
ALVVVTVVGGSTAFFAATVGLVQNDIKRVIAYSTCSQLGYMFFALGVGAYPVAIFHLFTHAFFKALLFLGSGSV
IHAMSEEQDMRRMGGIFKYVPQTAILMWVGSLALA--GLPI---------
FAGFYSKDAILESAFAAHTGAGDYAFYLGLAAAVMTAFYSWRLLFMTFHGIPR--ASQAVMSHV----------
---------------------------------------------------
HESPLVMLMPLYFLAAGSILSGILFVHYFVGD-GHQA---FW-
RGQAIFMAEGNDILESFHHVPIWVKVAPIAAGVFGIFVAWLFYIRN----------------------------
-------------------------------
QDLPAKLAKLHYPLYLFLLNKWYFDELYDLILVNPAKWLGRLLWKG-GDGKIIDGFGPDGIA-
ATVMNLARRASALQSGYLYHYAFAMLIGAVVLVTWYYTSTGGGH-----
>MBT3373516_NuoL_Rhodospirillaceae_co234_bin8
MY-------HAIVFLPLLAFLIVGTFG----RVLGDRPSQWITCGAVITSAILSLVAFWQVAL--GGQP-
VKVEIA-SWIVSGTFDVSWALRIDALTAVMLVVVNGVSALVHVYSVGYMSHDPH-------
KPRFMAYLSLFTFAMLMLITSDNFLQMFFGWEGVGLASYLLIGFWYHKPSANAAAIKAFLVNRVGDFGFGLGIM
ATFMVFGSLDF--DTVFAASPDQVGKT---
FLFLSWEWDVMTTICILLFIGAMGKSAQVPLHTWLPDAMEGPTPVSALIHAATMVTAGVFLVARCSPMFEFSPT
ALVVVTIVGGTTAFFAATVGLVQTDIKRVIAYSTCSQLGYMFFALGVGAYPVAIFHLFTHAFFKALLFLGSGSV
IHAMSDEQDMRRMGGIFKYVPQTAILMWIGSLALA--GLPI---------
FAGFYSKDAILESAFAAHTGAGDYAFYLGLAAAVMTAFYSWRLLFMTFNGTPR--ASQEVMSHV----------
---------------------------------------------------
HESPLVMLIPLYFLAAGSILSGILFVHYFVGD-GAEA---FW--
GQAIFMAEGNNILEAMHHVPMWVKIAPIAAGVIGILLAWQFYIRR-----------------------------
```

```
------------------------------
TDIPVKLAKMHHELYLFLLNKWYFDELYDLIFVNPAKWLGRVLWKG-GDGRIIDGFGPDGIA-
ATVMRVARRASALQSGYLYHYAFAMLIGVAVLVSWYYLSSGGGH-----
>MBL6951394_NuoL_Alphaproteobacteria_BS150m-G9_BlackSea
MY-------HAIVFLPLLAFLIVGSLG----RVLGDRPSQWITCGAVIISALLSLLAFWQVAL--GGQA-
VRVDIG-SWIVSGTFDVSWALRIDQLTAVMLVVVNGVSALVHVYSVGYMSHDPH-------
KPRFMAYLSLFTFAMLMLVTADNFLQLFFGWEGVGLASYLLIGFWYHRPTANAAAIKAFLVNRVGDFGLALGIM
AIFTVFGSLDF--DTVFAATPGQVGKS---
FNFLSWEWDVMTTICILLFIGAMGKSAQVPLHTWLPDAMEGPTPVSALIHAATMVTAGVFLVARCSPMFEFSPT
ALVVVTIVGGTTAFFAATVGLVQNDIKRVIAYSTCSQLGYMFFALGVGAYPMAIFHLFTHAFFKALLFLGSGSV
IHAMSDEQDMRRMGGIYKYVPQTTGLMWVGSLALA--GFWP---------
LAGYYSKDAILESAFAVHTGAGEYAFWLGMAAAVMTAFYSWRLLFMTFAGEPR--ASQEVMRHV----------
---------------------------------------------------
HESPLSMLIPLYFLAAGSILSGVLFVHYFVGE-GHVE---FW--
GQAIFMAPGNDVMEAVHHVPTWVKVAPLVAGIVGILVAWQFYIRR-----------------------------
------------------------------
RDIPAGLAKINHEVYEFLLNKWYFDELFELIFVRPAKWLGRLLWKT-GDGRIIDGFGPDGIA-
ASVLNLARRASALQSGYLYHYAFAMLIGVAALVSWYYISSGGGH-----
>MDP6874949_NuoL_Alphaproteobacteria_Arabian9_MAG_8_ODZ
MY-------HAIVFLPLLAFLIVGALG----RILGDKPSQWITCGAVITSAILSLVSFWQVAL--GGQP-
VKVEIA-SWIVSGTFDVSWALRIDALTAVMLVVVNGVSALVHVYSVGYMSHDPH-------
KPRFMAYLSLFTFAMLMLITSDNFLQMFFGWEGVGLASYLLIGFWYHKPSANAAAIKAFLVNRIGDFGFGLCIM
ATFMVFGSVDF--DTVFAAAPDQVGKT---
LHFLNWEWDVMTTICILLFIGAMGKSAQVPLHTWLPDAMEGPTPVSALIHAATMVTAGVFLVARCSPMFEYSPV
ALTVVTIVGASTAFFAATVGLVQNDIKRVIAYSTCSQLGYMFFALGVGAYPVAIFHLFTHAFFKALLFLGSGSV
IHAMSDEQDMRRMGGIFKYVPQTAIMMWIGSLALA--GLPI---------
FAGFYSKDAILESAFAAHTGAGDYAFYLGLAAAVMTAFYSWRLLFMTFNGTPR--ASQEVMSHV----------
---------------------------------------------------
HESPPSMLIPLYFLAAGSILSGILFSHYFIGE-GHEA---FW--
GQGIFMLPDNDILEALHHVPMWVKIAPIAAGIIGILVAWQFYIRR-----------------------------
------------------------------
TDLPAKLAKVHHEAYLFLLNKWYFDELYDLIFVNPAKWLGRLLWKG-GDGRIIDGFGPDGIA-
ATVMRVARRASALQSGYLYHYAFAMLIGVGVLVTWFYTSSGGGH-----
>MBM3510012_NuoL_Alphaproteobacteria_K_Offshore_80m_m2_115
MFY------AAIVFLPLVGAIAAGLFG----RLLGVRASQLVTCLALTLSGVLATWGLYETAV-IGPDK-
QQIVLF-TWIVSGDFDVSWTLRIDTLTAVMLFTVSVISSIIHWYSIGYMAHDPH-------
IPRFFAYLSLFTFAMLVLVTADNFLQLYFGWEGVGLCSYFLIGFWYDRPSANAAAIKAFVVNRVGDFGFALGIM
AMFFTFNSVNF--DTVFAGAPAVAGKT---
FEFLGAEYDILTVTTLLLFIGAMGKSAQVPLHTWLPDAMEGPTPVSALIHAATMVTAGVFLVARCSPLFEYAPT
TLAIVTVIGAFTAFFAATVGLVQNDIKRVIAYSTCSQLGYMFFACGVGAYAVAIFHLFTHAFFKALLFLGSGSV
IHGFSDEQDMRRMGGVWRKMPVTYAVMWVGSLALA--GIPF---------
FAGYYSKDLIIESAFAASTEVGYLAFYLGVGAALLTAFYSWRLLFMTFHGPTR--ANPEVYEHV----------
---------------------------------------------------
HESPKIMTVPLLALAVGALFAGVAGYDAFVGD-GRAA---FW--
REAIFLLPGHDVIEAAHHVPLWVKLLPIAIASAGIALAWQFYIRR-----------------------------
------------------------------
TELPGALAKTHADLYQFLLNKWYFDELYDAIFVRPAKFLGRALWHG-GDGRIIDGFGPNGIA-
QVVIQLARRASLLQTGYLYHYAFAMLIGVAALVTWFTRTGGGAP-----
```

```
>MBM3570281_NuoL_Alphaproteobacteria_M_DeepCast_65m_m2_129
MLY------GLIVFLPLIAAIVAGFGG----RALGDRGAQVVTCGALIVSALLSGVAFYQVAW--KGQP-
VTVELL-TWIDSGAFDVMWSLRIDQLTAVMLVVVTWVSAVVHVYSVGYMAHDPH-------
IPRFMAYLSLFTFAMLMLVTSDNFVQLYFGWEGVGLASYLLIGFWYHKPSANAAAIKAFVVNRIGDFGFGLGIC
GVFLVFGTVEF--DPVFQTAPGFAGKT---
FEFLGMKLDMLTVICLLLFMGAMGKSAQVPLHTWLPDAMEGPTPVSALIHAATMVTAGVFLVARCSPLFEYAPD
ALAVVACIGAFTAFFAATVGLCQNDIKRVIAYSTCSQLGYMFFAAGLSAYAAAIFHLFTHAFFKALLFLGSGSV
IHGMSGEQDMRRMGGLLAPLKITYVLMWIGSLALA--GVPL---------
FAGYYSKDAIVETAFMAGGKLGSFAFVMGVAAAFMTAFYSWRLLFMTFHGQNR--HPDRVIEHAH---------
---------------------------------------------------
HESPAVMLVPLYVLAAGSIFAGALFAQAFIGE-GSPI---FG--ASILVL-
PGHDALAQAHHAPLWVKLAPVAVGLAGIALAWWMYVRR------------------------------------
-----------------------PDLPGKLAAMHGDAHRFLLNKWYFDELYDALFVRPAKWLGHALWKR-
GDGKVIDGLGPDGIA-ALTLDIAKRATRLQTGYVYHYAFAMLIGVALLVSWYLY-RGGM------
>OUR77105_NuoL_Alphaproteobacteria_46_93_T64_Sneathiellales
MY-------VLIVFLPLLAALIAGFGA----RQLGDRGSQIVTSGAVSASAILSWVALFQVAF--GSAP-
TTIELF-TWINSGTFDVSWSLRVDSLTAVMLVVVNTVSALVHIYSIGYMSHDPH-------
KPRFMSYLSLFTFAMLMLVTSDNFLQMYFGWEGVGLASYLLIGFWFNKPSANAASIKAFVVNRVGDFGFALGIM
AIYLVFDSISF--DTVFAAVPEKVGET---
FNFLGYEVDVITTIALLLFLGAMGKSAQLGLHTWLPDAMEGPTPVSALIHAATMVTAGVFMVARCSPIFEFSPT
ALAVVTVVGASTAFFAATVGLVQNDIKRVIAYSTCSQLGYMFFAIGVGAYPVAIFHLFTHAFFKALLFLGSGAV
IHAVSDEQDMRKMGGLGKHIKITYAMMFIGTVALT--GVPF---------
FAGFYSKDAIIESAFMAGTDVGNYAFWCGIGAALMTAFYSWRLMFMTFHGKPR--ASKEVMSHV----------
---------------------------------------------------
HESPNVMLIPLYVLAAGAIGAGFVFYEPMVGH-GYKE---FW--
GNAILLRESNTIMTDFHNIPSWVFWAPMIAMIAGFLLAFNMYIRR-----------------------------
------------------------------
TDIPVQLAKTHSALYQFLLNKWYFDELYDVVFVRPARWIGSKLWTI-GDGKIIDGLGPDGIA-
ARVLDLAKRASMLQSGYLYHYAFAMMIGVAAFVSFFFL-AGGGH-----
>WP_144258814_NuoL_Ferrovibrio_terrae
MY-------HLIVFLPLLAAIIAGLFG----RRIGDVASMAVTCVAVIISAVLSWVAFYLVIS--KGMV-
VTVKVL-DWIDSGTFSVDWALKIDQLTAVMLVVVNTVSAVVHVYSVGYMSHDPH-------
RSRFFSYLSLFTFAMLMLITADNFVQLYMGWEGVGLASYLLIGFWFHKPSANAASIKAFVVNRVGDFGFALGIM
ALFFATGSVTF--EAVFAAAPDLAGKT---
FHFLWKDWDILTVATFLLFLGAMGKSAQLGLHTWLPDAMEGPTPVSALIHAATMVTAGVFMVARCSPLFEYAPD
TLAFVTVIGASTAFFAATVGLAQNDIKRVIAYSTCSQLGYMFFALGVSAYPAAIFHLFTHAFFKALLFLGAGSV
IHAVDGEQDMRRMGGLWKHIKITYAMMWIGSLALA--GFPI---------FAGYYSKDMILEVAYAA-
GGVGQYAYIMGMAAAILTAFYSWRLLFMTFHGKPR--ADHHVMEHV----------------------------
---------------------------------HESPMVMLIPLFVLATGAVLAGIVFYDGFVGH-HWKE---
FW--GSSILILENHKAMDEAHHVPFLVKVGPIIVGVIGIFIAWIAYIRD-------------------------
----------------------------------
TSLPGRTAAQHHMLYSFLLNKWYFDELYDRVFVRPTFWLGNLLWKG-GDGKIIDGLGPDGLA-
ATVVRLARRASILQSGYVVYHYAFAMLIGVAVLVTYFFSAMGH-------
>WP_206376079_NuoL_Sneathiella_chungangensis
MMY------VLIVFLPLLAALIAGLFG----RSLGDRGSQLITCGAVIISAILSWVALYQIAF--QQTT-
EAVELL-TWINSGDFDVNWALRIDSLTAVMLVVVNTVSAVVHVYSIGYMSHDPH-------
KPRFMAYLSLFTFAMLMLVTSDNFVQLYFGWEGVGLASYLLIGFWYKKPAANAAAIKAFVVNRVGDFGFALGIM
GIYFVFNSVSF--DEVFAAAPSQAGQT---
```

```
118… FEFLGYQVDILTTLCLLLFVGAMGKSAQIGLHTWLPDAMEGPTPVSALIHAATMVTAGVFMVARCSPIFEYSPE
   … ALMVVTVVGATTCFFAATIGLVQNDIKRVIAYSTCSQLGYMFFALGIGAYPAAIFHLFTHAFFKALLFLSAGSV
   … IHAVSDEQDMRKMGGLWKHIKVTYAMMWIGSLALA--GIPL---------
   … FAGYYSKDLIIESAFAAHTGVGDYAFWAGVAAALLTAFYSGRLIFMTFHGEPR--ASKEVMAHV----------
   … ---------------------------------------------------
   … HESPMVMLIPLGVLAAGAIFAGMLFAGPMTGE-DFRE---FW--
   … GASILMLENSTAMLDAHHVPLWVKLLPLVLGVFGIALTYQMYIRR-----------------------------
   … ------------------------------
   … TDIPVQLAKTHSALYQFLLNKWYFDELYDFLFVRPAKAIGRFFWTF-GDGKVIDGLGPDGIA-
   … GRVLVLARRASQLQSGYMYHYAFAMLLGVAAFVSYFFFAGGH-------
 119 >GAK32918_NuoL_alpha_proteobacterium_Q-1
 120 MSQF-----HIIVFGPLLGFLIAGLFA----KQIGDRGAMAVTTGLVTLSAILSAFVLKDVAF--DHNQ-
   … YRILVL-EWVRSGSLSFDWTLYIDSLTAVMLVVVNTVSALVHWYSIGYMSHDPH-------
   … RARFFAYLSLFTFAMLMLVTSDNLVQMFFGWEGVGLASYLLIGFWFKKPSANAAAIKAFVVNRVGDFGFSLGIF
   … ALFLLTGSVIF--EDIFQSLEGFADAR---
   … FVFLGMDVHALTTIAVLLFVGAMGKSAQLFLHTWLPDAMEGPTPVSALIHAATMVTAGVFLMARMSPLLELAPG
   … ALTLIMIIGGATAFFAASVGLVQNDIKRVIAYSTCSQLGYMFVAIGASAYGAAIFHLFTHAFFKALLFLGAGSV
   … IHAMSDEQDMRKMGGLSKMVPLTYLMMMVGTLALT--GFPY---------
   … TAGFFSKDMIIESAFATHNPAASFGFLMTVIAAFMTSFYSWRLVFMTFHGTPR--ADEKTLSHV----------
   … ---------------------------------------------------
   … HESPAVMWVPLAILAVGAVIAGVVFSDAFIGE-GRLA---FW--AGSLVA-
   … GEGDVMDEAHHIASWIKWLPTGMMALGFALSFLFYIAK------------------------------------
   … -----------------------PGLPAAMARTFPAAYQFLLNKWYFDELYDRIFIRPAFWIGRFLWVR-
   … GDQKTIDGLGPDGIA-AQVMAAARRVRVLQSGYVYHYAFAMLIGIALVITYFVLTGGGF------
 121 >WP_150006121_NuoL_Iodidimonas_muriae
 122 MSQFMSQ-FHIIVFGPLLGFLIAGLFG----RRLGDRGAMGVTTGLVTLSAVLSAFALKDVAF--DHNQ-
   … YRILVL-EWVRSGTLSFDWVLNIDTLTAVMLVVVNTVSALVHWYSIGYMSHDPH-------
   … RSRFFAYLSLFTFAMLMLVTSDNLVQMFFGWEGVGLASYLLIGFWFKKPSANAAAIKAFVVNRVGDFGFSLGIF
   … ALFMLTGSVVF--ADIFASLDGVADAR---
   … FIFLGYEVHALTTIAVLLFIGAMGKSAQLFLHTWLPDAMEGPTPVSALIHAATMVTAGVFLMARMSPILELAPG
   … AMTLIMVIGGATAFFAASVGLVQNDIKRVIAYSTCSQLGYMFVAIGASAYGAAIFHLFTHAFFKALLFLGAGSV
   … IHAMSDEQDMRKMGGLSKMIPLTYVMMMVGTLALT--GFPY---------
   … TAGFFSKDMIIEAAFATHNSMGSFGFLMTVVAAFMTSFYSWRLVFMTFHGTPR--ADEKVMSHV----------
   … ---------------------------------------------------
   … HESPAVMWVPLAILAVGAIAAGFVFSDYFIGH-DRFD---FW--AGALVT-
   … SHGDVMDEAHHIPGWIKWLPTGMMALGFLLSVLFYVVK------------------------------------
   … -----------------------PGIPKSMAATFPGAYKFLLNKWYFDELYDRIFVRPAFWLGRVLWIR-
   … GDQKTIDGFGPDGIA-ATVLAAAKRIRVLQSGYVYHYAFAMLIGIAIVISYFVLTGGGF------
 123 >RMF12532_NuoL_Alphaproteobacteria_J084
 124 MNQ-----FHVIVFGPLVGFLIAGLFG----RRIGDRASMLLTSLLVSLSAVLSIPVFIDTAF--SGGQ-
   … FSVPVL-EWVQVGMLHFTWSLHVDTLTAVMLVVVTVVSSLVHWYSIGYMDEDPD-------
   … KPRFFAYLSLFTFAMLMLVSADNLVQTFFGWEGVGLASYLLIGFWFKKPSANAAAIKAFVVNRVGDFGFSLGIF
   … ALFMVTGSVGF--EKIFAKVPSLAQAD---
   … FIFLGHQAHALTVIGVLLFIGAMGKSAQILLHTWLPDAMEGPTPVSALIHAATMVTAGVFLVARMSPLYEHAPA
   … AADLVTFIGATTAFFAATVGLVQNDIKRVIAYSTCSQLGYMFVAAGVSAYGAAIFHLFTHAFFKALLFLGAGSV
   … IHAMHHEQDMRRMGGLRPMIPLTYLMMIVGTLALT--GFPL---------
   … TAGYFSKDSIIEAAFAHGGSIGEYAFVMTVVAAALTSFYSWRLIFLTFHGRPR--ASEEVLAHV----------
   … ---------------------------------------------------
```

```
124… HESPAVMTIPLMVLAAGALFAGWGFHEFFVGE-ARGE---FW--NGALAA-
…    GGGRVLDHIHHLPWTITYLPTVMMVLGFVIAFWFYLAN------------------------------------
…    -----------------------PAVPVALAQRMNGLYRFLLNKWYFDELYDLLFVRPAFRIGRALWKI-
…    GDGRIIDGLGPDGVA-RAVMAGAARIRRIQTGCIYTYAFAMLIGVVLLVATVMLAERGF------
125  >NVJ99269_NuoL_Alphaproteobacteria_HF-Din21
126  MES------KFIVFLPLVGFLIAGMFG----RNIGDRASQVLTSSLVTIGAALSWYVFADVAF--VHNV-
…    YPVHVL-DWVKSGTLEFAWAFKIDTLTAVMLVVVNTVSALVHWYSMGYMEEDPD-------
…    KPRFFAYLSLFTFAMLMLVTSDNLVQMFFGWEGVGLASYLLIGFWFKKPSANAAAIKAFVVNRVGDFGFALGIY
…    AVFLMTGSVEF--DVIFANIGDYQTAT---
…    MTFLGHEYHAITVICLLLFVGAMGKSAQLGLHTWLPDAMEGPTPVSALIHAATMVTAGVFLVARFSPVFEYSEY
…    ALQVVTIVGASTAFFAASVGLVQNDIKRVIAYSTCSQLGYMFFALGVGAYGGAVFHLFTHAFFKALLFLGAGSV
…    IHALHHEQDMRNMGGLFKKIPFTGTVMIIGTLAIT--GVPF---------
…    LSGFYSKDLIIEAAYASDAGASQFAFVMGVAAAFMTSFYSWRLIFLTFFGETR--AKQKYFDHA----------
…    ---------------------------------------------------
…    HEGPIFIKVPLAILALGALGAGFIFKEFFIGH-ERVA---FW--NGALVT-
…    AMDDVMEAAHHVPGAVIAAPFVVMLLGFLLAFAFYGRK------------------------------------
…    -----------------------TDLPKRTAEMWDGLYAFLLNKWYWDELYNFLFVRPAFWLGNVFWKR-
…    GDEQTIDGFGPNGVS-AAVAAVAAKARKLQTGYVVHYAFAMIIGLAVVVTWFMAQSGAH------
127  >WP_121938164_NuoL_Eilatimonas_milleporae
128  MLY------KLIVFLPLIGFLVAGLFG----RRIGDRGSQLLTSGLVTISAILSWTAFVDVAL--LHNK-
…    YSIHVL-DWVTSGSLEFNWAFKIDTLTAVMLVVVNSVSALVHWYSMGYMHEDPD-------
…    KPRFFAYLSLFTFAMLMLVTADNLVQMFFGWEGVGLASYLLIGFWYTKPSANAAAIKAFVVNRIGDFGFSLGIF
…    AVFVLFNTVQF--DGIFTQVETYKDAT---
…    MVFLGQEYHALTVISLLLFVGAMGKSAQLGLHTWLPDAMEGPTPVSALIHAATMVTAGVFLVARFSPVFDLAPD
…    ALAVVTVIGAATAFFAATIGLVQNDIKRVIAYSTCSQLGYMFFALGVSAYGAAIFHLFTHAFFKALLFLGAGSV
…    IHAMHHEQDMRKMGGVRQYAPLTYGMMIIGTLALT--GFPF---------
…    TAGYYSKDMIIEAAYASHSAGAQFAFVMGVAAALMTSFYSWRLIFMTFTGECR--ADEHTRLHA----------
…    ---------------------------------------------------
…    HESPWIMLIPLVLLSLGALFAGFGFHEMFIGH-DRIA---FW--NGSLVT-
…    AAGDVIDEAHHVPFGVKLAPMVAMALGFLMAWIMYIRS------------------------------------
…    -----------------------TDIPGRLAKTWDGLYAFLLNKWYVDELYNLIFVRPAFWIGRQFWKR-
…    GDQQTIDGLGPNGVS-GAVAAIAARARKLQSGYLYHYAFAMIVGLALVVTWFMVRAGG-------
129  >WP_191253427_NuoL_Kordiimonas_sediminis
130  MMY------KLIVFLPLIGFLFAGLFG----SKFSDRISQVLTAGLVTVSAVLSWMAFADVAL--GGNQ-
…    YTVLVL-EWVSSGALSFDWAFKIDTLTVVMLVVVNSVSALVHWYSFGYMEDDPS-------
…    KPRFFGYLSLFTFAMLMLVTSDNLVQMFFGWEGVGLASYLLIGYYYKKPSANAAAIKAFVVNRVGDFGFSLGIF
…    GAFALFGSVMF--DDIFASADQYADYY---
…    INFLGTDFHAITVICILLFVGAMGKSAQLGLHTWLPDAMEGPTPVSALIHAATMVTAGVFLVARFSPVFEMSPT
…    ALAVVTIIGATTAFFAATVGLVQNDIKRVIAYSTCSQLGYMFFALGVSAYGAAIFHLFTHAFFKALLFLGSGAV
…    IHAMHHEQDMRNYGGLAKKIPFTFAMMTIGTIALT--GFPG---------
…    TAGFYSKDMILEAAYASHAFGAQYAFVLGIAAAFMTSFYSWRLAHLTFFGECR--ADEHTASHA----------
…    ---------------------------------------------------
…    HEAPWIMRLPLVILAVGALFAGVVFADMFVGE-GRFA---FW--NGALVT-
…    SVGDVLDEAHHVPFAVKAAPFVVMLLGFIFATVMYYKV------------------------------------
…    -----------------------TDIPERMAKTWDGLYAFLLNKWYFDELFNFIFIRPAFWLGRVFWKK-
…    GDEATIDGFGPNGVS-HAIAFVAARARKLQTGYVVHYAFAMIIGLSLAITWFMVTGSSH------
131  >WP_068306026_NuoL_Kordiimonas_lacus
132  MET----AAKFIVFLPLVGFLIAGMFG----RTIGDRVSQVLTSSLVTIGAILSWLVFVDVAL--AGNK-
```

132… FVVTLL-DWVKSGDLEFAWALQVDTLTAVMLVVVNTVSALVHWYSMGYMEEDPD-------
KPRFFAYLSLFTFAMLMLVTSDNLVQMFFGWEGVGLASYLLIGFWFKKPSANAAAIKAFVVNRVGDFGFALGIY
AVFLLTGSVQF--DVIFSTIGNYETAT---
LSFLGHEYHAITVVCLLLFVGAMGKSAQLGLHTWLPDAMEGPTPVSALIHAATMVTAGVFLVARFSPVFEYSAY
ALEVVAIVGATTAFFAASVGLVQNDIKRVIAYSTCSQLGYMFFALGVGAYGGAIFHLFTHAFFKALLFLGAGSV
IHALHHEQDMRKMGGLFKKIPITATVMTIGTLAIT--GFPL---------
LSGYYSKDLIIQAAFASDSSVAQYAFVMGVAAALMTSFYSWRLVWMTFFGETR--AKQKYFDAA----------
---------------------------------------------------
HEGPITIMGPLLILATGAVFAGAVFSSYFVGD-DRVA---FW--NGALVT-
QMGDVMDAAQSVPMGVTLAPFVVMVLGFFISMMFYAPRIAMPKFAAAGLAVLTLALFIAAKFTDLHHIGFMKYV
GYFAGFAFAIFYYAYLYAAGKVTNDLPQRTATMWNGLYAFLLSKWYWDELYDFLFVRPAFWLGRVFWKR-
GDEQTIDGFGPNGVS-STVALVAAKARKLQTGYVYHYAFAMIIGLAVVVTWFMAQSGAH------
133 >WP_068149388_NuoL_Kordiimonas_lipolytica
134 MET----AAKFIVFLPLVGFLIAGMFG----RQIGDRFSQVLTASLVTIGAILSWLVFVDVAL--AGNK-
FVVTLL-DWVKSGDLEFAWALKVDTLTAVMLVVVNSVSALVHWYSMGYMEEDPD-------
KPRFFAYLSLFTFAMLMLVTSDNLVQMFFGWEGVGLASYLLIGFWFKKPSANAAAIKAFVVNRVGDFGFALGIY
AVFLLTGSVQF--DVIFGSIGNYQEAT---
LSFLGHDYHAITVVCLLLFVGAMGKSAQLGLHTWLPDAMEGPTPVSALIHAATMVTAGVFLVARFSPVFEYSEY
ALQVVAIVGATTAFFAASVGLVQNDIKRVIAYSTCSQLGYMFFALGMGAYGGAIFHLFTHAFFKALLFLGAGSV
IHALHHEQDMRNMGGLFKKIPMTGTVMLIGTLAIT--GFPL---------
LSGYYSKDLIIEAAFASDSSVAQYAFVMGVAAALMTSFYSWRLVWLTFFGETR--AKQKYFDAA----------
---------------------------------------------------
HEGPIFIKAPLVILALGAVFAGAVFKEYFVGH-ERVA---FW--NGSLVT-
QMGDVIDAAHNVPMGVTLAPFVVMVLGFFISMMFYAPRIALPKLPAAGLAVLTLALFIAAKFTDLHHVAGMTYV
GYFSGFAFAIFFYAYLYAAGKVTNDLPQRTATMWSGLYAFLLNKWYWDELYDFLFVRPAFWLGRVFWKR-
GDEQTIDGFGPNGVS-ATVALVAAKARKLQTGYVYHYAFAMIIGLALVVTWFMAQSGAH------
135 >MBO6503706_NuoL_Kordiimonadaceae_HXMU1429-5
136 MLF------KLIVFLPLVGFLIAGMFG----RQIGDRASMILTSGLVSIAGILSWFAFFDVTA--GGNQ-
YSVHVL-SWVTSGTLTFDWAFHIDTLTAVMLIVVNTVSALVHWYSMGYMEEDPH-------
KPRFFAYLSLFTFAMLMLVTSDNLVQMFFGWEGVGLASYLLIGFWYKKPSANAAAIKAFVVNRVGDFGFALGIY
AVFLLVGSVEF--STIFARIGEFETAT---
VSFLGTEYHAITVAALLLFVGAMGKSAQLGLHTWLPDAMEGPTPVSALIHAATMVTAGVFLVARFSPLFDLSPV
ALSVVTYIGAATALFAASVGLVQNDIKRVIAYSTCSQLGYMFVALGVTAYGGAIFHLFTHAFFKALLFLGAGSV
IHAMHHEQDMRKMGGIYKKIPLTYIVMLIGTLAIT--GVPF---------
LSGYYSKDLIIEAAYAADAPGAQFAFVMTVVAALMTSFYSWRLIFLTFHGQTR--ADNHTFDHA----------
---------------------------------------------------
HEGPAVMTIPLIMLAVGALAAGGYFASSFIGD-GRFA---FW--NGALVT-
TAGDVMDLAHEVPSAVVWAPFIAMILGLFISACFYRPQIKMPAVAAFGLAVAAVAVLVLVPSQYA---------
GYAAVAALALSLYVLFGAWGWLSDDLPKRTAETWDGLYAFILNKWYFDELYDFLFVRPAFWLGKIFWKK-
GDEATIDGFGPNGVS-ALVAGVAARVRKLQTGYVYHYAFAMIIGLALVVTWFMMQGGAH------
137 >WP_194211999_NuoL_Kordiimonas_pumila
138 MI------PKAIVFLPLIGFLIVGILG----SKLGDKLSQIITASLVTVAAVFSWVVFADVTM--VHNT-
YDIHIL-DWVTSGTLSFNWALKIDTLTAVMLVVVNSVSALVHWYSMGYMSEDPN-------
KPRFFAYLSLFTFAMLMLVTSDNLVQMFFGWEGVGLASYLLIGFWYKKPSANAAAIKAFVVNRVGDFGFALGIY
AVFVLFGSVEF--ETIFANAGNYQDAT---
LNFLGHDYHALTVICLLLFVGAMGKSAQLGLHTWLPDAMEGPTPVSALIHAATMVTAGVFLVARFSPVFQYSPD
ALAVVTVVGAMTAFFAASVGLVQNDIKRVIAYSTCSQLGYMFFALGVGAYGGAIFHLFTHAFFKALLFLGAGSV
IHAMHHEQDMRNMGGLYKKIPFTFGVMLVGTLAIT--GFPF---------

```
LSGYFSKDMIIEAAYASHSAGSEFAFVMGVTAALMTSFYSWRLIYMTFFGKTR--ADHHTFDHA----------
---------------------------------------------------
HEGPWVMRIPLFLLAVGAVFAGGVFAGEFIGH-DKVE---FW--NNALVV-
AAGDVMDEAHHVPAAVKFAPFVAMVLGFVLATFFYLRK------------------------------------
-----------------------SDIAERTAETWDGLYKFLLNKWYFDELYNLIFVRPAFWLGRQLWKR-
GDEQTIDGFGPNGVS-HAVAVIATKVRKLQTGYVYHYAFAMIVGLAAVVTWFMVNTGGSH-----
>WP_201240297_NuoL_Rhodothalassium_salexigens
MI-------ELIVFLPLAGFLAAGLAG----QSLGDRGSMLLTSSFVTVSALLSVALFFEVS---
GNPDAFHTVTLGSWIASGDLAIDWALRVDTLTAVMLVVVNGVSALVHWYSVGYMAHDPS-------
KPRFFAYLSLFTFAMLMLVTADNLVQMFFGWEGVGLASYLLIGFWYQKPSANAASIKAFVVNRIGDFGYSLGIF
ATFLLFGSVQF--DTIFANVDAVKGTT---
LDFVGYDVPALTLVALLLFVGAMGKSAQLILHTWLPDAMEGPTPVSALIHAATMVTAGVFLIARMSPLFEAAPD
ALAVVTVIGAATAFFAATVGLVQNDIKRVIAYSTCSQLGYMFFALGVGAYGAGIFHLFTHAFFKALMFLSAGSV
IHALSDEQDMRRMGGLRPHLQTTWALMLVGTLALTGVGIPG------
LIGFSGFYSKDIVIEAAYAGEVGGSLFAFVMGIAAVFMTSFYSWRLMFMTFENKPR--ASKDVMGHI-------
------------------------------------------------------
HESPMVMLVPLFVLAGGAVFAGLAFQDWFVGD-GRLA---FW--AGSIVQ-
GEHDVIEQAHHVSKWVKWLPTAMMALGLGAAWLFYIRS------------------------------------
-----------------------PQLPGQVAASAKGLYQFLLNKWYFDELYRALIVRPVFALGRLLWIA-
GDQKTIDGLGPDGVS-RTLVAGGKRMRRVQSGYVYHYAFAMLVGVVILVSYIAFYSGGF------
>WP_132706734_NuoL_Rhodothalassium_salexigens
MI-------ELIVFLPLAGFLAAGLAG----QSLGDRGSMLLTSSFVTVSALLSVALFFEVS---
GNPDAFHTVTLGSWIQSGDLAIDWALRVDTLTAVMLVVVNGVSALVHWYSVGYMAHDPS-------
KPRFFAYLSLFTFAMLMLVTADNLVQMFFGWEGVGLASYLLIGFWYQKPSANAASIKAFVVNRIGDFGYSLGIF
ATFLLFGSVQF--DTIFANVDAVKGTT---
LDFVGYDVPALTLVALLLFVGAMGKSAQLILHIWLPDAMEGPTPVSALIHAATMVTAGVFLVARMSPLFEAAPD
ALAVVTVIGAATAFFAATVGLVQNDIKRVIAYSTCSQLGYMFFALGVGAYGAGIFHLFTHAFFKALMFLSAGSV
IHALSDEQDMRQMGGLRPHLRTTWALMLVGTLALTGFGIPG------
LIGFSGFYSKDIVIEAAYAGEVGGSLFAFVMGVAAAFMTSFYSWRLMFMTFENKPR--ASKDVMSHI-------
------------------------------------------------------
HESPMVMLVPLFVLAGGAVFAGFAFQDWFVGD-GRLA---FW--AGSIVQ-
GEHDVVDKAHHVSKWVEWLPTAMMALGLGAAWLFYIRS------------------------------------
-----------------------PQLPGQVAASAKGLYQFLLNKWYFDELYRALIVRPIFALGRLLWIG-
GDQKTIDGLGPDGVS-RTLVAGAKRMRRVQSGYVYHYAFAMLIGVVILVSYIAFYSGGF------
>PPR16856_NuoL_Alphaproteobacteria_MarineAlpha9_Bin3
ML-------KLLVFLPLMGSVLSGLFP----KFLGKKGVMWLPTLFSFLSLLFSIILLSQAI---TGES-
YITHLG-YWVNSEALTVSWSLRLDVLSAVMLFVVMLVSSLVHLYSIGYMDHDEH-------
KSRFFSYLSLFTFAMLMLVTSNNFLQLFFGWEGVGLCSYLLIGFWFKKQSANAAAIKAFIVNRVGDLGLILGIC
IIYKLTGSLDF--DKVFEKTSLFLESS---
ILFFNFSIPYIELACFLLFIGAMGKSAQIGLHTWLADAMEGPTPVSALIHAATMVTAGVFLVARCSPLFEYAPV
ALNFVILVGAITSIFAACIAITQNDIKKIIAYSTCSQLGYMFFACGVSAYSAGMFHLMTHAFFKALLFLGAGSV
IHALSDEQDIRKMGGLAKVLPYTCAAMWIGSLALA--GLPP---------
FAGFFSKDIILEAAWGHHSIFGYIAFYLGLIAAFLTAFYSWKILFMVFHGKPR-----TDISKA----------
---------------------------------------------------
HESPLYILIPLLILSIGAIISGWLGYSM-VDT-HHD----FW--
LESILILDNHQALVNAHHVPLLVKLSPIIVAILGIYFAWIFYIKK-----------------------------
------------------------------
VDLPKKNADKFPKLYNISKNKFWFDEIYDLILTRQLKNIGNILWTK-GDIRIIDRFGPNGIA-
```

```
YFCSRLASKIRSFQSGYVVYHYAFTMFFGLIILFSWQFLIYFGW-----
>AIL65955_NuoL_Rickettsiales_Ac37b
MKNEI----VFLVFIPLITSVISGLFM----QRVNAALAQFLTCIGMAISSFLSLIIFYNIIIK-GYNY-
NTVELL-RWIEINELVVNWALKVDALTAVMLVVVTVVSFLVHVYSVGYMHDDQH-------
KPRFMSYLSLFTFFMLMLVTSDNFLQLFLGWEGVGLCSYLLIGFWYQKNSANLAAMKAFIVNRVGDFGFILGIF
SIYLIFNSLQF--DQIFKLVNEYSYKT---
FNILGYNISAIEISCMLLFIGCMGKSAQLGLHIWLPDAMEGPTPVSALIHAATMVTAGVFLLVRCSWLFQYAPI
VLYIITVIGALTCIFAATIALTQNDIKKIIAYSTCSQLGYMFFACGVSAYNVALFHLATHAFFKALLFLGAGSV
IHAMSGEQDINKMGGIWRKIPFTYLMMWIGSLALA--GIYP---------
FAGYYSKDMIIEAAYLSPSESGRFAYWIGISAAFCTAFYSWRLLIKVFHGKFK--NEPTVWKRV----------
---------------------------------------------------
HESPLFMTIPLFILSLGAIFSGIIGEKLHIVD-NTNK---FW--HGAIEIFHSHS-----
HHIPVLYKFMPMGVGILGIVIAVIYYVVF---------------------------------------------
--------------KVLPDITKSTISPLYNLSYYKYYWDEIYEYIVIKPIRCLSVVLWKK-
IDNVIIDGLGPNGVV-KIINLLSTKVTKMQTGYLYHYAYIMLGAVMVLTSWYIFNIIN-------
>WP_041471749_NuoL_Rickettsia_conorii
MYQNICI--MMIIMLPLASSIINGLFL----RVIDKKLAQVIATGFLSLSALFSLIIFCDTGL--DGNI-
IHIKLL-PWIEVGTFKVNWSIYIDQLTSIMFIAVTWVSSIVHIYSLGYMAEDKG-------
IIRFLSFLSLFTFFMLMLVSSDNFLQLFFGWEGVGVCSYLLIGFWYSKESANKAAIKAFIINRASDFAFILGVI
TIIVYCGSANY--KDVLSSAELLSNIK----IFL--
HFSILDIICLLLFIGCMGKSAQIGLHVWLPDAMEGPTPVSALIHAATMVTAGVFLVARCSYLFEYSPLILQFIT
IIGGVTCLFAASIAIMHSDIKKIIAYSTCSQLGYMFMACGVSAYNSGIFHLVTHAFFKALLFLSAGSVIHAVH-
EQDIFKMGDLRNKMPVTYGNFLIGSLALI--GIYP---------LAGFYSKDSILEAAYSS----
GSFMFIFGIAAAILTAIYSMKIIMLVFHGKTK--LEKDVFEHA-------------------------------
------------------------------HEPAKVMNNPLILLVVGSFFSGMIGYYLLAMD-KPNG---YF--
HASLFNLHIYKLL--ISHPPLYIKLLPMAVGIVGIVTGIYLYKSSTV---------------------------
------------------------------
MSFPQKILRSSRGMTPLVLNKYYFDEIYNCLIVKPINCLASLFYL--GDQQIIDRFGPNGFS-
RVVNCFSVLTGKIQTGYVFNYALYIVSFIVVTISYFVWKNIMY------
>AEI88434_NuoL_Candidatus_Midichloria_mitochondrii
MNMYLHNLSLSVVFLPLLASIPVGLNT----NKVKPVIAQIITISFVSIAAIFSWIIFYYTCF--DGQI-
IHLKLI-NWLNLGELKADWSIYIDPLTAIMFLVVNTVSTLVHIYSVSYMSHDPN-------
QPRFFSYLSLFTFFMLVLVSADNFAQLFVGWEGVGLSSYLLIGFWFHKKSAYSAAMKAFLVNRVGDIGLAIGIF
LIAMKFGSVEY--ATVFSKTKYLSEET---
INFLSVDTRLLTVICIALFIGCMGKSAQLGLHTWLPDAMEGPTPVSALIHAATMVTAGVFLLARCSYLFEYSEI
ALDLVTIVGAATCLFAATIATAQNDIKKIIAYSTCSQLGYMFFACGVSAYSAGIFHLLTHAFFKALLFLGAGSI
IHALADEQNIKKMGGIWKKLPYTHALMWIGSIALA--GIPP---------
LSGFFSKDLILEAAYASDSRFGHLAYWLGVAAATLTAFYSCRLLFLVFYAPTN--ADKKIFTEI----------
---------------------------------------------------
HEAPQSMMIPLMVLGIGSLICGYIGYHIIDIS-NLS----FW--
SGAIFVLQEHDSIDKIDTIGHWVAQIPLLAALFAMLLAYYSYILK-----------------------------
------------------------------
PIVSSWSYKNFRNINSFLEKKWYFDEVYEFILIKPVKLLGTSLWRF-VDVGFIDGI-PNYSA-
RAVARFAKVACLLQTGYIYHYAMAMLIGVMILVYWYLPW----------
>WP_038464876_NuoL_andidatus_Paracaedibacter_acanthamoebae
--SVTHLSAMVAVFAPFIGFILSSLGI----RYRGDRFSQWVTCLFMLTATVAGGVLFYHIVF--MQGA-
TVIKLL-DWMNVGGFQAHWGFKLDTLSTTMIMVVNIVSLLVHVYSVGYMSQDKS-------
IPRFMGYLSFFTFTMLMLVTSPNLVQMFFGWEGVGVASYLLIGFWYERQSAGAAAMKAFIVNRVGDVGLALGIA
```

```
SLFLLTGTVEF--
DGALSQITTMAAGQRPELSFWGHSFDAINLIGILLFIGAMGKSAQLGLHTWLPDAMEGPTPVSALIHAATMVTA
GVFLVCRMSPLYEVAPFAREMIVIVGASTAFFAATVALTQTDIKRVIAYSTCSQLGYMFFAAGCSAYGAAMFHL
VTHAFFKALLFLGAGSVIHAMSDEQNMNRMGGIYKLIPTTYALMWIGSLALA--GIPL---------
FAGYYSKDAIIESAYLTGTSIGDYAFAIGLIVAFLTAFYSWRLLLLSFHGKQH--ADDTVMAHV----------
---------------------------------------------------
HESPVVMLLPLFVLSLGAIFSGYLANDWFVGH-KMAE---FW--GSAIAL-PRQP----
HHELPTWVIYSPLAAAVSGIVFAYLFYGGA--------------------------------------------
---------------RRLPQWIATKMSILYNFSYRKWFIDELYDILFVNRAFGLGQILWQK-
GDIGLIDRYGPDGVT-RLSLMVSRFMSYFQTGYVYHYAFAMIVGVLLLLTWYLRGEIG-------
>WP_033444476_NuoL_Candidatus_Odyssella_thessalonicensis
MH----LSAI-AVFAPLLGFILSSLGI----RYLGDRFSQFITCLLMLIAAVAGATLFYEVVF--LRKD-
TVIELL-SWINVGEFQANWGLKLDTLSTTMIGVVNTVSLLVHIYSIGYMSHDKS-------
IARFMSYLSFFTFTMLMLVTAPNLVQMFFGWEGVGVASYLLIGFWYERQSAGAAAMKAFIVNRVGDVGLALGIA
TAFLLTGTVDF--
DGVISTVTQIASAQRPSLNFWGYECDAINLLGILLFIGAMGKSAQLGLHTWLPDAMEGPTPVSALIHAATMVTA
GVFLVCRMSPVYEVAPVAREMIIIVGASTAFFAASIALTQTDIKRVIAYSTCSQLGYMFFAAGCSAYGAAMFHL
VTHAFFKALLFLGAGSVIHAMSDEQDINRMGGIYKLIPTTYVLMWIGSLALA--GIPL---------
FAGYYSKDAIIESAYLTGTGAGDYAFWIGLLVAFLTAFYSWRLLLLAFHGSPK--ADDRVMAHV----------
---------------------------------------------------
HESPFVMLLPLFVLSVGAIFSGYLAHDWFVGT-KMAE---FW--NGAIAL-PHHH----
HSNLPAWVVYSPLVAAVSGIVTAYLFYSKS--------------------------------------------
---------------QTVPQLIASKLSALYTFSYRKWFIDELYNVLFVNRAFGIGQMLWQK-
GDVGFIDRLGPNGIS-RVSLGLSRLVSYLQTGYVYHYAFAMILGVLLLLTWYIKGQMG-------
>MBN8803795_NuoL_Sphingopyxis_terrae
MI-------QAIVFLPLLAALVAGLGQ----RAIGTTAAKLVTTGALFASCALSWPIFLSFVA--
GTGEAGVTPVL-HWVSSGALQFNWELRVDTLTAVMLVVITTVSALVHLYSWGYMSEDPD-------
QPRFFAYLSLFTFAMLMLVTANNLVQMFFGWEGVGLASYLLIGFWYKKPSANAAAIKAFVVNRVGDLGFMLGIF
GTFLVFGTVSI--PEILAAAPGMAGST---
IGFLGHRFDTMTVLCLLLFIGAMGKSAQLGLHTWLPDAMEGPTPVSALIHAATMVTAGVFMVCRLSPMFEAAPV
AAGVVTFVGAATCFFAATVGTTQWDIKRVIAYSTCSQLGYMFFAAGVGAYGAAMFHLFTHAFFKALLFLGAGSV
IHAMHHEQDMRYYGGLRKHIPLTYWAMMMGTLAIT--GVGIAGV----AGFAGFYSKDGILEAAFAA-
GGGGQIAFWVGIFAALLTSFYSWRLVFLTFYGKPRWEQSEHIQHAVHDDHGHGHDD----------------
HAHAPHQ--DAHGDGTAGYHP--------------HESPINMLIPLGVLSLGAVFAGMLFNHQFIYPEEGAA--
-FW--KGSLAF-NEHL-MHAAHEVPLWVKWMPFTVMAIGLFLAWNSYIRN------------------------
-----------------------------------
TTLPARFVAQFSLLHKFLFNKWYFDELYNFLFVKPAFAIGRFFWKR-GDEGTIDRFGPNGVA-
ALVQGGTRLAVRLQSGYVYGYAFVMLLGLVGLASWAMVK-FQ-------
>WP_185664684_NuoL_Novosphingobium_flavum
MHS-----ILFIVFLPLLAAIVAGLGN----KALGNVPAKVITTGALFISCALSWPIFLSFVA--
GSAEASVTPVL-KWVQSGSMSFDWALRVDTLTAVMLVVITSVSALVHLYSWGYMDEEPD-------
QPRFFAYLSLFTFAMLMLVTADNLVQMFFGWEGVGLASYLLIGFWFRKPSACSAAIKAFVVNRVGDLGFMLGIF
GTWLVFQTTSI--PEILHAAPSMAGST---
IGFLGHRFDTMTVLCVLLFIGAMGKSAQLGLHTWLPDAMEGPTPVSALIHAATMVTAGVFMVCRLSPMFVVSPT
AMGVVTFVGAATCFFAATVGTTQWDIKRVIAYSTCSQLGYMFFAAGSGAFGAAMFHLFTHAFFKALLFLGAGSV
IHAMHHEQDMRYYGALRKEIPWTFWAMMAGTLAIT--GVGIVDV----
IGFAGFYSKDSILEAAYASGSEVGSFAFWCGITAALMTSFYSWRLVFLTFYGAPRWAASEHIQHAVHGEHEDAD
AEDGHDH----------DHSHELLD--PHHGEGTAGYHP--------------
```

```
158… HESPLVMLVPLIVLSLGAIFAGFLFTHSFISE-GEGE---FW--KGALVF-NEHL-
…    LHAMHGVPGWVKWAPFTVMAVGFGTAWYAYLKN-----------------------------------------
…    ------------------PRFPAAFVEQFGVVHRFVFNKWYFDELYNFLFVRPAFWLGKVFWKQ-
…    GDVGIIDRFGPNGAA-WVVAQGSRVAQKVQSGYLYSYALVMLIGLVVGISWVMVR----------
159  >WP_147034774_NuoL_Sphingomonas_aquatilis
160  MHA-----ILFIVFLPLLAAAVAGLGN----KAIGFVPAKVITTGALFVSCLLSWPIFLGFLA--
…    GTSTAQVVPVL-TWIVSGDMAVDWSLRVDQLTAIMLVVITSVSALVHLYSWGYMSEDPD-------
…    QPRFFAYLSLFTFAMLMLVTADTLVQMFFGWEGVGLASYLLIGFWFRKPSANAAAIKAFVVNRVGDLGFMLGIF
…    GTFLVFGTVSI--PAILAAAPNMAGST---
…    IGFLWFRADTMTVLCLLLFVGAMGKSAQLGLHTWLPDAMEGPTPVSALIHAATMVTAGVFMVCRLSPMFEMSPT
…    ALHFVTFIGAATCLFAATVGMVQTDIKRVIAYSTCSQLGYMFFAAGVGAYGAAMFHLFTHAFFKALLFLGAGSV
…    IHAMHHEQDMRYYGGLRKSIPLTFWGMMMGTLAIT--GVGLPAVFGNGIGFAGFHSKDAIIEASWAAGEPG---
…    VWFVGVLVALITSFYSWRLMFLTFWGKPRWAASEHIQHALHDAHGHGHDDHAHGHDAHAHDAHHADEAHGNPAQ
…    AEDAGDHPSTHGPAQHDLPAGTGGYHPHESPLPMLIPLIVLSIGAIAAGFVFHGYFIEPSAGES---FW--
…    KGDIAF-REHL-MHAMHEVPTWVKLSATVAMLTGLFTAWYAYIKA-----------------------------
…    ------------------------------
…    PTFPAAVAEQFRVLYRFLTHKYYFDELYNLLFVRPAFAIGRFLWHR-GDEKTIDRFGPNGSA-
…    WIVERSSRIAGRLQNGYVYTYAFVMLIGLTAVVTWAIAQ----------
161  >WP_066575997_NuoL_Sphingomonas_koreensis
162  MSS-----IHFIVFLPLLAAIVAGFSN----RAFGTAFPKLVTTGGLFVAAILSWMTFIPYLS--
…    GTAEPQVVQVL-TWMHSGSLNVDWALRVDSLTAVMLVVVTSVSALVHLYSWSYMEEDPD-------
…    QPRFFAYLSLFTFAMLMLVTADNLLQMFFGWEGVGLASYLLIGFWFKKPSANAAAIKAFVVNRVGDLGFMLGIF
…    GVFLVFGTISI--PEILAAAPSHAGST---
…    IGFLGYRFDTLTVLCLLLFIGAMGKSAQLGLHTWLPDAMEGPTPVSALIHAATMVTAGVFMVCRLSPLFEQSPV
…    ALGFVTFIGAATCLFAATVGTTQTDIKRVIAYSTCSQLGYMFFAAGVGMYGAAMFHLFTHAFFKALLFLGAGSV
…    IHAMHHEQDMRYYGGLRKHIPVTFWTMMAGTLAIT--GVGIYWLH---AGFAGFHSKDAILEAAYAAGGGGQG-
…    AFWVGVFAALLTSFYSWRLMFLTFWGKPRWTQSEHIQHAIHDSHGHDSHGHDDH-----------
…    AHAHDDHHAHADHGDGTGGYKP--------------HESPLPMLVPLLVLSVGAVFAGFAFNKFFIQPEAGEI-
…    --FW--KGAVAF-DEHL-AHAMHEVPLWVKLSATIVMLLGLFGAWVAYIKS-----------------------
…    ------------------------------------
…    PSIPARFTATFGLLYKFLLNKWYFDELYNVLFVKPAFWFGRIFWKK-GDEGTIDRFGPNGSA-
…    AVVKFGSILTGKLQSGYVYSYAFVMLIGLTLAVSWVFTR----------
163  >WP_011445932_NuoL_Novosphingobium_aromaticivorans
164  MSS-----ILIIVFLPLLAAIIAGLGN----KKLGAVPAKVVTTGALFISCALSWPIFLSFLG--
…    GHAEASVTPVL-HWVTSGDLDFSWALRVDTLTAVMLVVITSVSALVHLYSWGYMSEEPD-------
…    QPRFFAYLSLFTFAMLMLVTADNLVQMFFGWEGVGLASYLLIGFWFRKPSANAAAMKAFVVNRVGDLGFMMGIF
…    GTFLVFGTVSI--PAILEAAPGMAGST---
…    IGFLGHRFDTMTVLCLLLFIGAMGKSAQLGLHTWLPDAMEGPTPVSALIHAATMVTAGVFMVCRLSPMFEVSPT
…    ALSFVTFIGAMTCIFAATVGTTQTDIKRVIAYSTCSQLGYMFFAAGVSAYGAAMFHLFTHAFFKALLFLGAGSV
…    IHAMHHEQDMRYYGGLRKQIPLTYWAMMMGTLAIT--GVGIVDL----
…    FGFAGFYSKDAILEAAYAKGTELGHFAFFIGIFAALLTSFYSWRLVFLTFFGKPRWEQSEHIQHAVHDAHGHDD
…    HAHG------HAHGHAHGHDDHAHGHHDEHGDGTAGYHP--------------
…    HESPLNMLVPLGVLSLGAVFAGFVFHHAFVSEGTGH----FW--KGSLVF-SEHL-
…    IHAMHEVPTWVKWAPFAVMATGLLIAWYAYIKN-----------------------------------------
…    ------------------TKFPAAFVEQFAFLYKFLLNKWYFDELYHYIFVVPAMWLGRVFWKV-
…    GDQGIIDRFGPNGAA-WVVAQGSRAAAKLQSGYLYSYALVMLVGLVGAISWVIAQ----------
165  >WP_184002831_NuoL_Stakelama_sediminis
166  MIQ-------LIVFLPLLAAAFAGLSN----RAFGTTLPKLVTTGALFIACVLSWLTFIPFLE--
```

```
GTATSHVAPVL-NWMQSGSLDVAWQLRVDTLTAVMLVVVTTVSALVHLYSWGYMDEDPD-------
QPRFFAYLSLFSFAMLMLVTANNLIQMFFGWEGVGLASYLLIGFWFRKPSANAAAIKAFVVNRVGDLGFMLGIF
GTFLVFGTVSI--PDILAAAPHMAGST---
IGFLWFRADTTTVLCLLLFIGAMGKSAQIGLHTWLPDAMEGPTPVSALIHAATMVTAGVFMVCRLSPLFETSHT
AMAFVTAIGAITCFFAATIGTVQRDIKRVIAYSTCSQLGYMFFAAGVGAYGTAMFHLMTHAFFKALLFLSAGSV
IHAMHHEQDMRYYGGLRKHIPVTFWAMMAGTLAIT--GVGIFG-----IGFAGYYSKDAIIESAYAAG-
PGGYFAYWMGVIAALLTSFYSWRLMFLTFWGKPRWEQSEHIQHALHDAHGHGHDHDH----------------
-AGHDHHDAPEGTGGYKP--------------HESPVVMLIPLIVLSIGAVFAGAAFHHFFSDPGAGER---FW
--KGSLFF-NEHL-AHATEEIPAWAKYGATAAMLLGLIPAWFAYIRS--------------------------
--------------------------------
TDLPAKFADQWRVLYLFLLNKWYFDELYNWLFVRPAFAIGRLLWKR-GDEGTIDRFGPNGSA-
AIVALGSRIAGRLQSGYVYSYAFVMLIGLTAAITWVIAG----------
>WP_176868685_NuoL_Parasphingopyxis_algicola
MIQ-------LIVFLPLLAAIVAGLGN----RMLGNFVAKLLTTGALFVACALSWPIFLDFMI--
GGQQAYVAPVF-DWIRSGDMDVAWSLRVDTLTAVMLVVVTTVSALVHLYSWGYMEDDPD-------
QPRFFAYLSLFTFAMLMLVTSNNLVQMFFGWEGVGLASYLLIGFWYHKPSANAAAIKAFVVNRVGDFGFSLGIF
GTFLVFGTVSI--PEILEAAPAMAGST---
IGFLGYRVDTMTLLCLLLFIGAMGKSAQFGLHTWLPDAMEGPTPVSALIHAATMVTAGVFMVCRLSPMFETSDM
ALAVVTFFGLLTAFFAATVGTAQMDIKRVIAYSTCSQLGYMFVAAGVGAYGIAMFHLFTHAFFKALLFLGAGSV
IHAMHHEQDMRHYGALRKEIPLTFWAMIIGTLSIT--GVGIVGI----
FGFAGFYSKDLIIEAAFASATEMGGIAFFTLVFAALLTSFYSWRLIFLTFFGKPRWITSEHIQHSVHKTPET--
-----------------------------AEDATGGYHP--------------
HESPWSMLIPLGVLSLGAILAGYLFYYPFGYAEQGEI---FW--AGSIAL-DTHL-
LHAIHEVPTWVKYAPGAVMLVGLAIAWLAYIRY-----------------------------------------
------------------TDWPQKFVDQFRLLHAFLYNKWYIDELYDLIFVRPAFAIGRFLWKR-
GDKGFIDRFGPDGVS-ALVASGSGVTRRFQSGYLYTYALVMLIGLAAAATWAITR----------
>WP_072595757_NuoL_Tardibacter_chloracetimidivorans
MIQ-------ALVFLPLVAALIAGLGN----RAIGNVPAKLLTTGALFISCALAWGIFLPMMG--
GQTQPYVAHVL-DFIHSGDLNVAWSLRVDMLTAVMLVVVTTVSALVHLYSWGYMAEDPD-------
QPRFFAYLSLFTFAMLMLVTADNLVQMFFGWEGVGLASYLLIGFWYHKPSANAAAIKAFVVNRVGDFGFSLGIF
GTFLVFGTVSI--PAILAAAPDMAGST---
IGFLGMRMDTMTLLCLLLFIGAMGKSAQLGLHTWLPDAMEGPTPVSALIHAATMVTAGVFMVCRLSPMFETSVT
ATNVVIYVGAATALFAATVGTVQNDIKRVIAYSTCSQLGYMFFAAGVGAYSAAMFHLFTHAFFKALLFLGAGSV
IHAMHHEQDMRFYGGLRRHIPLTFWAMTAGTLAIT--GVGVIGV----
FGFAGFYSKDAIIEAAFAAGGQAGYAAFAVGVFAALLTSFYSWRLVFLTFFGQPRWAASEHIQHAVHGDHHDHP
DAEDAGQETE----------AHSHGHHVHPAKGTAGYHP--------------
HESPWTMLVPLGVLALGAVFAGFIFHHGFIDVEGN-----FW--NGSIAF-DAHL-
MHAMHEVPLWVKLSPAAVMLLGLFLAWNSYIRN-----------------------------------------
------------------PALPGAFVAQFRGLYAFLLNKWYFDELYNVLFVRPALAIGRFFWKK-
GDVGTIDRFGPDGMA-AVVAGGSVAARRLQTGYVYTYALVMLLGIAAAATWAMVR----------
>WP_079638416_NuoL_Sphingopyxis_flava
MIQ-------AIVFLPLLAALVAGLGQ----RFIGTAASKIVTTGALFASCALSWPIFLSFLG--
GDAEASVTPVL-HWVSSGALQFNWELRVDTLTAVMLVVITTVSALVHLYSWGYMEEDPD-------
QPRFFAYLSLFTFAMLMLVTANNLVQMFFGWEGVGLASYLLIGFWYKKPSANAAAIKAFVVNRVGDLGFMLGIF
GTFLVFNTVSI--PEILAAAPGMAGST---
IGFLGQRFDTMTVLCLLLFIGAMGKSAQLGLHTWLPDAMEGPTPVSALIHAATMVTAGVFMVCRLSPMFEVAPI
ALGVVTFVGAATCLFAATVGTTQWDIKRVIAYSTCSQLGYMFFAAGVGAYGAAMFHLFTHAFFKALLFLGAGSV
IHAMHHEQDMRYYGGLRKHIPLTYWAMMAGTLAIT--GVGIAGV----FGFAGFYSKDGILEAAYASGGGGQIA
```

```
-FWVGTFAALLTSFYSWRLVFLTFFGKPRWAASEHIQHAVHGDHHDRPADEHAGGDVHH-------------
AHAADAHEGTAGYHP--------------HESPLSMLIPLGVLSVGAIFAGYLFHHPFIYPEEGMA---FW--
NGSLAF-DAHL-MHAAHEVPTLIKWMPFAAMAIGLLIAWNSYIRN-----------------------------
------------------------------
TALPARFVAQFGLLHQFLFNKWYFDELYNLLFVKPAFAIGRFFWKR-GDEGTIDRFGPDGVA-
ALVQGGTRLAVRLQSGYVYGYAFVMLLGLVGLASWVMVRFL--------
>GGD46181_NuoL_Croceicoccus_pelagius
MIE-------IIVFAPLLAAIIAGLGN----RMIGNMPAKIVTTGALFLSCALSWPIFLGFMT--
GSETAYVDPVL-KWVQSGGMNFDWSLRVDTLTAVMLVVITSVSALVHLYSWGYMDEDPD-------
QPRFFAYLSLFTFAMLMLVTADNLVQMFFGWEGVGLASYLLIGFWFKKPSASAAAIKAFVVNRVGDLGFMMGIF
GVFLVFGTTSI--PEILDRAPEMAGST---
IGFLGMRVDTMTVLCLLLFIGAMGKSAQLGLHTWLPDAMEGPTPVSALIHAATMVTAGVFMVCRLSPLFETSQT
ALTFVTFIGAATCFFAATIGTTQTDIKRVIAYSTCSQLGYMFFAAGVGAYGAAMFHLFTHAFFKALLFLGAGSV
IHAMHHEQDMRYYGGLRKQIPITFWAMMAGTLAIT--GVGIWFLH---
IGFAGFYSKDAILESAFASGSEMGQFAFVMGIVAALLTSFYSWRLMFLTFWGKPRWIESEHIQHTVHKTPEEAG
A------------------------------DTTGGYHP--------------
HESPWVMLTPLVILSLGAVFAGAIWSHSFLDSAE------FW--NGSIFY-NEHL-
IHAMHNVPTWVKIAPGAVMLIGLLIAWNNYIRD-----------------------------------------
------------------PDAPARFVGMFGGIYTFLKNKWYFDELYNVIFVKPAFWLGRKFWKW-
GDVGTIDRFGPDGAA-WVVAQGSRYAQKVQTGYLYSYALVMLLGLLAAISWIMVR----------
>WP_066770280_NuoL_Croceicoccus_mobilis
MIQ-------ILVFLPLLAAIVAGLFN----RQLGNLPAKIITTGALFISCALAWPIFIGFLT--
GSAEAYVAPVL-KWVQSGDMSFDWALRVDTLTAVMLVVITSVSALVHLYSWGYMDEEPD-------
QARFFAYLSLFTFAMLMLVTADNLVQMFFGWEGVGLASYLLIGFWFRKPSANAAAMKAFVVNRVGDLGFMLGIF
GTFLVFGTTSI--SEILSAAPGMAGSS---
ITFMSMRLQTMDILCLLLFVGAMGKSAQLGLHTWLPDAMEGPTPVSALIHAATMVTAGVFMVCRLSPMFEAAPV
ALAFVTFIGAATCFFAATIGTTQTDIKRVIAYSTCSQLGYMFFAAGVGAYNAAMFHLFTHAFFKALLFLGAGSV
IHAMHHEQDMRYYGGLRKKIPLTFWAMMAGTLAIT--GVGLWFAH---
AGFAGFHSKDAILESAFAKGTELSQFAFFMGIVAALLTSFYSWRLMFLTFWGKPRWIESEHIQHSVHKTPEEAG
E------------------------------DTTGGYHP--------------
HESPLVMLIPLIVLSIGAVFAGFAFTHSFLDSSD------FW--DNSLFY-NEHL-
VHAMHEVPTWVKLAPGIVMLIGLAIAWNNYIRD-----------------------------------------
------------------PGAPARFVGQFGFVYTFLKNKWYFDELYHTIFVKPAFWLGRKFWKI-
GDVGIIDRFGPNGAA-WVVAASSRFASKVQTGYLYSYALIMLLGLVAAVSWIIATAGQ-------
>WP_119034702_NuoL_Hephaestia_caeni
GDRVTSTFILLIVFLPLLAAVIAGLGN----RALGNVAAKTVTTGALFVACALSWPIFFSYLA--
GTATATVVPVF-DWINSGSMHIGWALRVDALTAVMLVVVTTVSALVHLYSWGYMEEDPD-------
QPRFFAYLSLFTFAMLMLVTADNIVQMFFGWEGVGLASYLLIGFWFKKPSANAAAIKAFVVNRVGDLGFMLGIF
GTYLVFNTISI--PEILAAAPNMAGST---
IGFLWFRADTMTVLCLLLFVGAMGKSAQLGLHTWLPDAMEGPTPVSALIHAATMVTAGVFMVCRLSPMFEASPT
ALGVVTFIGAATCLFAATIGCVQTDIKRVIAYSTCSQLGYMFFAAGVGAFGGAMFHLFTHAFFKALLFLCAGSV
IVAMHHEQDMRFYGGLRKRIPVTFWAMVAGTLAIT--GVGLPGIGGLAFGFAGFHSKDAVLEASFASGYGGEI-
AWWVGTFAALLTSFYSWRLIFLTFFGKPRWVESEHIQHALHDEHHGAHHGDGAHG-------------------
-HENEAGTGGYRP--------------HESPLSILIPLVLLSVGAVAAGYAFQNYFIAPDAGET---FW--
KGALYH-SEHL-MHAMHEVPVWVKLGPTIVMLIGLYVAWLAYIRK-----------------------------
------------------------------
TDIPAHTAKAFEPVYRFLLNKWYVDELYDLLFVRPAFAIGRLFWKR-GDEGTIDRLGPNGIA-
AVIDGGSRVAGRLQSGYVYTYAFVMLIGLTAAVTWAIAG----------
```

```
>WP_156500484_NuoL_Croceicoccus_bisphenolivorans
MPV-----IKIIVFAPLLASIIAGLGN----RAMGNFAAKVVTTGALFLSCALSWPIFLGFMT--
GAESAYVEPVL-KWVQSGTMTFDWALRVDTLTAVMLVVITSVSALVHLYSWGYMDEDPD-------
QPRFFAYLSLFTFAMLMLVTADNLVQMFFGWEGVGLASYLLIGFWFRKPSASAAAIKAFVVNRVGDLGFMMGIF
GVFLVFGTTSI--PEILDRAPEMAGSS---
IGFLGYRVDTMTVLCILLFIGAMGKSAQLGLHTWLPDAMEGPTPVSALIHAATMVTAGVFMVCRLSPLFETSQM
ALTFVTFIGAATCFFAATIGTTQTDIKRVIAYSTCSQLGYMFFAAGVGAYGAAMFHLFTHAFFKALLFLGAGSV
IHAMHHEQDMRYYGGLRKHIPITFWAMMAGTLAIT--GVGIWFLH---
IGFAGFYSKDAILESAFASGTEMGQFAFFMGIVAALLTSFYSWRLMFLTFWGKPRWEQSEHIQHAVYGHHEEPS
EESDDHHEH----------------AHASHGDGTAGYHP--------------
HESPWVMLTPLVVLSLGAVLAGAIWSHSFLDSAE------FW--NGSIFY-NEHL-
IHAMHNVPTWVKIAPGAAMLIGLAIAWNNYIRD-----------------------------------------
------------------PGAAGRFVGMFGGVYTFLKNKWYFDEVYHTIFVKPAFWLGRKFWKW-
GDEGTIDRFGPNGAA-WVVAQGSRYAQKVQTGYLYSYALVMLLGLVAAISWVMANGLGH------
>RXR30042_NuoL_Sphingobium_fluviale
--------MLLIVFLPLLAAIIAGLGN----KALGKLPAKLITTGALFVSCALSWPIFLSYVA--
GDAGPTVVPVL-HWLSSGSFEAAWELRVDALTAVMLVVVTSVSALVHLYSWGYMDEEPD-------
QPRFFAYLSLFTFAMLMLVTANNLLQMFFGWEGVGLASYLLIGFWFRKPSASAAAIKAFVVNRVGDLGFMLGIF
GTYLVFDTISI--PDILAAAPGMAGST---
IGFLGYRFDTMTVLCLLLFVGACGKSAQLGLHTWLPDAMEGPTPVSALIHAATMVTAGVFMVCRLSPMFETSET
ALAVVTYVGAATCLFAATVGTTQNDIKRVIAYSTCSQLGYMFFAAGVGAYGAAMFHLFTHAFFKALLFLGAGSV
IHAMHHEQDMRYYGALRKEIPITFWAMTAGTLAIT--GVGIPGTL---FGFAGFFSKDSIIESAYAAGGPGTT-
AFLIGITAALLTSFYSWRLVFLTFFGKARWAGSEHIQHAVHDAHGHDSHGHDDHAHD-------------
DHHGHDAHAIGTAGYHP--------------HESPLVMLIPLIVLSIGAVAAGFVFHHAFIDAEGGAQ---FW-
-APSTLAFDAHL-MHASHEVPLWVKWSPFAVMATGLMIAYMAYIRH----------------------------
-------------------------------
TDWPAKFTAQFHVLYDFLLNKWYFDELYDMLFVKPAFAIGRFFWKR-GDEGTIDRFGPNGMA-
ALVVAGGRLTRRLQSGYLYTYALVMLLGLAAAVTWAITVS---------
>WP_046902666_NuoL_Altererythrobacter_atlanticus
MSS-----VLVIVFLPLLAAIIAGFGN----RAIGNVAAKVVTTGALLIACALSWPIFITFLA--
GDVPTTVVPVL-KWVQSGALTFDWALRVDTLTAIMLVVVTSVSALVHLYSWGYMSEDPD-------
QPRFFAYLSLFTFAMLMLVTADNLVQMFFGWEGVGLASYLLIGFWFRKPSASAAAIKAFVVNRVGDLGFMLGIF
GTFLVFQTTSI--AEILEAAPAMKGAS--
TIGFLGMQLDTMTIICLLLFVGAMGKSAQLGLHTWLPDAMEGPTPVSALIHAATMVTAGVFMVCRLSPMFEAAP
AALTFVTFVGGATCFFAATVGTTQWDIKRVIAYSTCSQLGYMFFAAGVGAYGAAMFHLFTHAFFKALLFLGAGS
VIHAMHHEQDMRYYGGLRKHIPITFWAMMAGTLAIT--GVGIYWLH---
AGFAGFHSKDAILEAAFASGTEMGRTAFWLGAFAALLTSFYSWRLMFLTFWGKPRWAESEHIQHAVHHGHAEPN
DANPPSQED-------AGHDVAHHVPSNDHEDGTAGYHP--------------
HESPWVMLVPLIVLSFGAVFAGYIFSPSFIESEA------FW--NGSIAF-NEHL-
IHSMHEVPLWVKLSASIVMLIGLAIAWYAYIRN-----------------------------------------
------------------TSVPQKAAAQLGPIYRFLFNKWYFDELYNFIFVKPAFWLGRQFWQR-
GDVGLIDRFGPNGAA-WVVEKGSVLAHKIQSGYLYSYALVMLIGLVAAISWVLV-----------
>WP_205441185_NuoL_Tsuneonella_flava
MHP-----ILLIVFLPLLAAVIGGLGN----KALGNLAVKSLTTALLFVSCGLSWPIFIGFLT--
GAETASVIPVL-QWVHSGSMTFDWALRVDTLTAVMLVVVTTVSALVHLYSWGYMADEPD-------
QPRFFAYLSLFTFAMLMLVTADNLVQMFFGWEGVGLASYLLIGFWFRKPSASAAAIKAFVVNRVGDFGFMLGIF
GTYLVFQTVSI--PEILAAAPGMAGST---
IGFLGYRVDTMDLLCILLFIGAMGKSAQLGLHTWLPDAMEGPTPVSALIHAATMVTAGVFMVCRLSPMFEAAPA
```

```
ALMFMTFVGAATCFFAATIGTTQWDIKRVIAYSTCSQLGYMFFAAGVGAYNVAMFHLFTHAFFKALLFLCAGSV
IHAMHHEQDMRYYGGLRKHIPITFWAMLAGTLAIT--GVGVAGV----
FGFAGFYSKDAILEAAFARGTNLGNLAFWMGTIAALLTSFYSWRLVFLTFYGKPRWIESEHIQHSVHKTPEQAG
EDT------------------------------TGGYHP--------------
HESPLPMLVPLLALSVGAVFAGFVFAPAFVESAE------FW--QGSVFF-NEGL-
MHAMHGIPAWAKWSATVAMLIGLAIAFQGYIRD-----------------------------------------
------------------KELPARIAGQLGPVYRFVYNKWYFDELYHFLFVKPAFWLGNKFWKW-
GDIGIIDRFGPNGAA-WLVSQGTAAAKKVQSGYLYSYALVMLIGVVVAISWVMVR----------
>WP_120110433_NuoL_Tsuneonella_suprasediminis
MHP-----ILLIVFLPLLAAVIGGLGN----KALGNLAVKSLTTAFLFVSCGLSWPIFIGFLT--
GAETASVIPVL-QWVHSGSMTFDWALRVDTLTAVMLVVVTTVSALVHLYSWGYMADEPD-------
QPRFFAYLSLFTFAMLMLVTADNLVQMFFGWEGVGLASYLLIGFWFRKPSASAAAIKAFVVNRVGDFGFMLGIF
GTYLVFQTVSI--PEILAAAPGMAGST---
IGFLGHRVDTMDLLCILLFIGAMGKSAQLGLHTWLPDAMEGPTPVSALIHAATMVTAGVFMVCRLSPMFEAAPA
ALMFMTFVGAATCFFAATIGTTQWDIKRVIAYSTCSQLGYMFFAAGVGAYNVAMFHLFTHAFFKALLFLCAGSV
IHAMHHEQDMRYYGGLRKHIPITFWAMLAGTLAIT--GVGLAGV----
FGFAGFYSKDAILEAAFARGTNLGNLAFWMGTIAALLTSFYSWRLVFLTFYGKPRWIESEHIQHSVHKTAEQAG
EDT------------------------------TGGYHP--------------
HESPLPMLVPLLALSVGAVFAGFVFAPAFVESAE------FW--QGSVFF-NEDL-
MHAMHGIPAWAKWSATVAMLIGLAIAFQGYIRD-----------------------------------------
------------------KELPARIAGQLGPVYRFVYNKWYFDELYHFLFVKPAFWLGNKFWKW-
GDIGIIDRFGPNGAA-WLVTQGTAAAKKVQSGYLYSYALVMLIGVVVAISWVMVR----------
>MXP14217_NuoL_Pseudopontixanthobacter_confluentis
MSS-----ILIIVFLPLLAIVAGLGN----RALGNTVSKSITTGALFISCALSWPIFLGFVA--
GSAEATVVPVL-QWVQSGDLSFDWALRVDTLTAIMLVVITTVSALVHLYSWGYMEEDPD-------
QPRFFAYLSLFTFAMLMLVTADNLVQMFFGWEGVGLASYLLIGFWFRKPSANAAAIKAFVVNRVGDLGFMLGIF
GTFLVFGTVSI--PEILEAAPSMSGAT---
IGFLGYRMQTMDILCILLFIGAMGKSAQLGLHTWLPDAMEGPTPVSALIHAATMVTAGVFMVCRLSPMFETAPV
ALGFVTFIGAATCLFAATVGTTQWDIKRVIAYSTCSQLGYMFFAAGVGAYGAAMFHLFTHAFFKALLFLGAGSV
IHSMHHEQDMRYYGGLRKKIPLTFFAMLAGTLAIT--GVGIYWLH---
AGFAGFHSKDAILEVAFARGTELSQFAFWVGCFAALLTSFYSWRLMFLTFWGKPRWIESEHIQHSVHKTPAEAG
EDT------------------------------TGGYHP--------------
HESPLSMLVPLGILSIGAVFAGFVFNPQFLDSAE------FW--AGSIYY--ENL-
IHAMHAVPLWVKLSATIVMLLGVLIAWMAYIRD-----------------------------------------
------------------TSIPGKFVEQFRNLHNFVYNKWYFDELYNLVFVRPAFWLGRKFWKI-
GDEGIIDRFGPNGAA-WLVSKGTVAAQKVQSGYLYSYALVMLLGLVAAVTWVLM-----------
>WP_130586631_NuoL_Qipengyuania_flava
MHP-----ILLIVFLPLLALVAGLTN----KAAPSVFAKAITTGALFVSAALSWPIFLGFVA--
GTYEPTVVPVL-KWVQSGSLSFDWALRVDTLTAIMLVVINTVSALVHLYSWGYMDEDPD-------
QPRFFAYLSLFTFAMLMLVTADNLVQMFFGWEGVGLASYLLIGFWFRKPSASAAAIKAFVVNRVGDLGFMLGIF
GTYLVFDTVSI--TEILAAAPGMSGAT---
IGFLGYNVYTMDVLCVLLFIGAMGKSAQLGLHTWLPDAMEGPTPVSALIHAATMVTAGVFMVCRLSPMFETAPV
ALGLVTFIGAATCIFAATVGTTQWDIKRVIAYSTCSQLGYMFFAAGVGAYGAAMFHLFTHAFFKALLFLGAGSV
IHAMHHEQDMRYYGELRKHIPLTFWAMMAGTLAIT--GVGIYHLG---
AGFAGFWSKDAILEVAYGRGTELGNFAFWMGTFAALLTSFYSWRLMFLTFWGKPRWIESEHIQHSVHKTPEEAG
ADT------------------------------TGGYHP--------------
HESPLTMLIPLGVLSIGAVLAGQVFAPTFLDDAA------FW--GSSIFY-NEPL-
IHAMHNVPYLVKYAALIVMVIGLVVAWYAYIKD-----------------------------------------
```

```
------------------TSIPAKTAEQLGPIYRFLYNKWYFDELYHYLFVVPAFWLGRQFWKI-
GDVGTIDRFGPNGIA-WVVEKGSVGARKFQTGYLYSYALVMLLGLVAAITWVLF-----------
>WP_120323177_NuoL_Altererythrobacter_spongiae
MSS-----ILIIVFLPLLASAIAGLGN----KALGNVPAKVITTGALFISCALSWPIFLSYLA--
GTAEPSVVPVL-MWVQSGDFAFDWALRVDTMTAIMLVVITSVSALVHLYSWGYMDDEPD-------
QPRFFAYLCLFTFAMLMLVTADNLVQMFFGWEGVGLASYLLIGFWFRKPSAGAAAIKAFVVNRVGDLGFMLGIF
GTFLVFGTVSI--PEILAAAPGMSGAT-----
FLGMEFHTMTILTLLLFIGAMGKSAQLGLHTWLPDAMEGPTPVSALIHAATMVTAGVFMVCRLSPMFQAAPDTM
TFVTLIGGLTCLFAATIGTTQWDIKRVIAYSTCSQLGYMFFAAGVGAYGVAMFHLFTHAFFKALLFLGAGSVIH
SMHHEQDMRYYGALRKQIPVTFWAMMAGTLAIT--GVGVYWLH---
AGFAGYHSKDAILEAAFASGTEMGRTAFWLGAIAALLTSFYSWRLMFLTFWGKPRWSQSEHIQAVHHGHDEPE
EANPARQED-------SGHDVSHHVPSPEEDDGTAGYHP--------------
HESGWVMLVPLILLSVGAVFAGFVFSPYFVESES------FW--QGSIVY-NEHL-
MHEMHGIPTWVKLSASIAMLIGLAVAWFAYIKD-----------------------------------------
------------------TSIPGKFVNQFRQVHAFLYNKWYFDELYNLIFVRPAFWLGKVFWKQ-
GDVGLIDRFGPNGAA-WVVERGAVVAKKIQSGYLYSYALIMLLGLIAAISWVMVR----------
>WP_170002558_NuoL_Pseudopontixanthobacter_vadosimaris
MSTS----ILIVVFLPLLAAIIGGLGN----RSLGNVAVKAITTGALFISCGLSWPIFLGFLS--
GETIAAVVPVL-DWVRSGDMEFGWALRVDTLTAVMLVVITTVSALVHLYSWGYMDEDPD-------
QPRFFAYLSLFTFAMLMLVTADNLVQMFFGWEGVGLASYLLIGFWFRKPSANSAAMKAFIVNRVGDLGFMLGIF
GTFLVFGTTSI--PVILEAAPAMSGAS---
VGFLGSRVMVMDVLCILLFIGAMGKSAQLGLHTWLPDAMEGPTPVSALIHAATMVTAGVFMVCRLSPMFETAPV
ALAFVTFIGAATCLFAATVGTVQWDIKRVIAYSTCSQLGYMFFAAGVGAYGAAMFHLFTHAFFKALLFLGAGSV
IHAMHHEQDMRYYGALRKKIPLTFWAMMAGTLAIT--GVGIYWLH---
AGFAGFHSKDAILEAAFGRGTELGQFAFWLGAFSALLTSFYSWRLMFLTFWGKPRWAQSEHIQAVHHGHDEPD
EANPARQEN-------AGHVAMHDVPSPEQAEGTAGYHP--------------
HESPVSMLIPLAVLSLGAIFAGWLFSEAFLDSAE------FW--NGSIYY-NDGL-
IHAMHATPLWVKLTATIVMLAGLLIAWFAYIRD-----------------------------------------
------------------TSIPGKFVEQFGLLHNFLFNKWYFDELYDRIFVRPAFWFGRKFWKL-
GDEGGIDRFGPDGAA-WLVAQGSAGARRFQSGYLYSYALIMLLGVVAAITWVLM-----------
>WP_202391155_NuoL_Pseudopontixanthobacter_sediminis
MP--------------LLAAIVGGLGN----RALGNLPVKLITTGALFVSCALSWPIFLGFIA--
GDAQATVVPVL-QWVQSGSLDFDWALRVDTLTAVMLVVITTVSALVHLYSWGYMDEDPD-------
QPRFFAYLSLFTFAMLMLVTADNLVQMFFGWEGVGLASYLLIGFWFRKPSANAAAIKAFVVNRVGDLGFMLGIF
GTFWVFGTVSI--PEILAAAPAMSGST---
IGFLGYRMQTMDILCILLFIGAMGKSAQLGLHTWLPDAMEGPTPVSALIHAATMVTAGVFMVCRLSPMFETAPI
ALGMVTFVGAATCIFAATIGTTQWDIKRVIAYSTCSQLGYMFFAAGVGAYGAAMFHLFTHAFFKALLFLGAGSV
IHAMHHEQDMRYYGGLRKRIPLTFFAMLAGTLAIT--GVGIYWLH---
AGFAGFHSKDAILEVAFARGTEMGQFAFWMGAFAALLTSFYSWRLMFLTFWGKPRWAESEHIQAVHHGHDDPE
AHNPPVQED-------AGHDAVHAVPSADQGDGTAGYHP--------------
HESPISMLVPLGVLTLGAVFAGFIFNPQFLDSAE------FW--GGSIYY-NENL-
IHAMHAVPLWVKLTASIVMLLGLFGAWLAYIRD-----------------------------------------
------------------TSIPGKFVEQFRLVHRFVYNKWYFDELYDLIFVRPAFWFGRKLWKI-
GDEGIIDRFGPNGAA-WLVLKGTGAAKRVQSGYLTSYALIMLLGLIAAVTWVLM-----------
>WP_176266180_NuoL_Actirhodobacter_atriluteus
MI-------TLIVFLPLLAAIIAGFGN----RALGNTLAKSVTTGALSIACALSWPIFLGFLS--
GTMEASVVQVL-PWVQSGTLTFDWSLRVDTLTAVMLVVVTTVSALVHLYSWGYMDEDPD-------
QPRFFAYLSLFTFAMLMLVTANNLVQMFFGWEGVGLASYLLIGFWFRKPSANAAAIKAFVVNRVGDLGFMLGIF
```

```
GTYLVFQTTSI--PEILEAAPAMSGAT---
IGFLGYRVATMDVLCILLFIGAMGKSAQLGLHTWLPDAMEGPTPVSALIHAATMVTAGVFMVCRLSPMFETAPV
ALDFVTFIGAATCIFAATVGTTQWDIKRVIAYSTCSQLGYMFFAAGVGAYGAAMFHLFTHAFFKALLFLGAGSV
IHAMHHEQDMRYYGALRKEIPFTFWAMLAGTLAIT--GVGVYHLG---
VGFAGFWSKDAILEVAYARGTDLGNFAFWMGTFAALLTSFYSWRLMFLTFWGKPRWADSEHIQHAVHHGHDEPE
EHNPAIQED-------AGHDVTHTVPSPNYSAGTGGYHP--------------
HESPISMLIPLGVLSLGAVLAGQLFAPAFLDSAA------FW--NGSIFY-NEPL-
IHAMHAVPTLVKYAAFIVMVLGLAAYLAYIKD----------------------------------------
------------------TSLPARGAEQLGPVYRFFYNKWYFDELYRMLFIIPAFAFGKAFWRF-
VDKGMIDRFGPDGAA-WIVTKGSAAAKRVQTGYVYSYALVMLLGLVAAITWVLF-----------
>WP_066965327_NuoL_Rhizorhabdus_dicambivorans
MI-------TLIVFLPLLAAIVAGLGN----RLIGNVPAKLVTTGALFASCAMSWPVFMGFMT--
GELTAHVHPVL-TFIQSGDLSVDWALRVDTLTAVMLVVVTSVSSLVHLYSWGYMAEDPS-------
QPRFFAYLSLFTFAMLMLVTSDSLVQMFFGWEGVGLASYLLIGFWYHKPSANAAAIKAFVVNRVGDFGFSLGIF
GTFLVFGTVSI--PEILAAAPGYANAS---
IGFLGARVDLMTLLCLLLFVGAMGKSAQLGLHTWLPDAMEGPTPVSALIHAATMVTAGVFMVCRLSPMFEVSET
AMHVVTYVGAATALFAATVGTTQTDIKRVIAYSTCSQLGYMFMAAGVGAYGAAMFHLFTHAFFKALLFLGAGSV
IHAMHHEQDMRYYGGLRKHIPFTYWAMMAGTLAIT--GVGIVG-V---
AGFAGFHSKDAIIEAAFANGSSSGGVAYAVAVFAALLTSFYSWRLVFLTFYGKPRWAGSEHIQHAVHDAHGHDH
HDEPAQEDA------GHDPHAHAAHDHGHPAEGDAGYHP--------------
HESPATMLIPLAVLSVGAIFAGFLWAPYFIDSHHGGA---FW--AGSLAF-DEHL-
MHAMHEVPWWVKWSATIVMITGFLIALWAYVLD----------------------------------------
------------------RTVPARFTAQFSVLYDFLLRKWYFDELYHVLFVVPAFWLGRFFWKK-
GDQGTIDRFGPNGIA-ALVSGSGAWARRAQTGYLYTYALVMLLGLAAAATWAMVG----------
>WP_183933181_NuoL_Sphingomicrobium_lutaoense
MQT----SIILIVFLPLLAAIVAGLGG----RIIGKFASKLVTTGALFLSCILSWPIFIGFLT--
GAETETVVPVL-DWIRSGDMVVDWALRVDTLTAVMLVVVTTVSSLVHLYSWGYMEEDPS-------
QPRFFSYLSLFTFAMLMLVTADSLVQMFFGWEGVGLASYLLIGFWYHKPSANAAAMKAFVVNRVGDFGFSLGIF
GVFLVFGTVSI--PAILEAAPGAVGTE---
IGFAGMRVDTLTLLCLLLFVGAMGKSAQLGLHTWLPDAMEGPTPVSALIHAATMVTAGVFMVCRLSPLFDAAPG
ALEVVTYVGAATALFAATVGTVQNDIKRVIAYSTCSQLGYMFFAAGVGAYGAAMFHLFTHAFFKALLFLGAGSV
IHAMHHEQDMRYYGALRKEIPLTFWAMIFGTLAIT--GVGIIG-
IFGGFYSKDAIIESAYAAAVMGGGNVSAGFAFAIALIAALLTSFYSWRLIFLTFYGKARWASSEHVQHALHGHH
EDESEVVADHDSA--------------DDHPVAVREGTGGYHP--------------
HESPWSMLVPLGALTIGAIFAGYAFYYPFFGTEEGAA---FW--AGSLVH-NAEL-
VEAAHHVPLWVKLSPALVMLIGLGIAYNNYIRK----------------------------------------
------------------PKNPAKFVAMFGGLHTFLMHKWYFDELYNLIFVKPAMWIGRIFWKR-
GDEQTIDRFGPHGAA-TAVGWGNRLTARLQSGYLYSYALVMLLGLIAAASWAYWWAR--------
>WP_108985294_NuoL_Can.Phycosocius_bacilliformis
ASGLAETLAHVVVLAPLLGALICGLFN----RFIGEKVAMFVATALLFVAAACAATIFIGHWN-
HSLHLPAKPIRLATWIDVGAFKSTWSIRLDAISAVMMIVVTGVSSLVHLYSWGYMAEDPH-------
KPRFFAYLSLFTFAMLSLVTAADFMQLFFGWEGVGLASYLLIGFWYHKRSANDAQIKAFVVNRVGDFGFALGIM
AVFFVFGSIEF--KQVFDQIPQKADMM---
LAWGVFHLPAIEIIAFLLFIGAMGKSAQFFLHTWLPDAMEGPTPVSALIHAATMVTAGVVLVCLCSPIYEAAPA
TAGFITVIGAVTALFAATVGIAQNDIKRVIAYSTCSQLGFMFFAAGIGAYQAAMFHLFTHAFFKALLFLGAGSV
IHGMHHEQDMRKMGDVARHMKITFVIMLIGTLAIT--GVGIPHTPFG---
FAGFFSKDAILETTFAAMTGNTQLAFWMALIAATLTSFYSWRLAFMTFHGTPKWKADADHGHDHDHGAGHETSD
SA--HAHAQPAVAHVGEVAHDAAGSHDDHAHHAWHGP--------------
```

```
HESPLVMLIPLIILAFGAVFAGSIFYDAFVGH-YAKE---FW--
GNAIYTAPDNNVLKDKYSVPTWVFYAPLVVMLIGLAAALYFYLFN-----------------------------
------------------------------
EGLGKKIADRKGLAHAFLENKWYFDELYEAVFVKGARALGDLFWKI-GDGKLIDGLGPNGIA-
AALNFGSKKAVKIQSGYVYHYAFLMLIAAVALGAFILYR-GA-------
>VDC50122_NuoL_Brevundimonas_Caulobacterales
MNL--HTLIILGIFAPLLGATVAGLFG----RRIGDIPSQTLTTGLLFFSCAVAWTVF-GQWTW-
GHLEPFTIRLA-PFINVGDFQSAWSIRIDALSATMLIVVTSVSSLVHLYSWGYMAEDDS-------
RPRFFAYLSLFTFMMLALVTAADFMQLFFGWEGVGLASYLLIGFWFKKPTASAAAIKAFVVNRVGDFGFVLGII
TIFWMYGTIEF--AELFPLVATKAGTT---
WEFLGVQWSALDLAGFLLFIGAMGKSAQFFLHTWLPDAMEGPTPVSALIHAATMVTAGVYMVCLLSPIYEYAPG
ASQIIAIIGAITALFAATVGLTQNDIKRVIAYSTCSQLGYMFFAAGVGAYQAAMFHLFTHAFFKALLFLGAGSV
IHGMHHEQDMRKMGGLWKLLPITYAVMTIGTIAITGLGIPGV-GG-----
FAGFYSKDSIIESAFASSGHSAMFAFSIGLIAAGLTAYYSWRLIFMTFHNKPVW--KEE-GDAHHADDHA-
SHAQLETHSE------------PVSDAHAHDDHAHDDHGHHGPLQP--------
HESPWVMLVPLILLSVGAVAAGFVFAPHFIGH-HEHE---FW--
RGAIFTGEHNHVLHESHDVPTWVKWSPLILTLTGTAFAYWIYVAR-----------------------------
------------------------------
EGMGRRMAERGGFLYNFLYNKWYFDELYDFVFVRGFKAVGDVFWKI-VDVKIIDGLGPNGAA-
WASLKSAARLGKLQSGFVYHYAFVMLLGVAGLLTFAILAWGA-------
>WP_004615182_NuoL_Caulobacter_vibrioides
MQT----LVTILVFAPLVGALIAGLFG----RRIGDVASQAVTTGLLILACALSWYTF-SQWTW-
GGLEAFTVRLL-PFIHIGDFQANWSIRIDALSATMLIVVTTVSALVHIYSWGYMAEDDS-------
KPRFFAYLSLFTFAMLSLVTAADFMQLFFGWEGVGLASYLLIGFWFKKPSASAAAIKAFVVNRVGDFGFALGIM
TTFWAFGSIQF--AEIFPQIAAHAGKT---
WVFAGHTFPLMDIACFLLFIGAMGKSAQFFLHTWLPDAMEGPTPVSALIHAATMVTAGVYMLCLLSPMFEYAPI
AKNIVTVIGAVTALFAATVGLTQNDIKRVIAYSTCSQLGYMFFAAGVGAYQAAMFHLFTHAFFKALLFLGAGSV
IHGMHHEQDMRKYGALAKLLPITFIAMTIGTIAITGLGIPPLELG-----
FAGFYSKDTIIEAAYAAGQHNPMFAWVIGVLVAGLTSFYSWRLAFFTFNGKARWGHDDH--
HAHAHDAHGHDAHADETHDE------------PLPD---DDHGHGHDHKP--------------
HESPWVMLFPLVVLSIGAVAAGFVFTGYFVGH-HQEE---FW--
RGAIYNAPTNHVLHEAHGVAEWVKYSPLIATILGLLIAAYVYLLKGD---------------------------
------------------------------
ERLGLKLAERKGPLYVFFYNKWFFDELYDATFVRLAKFLGDLFWKGGGDQKIIDGLGPDGVS-
AVSYEVGKRTGKLQTGYLYHYAFVMLLGVAGLLTYALFKFH--------
>WP_014891219_NuoL_Methylocystis_sp._SC2
MI-------YAIVFLPLVGFLIAGALG----PWIGARASELVTTGLLLICAVLSWIVFFDVAL--GHDQ-
GYAPIIGNWMTVGDLKVDWALRVDTLTAVMLVVVNTVSSLVHLYSIGYMHEDPD-------
RPRFFAYLSLFTFAMLMLVTADNLVQMFFGWEGVGLASYLLIGFWYQKPSANAAAIKAFVVNRVGDFGFALGIF
LVFQLTKSLGF--EEVFAAVPGLAGKT---
IHVFGMDVDALTLATFLLFIGAMGKSAQFLLHTWLPDAMEGPTPVSALIHAATMVTAGVFMVARLSPIFEYAPA
TLEFVTLVGAVTAVFAATVGLVQNDIKRVIAYSTCSQLGYMFVAEGVGAYSIGVYHLFTHAFFKALLFLGAGSV
IHAMHHEQDMRNMGGLRKDIPFTFAMMIIGTLALT--GFPF---------
TAGFYSKDAIIEAAFAASEHHHMFAFVATVAAAGLTSFYSWRLVFMTFFGARK---DHA-
VQTDHDKTTAAASAHADGDHA------------HSHDD--GHGHGHGHHTP--------------
HESPVVMLAPLAVLGLGALGAGIVFGKYFIGH-DYDE---FW--
KGALFTGRDNHIIHEFHDVALGVGLAPTVVMVLGFAVALYFYVLR-----------------------------
------------------------------
```

```
PGTAQALARAFPRLYRFLLNKWYFDELYDFLFVRPAFALGRLFWKGGGDGAIIDGLGPDGVA-
ARVADGARLAVRLQTGYVYHYAFAMLIGVAAVVTYFVAGGLR-------
>WP_102843351_NuoL_Methylocella_silvestris
MY-------AAILFLPLIGFLIAGPFG----RQLGARPSELVTTSLLFVAALLSWIAFANVAL--GEEP-
GSVALLGEWFSSGALRAEWTVRVDSLTAVMLVVVTTVSALVHLYSIGYMSDDPS-------
RPRFFSYLSLFTFAMLALVTADNLLQMFFGWEGVGLASYLLIGFWYQKPSANAAAIKAFVVNRVGDFGFLLGIF
MVFVLTRSINF--EQIFAAAPGLANTT---
IHVFGAEWDAMTITCLLLFMGAMGKSAQFLLHTWLPDAMEGPTPVSALIHAATMVTAGVFMVARLSPLFEQAPH
ALSFVIFIGATTAFFAATVGLVQNDIKRVIAYSTCSQLGYMFVALGVGGYSIGIFHLFTHAFFKALLFLGAGSV
IVAMHHEQDMRNMGGLWRKIPFTFAMMTIGTLALT--GFPY---------
TSGYFSKDAIIEAAYASHRPMAFYGYLMTVIAAALTSFYSWRLVFLTFFGKAQW-
ADEGHGAAAHSVGAAAVAGHAPDAHA------------ADAH---DDHGHHGPLNP--------------
HEAPIVMLIPLAVLAVGALFAGLIFHSDFIGE-GFNE---FW--
KGSLFLGPDNHILHEMEEIPHLAALMPTIMMVAGFLIALYMYILA-----------------------------
------------------------------
PATPARLAAAMPALYRFLLNKWYFDELYDKIFVRPAFWLGNLFWRGGGDGAIIDRLGPDGVA-
ARVVDVTGRVVRLQSGYIYHYAFAMLIGLAAIITWYIVGGVR-------
>WP_102958287_NuoL_Mangrovicella_endophytica
MY-------QAIVLLPLVGFLIAGLFG----RSIGAKASEYVTSSLLIVAAVLSWIAFISFGF--
GEGETLRVTML-RWMQVGSLDIDWSLRIDRLTLVMLVVVNTVSALVHVYSIGYMHHDPH-------
RPRFFAYLSLFTFAMLMLVTSNNLVQMFFGWEGVGLASYLLIGFWYKKPSASAAAMKAFIVNRVGDFGFALGIF
GLFVLFGSVNF--
DTIFAGAADVAGRMLVFAGYSLSMAGALTVICLLLFMGAMGKSAQLGLHTWLPDAMEGPTPVSALIHAATMVTA
GVFMLARMSPVFELSHEALTVVTFVGATTAFFAATVGLVQNDIKRVIAYSTCSQLGYMFVALGVGAYGAAIFHL
FTHAFFKALLFLGAGSVIHAVSDEQDMRNMGGLRKHIPITYWMMVIGTLALTG--FPF---------
TAGYFSKDAIIESAYAGHNSFAAYGFGMTVIAALLTSFYSWRLIFMTFHGKPR--ASHEVMHHV----------
---------------------------------------------------
HESPMVMLVPLFVLAAGALLAGLLFEPYFLGE-GYEE---FW--
VSALYSGAENHVLHDMHATPFLIGLLPTIMMVIGLVLAWLFYIRS-----------------------------
------------------------------
PQTPARIAERHSGLYKFLLNKWYFDELYDVLFVRSAKALGRFLWKK-GDVGTIDRLGPDGIS-
ARVLDVTDRVVRLQTGYLYHYAFAMLIGVAALVTWMMFGGAR-------
>WP_090676648_NuoL_Aureimonas_jatrophae
MY-------TLIVLLPLIGFLVAGLFG----NAIGAKNSEYVTSGLLIVAAVLSWIAFLIGV---DGVESIKVL
---QWIKAGTLDVSWSLRIDRLTLVMLVVVNTVSALVHVYSIGYMHHDDS-------
RPRFFAYLSLFTFAMLTLVTADNLVQMFFGWEGVGLASYLLIGFWYQKPSANAAAMKAFVVNRVGDFGFALGIF
GLFVVFGSVNL--
TDIFAGAGQMVAQLLQFAGYSLSASGAMTAICLLLFMGAMGKSAQIGLHTWLPDAMEGPTPVSALIHAATMVTA
GVFMLARMSPVFELSHSALTFVTFIGATTAFFAATVGLVQNDIKRVIAYSTCSQLGYMFVALGVGAYGAAVFHL
FTHAFFKALLFLGAGSVIHAVDGEQDMRHMGGLRKHIPVTYWTMMIGTLALTG--FPF---------
TAGYYSKDAIIESAFAGENAFAMYGYIMTVIAAALTSFYSWRLVFLTFHGKPR--ASHEVMHHV----------
---------------------------------------------------
HESPQVMLVPLYILSAGALLAGIVFYPFFLGG-SYDT---FW--
NGALFSGPQNHILHESHEVPGLVVWSPTIAMALGFIVAYIFYVRS-----------------------------
------------------------------
PEKPAQLAARNSGLYQFLLNKWYFDELYDFLFVRSAKWLGRFLWKK-GDGAVIDRLGPDGVS-
ARVLDVTNRVVQLQTGYLYHYAFVMLIGVAALVTFMMFGGMR-------
>WP_090959385_NuoL_Aureimonas_phyllosphaerae
```

```
MY-------TLIVLLPLVGFIVAGLFG----NRIGAKASEYVTSGLLIAAALLSWIGFLSFGG-----
ETQHVQVL-QWMKAGTLDISWQLRIDRLTLVMLVVVNTVSALVHVYSIGYMHHDDS-------
RPRFFAYLSLFTFAMLTLVTADNLVQMFFGWEGVGLASYLLIGFWYQKPSANAAAMKAFVVNRVGDFGFLLGIF
GLFMVFGSANL--
DTIFAGAGQMVGTILNFAGYSLPAAGAMTAICLLLFMGAMGKSAQIGLHTWLPDAMEGPTPVSALIHAATMVTA
GVFMLARLSPVFELSHSALTFVTFIGATTAFFAATIGLVQNDIKRVIAYSTCSQLGYMFVALGVGAYGAAIFHL
FTHAFFKALLFLGAGSVIHAVDGEQDMRHMGGLRKHIPITYWTMMIGTLALTG--FPF---------
TAGYYSKDAIIESAFAGENAFAMYGYMMTVIAAALTSFYSWRLVFLTFHGKPR--ASHEVMHHV----------
---------------------------------------------------
HESPLVMTTPLFILSAGALLAGIVFYPFFLGG-SYET---FW--
HTALFAGPENHILHEAHEVPALVVWSPTIAMVLGFVVAYIFYIRQ-----------------------------
------------------------------
PQRPAQVAARNPSLYKFLLNKWYFDELYDVLFVRSAKWLGRFLWKK-GDVAVIDRLGPDGIS-
ARALDVTGWVVRLQTGYLYHYAFVMLIGVAALVTFMMFGGMR-------
>MBB4001705_NuoL_Aurantimonas_endophytica
MY-------QIIVLLPLAGFLVAGLFG----NAIGAKAAEYITSAFLIVAAALSWFAFITFGF--
GEGETLRVTLL-TWMQSGGLDIAWSLRIDRLTLVMLVVVNTVSALVHVYSIGYMHHDPH-------
RPRFFAYLSLFTFAMLMLVTSNNLVQMFFGWEGVGLASYLLIGFWYKKPSANAAAMKAFIVNRVGDFGFILGIF
GVYVLFGSVNL--
DTIFAGAADLMAQMLSFAGYALGMGAALTTVCLLLFMGAMGKSAQLGLHTWLPDAMEGPTPVSALIHAATMVTA
GVFLVARMSPLFELSHTALTVVTFVGATTAIFAATVGLVQNDIKRVIAYSTCSQLGYMFVALGVGAYSVAIFHL
FTHAFFKALLFLGAGSVIHAVSDEQDMRRMGGLRKHIPKTYWMMVIGTLALTG--FPY---------
TAGYFSKDAVIEAAFVGHNSMAMYGWGLTVLAALLTSFYSWRLIFMTFHGKPR--ASADVMHHI----------
---------------------------------------------------
HESPPVMLVPLYVLAVGALLAGIVFADFFIDG-GYDA---FW--
RSALFTGPENHVLHDMHEIPFAIVFLPTVMMALGFVLAWLFYIRS-----------------------------
------------------------------
PERPRQLAERHHGLYRFLLNKWYFDELYNFLFVRPAIAFGRILWRV-GDGRIIDGLGPDGVS-
ARVLDVTNRVVRLQTGYLYHYAFAMLIGVAALVTWTMLGGAN-------
>NDV85086_NuoL_Aurantimonas_aggregata
MY-------QIIVLLPLAGFLVAGLFG----NAIGAKEAEYITSAFLVVAAALSWFAFVTFGF--
GNGETLRVTLL-TWIQSGGLDISWSLRIDRLTLVMLVVVNTVSALVHIYSIGYMHHDPH-------
RPRFFAYLSLFTFAMLMLVTSNNLVQMFFGWEGVGLASYLLIGFWYKKPSANAAAMKAFIVNRVGDFGFILGIF
GIYVLFGSVNL--
DTIFAGAADLMAQMLSFAGYSLGMGAALTTVCLLLFMGAMGKSAQLGLHTWLPDAMEGPTPVSALIHAATMVTA
GVFLVARMSPLFELSHTALTVVTFVGATTAIFAATVGLVQNDIKRVIAYSTCSQLGYMFVALGVGAYSVAIFHL
FTHAFFKALLFLGAGSVIHAVSDEQDMRRMGGLRKHIPKTYWMMVIGTLALTG--FPY---------
TAGYFSKDAVIEAAFVGHNSFAMYGWALTVLAALLTSFYSWRLIFMTFHGKPR--ASADVMHHI----------
---------------------------------------------------
HESPPVMLVPLYVLAIGALLAGIVFADSFIDG-GYDA---FW--
RSALFTGPENHVLHDMHEIPFAIVFLPTVMMALGFALAWLFYIRF-----------------------------
------------------------------
PEQPRRLAERHRGLYQFLLNKWYFDELYNFLFVRPAIALGRMLWKT-GDGRIIDGLGPDGVS-
ARVLGVTNHVVRLQTGYLYHYAFAMLIGVAALVTWMMLGGVN-------
>NDW06534_NuoL_Jiella_pacifica
MY-------TIIVLLPLAGFLIAGLFG----KSIGAKACEYTLSGFMVVAAILSWIAFISFGF--
GEGETLRVPLL-QWMQSGTLDVAWSLRIDRLTLVMLVVVNTVSSLVHVYSIGYMHHDPH-------
RARFFAYLSLFTFAMLMLVTSNNLVQMFFGWEGVGLASYLLIGFWYKKPSANAAAMKAFVVNRVGDFGFALGIF
```

226 GVFALFGSVNL--
DTIFAGAADVAGSLMNFAGYALTMGSALTVVCLLLFMGAMGKSAQIGLHTWLPDAMEGPTPVSALIHAATMVTA
GVFMLARMSPLFELSSTALTFVTFIGATTAFFAATIGLVQNDIKRVIAYSTCSQLGYMFVALGVGAYGPAIFHL
FTHAFFKALLFLGAGSVIHAVSDEQDMRRMGGLRKHIPKTYWMMVIGTLALTG--FPF---------
TAGYYSKDAIIEAAFVGHNSFALYGWGMTVIAAALTSFYSWRLIFLTFHGKPR--ASADVMHHI----------
---------------------------------------------------
HESPWVMLVPLFVLAAGALFAGILFEGAFIGE-GYGE---FW--
HAALFTGSDNHVLHDMHEISFWIAFLPTLCMVVGFVTAYFFYIAR-----------------------------
------------------------------
PSLAKQTADRNRGLYQFLLNKWYFDELYDFLFVQPAQRVGRFLWRT-GDGKVIDGFGPDGVS-
ARVMDVTGRVVKLQSGYLYHYAFAMLIGVAALVTWMMLGGAS-------
227 >WP_08440924_NuoL_Fulvimarina_manganoxydans
228 MY-------QIIVLLPLAGFLIAGLAG----KAIGAKASEYTTSSFLVIAAVLSWIGFLSFGW--
GEGETIRVQML-QWMQSGGLDVSWSLRIDRLTLIMLVVVNTVSALVHIYSIGYMHHDPH-------
RPRFFAYLSLFTFAMLMLVTSNNLVQMFFGWEGVGLASYLLIGFWYKKPSANAAAMKAFVVNRVGDFGFILGIF
GVFVLFGSVNL--
DTIFAGAADIAATMLNFAGYALTMGGALTAVCLLMFVGAMGKSAQIGLHTWLPDAMEGPTPVSALIHAATMVTA
GVFMVARMSPVFELSHTALTFITFIGATTAFFAATIGLVQNDIKRVIAYSTCSQLGYMFVALGVGAYSVAIFHL
FTHAFFKALLFLGAGSVIHSVSDEQDMRRMGGLRKHIPITYWTMVIGTLALTG--FPF---------
TAGYFSKDAIIEAAFVGHNAFATYGWGMTVIAALLTSFYSWRLIFMTFHGKPR--ASADVMHHV----------
---------------------------------------------------
HESPMVMLVPLFVLSAGALLAGILFEHSFVGE-GYEH---FW--
HAALFTGAENHVLHDMHEVAFGIAILPTVMMALGFVVAYWMYVKD-----------------------------
------------------------------
KAAPKRLAETHRGLYRFFLNKWYFDELYNVLFVRPAMAVGRTLWKT-GDGRIIDGLGPDGVS-
ARVVDVTNRVVRLQTGYLYHYAFAMLIGVAALVTWMMLGGAN-------
229 >WP_007065974_NuoL_Fulvimarina_pelagi
230 MY-------QIIVLLPLAGFLIAGLAG----NTIGAKASEYITSSFLVVAAILSWIGFLTFDA---
DEPMAPITML-QWMQSGGLDVAWALKIDRLTLVMLVVVNTVSALVHIYSIGYMHHDPH-------
RPRFFAYLSLFTFAMLMLVTADNLVQMFFGWEGVGLASYLLIGFWYKKPSANAAAMKAFIVNRVGDFGFILGIF
GVFAIFQSVNL--
DTIFAGAADVASRMLNFAGYSLTMGGAITCVCLLLLLGAMGKSAQLGLHTWLPDAMEGPTPVSALIHAATMVTA
GVFMLARMSPIFELSHTALTVVTFVGASTAIFAATVGLVQNDIKRVIAYSTCSQLGYMFVALGVGAYSVAIFHL
FTHAFFKALLFLGAGSVIHAVSDEQDMRRMGGLRKHIPVTYWTMIIGTLALTG--FPF---------
TAGYFSKDAVIEAAYVGHNAFALYGWGLTVIAALLTSFYSWRLIFLTFHGKPR--ASADVMHHV----------
---------------------------------------------------
HESPSVMIVPLFFLAAGAILAGLLFEHEFVGD-AYEH---FW--
GASLFTSAENHVLHDLHEVAFGVAIIPTVMMAVGFALAYYFYIAN-----------------------------
------------------------------
PAMPKQLSQTHRGLYKFFLNKWYFDELYDFLFVRSSKWIGRTLWKT-GDGRIIDGLGPDGVS-
ARVIDVTNRVVKLQTGYLYHYAFAMLIGVAALVTWMMIGGIS-------
231 >WP_116682147_NuoL_Fulvimarina_endophytica
232 MY-------QIIVLLPLAGFLIAGLGG----NMIGAKASEYVTSSFLVVAALLSWIGFLSFDA---
AEPMAPITLL-HWMQSGDFDVSWALKIDRLTLVMLVVVNTVSALVHIYSMGYMHHDPH-------
RPRFFAYLSLFTFAMLMLVTADNLIQMFFGWEGVGLASYLLIGFWYKKPSANAAAMKAFVVNRVGDFGFLLGIF
GVFVLFGSVNL--
DTIFAGAADIAAQMLSFAGYSLTMGSALTVVCLLLFVGAMGKSAQIGLHTWLPDAMEGPTPVSALIHAATMVTA
GVFMVARMSPIFELSHTALTFITFIGATTAFFAATIGLVQNDIKRVIAYSTCSQLGYMFVAIGVGAYSVAIFHL

```
232… FTHAFFKALLFLCAGSVIHSVSDEQDMRRMGGLRKHIPVTYWTMVIGTLALTG--FPF---------
TAGYFSKDAVIEAAFVGHNAFAGYGWGLTVIAALLTSFYSWRLIFMTFHGKPR--ASADVMHHV----------
---------------------------------------------------
HESPMVMLVPLFILSIGALFAGMMFEHSFVGE-GYEH---FW--
QAALYMGPDNHILHDLHEVAFGIAIIPTIMMALGFVIAYWLYIVN-----------------------------
------------------------------
PAKPKQLSETHRGLYKFFLNKWYFDELYDFLFVRPAKALGRTLWKT-GDGRIIDGLGPDGVS-
ARVLDVTNRVVRLQTGYLYHYAFAMLIGVAALVTWMMIGGIS-------
233 >WP_029057157_NuoL_Stappia_stellulata
234 MY-------QAIVFLPLVGFLVAGLFG----RSIGAKASEYITSGLLIVAALLSWVAFFSIGF--
GETPVVRETLL-RWMTSGTFSVDWTIRVDTLTAVMLVVVNSVSALVHVYSIGYMHDDPH-------
RPRFFAYLSLFTFAMLMLVTSDNLLQMFFGWEGVGLASYLLIGFWYKKPSANAAAMKAFVVNRVGDFGFILGIL
GVFYLFNSTDY--
DTIFANAQAFAATVLTFLGAELSQDAALTVVCLLLFMGAMGKSAQFLLHTWLPDAMEGPTPVSALIHAATMVTA
GVFMVARLSPIFELSHTAMTVVTLFGATTAFFAATVGLVQNDIKRVIAYSTCSQLGYMFVALGVGGYSIAIFHL
FTHAFFKALLFLGAGSVIHAVSNEQDMRKMGGLRKHIPLTYWMMVIGTLALTG--FPL---------
TAGFFSKDAVIEAAYVGHNAFAQYAFWLTVAAAALTSFYSWRLAFMTFHGKPR--ASVDVMKHV----------
---------------------------------------------------
HESPLVMTVPLMILAAGALLAGFVFKGVFIGD-GYDA---FW--
KGALFVSEENHVLHDIHEVPLWVKASPFVMMLGGFAVAWMFYIRS-----------------------------
------------------------------
PEIPRQLAERHDGLYRFLLNKWYFDELYDVIFVRPAMWIGRQLWKK-GDGVVIDGMGPDGVA-
ARVQNVTTWVVRLQTGYLYHYAFAMLIGVAALITWSMFSVGGAH-----
235 >WP_107989924_NuoL_Breoghania_corrubedonensis
236 MY-------TAIVLLPLLGFLVAGIFG----RVIGAKASEYITSGFLIIAAILSWVAFISVGL--
GDGETQIIPIA-TWIVSGALDVQWAIRVDTLTAVMLVVVNTVSALVHVYSIGYMHHDPH-------
KSRFFAYLSLFTFAMLTLVTSDNLVQMFFGWEGVGLASYLLIGFWYQRPSANAAAMKAFVVNRIGDFGFALGIF
GVYYLFDSVNL--
ATIFANAQGFVESVLTFLGTPYEPATAMTVVCLLLFMGAMGKSAQFLLHTWLPDAMEGPTPVSALIHAATMVTA
GVFMVARLSPLFELSHTALTFVTFIGATTAFFAATIGLVQTDIKRVIAYSTCSQLGYMFVALGVGAYGVGVFHL
FTHAFFKALLFLGAGSVIHAVSDEQDMRKMGGLRKHIPLTYWMMIVGTLALTG--FPL---------
TAGYFSKDAIIEAAFVGDNAFAQYGFWMTVIAALMTSFYSWRLIFLTFHGKPR--ASVDVMKHV----------
---------------------------------------------------
HESPPVMTVPLMVLALGALVAGFVFSGSFIGH-HQGE---FW--
KTALFMSADNHVLEEMHHVPQWVKWSPFVMMVIGLVVAWYFYIKA-----------------------------
------------------------------
PDAPKRLAARHDGLYKFLLNKWYFDELYDFLFVRPAMKLGRFLWKR-GDGTVIDGFGPDGIS-
TLVQDVTARVVRLQTGYIYHYAFAMLIGVALLVTWTMFAGGGAH-----
237 >WP_136353129_NuoL_Mesorhizobium_composti
238 MY-------QAIVFLPLIGFLVVGLFG----NSLGAKASEFITSGLLVISAVLSWVAFFTVAF--
GDGEVFTVPVL-RWIQSGGIDTSWALRIDTLTAVMLVVVNTVSSLVHIYSIGYMHHDPN-------
RPRFFAYLSLFTFAMLMLVTADNLIQMFFGWEGVGLASYLLIGFWYKKPSANAAAIKAFVVNRVGDFGFILGIF
GVFVLFGSVNL--
TTIFANAASYLPAVLTFLGYALDKGAAMTVVCLLLFMGAMGKSAQVPLHTWLPDAMEGPTPVSALIHAATMVTA
GVFMLARLSPLFELSHTALTVVTFIGAITAFFAATVGLVQNDIKRVIAYSTCSQLGYMFVALGVGAYGAAIFHL
FTHAFFKALLFLGSGSVIHAMSDEQDMRKMGGIRKLIPNTYWMMVIGTLALTGVGIPMTLVG-----
FAGFFSKDAIIEASFVGENAVATFAFVLLVIGAAFTSFYSWRLIFMTFHGEPR--ASHEVMHHV----------
---------------------------------------------------
```

```
238 HESPPVMLVPLYVLAAGAIFAGVLFSTYFVGE-GYEH---FW--
KAALFTGPDNHVLHEMHGVPLWVKLAPIVVTVLGFLVAWQFYIRK-----------------------------
------------------------------
PALPAEVAGRHRGLYAFLLNKWYFDEVYDFLFVRFAKWLGYTLWKK-GDGAVIDGLGPDGVS-
ARVIDVTNRVVKLQTGYLYHYAFAMLIGVAALVTWMML-----------
239 >WP_035023849_NuoL_Aquamicrobium_defluvii
240 MY-------QAIVFLPLIGFLIVGLFG----TSLGAKASEYITSGLVVVSAALSWIAFFSVGF--
SDAETFTVPLL-RWLQTGGLDAAWALRIDTLTVVMLVVVNTVSALVHIYSIGYMHHDPH-------
RPRFFAYLSLFTFAMLMLVTSDNLVQMFFGWEGVGLASYLLIGFWYKKPSASAAAMKAFIVNRVGDFGFLLGLF
GVFVLFGSVNF--
STIFAGAATYLPAVLTFLGYSLDRTAAITVVCLLLFMGAMGKSAQVPLHTWLPDAMEGPTPVSALIHAATMVTA
GVFMLARLSPLFELSHSALTVVTFIGATTAFFAATVGLVQNDIKRVIAYSTCSQLGYMFVALGVGAYGAAIFHL
FTHAFFKALLFLGSGSVIHAMSDEQDMRKMGGIRKLIPTTYWMMIIGTVALTGLGIPATVIG-----
TAGFFSKDAIIESAFVGHNAVAGFAFALLVIAACFTSFYSWRLIFMTFHGKSR--ASEEVLHHV----------
---------------------------------------------------
HESPPVMLVPLYILAVGALFAGFVFHDQFIGD-AYEA---FW--
KTALFTSADNHVLHDMHNVPMWVKMSPFVAMVVGFLLSYQFYIRS-----------------------------
------------------------------
PETPKRLAAQHRGLYQFLLNKWYFDELYDFLFVRSAKALGRFLWKT-GDGKIIDGLGPDGVS-
ARVVDVTQRVVRLQSGYLYHYAFAMLIGVAALATWMML-----------
241 >WP_183830913_NuoL_Aquamicrobium_lusatiense
242 MY-------QAIVFLPLIGFLIVGLFG----NAIGARASEYLTSGLMVIVAALSWVAFFTVGF--
SETETFTVPLL-RWLQSGGLDADWALRIDTLTVVMLVVVNTVSALVHIYSIGYMHHDPH-------
RPRFFAYLSLFTFAMLMLVTSDNLVQMFFGWEGVGLASYLLIGFWYKKPSASAAAMKAFIVNRVGDFGFLLGLF
GVFVLFGSVNF--
STIFASTANYLPAVLTFLGYSLDKTAAMTIVCLLLFMGAMGKSAQVPLHTWLPDAMEGPTPVSALIHAATMVTA
GVFMLARLSPLFELSHSALTVVTFIGATTAFFAATVGLVQNDIKRVIAYSTCSQLGYMFVALGVGAYGAAVFHL
FTHAFFKALLFLGSGSVIHAMSDEQDMRKMGGIRKLIPTTYWMMIIGTIALTGLGIPATVIG-----
TAGFFSKDAIIESAFVGHNAVAGYAFALLVIAACFTSFYSWRLIFMTFHGKSR--ASEDVLHHV----------
---------------------------------------------------
HESPPVMLVPLYILAVGALFAGFVFHDQFIGE-AYEA---FW--
KTALFTSADNHVLHEMHNVPMWVKMSPFVAMVVGFLLSYQFYIRS-----------------------------
------------------------------
PETPKRLAAQHRGLYQFLLNKWYFDELYDFLFVRSAKAIGRFLWKT-GDGKIIDGLGPDGIS-
ARVVDVTQRVVKLQSGYLYHYAFAMLIGVAALATWMML-----------
243 >WP_013894820_NuoL_Mesorhizobium_opportunistum
244 MY-------QAIVFLPLLGFLIVGLFG----TSLGAKASEYITSGFLVISALLSWVAFFSVGF--
GHGEVFTVPVL-HWIQSGGLDVSWALRIDTLTVVMLVVVNTVSALVHIYSIGYMHHDPN-------
RPRFFAYLSLFTFAMLMLVTADNLVQMFFGWEGVGLASYLLIGFWYKKPSANAAAIKAFVVNRVGDFGFALGIF
GVFVLFGSVNL--
GTIFANAATFVPAVLTFLGHALDKQAAMTVVCLLLFMGAMGKSAQVPLHTWLPDAMEGPTPVSALIHAATMVTA
GVFMLARLSPLFELSHSALTVVTFIGAFTAFFAATVGLVQNDIKRVIAYSTCSQLGYMFVALGVGAYGAAIFHL
FTHAFFKALLFLGSGSVIHAVSDEQDMRKMGGLRKLIPATYWMMVIGTLALTGVGIPVTVIG-----
TAGFFSKDAIIETAFASHNSVAGLAFVLLVIAAGFTSFYSWRLIFMTFHGKPR--ASHEVMHHV----------
---------------------------------------------------
HESPPVMLVPLFILAAGALFAGIIFHNSFIGE-GYAE---FW--
KASLFTLPDNHILHEIHELPLWVELSPFIAMLVGLALAWKFYIRS-----------------------------
------------------------------
```

```
PEMPVNLAAQHRGLYAFLLNKWYFDELYDFLFVRPAKRLGSFLWKT-GDGTIIDGLGPDGIS-
ARVVDVTNRVVKLQTGYLYHYAFAMLIGVAAFVTWMML-----------
>WP_192362998_NuoL_Mesorhizobium_mediterraneum
MY-------QAIVFLPLLGFLIVGLFG----TSLGAKASEYITSGLLVISAVLSWVALFAVGF--
GEGEVFTVPVM-RWIQSGGLEAAWALRIDTLTVVMLVVVNTVSALVHIYSIGYMHHDPN-------
RPRFFAYLSLFTFAMLMLVTADNLVQMYFGWEGVGLASYLLIGFWYKKPSANAAAIKAFVVNRIGDFGFALGIF
GVFVLFGSVNL--
GTIFANAATFIPAVLTFLGYALDKQAAMTVVCLLLFMGAMGKSAQVPLHTWLPDAMEGPTPVSALIHAATMVTA
GVFMLARLSPLFELSHSALIVVTFIGAFTAFFAATVGLVQNDIKRVIAYSTCSQLGYMFVALGVGAYGAAIFHL
FTHAFFKALLFLGSGSVIHAVSDEQDMRKMGGLRTLIPKTYWMMVIGTLALTGVGIPVTVIG-----
TAGFFSKDAIIESAFAAHNPVAGLAFVLLVVAACFTSFYSWRLIFMTFHGQPR--ASHEVMHHV----------
---------------------------------------------------
HESPPVMLVPLFILAAGSLFAGVIFHGAFIGE-GYAE---FW--
KASLFTLPENHILHDIHEVPLWVKLAPFVAMLIGFAIAWQFYIRA-----------------------------
------------------------------
PEMPKNLAAQHRGLYAFLLNKWYFDELYDLLFVRPAKRLGHFLWKT-GDGTVIDGLGPDGVS-
ARVVDVTNRVVKLQTGYLYHYAFAMLIGVAALVTWMML-----------
>WP_047144650_NuoL_Aquamicrobium_sp._LC103
MY-------HAIVFLPLLGFLIAGLFG----RAIGAKASEYVTSGFLVISAILSWVAFFTVAL--
GETEAFTVPVL-RFIQAGGLDVSWALRIDTLTAVMLVVVNTVSALVHIYSIGYMHHDPH-------
RPRFFAYLSLFTFAMLMLVTSDNLVQMFFGWEGVGLASYLLIGFWYKKPSANAAAIKAFVVNRVGDFGFLLGLF
GVFVLFGSVNF--
DTIFANAAAMIPAVFNFLGYALNNQTALTVICLLLFMGAMGKSAQVPLHTWLPDAMEGPTPVSALIHAATMVTA
GVFMLARLSPIFELSHSALLVVTFVGAFTAFFAATVGLVQNDIKRVIAYSTCSQLGYMFVALGLGFYSAAIFHL
FTHAFFKALLFLGSGSVIHAVSDEQDMRKMGGLRKLIPATYWMMVIGTIALTGVGIPATIIG-----
TAGFFSKDAIIEGAFVGHNALATFAFILLTIAAVFTSFYSWRLIFMTFHGRPR--ASADVMHHV----------
---------------------------------------------------
HESPPVMLVPLFILAVGALFAGLVFHDWFIGH-HYDE---FW--
KGALFTLPDNHILHEFHDVPMWVKLAPFLAMLIGFFVAYQFYIRS-----------------------------
------------------------------
PETPKQLAAQHEGLYRFLLNKWYFDELYDFLFVRPAMRLGRFLWKK-GDGAIIDGLGPDGIS-
ARVVDVTQRVVKLQTGYLYHYAFAMLIGVAALVTWMMLGSSF-------
>WP_009450014_NuoL_Nitratireductor_indicus
MY-------HAIVFLPLVGFLIAGLFG----RSIGAKASEYITSGLMVIAALLSWVAFFSVAM--
GSGEAFTVPVL-RFITSGSLDMSWALRIDTLTVVMLIVVNTVSALVHIYSIGYMHHDPH-------
RPRFFAYLSLFTFAMLMLVTSDNLIQMFFGWEGVGLASYLLIGFWFKKPSANAAAMKAFVVNRVGDFGFLLGIF
GLFMLFGSVNF--
STIFANAATYLPAVLTFLGYALDKGAALTVICLLLFMGAMGKSAQVPLHTWLPDAMEGPTPVSALIHAATMVTA
GVFMVARLSPIFELSHTALTVVTFVGAFTAFFAATVGLVQNDIKRVIAYSTCSQLGYMFVALGTGFYGAAIFHL
FTHAFFKALLFLGSGSVIHAVSDEQDMRKMGGLRKLIPTTYWMMVIGTIALTGVGIPLTVIG-----
TAGFFSKDAIIEGVFAGENYFSGFAFVLLTVAAVFTSFYSWRLIFMTFHGKPR--ASADVMHHV----------
---------------------------------------------------
HESPLVMLVPLFILAAGALFAGVAFKEFFIGH-DYEH---FW--
KAALFTLPENHILEEFHHVPLWVKLAPFVAMLLGLYVAWVFYIRS-----------------------------
------------------------------
PERPKALARRFSGLYAFLLNKWYFDEAYDFLFVRPAKRLGHFLWKK-GDGIVIDGLGPDGIS-
ARVVDVTNRIVKLQSGYLYHYAFAMLIGVAALVTWMMLGGTF-------
>WP_206554529_NuoL_Nitratireductor_aquibiodomus
```

```
MY-------QAIVFLPLIGFLIAGLFG----RSIGAKASELITSGLMVITALLSWVAFFSVAL--
GSGEAFTVPLL-SFITSGNLDVSWAIRVDTLTAVMLIVVNTVSALVHIYSIGYMHHDPH-------
RPRFFAYLSLFTFAMLMLVTSDNLVQMFFGWEGVGLASYLLIGFWFKKPSANAAAMKAFVVNRVGDFGFLLGIF
GIYMLFGSVNF--
STIFANAATYLPAVLTFLGYALDKGGALTAVCLLLFMGAMGKSAQVPLHTWLPDAMEGPTPVSALIHAATMVTA
GVFMVARLSPVFELSHTALTVVTFVGAFTAFFAATVGLVQNDIKRVIAYSTCSQLGYMFVALGTGFYSAAIFHL
FTHAFFKALLFLGSGSVIHAVSDEQDMRKMGGLRKLIPQTYWMMIIGTIALTGVGIPMTVIG-----
TAGFFSKDAIIEGAFAGQNYFAMFAFVMLTIAAVFTSFYSWRLIFMTFHGKPR--ASADVMSHV----------
---------------------------------------------------
HESPLVMLVPLFILAVGALFAGVAFKGFFIGD-AYDQ---FW--
KGALFTLPGNHMLEEFHHVPVWVKLAPFVAMLLGLYVAWVYYIRS-----------------------------
------------------------------
PERPKRLAKQFSGLYAFLLNKWYFDELYDFLFVRPAMRLGRFLWKK-GDGAVIDGLGPDGIS-
ARVVDVTNRIVKLQSGYLYHYAFAMLIGVAALVTWMMLGGTF-------
>WP_209336310_NuoL_Tianweitania_sediminis
MY-------QAIVFLPLIGFLVAGLFG----RSIGAKASEYVTTGLLIVSAVLSWIAFFSVAL--
GHGEPFTVPIL-RWIQVGGVDVAWSLRIDTLTAVMLIVVNTVSALVHVYSIGYMHHDPH-------
RPRFFAYLSLFTFAMLMLVTSDNLIQMFFGWEGVGLASYLLIGFWYKKPSASAAAMKAFIVNRVGDFGFALGIF
GVFVLFGSVNF--
DTIFANAATFIPAALNFLGYALSNQAALTVVCLLLFMGAMGKSAQVPLHTWLPDAMEGPTPVSALIHAATMVTA
GVFMLARLSPIFEHSHTALTFVTFIGATTAIFAATVGLVQNDIKRVIAYSTCSQLGYMFVALGTGFYSAAIFHL
FTHAFFKALLFLGSGSVIHAVSDEQDMRKMGGLRKLIPTTYWMMVIGTIALTGVGIPTTILG-----
TAGFFSKDAIIEGAFAAHNPMAGYAFVLLVLAAVFTSFYSWRLIFMTFHGTPR--ASADVMHHV----------
---------------------------------------------------
HESPPVMLVPLYILAAGALFAGVIFHDFFIGH-IEEYNE-FW--
KGALFTLPENEILDHFHHVPLWVKLSPLVAMVIGFALAYQFYIRS-----------------------------
------------------------------
PETPQRLAAQHQGLYKFLLNKWYFDELYDFLFVRPAMRLGHFLWKQ-GDGRVIDGLGPDGIA-
ARVADVTRGVVRLQTGYLYHYAFVMLIGVAALVTWMMLGNTF-------
>WP_114442382_NuoL_Phyllobacterium_salinisoli
MLY------YAIVFLPLAGFLAAGLLG----KQLGVKACEYITSGFLVIAAALSWVAFFQVAL--
GHGETLRIPVM-HWVTSGTLSFDWAFRIDTLTAVMLVVVNTVSALVHIYSIGYMHHDPH-------
RPRFFAYLSLFTFAMLMLVTSDNLVQMFFGWEGVGLASYLLIGFWFKKPSANAASMKAFIVNRVGDFGFLLGIF
GVFVLFNSVSY--
DTIFAAAANYLPAAVHFLGYSLDRQGAITVVCLLLFMGAMGKSAQFLLHTWLPDAMEGPTPVSALIHAATMVTA
GVFMVARMSPLFELSQTALTFVTIIGATTAFFAATVGLVQNDIKRVIAYSTCSQLGYMFVALGVGAYGAGVFHL
FTHAFFKALLFLGAGSVIHAVSDEQDMRRMGGLRKLIPTTYWMMIIGTVALTGLGIPGTSIG-----
TAGFFSKDAIIESSYASHGPAAGYAFVMLTVAALFTSFYSWRLIFMTFHGKPR--ATADVMHHV----------
---------------------------------------------------
HESPPVMLVPLFVLGAGALIAGFLFHDYFFGH-HYVE---FW--
KGALFTGPHNEILEEYHHVPFLVKIAPFLAMAVGFVIAWIFYIRS-----------------------------
------------------------------
PEMPKQLARRHPGLYQFLLNKWYFDELYNFLFVRPTLALGRILWKG-GDGWLIDGFGPDGIS-
ARVVDVTNRVVKLQTGYLYHYAFAMLIGVAALVTWMMLGSSF-------
>WP_147835202_NuoL_Phyllobacterium_endophyticum
ML-------LAIVFLPLLGFLIAGLFG----SSIGAKASEYITTGLLIICAVLSWIAFGTVAL--
GHAEAFKVPVM-RWVESGSLSFDWSLRIDTLTAVMLVVVNTVSALVHTYSIGYMHHDPN-------
RPRFFAYLSLFTFAMLMLVTGDNLVQMFFGWEGVGLASYLLIGFWFKKPSANAAAMKAFIVNRVGDFGFLLGIF
```

```
ALFVLFDTVQF--
DTLFAAAREYLPAVLNFLGYALDKQGAITITCLLLFMGAMGKSAQFLLHTWLPDAMEGPTPVSALIHAATMVTA
GVFMVARLSPIFELSHTALIFVTFIGATTAFFAATIGLVQNDIKRVIAYSTCSQLGYMFVALGLGAYGAGIFHL
FTHAFFKALLFLGAGSVIHAVSDEQDMRHMGGLRTFIPKTYWMMIVGTVALTGLGIPGTVIG-----
TAGFFSKDMIIESAFASHSPMASYAFILLVVAAIFTSFYSWRLIFMTFHGKPR--ASHDVMHHV----------
---------------------------------------------------
HESPYVMLVPLFILAAGALFAGVMFREYFFGH-EYLE---FW--
KGAIFTGPENEILEEHHHVPFLVALSPFLAMAAGFVVAWYFYIRS-----------------------------
------------------------------
PETPKRLAAQHPGLYQFLLNKWYFDELYDFLFVRPAKALGRFLWKR-GDGWLIDGHGPDGIS-
ARVVDVTNRVVRLQSGYLYHYAFAMLIGVAALVTWMMLGSNF-------
>WP_162293147_NuoL_Phyllobacterium_zundukense
MLY------LAIVFLPLLGFLIAGLFG----SSIGAKASEYVTTGLLIVCAVLSWIAFGTVAL--
GHGEAFKVPVM-HWVESGSLSFDWALRIDTLTAVMLVVVNTVSALVHTYSIGYMHHDPD-------
RPRFFAYLSLFTFAMLMLVTGDNLVQMFFGWEGVGLASYLLIGFWFKKPSANAAAIKAFVVNRVGDFGFLLGIF
ALFVLFDTVQF--
DTLFTSAAQYLPAVLNFLGYALDKQGAITIACLLLFMGAMGKSAQFLLHTWLPDAMEGPTPVSALIHAATMVTA
GVFMVARLSPIFELSHTALIFVTFIGATTAFFAATVGLVQNDIKRVIAYSTCSQLGYMFVALGLGAYGAGIFHL
FTHAFFKALLFLGAGSVIHAVSDEQDMRHMGGLRKLIPQTYWLMVVGTVALTGLGIPGTLIG-----
TAGFFSKDMIIESAYASHSPVAGYAFILLVIAAIFTSFYSWRLIFMTFHGKPR--ASHDVMHHV----------
---------------------------------------------------
HESPYVMLVPLFILAFGALFAGVGFHEYFFGH-EYAE---FW--
KGAIFTGPENEILEEHHHVPFLVAMSPFLAMAAGFLVSWYFYIRS-----------------------------
------------------------------
PETPKRLAAQHPGLYQFLLNKWYFDELYDFLFVRPAKALGRFLWKQ-GDGWLIDGHGPDGIS-
ARVVDVTNRVVRLQTGYLYHYAFAMLIGVAALVTWMMLGSNF-------
>WP_105735480_NuoL_Phyllobacterium_myrsinacearum
MLY------LAIVFLPLLGFLIAGLFG----SSIGAKASEYVTTGLMIVVAVLSWIAFTTVAL--
GHGEAFKVPVM-HWVESGSLSFDWALRIDTLTAVMLIVVNTVSALVHTYSIGYMHHDPH-------
RPRFFAYLSLFTFAMLMLVTGDNLVQMFFGWEGVGLASYLLIGFWFKKPSANAAAMKAFIVNRVGDFGFLLGIF
ALFVLFDTVQF--
DTLFQSAAQYLPAVLNFLGYALGKQEAITVACLLLFMGAMGKSAQFLLHTWLPDAMEGPTPVSALIHAATMVTA
GVFMVARLSPIFELSHTALIVVTLVGATTAFFAATVGLVQNDIKRVIAYSTCSQLGYMFVALGLGAYGAAIFHL
FTHAFFKALLFLGAGSVIHAVSDEQDMRHMGGLRKLIPQTYWMMIIGTVALTGLGIPGTAIG-----
TAGFFSKDAIIESAYASTSPAAGYAFILLVIAALFTSFYSWRLIFMTFHGKPR--ASHDVMHHV----------
---------------------------------------------------
HESPYVMLVPLFILAIGALFAGVIFREYFFGH-EYAE---FW--
KGALFTGKENEILELHHHVPALVAASPFLAMLAGFILSWYFYIRS-----------------------------
------------------------------
PQTPVRLAEQHRGLYQFLLNKWYFDELYDFLFVRPAKALGKFLWKK-GDGWLIDGHGPDGIS-
ARVVDVTNRVVRLQTGYLYHYAFAMLIGVAALVTWMMLGSNF-------
>PSH67906_NuoL_Phyllobacterium_brassicacearum
MY-------LAIVFLPLLGFLIAGLFG----SSIGAKASEYVTTGLLIVCAVLSWIAFGTVAL--
GHGEAFKVPVA-LWVESGSLSFDWALRIDTLTAVMLVVVNTVSALVHTYSIGYMHHDPD-------
RPRFFAYLSLFTFAMLMLVTGDNLVQMFFGWEGVGLASYLLIGFWYKKPSANAAAIKAFVVNRVGDFGFLLGIF
ALFVLFDTVQF--
DTLFTSAAQYLPAVLNFLGYSLDKQGAITITCLLLFMGAMGKSAQFLLHTWLPDAMEGPTPVSALIHAATMVTA
GVFMVARLSPIFELSHAALIFVTFIGATTAFFAATVGLVQNDIKRVIAYSTCSQLGYMFVALGLGAYGAGIFHL
```

```
FTHAFFKALLFLGAGSVIHAVSNEQDMRHMGGLRKLIPQTYWMMIIGTIALTGLGIPGTLIG-----
TAGFFSKDMIIESAYASHSPVAGYAFILLVVAAIFTSFYSWRLIFMTFHGKPR--ASHDVMHHV----------
---------------------------------------------------
HESPYVMLIPLFILAFGALFAGVGFREYFFGH-EYAE---FW--
KGAIFTGPENEILEEHHHVPFLVAMSPFLAMAAGFIVAWYYYIRS-----------------------------
------------------------------
PETPKQLAANHPGLYQFLLNKWYFDELYDFLFVRPAKSLGRFLWKK-GDGWLIDGHGPDGIS-
ARVVDVTNRVVRLQTGYLYHYAFAMLIGVAALVTWMMLGSNF-------
>WP_162701970_NuoL_Phyllobacterium_trifolii
MLY------LAIVFLPLLGFLIAGLFG----SSIGAKASEYVTTGLLIVCAVLSWIAFATVAL--
GHGEAFKVPVA-PWVESGSLSFDWALRIDTLTVVMLVVVNTVSALVHTYSIGYMHHDPD-------
RPRFFAYLSLFTFAMLMLVTGDNLVQMFFGWEGVGLASYLLIGFWFKKPSANAAAIKAFVVNRVGDFGFLLGIF
ALFVLFDTVQF--
DTLFASAAQYLPAVLNFLGYTLDKQGAITIACLLLFMGAMGKSAQFLLHTWLPDAMEGPTPVSALIHAATMVTA
GVFMVARLSPIFELSHTALIFVTFIGATTAFFAATVGLVQNDIKRVIAYSTCSQLGYMFVALGLGAYGAGIFHL
FTHAFFKALLFLGAGSVIHAVSDEQDMRHMGGLRTLIPKTYWMMIVGTVALTGLGIPGTLIG-----
TAGFFSKDMIIESAFASHSPMASYAFILLVVAAIFTSFYSWRLIFMTFHGKPR--ASHDVMHHV----------
---------------------------------------------------
HESPYVMLVPLFILAAGALFAGVIFREYFFGH-EYAE---FW--
KGAIFTGPENEILEEHHHVPFLVAMSPFLAMAVGFIVAWYFYIRS-----------------------------
------------------------------
PETPRRLAAQHPGLYQFLLNKWYFDELYDFLFVRPTKALGRFLWKQ-GDGWLIDGHGPDGIS-
ARVVDVTNRVVRLQSGYLYHYAFAMLVGVAALVTWMMLGSNF-------
>WP_110754317_NuoL_Phyllobacterium_leguminum
MLY------YAIVFLPLLGFLVAGLFG----KQLGAKACEFITSGFLVIAAVLSWVAFFQVAL--
GHGETVRIPVL-HWVTVGSLSFDWALRIDTLTAVMLVVVNTVSALVHIYSIGYMHHDPH-------
RPRFFAYLSLFTFAMLMLVTSDNLVQMFFGWEGVGLASYLLIGFWFQKPSANAAAMKAFVVNRVGDFGFLLGIF
GVFVLFNAVNY--
DTIFPAASNYLSGVLNFLGYSLDRQGAITVICLLLFMGAMGKSAQFLLHTWLPDAMEGPTPVSALIHAATMVTA
GVFMVARMSPLFELSQTALTLITIIGATTAFFAATVGLVQNDIKRVIAYSTCSQLGYMFAALGVGAFGAGVFHL
FTHAFFKALLFLCAGSVIHAVSDEQDMRNMGGLRKLIPTTYWMMIIGTVALTGLGIPFTEIG-----
TAGFFSKDAIIESTYVSHNPAAGYAFWLLVIAALFTSFYSWRLIFMTFHGKPR--ASHEVMHHV----------
---------------------------------------------------
HESPPVMLFPLFVLAIGALFAGFVFREYFFGH-EYAE---FW--
KGALFTAPGNQMLEEHHHVPAWVAASPFVAMVLGFVIAWVFYIRS-----------------------------
------------------------------
PEMPKELARRHPGLYQFLLNKWYFDELYDFIFVRPTLWLGRVLWKD-GDGKIIDGLGPNGIS-
ARVVDVTNRVVKLQSGYLYHYAFAMLIGVAALVTWMMIGSSI-------
>WP_189501590_NuoL_Corticibacterium_populi
MY-------QAIVFLPLLGFLIAGLFG----RSIGAKASEYVTSGFLVVSAVLSWIAFFTVAL--
GHVEAFTVPVL-RFIQAGGIDVSWALRIDTLTAVMLIVVNTVSALVHIYSIGYMHHDPH-------
RPRFFAYLSLFTFAMLMLVTSDNLIQMFFGWEGVGLASYLLIGFWYKKPSASAAAMKAFIVNRVGDFGFALGIF
GVFVLFGSVNF--
DTIFANAATFIPAALTFLGYALNNQAALTIVCLLLFMGAMGKSAQVPLHTWLPDAMEGPTPVSALIHAATMVTA
GVFMLARLSPIFEHSATALTVVTFIGATTAIFAATVGLVQNDIKRVIAYSTCSQLGYMFVALGTGFYGAAVFHL
FTHAFFKALLFLGSGSVIHAVSDEQDMRRMGGLRKLIPTTYWMMVIGTIALTGVGIPTTMIG-----
TAGFFSKDAIIEGSFAAHNPVTGYAFIMLVIAAVFTSFYSWRLIFMTFHGKPR--ASADVMHHV----------
---------------------------------------------------
```

```
HESPPVMLVPLFILAVGALLAGVLFEGYFVGH-EYNE---FW--
KGALFTLPENEILEHFHHVPLWVKLSPFIAMIVGFLLAYQFYIRS-----------------------------
------------------------------
PETPKRLAAQHHGLYQFLLNKWYFDELYDFLFVRPAMRIGRFLWKQ-GDGRVIDGLGPDGIA-
ARVTDVTRGVVRLQTGYLYHYAFVMLIGIAALVTWMMLGNSF-------
>WP_055969364_NuoL_Aminobacter_sp._DSM101952
MY-------QLIVFLPLLGFLIAGLFG----RSIGAKASEYITSGLLVVSAALSWVAFITVGL--
GDVEVFTVPVL-RFIDSGGLQANWALRIDTLTTVMLVVVNTVSALVHIYSIGYMHHDPH-------
RPRFFAYLSLFTFAMLMLVTSDNLVQMFFGWEGVGLASYLLIGFWYKKPSANAAAIKAFVVNRVGDFGFLLGIF
GLFVLFGSVEF--
STIFAGAASYLPAVLTFLGYALDKQGALTIVCLLLFMGAMGKSAQVPLHTWLPDAMEGPTPVSALIHAATMVTA
GVFMLARMSPVFELSHSALTVVTFIGAFTAFFAATVGLVQNDIKRVIAYSTCSQLGYMFVALGMGFYSAAIFHL
FTHAFFKALLFLGSGSVIHAVSDEQDMRKMGGLRKLIPTTYWMMVIGTLALTGVGIPATIIG-----
TAGFFSKDAIIEGAFVGHNAVAGLAFAALVIAACFTSFYSWRLIFMTFHGKPR--ATADVMHHV----------
---------------------------------------------------
HESPPVMLVPLFILAVGALFAGVVFHDQFIGE-GYNE---FW--
KGALFTLETNHILHDFHGVPMWVKLAPFVAMIAGFLMAYLFYIRS-----------------------------
------------------------------
PEMPKVLAERHRGLYAFLLNKWYFDELYDFLFVRPAKWLGRFLWKK-GDGWLIDGFGADGIS-
ARVVDVTNRVVKLQTGYLYHYAFAMLIGVAALVTWMMLGSSF-------
>WP_184698080_NuoL_Aminobacter_aganoensis
MY-------HFIVFLPLLGFLIAGLFG----RSIGAKASEYITSGLLVVSAALSWVAFITVGL--
GDVEVFTVPVL-RFIDSGGLQANWALRIDTLTTVMLVVVNTVSALVHIYSIGYMHHDPH-------
RPRFFAYLSLFTFAMLMLVTSDNLVQMFFGWEGVGLASYLLIGFWYKKPSANAAAIKAFVVNRVGDFGFLLGIF
GLFVLFGSVEF--
STIFAGAASYLPAVLTFLGYALDKQGALTIVCLLLFMGAMGKSAQVPLHTWLPDAMEGPTPVSALIHAATMVTA
GVFMLARMSPVFELSHSALTVVTFIGAFTAFFAATVGLVQNDIKRVIAYSTCSQLGYMFVALGMGFYSAAIFHL
FTHAFFKALLFLGSGSVIHAVSDEQDMRKMGGLRKLIPTTYWMMVIGTLALTGVGIPATIIG-----
TAGFFSKDAIIEGAFVGHNAVAGLAFAALVIAACFTSFYSWRLIFMTFHGKPR--ATADVMHHV----------
---------------------------------------------------
HESPPVMLVPLFILAVGALFAGVVFHDQFIGE-GYNE---FW--
KGALFTLETNHILHDFHGVPMWVKLAPFVAMIAGFLMAYLFYIRS-----------------------------
------------------------------
PEMPKVLAERHRGLYAFLLNKWYFDELYDFLFVRPAKWLGRFLWKK-GDGWLIDGFGADGIS-
ARVVDVTNRVVKLQTGYLYHYAFAMLIGVAALVTWMMLGSSF-------
>WP_115731531_NuoL_Aminobacter_aminovorans
MTY------HFIVFLPLIGFLIAGLLG----RSIGAKASEYITSGLLVVSAVLSWVAFFTVAL--
GDVETFTVPVL-RFLDVGGIQADWALRIDTLTTVMLVVVNTVSALVHIYSIGYMHHDPH-------
RPRFFGYLSLFTFAMLMLVTSDNLVQMFFGWEGVGLASYLLIGFWYKKPSANAAAIKAFVVNRVGDFGFLLGIF
GLFVLFGSVEF--
STIFAGAASYLPAVLTFLGYALDKHNALTVVCLLLFMGAMGKSAQVPLHTWLPDAMEGPTPVSALIHAATMVTA
GVFMLARLSPVFELSHTALTVVTFIGAITAFFAATVGLVQNDIKRVIAYSTCSQLGYMFVALGMGFYSAAIFHL
FTHAFFKALLFLGSGSVIHAVSDEQDMRKMGGLRKLIPTTYWMMVIGTLALTGVGIPATIIG-----
TAGFFSKDAIIEGAFVGHNAVAGFAFAMLVIAACFTSFYSWRLIFMTFHGKPR--ATADVMHHV----------
---------------------------------------------------
HESPPVMLVPLFILAAGALFAGVIFHDQFIGE-AYAE---FW--
KGALFTLDTNHILHDFHSVPMWVKLAPFAAMIGGFAMAYLFYIRS-----------------------------
------------------------------
```

```
PEMPKVLAERHRGLYAFLLNKWYFDELYDFLFVRPAKWLGSFLWKK-GDGWLIDGFGADGIS-
ARVVDVTNRVVKLQTGYLYHYAFAMLIGVAALVTWMMLGSSF-------
>MBA8904744_NuoL_Aminobacter_ciceronei
MMY------HFIVFLPLLGFLIAGLLG----RSIGAKASEYVTSGFLVISAVLSWIAFVTVAL--
GDVEAFTVPVL-RFLDVGGLQADWALRIDTLTAVMLVVVNTVSALVHIYSIGYMHHDPH-------
RPRFFAYLSLFTFAMLMLVTSDNLVQMFFGWEGVGLASYLLIGFWYKKPSANAAAIKAFVVNRVGDFGFLLGIF
GLYVLFGSVEF--
STIFASAASYLPAVLNFLGYALDKHSALTIVCLLLFMGAMGKSAQVPLHTWLPDAMEGPTPVSALIHAATMVTA
GVFMLARLSPVFELSHTALTVVTFIGAITAIFAATVGLVQNDIKRVIAYSTCSQLGYMFVALGMGFYSAAIFHL
FTHAFFKALLFLGSGSVIHAVSDEQDMRKMGGLRKLIPTTYWMMVIGTLALTGVGIPATIIG-----
TAGFFSKDAIIEGAFVGHNAVAGFAFAMLVIAACFTSFYSWRLIFMTFHGKPR--ATADVMHHV----------
---------------------------------------------------
HESPPVMLVPLFVLAVGALLAGVVFHDQFIGE-AYNE---FW--
KGALFTLDTNHILHDFHGVPMWVKLAPFAAMIGGFAMAYLFYIRS-----------------------------
------------------------------
PEMPKVLAERHRGLYAFLLNKWYFDELYDFLFVRPAKWLGSFLWKK-GDGWLIDGFGADGIS-
ARVVDVTNRVVKLQTGYLYHYAFAMLIGVAALVTWMMLGSSF-------
>MBB6464676_NuoL_Aminobacter_lissarensis
MMY------HFIVFLPLVGFLIAGLFG----RSIGAKASEYITSGLLVVSAVLSWFAFATVAL--
GDVEVFTVPVL-RFLDVGGIQADWALRIDTLTTVMLVVVNTVSALVHIYSIGYMHHDPH-------
RPRFFGYLSLFTFAMLMLVTSDNLVQMFFGWEGVGLASYLLIGFWYKKPSANAAAIKAFVVNRVGDFGFLLGIF
GLYVLFGSVEF--
STIFATAASYLPAVLNFLGYALDKHNALTIVCLLLFMGAMGKSAQVPLHTWLPDAMEGPTPVSALIHAATMVTA
GVFMLARLSPVFELSHTALTVVTFVGAFTAFFAATVGLVQNDIKRVIAYSTCSQLGYMFVALGMGFYSAAIFHL
FTHAFFKALLFLGSGSVIHAVSDEQDMRKMGGLRKLIPTTYWMMVIGTLALTGVGIPATIIG-----
TAGFFSKDAIIEGAFVGHNAVAGFAFAMLVIAACFTSFYSWRLIFMTFHGKPR--ATADVMHHV----------
---------------------------------------------------
HESPPVMLVPLFILAVGALFAGVIFHDQFIGE-AYNE---FW--
KGALFTLDTNHILHDFHSVPMWVKLAPFAAMIGGFAMAYLFYIRS-----------------------------
------------------------------
PEMPKVLAERHRGLYAFLLNKWYFDELYDFLFVRPAKWLGRFLWKK-GDGWLIDGFGADGIS-
ARVVDVTNRVVKLQTGYLYHYAFAMLIGVAALVTWMMLGSSF-------
>WP_068881841_NuoL_Paramesorhizobium_deserti
MLY------YAIVFLPLLGFLAAGLLG----KQLGVKACEYITSGFLVISAALSWVAFFQVAL--
GHGETLRIPVL-HWVNSGALTFDWAFRIDTLTAVMLVVVNTVSALVHIYSIGYMHHDPN-------
RPRFFAYLSLFTFAMLMLVTADNLVQMFFGWEGVGLASYLLIGFWYKKPSANAAAMKAFIVNRVGDFGFLLGIF
GVFVLFSAVNY--
DTIFAAAAQYLPAVLNFLGYSLDRQGAITIVCLLLFMGAMGKSAQFLLHTWLPDAMEGPTPVSALIHAATMVTA
GVFMVARMSPLFELSQTALTFVTIIGATTAFFAATVGLVQNDIKRVIAYSTCSQLGYMFVALGVGAYGAGIFHL
FTHAFFKALLFLGAGSVIHAVSDEQDMRRMGGLRKLIPTTYWMMIIGTVALTGLGIPFTSIG-----
TAGFFSKDAIIESSYASHAPAAGYAFVLLVVAALFTSFYSWRLIFMTFHGKPR--ATADVMHHV----------
---------------------------------------------------
HESPPVMLVPLFVLALGALLAGFLFHDYFFGH-HYVE---FW--
KGALFTGPENEILEEYHHVPFLVKIAPFVAMAIGFVIAWIFYIRS-----------------------------
------------------------------
PETPKELARRHPGLYQFLLNKWYFDELYDFLFVRPAKALGRLLWKG-GDGWLIDGFGPDGIS-
ARVVDVTNRVVKLQSGYLYHYAFAMLIGVAALVTWMMLGSSF-------
>WP_126010653_NuoL_Georhizobium_profundi
```

MY-------HAIVFLPLIGAIIGLFG----RSLGAKACEYITSGFLVFAAFLSWIAFFSVAY--
GDTDVIRVPVL-QWIQSGSMSVDWSFRIDTLTAVMLVVVNTVSALVHIYSIGYMHHDPH-------
RPRFFAYLSLFTFAMLMLVTSDNLLQMFFGWEGVGLASYLLIGFWYKKPTATAAAMKAFIVNRVGDFGFALGIF
GIFVLFGSISF--
DTIFAGAADFMPVVLNFLGYSLTRADALTVTCLLLFMGAMGKSAQFLLHTWLPDAMEGPTPVSALIHAATMVTA
GVFLVARMSPVFELSIDARTVVVIIGAITAFFAATVGLVQNDIKRVIAYSTCSQLGYMFVALGIGAYGAAIFHL
FTHAFFKALLFLGAGSVIHAVSDEQDMRRMGGLRKYIPRTYWMMIIGTVALTGVGIPGLYFG-----
TAGFFSKDAIIESAFVAENAAAGFAFTMLVIAALFTSFYSWRLIFMTFHGKPR--ASADVMHHV----------
---------------------------------------------------
HESPAIMMVPLYILAVGALLAGILFHKQFFGD-YYEG---FW--
QGALFTLPTNQILEEFHHVPFWVKLSPFMAMLIGLVTAWYMYIRS-----------------------------
------------------------------
PETPKELAAQHRGLYQFLLNKWYFDELYDFLFVRSSKALGRFLWKE-GDGRVIDGFGPNGVA-
ARVQDVTGRVVRLQSGYLYHYAFVMLIGVAALVTWMMLGSAF-------
>MBE7185388_NuoL_Methylobacterium_mesophilicum
MY-------QAIVFLPLLGFLVVGLFG----KRLGARPSELITSGFMVVAAVLSWLAFFTVAL--
GESEPVIVPLL-RFLDIGALQADWTLRIDTLTAVMLIVVNTISTLVHIYSIGYMHHDPH-------
RPRFFAYLSLFTFAMLMLVTSDNLVQMFFGWEGVGLASYLLIGFWYQKPSANAAAMKAFIVNRVGDFGFLLGIF
GVFVLFGTVDF--
STIFANAQTYIPAVFNFLGYQLDKQSALTVTCLLLFMGAMGKSAQVPLHTWLPDAMEGPTPVSALIHAATMVTA
GVFMLARLSPLFEHSQTALTVVTFIGALTAIFAATVGLVQNDIKRVIAYSTCSQLGYMFVALGLGFYSAAIFHL
FTHAFFKALLFLGSGSVIHAVSDEQDMRKMGGLRKLIPTTYWMMVIGTLALTGVGIPATLIG-----
TAGFFSKDVIIEGAFVGQNIFATMAYVLLVVAAVFTSFYSWRLIFMTFHGKPR--ASHEVMHHV----------
---------------------------------------------------
HESPPVMLVPLFVLAVGALLAGVLFEGYFVGE-EYAE---FW--
KGALFTLPENEILEAFHTVPLWVKLSPFVAMLIGFFTAYQFYIRS-----------------------------
------------------------------
PETPRRLAEAQPGLYQFLLNKWYFDELYDFLFVRPAKRIGHFLWKT-GDGRIIDGLGPDGLS-
ARVQDITRSVVRLQTGYLYHYAFVMLIGVAALVTWMMVGNNF-------
>WP_099304717_NuoL_Zhengella_mangrovi
MY-------HAIVFLPLIGFLIAGLFG----
RSIGAKASEYITSGLLVVSAVLSWVAFFTVALEGHGTQAMIIPVL-TFIQSGDLAVDWAL------
AVMLVVVNTVSALVHIYSIGYMHHDPD-------
RPRFFAYLSLFTFAMLMLVTSDNLVQMFFGWEGVGLASYLLIGFWYKKPSANAAAMKAFIVNRVGDFGFALGIF
GVFVLFGSVTF--
ETIFANAATYLPAVLSFLGHALDRQGAITIVCLLLFMGAMGKSAQFLLHTWLPDAMEGPTPVSALIHAATMVTA
GVFMVARLSPLFELSHTALVVVTVIGSITAFFAATVGLVQNDIKRVIAYSTCSQLGYMFAALGVGAYGAAIFHL
FTHAFFKALLFLGAGSVIHAVSDEQDMRRMGGLRKLIPTTYWMMIVGTIALTGLGIPGTFIG-----
TAGFFSKDIIIESVFASGSPVSAFSFTMLAVAALFTSFYSWRLIFMTFHGKPR--ATADVMHHV----------
---------------------------------------------------
HESPPVMLVPLFVLAAGALFAGVLFEGYFYGH-EYLA---FW--
KGALFTGPENELLNEFHHVPLLVKLSPFVAMVAGFLIAWWFYISN-----------------------------
------------------------------
PEMPRQVAQRHHGLYQFLLNKWYFDELYDFIFVRPAMWLGRFLWKK-GDGAVIDGLGPDGVA-
ARVVDVTNRVVKLQSGYLYHYAFAMLIGVAALVTWMMLGSAF-------
>WP_139975675_NuoL_Ochrobactrum_sp._CGA5
MLY------YAIVFLPLLGFLIAGLFG----NQIGAKASEYITSGFMVIVAILSWVVFFQIPL--
GHDAETVRIPVLHWVTSGTLSFDWALRIDTLTGVMLVVVNSVSALVHIYSIGYMHHDPH-------

```
RPRFFAYLSLFTFAMLMLVTSDNLVQMFFGWEGVGLASYLLIGFWFQKPSANAAAMKAFVVNRVGDFGFLLGIF
GVFALFQSVDY--
NTIFAAAANYLPAVLNFLGYQLDKQSAITITCLLLFMGAMGKSAQFLLHTWLPDAMEGPTPVSALIHAATMVTA
GVFMVARMSPIFELSQTALLVVTIIGATTAFFAATVALVQNDIKRVIAYSTCSQLGYMFAALGVGAYGAAVFHL
FTHAFFKALLFLCAGSVIHAVSEEQDMRRMGGLRKLIPITYWMMVIGTVAITGLGIPGTVIG-----
TAGFFSKDAIIESVFASHSAASGYASTLLIVAALFTSFYSWRLIFMTFHGKPR--ASAEVMHHV----------
---------------------------------------------------
HESPAVMFVPLLILGVGALFAGVVFKEFFFGH-EYAE---FW--
KGALFTSAANQILEEYHHVPLWVKLSPFISMVLGFIVAWIFYIRS-----------------------------
------------------------------
PEMPKALAARHRGLYQFLLNKWYFDELYDFLFVRPARWLGRVFWKG-GDGWLIDGFGPDGIS-
ARVLDVTNRVVKMQSGYLYHYAFAMLIGVAALVTWMMLGSSF-------
>WP_173191174_NuoL_Mesorhizobium_sediminum
MY-------HAIVFLPLLGFLIAGLFG----RSIGAKASEYITSGFLIIAAALSWLAFFTVGF--
GDGEAFTVPVL-RFIQSGGLDVAWALRIDTLTVVMLVVVNTVSALVHVYSIGYMHHDPH-------
RPRFFAYLSLFTFAMLMLVTADNLVQMFFGWEGVGLASYLLIGFWFKKPSANAAAIKAFVVNRVGDFGFLLGLF
GVFVLFGSVNF--
DTIFASAAAVAEAIFTFLGWSLTQGGALTAICLLLFMGAMGKSAQVPLHTWLPDAMEGPTPVSALIHAATMVTA
GVFMLARLSPIFELSHAALLFVTFIGAFTAFFAATVGLVQNDIKRVIAYSTCSQLGYMFVALGMGFYSAAIFHL
FTHAFFKALLFLGSGSVIHAVSDEQDMRKMGGLRKLIPTTYWMMVIGTLALTGVGIPLTVIG-----
TAGFFSKDAIIEGAFAGHNGLAGFAFTLLVVAAVFTSFYSWRLIFMTFHGKPR--ASSEVMHHV----------
---------------------------------------------------
HESPPVMLVPLFILAVGALFAGIVFHDWFIGY-HHDA---FW--
KGALFILPDNHILHEFHEVPLWVKLAPLVAMIVGFVTAYRFYIQS-----------------------------
------------------------------
PDAPRQLAERHQGLYQFLLNKWYFDELYDFMFVRPAKRLGRFLWKK-GDGTVIDGMGPDGIA-
ARVVDITDRVVKLQTGYLYHYAFAMLIGVAALVTWMMLGGSY-------
>WP_110030771_NuoL_Hoeflea_marina
MY-------HAIVFLPLIGAIIAGLFG----RSIGAKASEYVTTLFLIVAAVLSWVAFFTVAM--
GDAETIKIPVM-RWIDSGTLNVEWAFRIDTLTAVMLVVVNTVSSLVHVYSIGYMHHDPH-------
RPRFFAYLSLFTFAMLMLVTSDNLVQMFFGWEGVGLASYLLIGFWYKKPSAAAAAMKAFIVNRVGDFGFALGIF
GVFVLFGSVGF--
DTIFAGAADFIPQEMTLFGMSLDKGHAMTGVCLLLFMGAMGKSAQFLLHTWLPDAMEGPTPVSALIHAATMVTA
GVFLVARMSPLFELSQSALTVVVIIGAITAFFAATVGLVQNDIKRVIAYSTCSQLGYMFVALGIGAYGAAIFHL
FTHAFFKALLFLGAGSVIHSVSDEQDMRRMGGLRKHIPATYWMMIIGTVALTGLGIPGTIIG-----
TSGFFSKDAIIESAFMAHNPAAGFAFTLLVVAALFTSFYSWRLIFMTFHGKPR--ASADVMHHV----------
---------------------------------------------------
HESPPVMLVPLYILAAGALLAGFIFKDYFFGH-HYAE---FW--
KGALFTLPENEIVEEYHHVPLWVALSPFLAMLIGFATAWFFYIRS-----------------------------
------------------------------
PETPRRLAGEHQGLYQFLLNKWYFDELYDFLFVRPAKWLGTFLWKQ-GDGRVIDGFGPDGIA-
ARVQDISGRAVRLQTGYLYHYAFVMLIGIAALVTWMMLGSAF-------
>WP_066175677_NuoL_Hoeflea_olei
MY-------QAIVFLPLIGALIAGLGG----RAIGAKASEYVTTLFLIVAALLSWVAFFSVAL--
GDGEVVRIQVL-RWMDSGSMTADWALRIDTLTAVMLVVVNTVSCLVHVYSIGYMHHDPH-------
RPRFFAYLSFFTFAMLMLVTSDNLLQMFFGWEGVGVASYLLIGFWFKKPSANAAAMKAFIVNRVGDFGFLLGMF
GLFVLFGSISF--
DTIFASVADYLPAEITLFGMELDKAHALTAVCLLLFMGAMGKSAQFLLHTWLPDAMEGPTPVSALIHAATMVTA
```

```
GVFLVARMSPVFEHSPDALTFVTWIGAITAFFAATVGLVQNDIKRVIAYSTCSQLGYMFVALGIGAYGGAVFHL
FTHAFFKALLFLGAGSVIHAVSDEQDMRRMGGLRKHIPLTYWMMIIGTIALTGVGIPGTMIG-----
TAGFFSKDIIIESAYVAHNSASGVAFGLLVVAALFTSFYSWRLIFMTFHGKPR--ASSDVMHHV----------
---------------------------------------------------
HESPMVMLVPLFILAVGALFAGVLFEGYFFGH-HYAE---FW--
KGALFTGPENEILEEFHHVPLWVKLSPFVAMLLGLVTAWFFYIRS-----------------------------
------------------------------
PETPKRLAEDHQMLYRFLLNKWYFDELYDFIFVRPAKALGRLLWKQ-GDVRIIDGFGPNGVA-
ARAQDITGWVVRLQTGYLYHYAFVMLIGIAALVTWMMLGSAF-------
>WP_097104131_NuoL_Hoeflea_halophila
MY-------HAIVFLPLIGALIAGFGG----RAIGAKASEYVTTLFLIVAALLSWVAFLAVAM--
GDGEMIRVPIM-RWIDSGSLNVEWALRIDTLTAVMLVVVNTVSCLVHVYSIGYMHHDPH-------
RPRFFAYLSFFTFAMLMLVTSDNLLQMFFGWEGVGVASYLLIGFWYKKPSANAAAMKAFIVNRVGDFGFLLGIF
GLFVLFGSASF--
DAIFAGVADYLPSEITLFGMHLDKANALTGVCLLLFMGAMGKSAQFLLHTWLPDAMEGPTPVSALIHAATMVTA
GVFLVARMSPVFEYAPDALTVVTWIGAITAFFAATVGLVQNDIKRVIAYSTCSQLGYMFVALGIGAYGGAIFHL
FTHAFFKALLFLGAGSVIHAVSDEQDMRRMGGLRKKIPMTYWMMIIGTIALTGVGIPGTLIG-----
TAGFFSKDIIIESAFVAQNSAAGVAFGLLVIAALFTSFYSWRLIFMTFHGKPR--ASADVMHHA----------
---------------------------------------------------
HESPPVMLVPLFILAVGALLAGVVFKEFFFGH-DYDA---FW--
KGALFTLPGNTIVEEFHYVPLWVKLSPFVAMLIGLVSAWYFYIRS-----------------------------
------------------------------
PETPKRLAEDHQMLYQFLLNKWYFDELYDFIFVRPAKWLGSVLWKQ-GDVRVIDGFGPNGIA-
ARVQDVTGRITRLQTGFLYHYAFVMLIGIAALVTWMMLGSAF-------
>WP_007197627_NuoL_Hoeflea_phototrophica
MY-------QAIVFLPLIGALIAGFGG----RAIGAKASEYVTTLFLIVAAVLSWVAFFTVAM--
GETEMIRVPVM-RWIDSGSFSVDWALRIDTLTAVMLVVVNTVSSLVHLYSIGYMHHDPH-------
RPRFFAYLSLFTFAMLMLVTSDNLLQMFFGWEGVGVASYLLIGFWYKKPSANAAAMKAFIVNRVGDFGFLLGMF
GVFMLFGSISY--
DAIFAGAADFIPVEITLFGMHLDKAHALTAVCLLLFMGAMGKSAQFLLHTWLPDAMEGPTPVSALIHAATMVTA
GVFLVARMSPLFELSVDALTVVVITGAITAFFAATVGLVQNDIKRVIAYSTCSQLGYMFVALGIGAYGAAVFHL
FTHAFFKALLFLGAGSVIHAVSDEQDMRRMGGLRKKIPITYWMMIIGTIALTGVGIPGTVIG-----
TAGFFSKDAIIESAYVAHNPAAGFAFGLLVIAALFTSFYSWRLIFMTFHGKPR--ASADVMHHA----------
---------------------------------------------------
HESPPVMLVPLFVLAVGALFAGVIFKEYFFGH-HYTE---FW--
GSALFTLPDNKIVEEFHHVALWVKLSPFVAMLLGLVTAWYFYIRS-----------------------------
------------------------------
PGTPKQLAEDHSLLYQFLLNKWYFDELYDFIFVRPAKWLGRFLWKQ-GDGRVIDGYGPDGIA-
ARVQDITGRVVKLQTGYLYHYAFVMLIGIAALVTWMMLGSAFR------
>WP_011580373_NuoL_Chelativorans
MY-------QAIVFLPLAGFLIAGLFG----RSIGAKGSEYITSGLLVVSAFLSWIAFFQVGL--
GDGEAITIPLL-RFISSGALDVDWALRIDTLTAVMLVVVNTVSALVHIYSIGYMHHDPD-------
RPRFFAYLSLFTFAMLMLVTSDNLVQMFFGWEGVGLASYLLIGFWYKKPSANAAAMKAFIVNRVGDFGFALGIF
GVFVLFGSINF--
DTIFANTAAFMPGELTFLGYALDPHSALTVVCLLLFLGAMGKSAQVPLHTWLPDAMEGPTPVSALIHAATMVTA
GVFMVARLSPIFELSQTALTVVTAVGAFTAFFAATVGLVQNDIKRVIAYSTCSQLGYMFVALGTGFYSAAIFHL
FTHAFFKALLFLGSGSVIHAVSDEQDMRKMGGLRKLIPQTYWMMIIGTLALTGVGIPGTLIG-----
TAGFFSKDAIIEGSFVAHNAVAPIAFTLLVVAAVFTSFYSWRLIFMTFHGKPR--ASSDVMHHV----------
```

```
---------------------------------------------------
HESPYVMLVPLFLLAIGAIFAGLVFEGFFVGH-EYDH---FW--
KGALFTLPDNHMVEEFHHVPLWVKLSPFVAMLFGLLVAWQFYIRS-----------------------------
------------------------------
PESPKRLAERHRALYAFLLNKWYFDEVYDFLFVRPAQRLGRFLWKK-GDGWFIDGFGPDGIS-
ARVVDVANRVVKLQTGYLYHYAFAMLIGIAAFVTWMMLGSSF-------
>WP_198476026_NuoL_Aquamicrobium_sp._cd-1
MY-------HAIVFLPLAGFLIAGIFG----RAIGAKASEFVTSGFLLVAALLSWVAFFQVGL--
GDTAAFSVPVL-RFIEVGSLDVSWAFRIDTLTTVMLVVVNSVSALVHAYSIGYMHHDPH-------
RPRFFAYLSLFTFAMLMLVTSDNLVQMFFGWEGVGLASYLLIGFWFKKPSANAAAIKAFVVNRVGDFGFLLGMF
GVYVMFGSVNF--
DTIFANAAAIAQNPFQFLGWSLTQAGALTAICLLLFMGAMGKSAQVPLHTWLPDAMEGPTPVSALIHAATMVTA
GVFMLARLSPVFELSDTALLVVTIIGAITAFFAATVGLVQNDIKRVIAYSTCSQLGYMFVALGLGFYSAAVFHL
FTHAFFKALLFLGSGSVIHAVSDEQDMRKMGGLRKLIPTTYWMMIIGTIALTGVGLPATYIG-----
TAGFFSKDAIIEGAFASSSAAATFAFIMLVVAAMFTSFYSWRLIFLTFHGQPR--ASEEVMHHV----------
---------------------------------------------------
HESPPVMLVPLYLLAVGALFSGFVFYGYFLGD-NYDG---FW--
QGALFTLEDNHILHDYHSVPFWVKASPFAAMILGFLLAYQFYIRS-----------------------------
------------------------------
PETPKRLAAQHSGLYQFLLNKWYFDELYDFIFVRPAKRLGHFLWKK-GDGVVIDGFGPNGIA-
ARVVDITDRVVKMQTGYLYHYAFAMLIGVAALVTWMMLGGSL-------
>WP_053998662_NuoL_Ahrensia_marina
MY-------HAIVFLPLLGAIVAGFGG----RAIGAKASEYITSGFMVIAAILSWVAFLTVAM--
AHDDHSATVQVMRWIQVGSLDVNWAFRIDTLTAVMLVVVNTVSTLVHIYSIGYMHHDPH-------
RPRFFAYLSLFTFAMLMLVTSDNLVQMFFGWEGVGLASYLLIGFWYKKPSANSAAMKAFIVNRVGDFGFALGIF
GVFVLFGSVNF--
DTIFANAATYLPAELTFLGYSLDKSHALTVVCLLLFMGAMGKSAQFLLHTWLPDAMEGPTPVSALIHAATMVTA
GVFLVARMSPIFELSQTALTVVVIIGATTAFFAATVGLVQNDIKRVIAYSTCSQLGYMFVALGIGAYGAAVFHL
FTHAFFKALLFLGAGSVIHAVSDEQDMRRMGGLRKLIPTTYWMMIIGTVALTGVGIPGTLIG-----
TAGFFSKDAIIESAFVAQNSAAGYAFVMLVVAALFTSFYSWRLIFMTFHGKPR--ATADVMHHV----------
---------------------------------------------------
HDSPPVMLVPLFILAVGAILAGVVFHGYFFGH-HYDE---FW--
KGALFTLPDNNIVDEFHNVPFLVKWSATIVMLLGLLGAWYMYIRS-----------------------------
------------------------------
PQTPASLAAQHNGLYKFLLNKWYFDELYDFLFVRPAKRLGSFLWKR-GDGWLIDGFGPNNIA-
ARVSGLAQRAVKVQTGFLYHYAFAMLIGVAALVTWIMLGGIH-------
>WP_127709278_NuoL_Sinorhizobium_meliloti_USDA1025
MDTIV----KAIVFLPLIGFLIAGLLG----TQIGAKASEYVTSGLMIVTAALSWFVFFHVAL--
GEEEMIKVSVL-RWIQSGSFDVEWAFRVDTLTAVMFVVVNTVSTLVHIYSIGYMHHDPN-------
RPRFFAYLSLFTFAMLMLITSDNLLQMFFGWEGVGLASYLLIGFWYKKPSANAAAMKAFIVNRVGDFGFSLGIF
CVFVLFGSINF--
ETIFAAAQNYLPAEINLFGMQLDKGHALTATCLLLFMGAMGKSAQFLLHTWLPDAMEGPTPVSALIHAATMVTA
GVFLVARMSPLFELSPDALTVVTLIGAITAFFAATVGLVQNDIKRVIAYSTCSQLGYMFVALGVGAYGAAIFHL
FTHAFFKALLFLGAGSVIHAVDGEQDMRYMGGLRTHIPVTYWMMFIGTIALTGVGIPGTVIG-----
TAGFFSKDAIIESTFASHSVVSGLAFALLVIAALFTSFYSWRLTFMTFHGKPR--ASSDVMHHV----------
---------------------------------------------------
HESPQVMLVPLYILAAGALVAGFLFHDYFFGH-HYAE---FW--
QGALFTSAENELLEEYHHVPLWVKWSPFAAMALGLFTAWYMYIRS-----------------------------
```

```
------------------------------
PETPKYLAEQHRGLYQFLLNKWYFDELYDFLFVRSAKRLGTFLWKE-GDGRVIDGYGPNGIA-
ARVLDVTDRVVRLQTGYLYHYAFAMLIGIAALVTWMMLGSSF-------
>WP_132561602_NuoL_Rhizobium_sullae
MFLY-----KAIVFLPLIGAIIAGLFG----
RAIGAKASEYVTCGLMIVAAVLSWYVFVTVGMGHLEGGPIKVEVL-
RWIQSGGIDASWSLRIDTLTSVMLIVVNTVSTLVHVYSIGYMHHDPH-------
RPRFFAYLSLFTFAMLMLVTSDNLAQMFFGWEGVGLASYLLIGFWYKKPSANAAAIKAFVVNRVGDFGFVLGIA
GIFVLFGSINL--
DTIFANASSFAPQELNLFGMQLDKAHALTGICLLLFMGAMGKSAQFLLHTWLPDAMEGPTPVSALIHAATMVTA
GVFLVARMSPLFELSPDALVFVTFIGAITAFFAATVGLVQNDIKRVIAYSTCSQLGYMFVALGVGAYGAAIFHL
FTHAFFKALLFLGAGSVIHAVDGEQDMRYMGGLRTHIPVTFWMMTIGTLALTGVGIPFTPVG-----
FAGFFSKDVIIEATYASHSPVAGFAFTLLVIAALFTSFYSWRLAFLTFFGTPR--ASHEVMHHV----------
---------------------------------------------------
HESPQVMLVPLYLLAIGAVLSGVVFEGYFYGH-HYVE---FW--
KGALFTGAENELLEEFHHVPAWVALSPFIAMLLGFVTAWYLYIQS-----------------------------
------------------------------
PSTPRVLAQQHRVLYQFLLNKWYFDELYDFLFVRSAKALGRFLWKK-GDIGVIDAYGPNGVA-
ARVVDVTNRVVRLQTGYLYHYAFAMLIGIAALVTWMMLGSSF-------
>WP_020921010_NuoL_Rhizobium_etli_bv.mimosae
MFLY-----KAIVFLPLIGAIVAGLFG----
RAIGAKASEYVTSGLMIVAGILSWIVFFNVGMGHLEGGPIKVEVL-
RWIQSGGIDVSWSLRVDTLTSVMLIVVNTVSTLVHIYSIGYMHTDPH-------
RPRFFAYLSLFTFAMLMLVTADNLAQMFFGWEGVGLASYLLIGFWFKKPSATAAMKAFIVNRVGDFGFVLGIA
GVFVLFGSINL--
DTIFANASNFAPHELNLFGMQLDKAHALTGICLLLFMGAMGKSAQFLLHTWLPDAMEGPTPVSALIHAATMVTA
GVFLVARMSPLFELSPDALTVVTVIGAITAFFAATVGLVQNDIKRVIAYSTCSQLGYMFVALGVGAYGAAIFHL
FTHAFFKALLFLCAGSVIHAVDGEQDMRYMGGLWPHIKVTGGLMIIGTLAITGVGIPFTPFG-----
LAGFFSKDVIIEATYASHSPVAGFAFSLLVIAALFTSFYSWRLIFMTFFGKPR--ASHEVMHHV----------
---------------------------------------------------
HESPQVMLVPLYLLAVGAIFAGVIFEGRFYGE-EYAE---FW--
KGALFTGAENELVEEFHHVPALVGLSPFIAMLIGFVIAWYMYIRS-----------------------------
------------------------------
PQTPRVLAKQHRVLYHFLLNKWYFDELYDFLFVRTARALGRFLWKK-GDVGVIDTYGPNGVA-
ARVVAVTDRVVRLQTGYLYHYAFAMLLGIAALVTWMMLGSSF-------
>WP_183260520_NuoL_Aminobacter_niigataensis
MY-------QLIVFLPLLGFLIAGLFG----RSIGAKASEYITSGLLVVSAVLSWVAFITVGL--
GDVEVFTVPVL-RFIDSGGLQANWALRIDTLTVVMLVVVNTVSALVHIYSIGYMHHDPH-------
RPRFFAYLSLFTFAMLMLVTSDNLVQMFFGWEGVGLASYLLIGFWYKKPSANAAAIKAFVVNRVGDFGFLLGIF
GLFVLFGSVEF--
STIFAGAASYLPAELTFLGYALDKQGALTIVCLLLFMGAMGKSAQVPLHTWLPDAMEGPTPVSALIHAATMVTA
GVFMLARMSPVFELSASALTVVTFIGAFTAFFAATVGLVQNDIKRVIAYSTCSQLGYMFVALGMGFYSAAIFHL
FTHAFFKALLFLGSGSVIHAVSDEQDMRKMGGLRRLIPTTYWMMVIGTLALTGVGIPATIIG-----
TAGFFSKDAIIEGAFVGHNAVAGLAFAMLVVAACFTSFYSWRLIFMTFHGKPR--ATADVMHHV----------
---------------------------------------------------
HESPPVMLVPLFVLAVGALFAGVVFHDQFIGE-AYNE---FW--
KGAIFTLETNHILHDFHGVPMWVKLAPFAAMISGFLVAYLFYIRS-----------------------------
------------------------------
```

```
PEMPKVLAERHRGLYAFLLNKWYFDELYDFLFVRPAKWLGRFLWKK-GDGWLIDGFGADGIS-
ARVVDVTNRVVKLQTGYLYHYAFAMLIGVAALVTWMMLGSSF-------
>WP_109613072_NuoL_Pseudaminobacter_salicylatoxidans
MY-------QAIVFLPLLGFLIVGLFG----NSLGARASEYITSGFLVIAAVLSWVAFFSVGI--
GSTEAFTVPLL-RFIDSGTLQSDWALRVDTLTVVMLVVVNTVSALVHIYSIGYMHHDPH-------
RPRFFAYLSLFTFAMLMLVTSDNLVQMFFGWEGVGLASYLLIGFWYKRPSANAAAIKAFVVNRVGDFGFILGIA
GVFLLFGSVNF--
DTIFATAASYVPVEFNFLGWDVDRAGALTVICLLLFMGAMGKSAQVPLHTWLPDAMEGPTPVSALIHAATMVTA
GVFMLARLSPIFELSHTALLVVTFIGAITAFFAATVGLVQNDIKRVIAYSTCSQLGYMFVALGMGFYSAAIFHL
FTHAFFKALLFLGSGSVIHAVSDEQDMRKMGGLKKLIPTTYWMMVIGTLALTGVGIPGTIIG-----
TAGFFSKDAIVEGAFAGIGHNAVAGFAFTLLVIIFTSFYSWRLIFMTFHGKPR--ATADVMHHV----------
-----------------------------------------------------
HESPPVMLVPLFILAVGALMAGVVFHDQFVGE-GYDA---FW--
KTALYMLPENHILHEVHDVPLWVKLSPFIAMVIGFVVAYQFYIRK------------------------------
-------------------------------
PETPKYLAAQHRGLYAFLLNKWYFDELYDFLFVRTAKRLGTFLWKK-GDGAVIDGLGPDGVS-
AWVIDVTNRVVKLQTGYLYHYAFVMLIGVAALVTWMMLGSSF-------
>WP_189486644_NuoL_Limoniibacter_endophyticus
MF-------HAIVFLPLVGFLIAGLFG----RKIGATASEYVTSAFVGIAALLSWVVFFQYAL--
GDAEKVSVPVL-TFLQSGGLDVSWALRIDTLTAVMLVVVNSVSTLVHVYSIGYMHHDPH-------
RPRFFAYLSLFTFAMLSLVTADNLVQMFFGWEGVGLASYLLIGFWFKKQSAGAAAMKAFIVNRVGDFGFLLGIF
GLYVLTGSVNF--
DTIFANIQSMYAGELNFLGYALNPQAALTIVCLLLFMGAMGKSAQVPLHTWLPDAMEGPTPVSALIHAATMVTA
GVVMLARMSPVFEFSHTALTVVTIIGAITAIFAATVGLVQNDIKRVIAYSTCSQLGYMFVALGLGFYSAAVFHL
FTHAFFKALLFLGSGSVIHAVSDEQDMRKMGGLRKLIPNTYWMMIIGTIALTGVGIPGTVIG-----
TAGFFSKDAIIEGAFSAHGAAAGFAFILLTVAACMTSFYSWRLIFMTFHGKPR--ATADVMSHV----------
-----------------------------------------------------
HESPPVMLIPLYILAVGALFAGFLFHDAFIGE-AYGE---FW--
KVSLFTLESNHILHEFHHVPFWVKLAPFVAMLLGFALAYQFYIRS------------------------------
-------------------------------
PETPVKLAERHRGLYAFLLNKWYFDELYDFLFVRPANCLGRILWKT-GDGTLIDGLGPNGVA-
ARVVDVTNRVVKMQTGYLYHYAFAMLIGVAALVTWMMLGSSF-------
>WP_163269623_NuoL_Chelativorans_alearense
MY-------QAIVFLPLAGFLIAGLFG----RSIGAKGAEYITSGLLVISALLSWVAFFTFAL--
GDGEAFTVPVL-RFISSGALDVDWALRIDTLTVVMLVVVNTVSSLVHIYSIGYMHHDPH-------
RPRFFAYLSLFTFAMLALVTADNLVQMFFGWEGVGLASYLLIGFWYKKPSANAAAIKAFIVNRVGDFGFALGIF
GVFVLFGSINL--
DTIFANAATFLPAELNFLGYALTNQAALTVVCLLLFLGAMGKSAQVPLHTWLPDAMEGPTPVSALIHAATMVTA
GVFMVARLSPLFELSSTALTVVTAVGAFTAFFAATVGLVQNDIKRVIAYSTCSQLGYMFVALGTGFYGAAIFHL
FTHAFFKALLFLGAGSVIHAVSDEQDMRKMGGLRRLIPRTYWMMIIGTLALTGVGIPGTLIG-----
TAGFFSKDAIIEGSFVAHNAVAPIAFGLLVVAAIFTSFYSWRLIFMTFHGKPR--ASADVMHHV----------
-----------------------------------------------------
HESPLVMLVPLFLLAAGAVFAGILFEGFFVGH-EYDH---FW--
KGALFTLPGNEILEEFHHVPLWVKLSPFAAMLLGLLVAWQFYIRS------------------------------
-------------------------------
PGSPKELAKRQRVLYAFLLNKWYFDEVYDFLFVNPAKRLGTFLWKR-GDGWLIDGFGPDGIS-
ARVIDVTNRVVKMQTGYLYHYAFAMLIGIAALVTWMMLGSSF-------
>WP_159586339_NuoL_Chelativorans_xinjiangense
```

```
MY-------QAIVFLPLAGFLIAGLFG----RSIGAKGAEYITSGLLVISALLSWIAFFTFAL--
GDGEAFTVPML-RFISSGALDVDWALRIDTLTVVMLVVVNTVSSLVHIYSIGYMHHDPH-------
RPRFFAYLSLFTFAMLALVTADNLVQMFFGWEGVGLASYLLIGFWYKKPSANAAAIKAFIVNRVGDFGFALGIF
GVFVLFGSINL--
DTIFANAATFLPAELNFLGYALTNQAALTVVCLLLFLGAMGKSAQVPLHTWLPDAMEGPTPVSALIHAATMVTA
GVFMVARLSPLFELSSTALAVVTAVGAFTAFFAATVGLVQNDIKRVIAYSTCSQLGYMFVALGTGFYSAAIFHL
FTHAFFKALLFLGAGSVIHAVSDEQDMRKMGGLRRLIPQTYWMMIAGTLALTGVGIPGTIIG-----
TAGFFSKDAIIEGSFVAHNAVAPIAFGLLVVAAIFTSFYSWRLIFMTFHGKPR--ASADVMHHV----------
---------------------------------------------------
HESPLVMLVPLFLLAAGAVFAGVVFEGFFVGD-EYDH---FW--
KGALFTLPGNEILEEFHHVPLWVKLSPFAAMLLGLFVAWQFYIRS-----------------------------
------------------------------
PGSPKELAKRQRVLYAFLLNKWYFDEIYDFLFVNPAKRLGTFLWKR-GDGWLIDGFGPDGIS-
ARVIDVTNRVVKMQTGYLYHYAFAMLIGVAALVTWMMLGSSF-------
>WP_121644644_NuoL_Notoacmeibacter_ruber
MF-------HLIVFLPLIGFLIAGLFG----RSIGAKASEYVTTGFLYVAAVLSWIAFFDFGL--
GHEEAMVVPVM-QWMNIGSLAVDWAFRIDTLTVVMLVVVNTVSALVHTYSIGYMHHDPQ-------
RPRFFAYLSLFTFAMLMLVTSNNLVQMFFGWEGVGLASYLLIGFWYKKPSANAAAMKAFIVNRVGDFGLLGIF
GIFLLFGSVNF--
DTIFANARQFDPSGMSFFGYGVPAATAITVTCLLLFMGAMGKSAQVPLHTWLPDAMEGPTPVSALIHAATMVTA
GVVMVARLSPIFELSQTALVVVTVVGAITAFFAATVGLVQNDIKRVIAYSTCSQLGYMFVALGVGAYGAAIFHL
FTHAFFKALLFLGAGSVIHAVSDEQDMRFMGGLRKHIPATYWMMIIGTLALTGVGIPFTAIG-----
TAGFFSKDVIIETAYASHSVVSTFAFVMLVVAACFTSFYSWRLIFMTFHGTPR--ATADVMHHV----------
---------------------------------------------------
HESPWVMLIPLFILAAGAIFAGLLFEGFFYGH-EYGE---FW--
KESLFVAGGNELLEEFHHVPVWVALSPFIAMLLGLVFAYWFYIRS-----------------------------
------------------------------
PGTPKALAQSQSGLYAFLLNKWYFDELYDFLFVRPAKWLGRFLWKR-GDGTVIDGLGPDGIS-
ARVVDVTNRVVRLQSGYLYHYAFAMLIGVAALVTWMMLGSAF-------
>WP_131616112_NuoL_Roseitalea_porphyridii
MY-------HAIVFLPLLGALIAGFGG----RALGAKACEYVTSGFLVIAAILSWIALITVGF--
SEADPRTVQVM-RFMDSGNLDVDWAFRIDTLTVVMLVVVTTVSSLVHIYSIGYMHHDPH-------
RPRFFAYLSLFTFAMLMLVTADNLIQMFFGWEGVGLASYLLIGFWYKKASANNAAMKAFIVNRVGDFGFILGIF
GVFVLFGSVNF--
DTIFANAAAFASVDLTFLWMDLTAGEALTAVCLLLFLGAMGKSAQFLLHTWLPDAMEGPTPVSALIHAATMVTA
GVFMVARLSPIFEYSQDALTVVTIVGAITAFFAATVGLVQNDIKRVIAYSTCSQLGYMFVALGIGAYGAAMFHL
FTHAFFKALLFLGAGSVIHAVSDEQDMRKMGGLRKHIPMTYWMMIIGTVALTGVGIPGTYIG-----
TAGFFSKDAIIESAFVGQNAAAGIAFGLLVIAALFTSFYSWRLIFMTFHGKPR--ATADVMHHV----------
---------------------------------------------------
HESPPVMLVPLYVLAAGALFAGVLFYSAFVGH-GHHEYNDFF--
RTALFAGPDNHVLDEFHDVPYWVKWSPFVVMAIGLVTAWYMYIRS-----------------------------
------------------------------
PETPAALARQHAGLYRFLLNKWYFDELYDFLFVRPAKRLGTFLWKR-GDGWLIDGFGPDGVA-
RRVTDVTGRVVKWQTGYIYHYAFAMVIGIAALVTWMMLGGVL-------
>WP_109765985_NuoL_Oceaniradius_stylonematis
MY-------HAIVFLPLLGALIAGFGG----RALGAKACEYITSGFLVIAAILSWIALITVGF--
SEAEPRTVQVM-RFIGSGNLDVNWAFRIDTLTVVMLVVVNTVSSLVHIYSIGYMHDDPH-------
RPRFFAYLSLFTFAMLMLVTADNLVQMFFGWEGVGLASYLLIGFWYKKASANNAAMKAFIVNRVGDFGFILGIF
```

```
GVFVLFGSVNL--
DTIFANAAAFASVDFRFLWMDLTAGEALTVVCLLLFLGAMGKSAQFMLHTWLPDAMEGPTPVSALIHAATMVTA
GVFMVARLSPIFEFSPDALTVVTIVGAITAFFAATVGLVQNDIKRVIAYSTCSQLGYMFVALGVGAYGAAMFHL
FTHAFFKALLFLGAGSVIHAVSDEQDMRKMGGLRKHIPMTYWMMIIGTIALTGVGIPGTYIG-----
TAGFFSKDAIIESAFVGHNAAAGFAFGMLLIAAVMTSFYSWRLIFMTFHGKPR--ATADVMHHV----------
---------------------------------------------------
HESPPVMLVPLYILAVGALLSGFFFYSAFVGH-GHHEYNDFF--
RTALFAGPDNHILEEFHEVAFWVKASPAVAMLIGLVTAWYMYIRS-----------------------------
------------------------------
PETPAALARQHRGLYQFLLNKWYFDELYDFLFVNPAKRLGTFLWKR-GDGWLIDGFGPNGVA-
KRVTDVTGRVVRLQSGYLYHYAFAMMIGLAALITWMMLGGAL-------
>WP_094078056_NuoL_Notoacmeibacter_marinus
MF-------HLIVFLPLIGFLIAGLFG----RSIGAKASEYLTTGLLYVSAVLSWIAFFDFGL--
GHEEAMVVPVM-TWMNVGSLAVDWSFRIDTLTVVMLVVVNTVSALVHTYSIGYMHHDPH-------
RPRFFAYLSLFTFAMLMLVTSNNLVQMFFGWEGVGLASYLLIGFWYKKPSANAAAMKAFIVNRVGDFGFALGIF
GIFLLFGSVNF--
DTIFANARQFDPAGIDFFGYAVPAATAITATCLLLFMGAMGKSAQVPLHTWLPDAMEGPTPVSALIHAATMVTA
GVVMVARLSPIFELSTTALVVVTVVGAITAFFAATVGLVQNDIKRVIAYSTCSQLGYMFVALGVGAYGAAIFHL
FTHAFFKALLFLGAGSVIHAVSDEQDMRYMGGLRKHIPATYWMMIIGTIALTGVGIPFTMIG-----
TAGFFSKDIVIETAFASHSVVSTFAFVMLVIAACFTSFYSWRLIFMTFHGTPR--ATADVMHHV----------
---------------------------------------------------
HESPWVMLAPLFVLAAGALFAGVLFKEFFYGH-EYAE---FW--
KEALFVAGGNELLEEFHHVPAWVALSPFIAMIVGLLVAYWFYIRS-----------------------------
------------------------------
PGMPKALAQSQSGLYKFLLNKWYFDELYDFLFVRPAMWLGRFLWKR-GDGTVIDGMGPDGIS-
ARVVDITNRVVRLQSGYLYHYAFAMLIGVAALVTWMMLGSAL-------
>WP_170141804_NuoL_Ciceribacter_lividus
MILY-----KAIVFLPLIGALIAGLLG----RQIGAKASEYITSGLMIVTAVLSWVVFVNVGF--
GEGHEVIKVSVLRWIQSGGIDVEWSLRIDTLTAVMLVVVNSVSTLVHVYSIGYMHHDPD-------
RPRFFSYLSLFTFAMLMLVTSDNLLQMFFGWEGVGLASYLLIGFWFKKPSACAAAMKAFIVNRVGDFGFILGIA
GIFVLFGSINF--
ETIFAAAQSYLPAEINLFGMSLDKAHAMTAVCLLLFMGAMGKSAQFLLHTWLPDAMEGPTPVSALIHAATMVTA
GVFLVARMSPLFELSPNALTFVTLIGAITAFFAATVGLVQNDIKRVIAYSTCSQLGYMFVALGVGAYGAAIFHL
FTHAFFKALLFLGAGSVIHAVDGEQDMRYMGGLRKHIPLTFWAMTIGTLALTGVGIPGTMIG-----
FAGFFSKDVIIESAYASHSAVSGFAFTLLVIAALFTSFYSWRLAFLTFFGKPR--ASADVMHHV----------
---------------------------------------------------
HESPMVMLAPLALLALGAVFAGVAFEGYFFGH-EYGE---FW--
KGALFTLPENEILEQFHGVPTWVKWSPFVAMLTGFVTAWYMYIKS-----------------------------
------------------------------
PSTPKVLAEQHRVLYQFLLNKWYFDELYDLLFVRSAKALGRFLWKK-GDVGVIDTYGPNGVA-
AAVVDVTQRVVRLQSGYLYHYAFAMLIGVAALITWMMLGSSF-------
>WP_068082792_NuoL_Pseudovibrio_stylochi
MY-------SAIVFLPLVGFLVVGLFG----RALGAKASEFITSSLLVIAAILSWIAFFSVGF--
GSTEKVQLEIL-QWISSGDLSINWTIRVDTLTAVMLVVVNTVSALVHIYSIGYMHHDPH-------
RPRFFAYLSLFTFAMLSLVTSDNLLQMFFGWEGVGLASYLLIGFWFKKPSANAAAMKAFVVNRVGDFGFALGIC
GIYILFGSIDY--
DVIFQNVNSVAGQELTFLGTGLAADSAITVICLLLFMGAMGKSAQLLLHTWLPDAMEGPTPVSALIHAATMVTA
GVFMVARLSPIFEYSHTALTVVTFVGATTAFFAGTIGCVQNDIKRVIAYSTCSQLGYMFAALGVGAYGAAVFHL
```

```
FTHAFFKALLFLCAGSVIHAVSDEQDMRRMGGLRKLIPVTYWTMFIGTLALTGFGIPLAHLGELYIGTAGFVSK
DAIIESTFAGHNMFSQYAFWMTIIAAFLTSFYSWRLTFMTFHGKPR--ASVDVMKHV-----------------
--------------------------------------------
HESPLVMTMPLFILAFGALFAGMAFQSGFIGE-AYDA---FW--
KGALFTGAENHVLHEMHHVPTWVKLSPFVMMVAGFLLAYQFYIRA-----------------------------
------------------------------
PEMPKKLAERHDALYQFLLNKWYFDELYNFLFVKPSMKIGRFLWKR-GDGAVIDGLGPDGVA-
SRVQKLSAQIVRLQSGYIYHYAFAMLIGVTFFVTLSMFS----GGGAH-
>WP_068316307_NuoL_Pseudovibrio_hongkongensis
MY-------SAIVFLPLVGFLIAGLFG----RAIGAKASEYVTSSLLVVAALLSWVAFFDVAL--
GTEALVQIPVL-QWISSGALDVTWTLRIDTLTVVMLVVVNSVSALVHIYSIGYMHHDPH-------
RPRFFAYLSLFTFAMLSLVTSDNLLQMFFGWEGVGLASYLLIGFWFKKPSANAAAMKAFVVNRVGDFGFLLGIF
GVYMMFGTISY--
DDIFAQAGAIAAGTLEFLGYSLSFDSAITVTCLLLFLGAMGKSAQLLLHTWLPDAMEGPTPVSALIHAATMVTA
GVFMVARLSPLFELSHTALTVVTFVGASTALFAGTIGCVQNDIKRVIAYSTCSQLGYMFVALGIGAYGVAVFHL
FTHAFFKALLFLCAGSVIHAVSDEQDMRRMGGLRKHIPVTYWTMFIGTLALTGFGIPLVHIGHTYLGTAGFISK
DAIIEATYAGHNAASNFAFWMTIAAAFLTSFYSWRLTFMTFHGKPR--ASVDVMKHV-----------------
--------------------------------------------
HESPLVMTVPLFVLAVGALFAGMAFQSGFIGE-TYDT---FW--
KGALFTGADNHVLHDMHHVPIWVKLMPFIMMVAGFLLAVQFYIRK-----------------------------
------------------------------
PDAPKKLAERHDALYQFLLNKWYFDELYDFLFVKPSLKLGRFLWKR-GDGAVIDGMGPDGIA-
ARVQQITAQVVRLQSGYIYHYAFAMLIGVAFFITLSMFS----GGGAH-
>WP_207140330_NuoL_Labrenzia_aggregata
MY-------SAILFLPLIGFLIAGLFG----RSIGAKACEYVTSGLVILAALLSWITFFGFWL--
GGSELQVVELF-RWMDAGDLKVSWSIRVDTLTAVMLVVVNTVSALVHVYSIGYMHHDPH-------
RPRFFAYLSLFTFAMLSLVTADNLLQMFFGWEGVGLASYLLIGFWYQKPSANAAAMKAFVVNRVGDFGFALGIF
GVFYLFQSIDF--
TTIFNDAPSFIEEHLTFLGYHLDPHAAMTVVCLLLFMGAMGKSAQFLLHTWLPDAMEGPTPVSALIHAATMVTA
GVFMVARLSPLMELSHTALAVIVFFGATTAFFAATVGLVQNDIKRVIAYSTCSQLGYMFVALGVGAYSIGIFHL
FTHAFFKALLFLCAGSVIHAVSDEQDMRKMGGLRKHIPITYWTMMIGTLALTGVGIPFTHLG-----
FAGFISKDAIIEAAFAGAGGEHANPMALYGFWMTVISFYSWRLTFMTFHGKPR--APVDVMKHV----------
---------------------------------------------------
HESPLVMTVPLFILSVGAALAGMVFYGSFYGE-GYAE---FW--
KGALFNSETNHILHEMHHVPGWVKVSPFVMMVLGFVVAWVFYVRS-----------------------------
------------------------------
PEMPKQLAQRHSGLYQFLLNKWYFDELYDFIFVRPALWIGRVLWKK-GDGTVIDGYGPNGIA-
ARVQNVTAWVVRLQTGYLYHYAFAMLIGVAALITWSMFTGAGTGGGAH-
>WP_055121168_NuoL_Labrenzia_alba
MY-------SAILFLPLIGFLIAGLFG----RAIGDKACEYLTSGLVILAALLSWITFFGFWL--
GGSELQIEPLF-RWMDAGDLKVMWSIRVDTLTAVMLVVVNTVSALVHVYSIGYMHHDPH-------
RPRFFAYLSLFTFAMLSLVTADNLLQMFFGWEGVGLASYLLIGFWYKKPSANAAAIKAFVVNRVGDFGFALGIF
GIFFIFQSIEF--
TTIFNDAPSFIEGQMTFLGYHLDPQAAMTVICLLLFMGAMGKSAQFLLHTWLPDAMEGPTPVSALIHAATMVTA
GVFMVARLSPLMELSQTALTVIIFFGATTAFFAATIGLVQNDIKRVIAYSTCSQLGYMFVALGVGAYSIGIFHL
FTHAFFKALLFLCAGSVIHAVSDEQDMRKMGGLRKHIPITYWTMMIGTLALTGVGIPLTHLG-----
FAGFISKDAIIEAAYASAGGEHANPMALYGFWMTVISFYSWRLTFMTFHGKPR--APVDVMKHV----------
---------------------------------------------------
```

```
HESPLVMTIPLFVLSAGAALAGMIFYSSFYGE-GYEA---FW--
KGSLFTSEANHVLHEMHSVPAWVKVSPFVMMALGFLVAYQFYIRS-----------------------------
------------------------------
PETPKKLAERHQGLYQFLLNKWYFDEIYDFLIVRPTMWLGRVLWKK-GDGTVIDGYGPDGIS-
ARVQNVTSWVVRLQTGYLYHYAFAMLIGVAALITWSMFTGAGTGGGAH-
>MBD8891079_NuoL_Labrenzia_suaedae
MY-------SAIVFLPLIGFLIAGLFG----RALGAKACEFITSGLVVVAALLSWVAFFSIGY--
GETNLVQVEVL-RWITIGALDVSWSLRIDTLTVVMLVVVNSVSALVHIYSIGYMHHDPH-------
RPRFFAYLSLFTFAMLSLVTSDNLLQMFFGWEGVGLASYLLIGFWFQKPSANAAAMKAFVVNRVGDFGFALGIF
GVFVLFDSIDL--
STIFAKAPAVIGAALNFLGYHLDSHSAMTVVCLLLFMGAMGKSAQFLLHTWLPDAMEGPTPVSALIHAATMVTA
GVFMVARLSPLFELSHTALTVITFFGATTAFFAATVGLVQNDIKRVIAYSTCSQLGYMFVALGVGAYSVAVFHL
FTHAFFKALLFLCAGSVIHAVSDEQDMRKMGGLRKHIPVTYWTMFIGTVALTGVGIPFTHIG-----
FAGFISKDAVIEAAYAGHNAMAQYGFWLTVFAAGLTSFYSWRLTFMTFHGKPR--ASVDVMKHV----------
---------------------------------------------------
HESPLVMTAPLYILAVGATLAGMVFFGSFYGE-GYDA---FW--
KGALPSFEASHVLHAMHEVPGWVKVSPFVMMVGGFLVAYQFYIRS-----------------------------
------------------------------
PETPKKLAEDFSGLYQFLLNKWYFDELYDFLFVRPALWIGRQLWKK-GDGVVIDGYGPNGVA-
ARVQNVTTWVVKLQTGYLYHYAFAMLIGVAALITWSMFTGAGTVGGAH-
>WP_055455499_NuoL_Pannonibacter_indicus
MY-------SAIVFLPLIGFLIAGLFG----RVIGHTASEYITSGLVIVSAVLSWIAFFTIGY--
GETALVEVEVL-RWISSGALNVAWSLRIDTLTVVMLVVVNTVSALVHVYSIGYMHHDPH-------
RSRFFAYLSLFTFAMLSLVTSNNLLQMFFGWEGVGVASYLLIGFWYKKPSANAAAMKAFVVNRVGDFGFALGIF
GVFMMFGTINL--
PEIFANAQAVVGGHLNFLGHPLDAHTAMTVVCLLLFMGAMGKSAQFLLHTWLPDAMEGPTPVSALIHAATMVTA
GVFMVARLSPLFELSHTALTVVTFFGATTAFFAATVGLVQNDIKRVIAYSTCSQLGYMFVALGVGAYSIAVFHL
FTHAFFKALLFLCAGSVIHAVSDEQDMRKMGGLRKHIPFTYWTMFIGTLALTGVGIPLTHIG-----
LAGFLSKDLVIEAAYAASSGEHANAFALYGFWMTVVSFYSWRLTFMTFHGKPR--APVDVMKHV----------
---------------------------------------------------
HESPMVMLAPLFILSIGAVLAGMVFYSQFAGE-GYDA---FW--
KGALFTGAENHILHAIHEVPKWVKVSPFVMMVLGLATAWYFYIRS-----------------------------
------------------------------
PETPKRLAAEHEGLYKFLLNKWYFDELYNIIFVRPAMWTGRQLWKK-GDGKVIDGYGPDGVA-
ARVQNVTSWVVRLQTGYLYHYAFAMLIGVAALVTWSMFTGVLTGGGAH-
>WP_050473307_NuoL_Pannonibacter_phragmitetus
MY-------SAIVFLPLIGFLIAGLFG----RVIGHTASEYITSGLVIVSAVLSWIAFFTIGY--
GETALVEVEIL-RWISSGALDVAWSLRIDTLTVVMLVVVNTVSALVHVYSIGYMHHDPH-------
RSRFFAYLSLFTFAMLSLVTSNNLLQMFFGWEGVGVASYLLIGFWYKKPSANAAAMKAFVVNRVGDFGFALGIF
GVFMMFGTINL--
PEIFANAQAVVGGHLNFLGHPLDAHTAMTVVCLLLFMGAMGKSAQFLLHTWLPDAMEGPTPVSALIHAATMVTA
GVFMVARLSPLFELSHTALMVVTFFGATTAFFAATVGLVQNDIKRVIAYSTCSQLGYMFVALGVGAYSIAVFHL
FTHAFFKALLFLCAGSVIHAVSDEQDMRKMGGLRKHIPFTYWTMFIGTLALTGVGIPLTHIG-----
LAGFLSKDLVIEAAYAASSGAHANAFALYGFWMTLTSFYSWRLTFMTFHGKPR--APVDVMKHV----------
---------------------------------------------------
HESPMVMLAPLFILSIGAVLAGMLFYSQFAGE-GYDA---FW--
KGALFTGTENHILHAIHDVPKWVKVSPFVMMVLGLATAWYFYIRS-----------------------------
------------------------------
```

```
PETPKRLAAEHEGLYKFLLNKWYFDELYNLIFVRPAMWIGRQLWKK-GDGKIIDGYGPDGIA-
ARVQNVTSWVVRLQTGYLYHYAFAMLIGVAALVTWSMFTGVSTGGGAH-
>WP_149891971_NuoL_Roseibium_aestuarii
MY-------TAIVFLPLIGFLIAGLFG----RMIGPKASEIVTATLVTASAILSWVAFFTIGY--
GDTKLVQIEIM-RWISAGALDVTWGIRVDTLTVVMLVVVNSVSALVHWYSRGYMHHDPH-------
RPRFFAYLSLFTFAMLSLVTSDNLLQMFFGWEGVGLASYLLIGFWFHKPSANAAAMKAFVVNRVGDFGFALGIF
GTFVLFDSIDL--
QTIFNQAIALGGQGFEFLGYHLDTHAAMTVVCLLLFMGAMGKSAQFLLHTWLPDAMEGPTPVSALIHAATMVTA
GVFMVARLSPLFELSHTALTVVTFFGATTAFFAATVGLVQNDIKRVIAYSTCSQLGYMFVALGVGAYSIGVFHL
FTHAFFKALLFLCAGSVIHAVSDEQDMRKMGGLRKHIPWTYWTMFIGTVALTGVGIPFTHIG-----
FAGFISKDAVIEAAYAGHNAMSQYAFWLTVIAAGLTSFYSWRLTFMTFHGKPR--APVDVMSHV----------
---------------------------------------------------
HESPWIMLFPLVLLSVGAVLAGMVFYGVFVGE-GFEH---FW--
KGAIFMGPENHILHAMHEVPGWVKVSPFVMMAGGFVVAYIFYLRK-----------------------------
------------------------------
PDLPKKLAEQHSGLYQFLLNKWYFDEAYDFLFVRPAKWLGRTLWKK-GDGTVIDGFGPNGIA-
ARVQNVTTWVVKLQTGYLYHYAFAMLIGVAALVTWSMFTGAGTG-GAH-
>NBN77992_NuoL_Microvirga_tunisiensis
MY-------QAIVFLPLIGFLIAGLFG----RAIGHTASELVTTTLVVISAVLSWVAFFQIGY--
GETALVQVEIL-RWITSGALDVAWSLRIDTLTVVMLVVVNSVSALVHIYSIGYMHDDPH-------
RSRFFAYLSLFTFAMLSLVTADNLLQMFFGWEGVGLASYLLIGFWYQKSSANAAAMKAFVVNRVGDFGFALGIF
GVFVMFDTINL--
SAIFANAPAVVGAVLDFLGYSLDAGTAMTVVCLLLFMGAMGKSAQFLLHTWLPDAMEGPTPVSALIHAATMVTA
GVFMVARLSPLFELSHTALTVVTFFGATTALFAATVGLVQNDIKRVIAYSTCSQLGYMFVALGVGAYSVGVFHL
FTHAFFKALLFLCAGSVIHAVSGEQDMRRMGGLRKHIPVTYWTMMIGTLALTGVGIPLTYIG-----
FAGFISKDAVIEAAFAGHNAASQYAFWLTVAAAGLTSFYSWRLTFLTFHGKPR--APVDVMKHV----------
---------------------------------------------------
HESPMVMLVPLFILSVGAIAAGMAFKAPFFGD-GYDA---FW--
KGALFTGPDNHILHALHDVPKWVKVSPFVMMATGFIVAWYFYIRS-----------------------------
------------------------------
PETPKRLAAEHEGLYKFLLNKWYIDELYDWIFVRPAMWLGRTLWKK-GDGVVIDGCGPDGVA-
ARVQNVTTWVVRLQTGYLYHYAFAMLIGVAALVTWSMFSGVSTGGGAH-
>WP_106753365_NuoL_Pannonibacter_carbonis
MF-------QAIVFLPLIGFLIAGLFG----RAIGHKASELVTSTLVVIAALLSWVAFFQIGY--
GETALVQIEIL-RWITSGTLDVSWSLRIDTLTVVMLVVVNSVSALVHIYSIGYMHHDPH-------
RSRFFAYLSFFTFAMLSLVTADNLLQMFFGWEGVGVASYLLIGFWYQKPSANAAAMKAFVVNRVGDFGFILGIF
GVFVMFDTINL--
STIFANAPAVVGVLLNFLGYHLDAQSAITVVCLLLFVGAMGKSAQFLLHTWLPDAMEGPTPVSALIHAATMVTA
GVFMVARMSPLFELSTTALTVVTIFGATTAFFAATVGLVQNDIKRVIAYSTCSQLGYMFVALGVGAYSVGIFHL
FTHAFFKALLFLCAGSVIHAVSDEQDMRRMGGLRKHIPYTYWTMMIGTLALTGVGIPLTYIG-----
FAGFISKDAVIEAAFAGHNPAAQYAFWMTVVAAGLTSFYSWRLTFMTFHGKPR--APVDVMKHV----------
---------------------------------------------------
HESPMVMLVPLFLLSVGAIASGMVFKPQFIGD-AYDA---FW--
KGALFTGPDNHILHAIHDVPKWVKVSPFVMMAGGFIVAWYFYIRS-----------------------------
------------------------------
PETPKRLAAEQEGLYKFLLNKWYIDELYDWIFVRPAMWIGRMLWKK-GDGVVIDGCGPDGVA-
ARVQNVTTWVVRLQTGYLYHYAFAMLIGVAALVTWSMFTGVSTGGGAH-
>WP_189437262_NuoL_Pseudovibrio_japonicus
```

```
MF-------SAIVFLPLLGFLIVGMFG----NRLGVKASEYITSGFMIVAALLSWVVFFQYGL--
GSETTEIVTIF-RWITSGDLAVNWTIRVDTLTAVMLVVVNSVSCLVHIYSIGYMHHDPH-------
RPRFFAYLSLFTFAMLTLVTADDLLQMFFGWEGVGLASYLLIGFWYQKPSANAAAMKAFIVNRVGDFGFALGIF
GVFYLYGSIDF--
TSIFSSGPTLLEGTLNFLGFELMPEGALTVVCLLLFMGAMGKSAQFLLHTWLPDAMEGPTPVSALIHAATMVTA
GVFMVARLAPVFELAPTAMAVVTFFGATTAFFAATIGCVQNDIKRVIAYSTCSQLGYMFVALGVGAFGVGIFHL
FTHAFFKALLFLCAGSVIHAVSDEQDMRKMGGLRKHIPWTYWTMMIGTLALTGFGIPFIGIG-----
TAGFVSKDAIIEAAYVGHNAFASYAYWATVIAALLTSFYSWRLVFMTFHGKPR--ASVDVMKHI----------
---------------------------------------------------
HESPWVMLIPLLLLSIGALFAGMVFKEAFMGH-DYGH---FW-
KGVFGGTNSELAVIEEMHHVHWMVALSPTLAMIGGFIVAYMFYIAK----------------------------
-------------------------------
PDMPKVLAERHQPLYKFLLNKWYFDELYDFLFVKPAMALGRFLWKR-GDGTIIDGLGPDGVA-
SRVQQVTGWVVRLQSGYIYHYAFAMLIGVAFFVTLAMFSG----GGVH-
>WP_068001063_NuoL_Pseudovibrio_axinellae
MF-------SAIVFLPLLGFLIVGLFG----NRLGVKASEYITSGFMIVAALLSWVVFFQYGL--
GSESTQIVTIF-RWITSGDLAVNWTIRVDTLTAVMLVVVNSVSCLVHIYSIGYMHHDPH-------
RPRFFAYLSFFTFSMLALVTADDLLQMFFGWEGVGLASYLLIGFWYKKPSANAAAMKAFIVNRVGDFGFALGIF
GVFYLYGSIDF--
TTIFSSGPSLLENKLNFLGYELMPEAALTVVCLLLFMGAMGKSAQFLLHTWLPDAMEGPTPVSALIHAATMVTA
GVFMVARLAPIFELAPTALTVVTFFGATTAFFAATIGCVQNDIKRVIAYSTCSQLGYMFVALGVGAFGVGIFHL
FTHAFFKALLFLCAGSVIHSVSDEQDMRKMGGLRKHIPWTYWTMMIGTLALTGFGIPFTYIG-----
TAGFISKDAIIEAAYVGHNMFATYAFWATVLAAALTSFYSWRLVFMTFHGKTR--ASVDVMKHI----------
---------------------------------------------------
HESPWVMLVPLLLLALGAALAGMVFKEAFIGH-DYVH---FW-
KNVFGGTSSELAVIEEMHHVHWLVKLSPWLAMAGGFIIAYLFYIAK----------------------------
-------------------------------
PDMPKALAQRHKPLYLFLLNKWYFDELYDFLFVRPSMALGRFLWKR-GDGTVIDGLGPDGVA-
ARVQQVTGWVVRLQSGYIYHYAFAMLIGVAFFVTLAMFSG----GGVH-
>WP_093521060_NuoL_Pseudovibrio_ascidiaceicola
MF-------SAIVFLPLLGFLIVGMFG----NRLGVKASEYITSGFMIVAALLSWVVFFQYGL--
GSESTQIVTIF-RWITSGDLSVNWTIRVDTLTAVMLVVVNSVSCLVHIYSIGYMHHDPH-------
RPRFFAYLSFFTFSMLALVTSDDLLQMFFGWEGVGLASYLLIGFWYKKPSANAAAMKAFIVNRVGDFGFALGIF
GVFYLYGSIDF--
TTIFSSGPSLLENKLNFLGYELMPEAALTVVCLLLFMGAMGKSAQFLLHTWLPDAMEGPTPVSALIHAATMVTA
GVFMVARLAPIFELAPTALTVVTFFGATTAFFAATIGCVQNDIKRVIAYSTCSQLGYMFVALGVGAFGVGIFHL
FTHAFFKALLFLCAGSVIHSVSDEQDMRKMGGLRKHIPWTYWTMMIGTLALTGFGIPFTYIG-----
TAGFISKDAIIEAAYVGHNMFASYAFWATVLAAALTSFYSWRLVFMTFHGKTR--ASVDVMKHI----------
---------------------------------------------------
HESPWVMLVPLLLLALGAAFAGMVFKEAFIGH-DYLH---FW-
KNVFGGTNSELAVIEEMHHVHWLVKLSPWLAMAGGFIIAYLFYIAK----------------------------
-------------------------------
PDMPKALAERHKPLYLFLLNKWYFDELYDFLFVKPSLALGRFLWKR-GDGTVIDGLGPDGVA-
SRVQQVTGWVVRLQSGYIYHYAFAMLIGVAFFVTLAMFSG----GGVH-
>WP_00803949_Bartonella_tamiae
MF-------YAIVFLPLLGFLIAGLGG----KTLGVRGAELVTCTFMVIVAVLSWIAFFNIAI--
GNGSVVNVPVL-HWLSVGNLSFNWSLHIDTLTAVMLVVVNTVSALVHIYSIGYMHHDPS-------
RPRFFAYLSFFTFMMLMLVTANNLIQMFFGWEGVGLASYLLIGFWFQRPSANKAAIKAFVVNRVGDFGFLLGIF
```

```
SIFVLFKAVDF--
ETIFQKATDDSFVQMTFLSWHLDANIALTVTCLLLFVGAMGKSAQFLLHTWLPDAMEGPTPVSALIHAATMVTA
GVFMISRMSPIFELSTIALTVIVVIGATTAFFAATVGLVQNDIKRVIAYSTCSQLGYMFVALGVGAYGAAVFHL
FTHAFFKALLFLGAGSVIHAVSDEQDMRKMGGLRKYIPTTYWMMLIGTFALTGLGIPGTIFG-----
TAGFFSKDSIVESAFASYNPAAGYAFWLLVIAALFTSFYSWRLIFMTFHGKPR--ATADVMHHV----------
---------------------------------------------------
HESSAVMLIPLFVLAIGALFAGVIFQPYFFGD-LYIG---FW--
KGALFTGPENHIMHEAHNVPMWVKTSPFIAMVIGFIFAYLFYILA-----------------------------
------------------------------
PSIPKALANAFPFGYRFLLKKWYFDELYNVLFVRPTFALGRFFWKV-GDIAIIDGLGPNGIA-
ARVVNITNRVVRMQTGYLYHYAFAMLIGVAALITWMMIGSIN-------
>WP_108004655_NuoL_Mycoplana_dimorpha
MDTII----KAIVFLPLIGFLIAGLGG----NTIGAKASEYVTSGFLVIACALSWLVFFQVAL--
GEEELIKVPVL-HWIQSGSFDVEWAFRVDTLTAVMFVVVNTVSMLVHIYSIGYMHHDPH-------
RPRFFAYLSLFTFAMLMLITADNLLQLFFGWEGVGLASYLLIGFWYKKASANAAAMKAFIVNRVGDFGFLLGIA
AVFVLFGSINF--
ETIFATAATYLPAEITLFGMPLDKADALTGACLLLFMGAMGKSAQFLLHTWLPDAMEGPTPVSALIHAATMVTA
GVFLVARMSPLYELSPDALTVVTIVGAITAFFAATVGLVQNDIKRVIAYSTCSQLGYMFVALGIGAYGAAIFHL
FTHAFFKALLFLGAGSVIHAVDGEQDMRYMGGLRTHIPVTFWMMTIGTLALTGVGIPGTMIG-----
FAGFFSKDAIIESAFASHSAVSNFAFILLVVAALFTSFYSWRLAFMTFFGRPR--ASSDVMHHV----------
---------------------------------------------------
HESPAVMLVPLYILAVGAIFSGFIAVEYFYGH-HYEE---FW--
QGAIFTGVENKLIEEFHHVPLWVKWSPFVAMALGLFTAWYMYIRR-----------------------------
------------------------------
PDLPKYLAEQHRGLYQFLLNKWYFDELYDFLFVRPALWLGRLFWKG-GDGAVIDRLGPNGLA-
ARVLDVTDRVVRLQTGYLYHYAFAMLLGIAALVTWMMLGSSI-------
>WP_18348086_NuoL_Martelella_radicis
MIY------KLIIFLPLLGFLIAGLFG----NRIGAKATEWVTIAFMAVAAILSWYVFISVGF--
LSHAEDLVIQVMQWISSGDIDVAWSLRIDTLTAVMLVVVNTVSCLVHLYSVGYMHDDPH-------
RPRFFAYLSLFTFAMLSLVTSDNLLQMFFGWEGVGLASYLLIGFWYKKPSACAAAMKAFIVNRVGDFGFLLGIA
GLFFLAHTINF--
DPIFAAAERIADNHLHFLGMDLTLMHAITAICLLLFMGAMGKSAQFLLHTWLPDAMEGPTPVSALIHAATMVTA
GVFLVARMSPLFELSPSALTFVTLIGSITALFAATIGLVQNDIKRVIAYSTCSQLGYMFAALGAGAYGAAVFHL
FTHAFFKALLFLCAGSVIHAVSGDQDMRRMGGLRKHIPITFWGMIFGTLAITGVGIPGTVIG-----
AAGFFSKDAIIEAAFVSASPLANFSFWALVIAAVLTAFYSWRLIFMTFFGKPR--ASSDVMHHV----------
---------------------------------------------------
HESPWVMLIPLIVLSVGALFAGVVFAEFFYGH-HYAE---FW--
KGALFTGPHNEVLEEHHHVALWVKLSPFFAMLIGTVVAIYMYLIK-----------------------------
------------------------------
PASAPAMARAFPRIYAFLLNKWYFDELYDFLFVRPAKKLGHFLWQK-GDVGVIDRFGPNGIA-
ALVVDVTNRVVKLQSGYVYHYAFAMLIGVAALVTWMMLGGAL-------
>WP_045681980_NuoL_Martelella_endophytica
MIY------KLIIFLPLIGFLIAGLFG----NRIGAKATEWLTITLMAIAAVLSWYVFISVGF--
LQHEEGYVLHVMQWINSGGINATWSLRIDTLTAVMLVVVNTVSCLVHLYSVGYMHHDPH-------
RPRFFAYLSFFTFAMLSLVTADNLLQMFFGWEGVGLASYLLIGFWYKKPSACAAAMKAFIVNRVGDFGFLLGIA
GLFFLAHTIEF--
DPIFEAARNLADNHLHFLGMNLDVMQATTAICLLLFMGAMGKSAQFLLHTWLPDAMEGPTPVSALIHAATMVTA
GVFLVARMSPVFELSPTALTFITLVGSITAFFAATVGLVQNDIKRVIAYSTCSQLGYMFAALGAGAFGAGVFHL
```

FTHAFFKALLFLCAGSVIHAVSDEQDMRRMGGLRKHIPVTFWAMVIGTLAITGVGIPFTEIG-----
AAGFFSKDAIIESAYVSASPLASFSFWMLVIAAAFTAFYSWRLIFMTFFGKPR--ASSDVMHHV----------
---------------------------------------------------
HESPWIMLTPLVILSVGALFAGVVFLPYFFGH-DYSE---FW--
KGALFTGPHNEVLDEHHHVPAWVKLSPFIAMLLGTVTAIYMYLIN-----------------------------
------------------------------
PSSAPKLAQTFPRLYAFLLNKWYFDELYDFLFVRPSKRLGYFLWQK-GDVGFIDRYGPNGIA-
SLVSDITDRVVKLQSGYVYHYAFAMLIGLAALVTWMMLGGAL-------
>WP_138748498_NuoL_Martelella_lutilitoris
MIY------KLIIFLPLLGFLIAGLFG----NRIGAKATEWVTIAFMVVAAILSWYVFISVGF--
LSHAEDLVIQVMQWISSGDIDVAWSLRIDTLTAVMLVVVNTVSCLVHLYSVGYMHDDPH-------
RPRFFAYLSLFTFAMLSLVTSDNLLQMFFGWEGVGLASYLLIGFWYKKPSACAAAMKAFIVNRVGDFGFLLGIA
GLFFLAHTINF--
DPIFAAAERIADDHLHFLGMDLTLMHAITAICLLLFMGAMGKSAQFLLHTWLPDAMEGPTPVSALIHAATMVTA
GVFLVARMSPLFELSPSALTFVTLIGSITALFAATIGLVQNDIKRVIAYSTCSQLGYMFAALGAGAYGAAVFHL
FTHAFFKALLFLCAGSVIHAVSGDQDMRRMGGLRKHIPITFWGMIFGTLAITGVGIPGTVIG-----
AAGFFSKDAIIEAAFVSASPLAGFSFWALVIAAVLTAFYSWRLIFMTFFGKPR--ASSDVMHHV----------
---------------------------------------------------
HESPWVMLIPLIVLSVGALFAGVVFAEFFYGH-DYSE---FW--
KGALFTGPHNEVLEEHHHVALWVKLSPFFAMLIGTAVAIYMYLIK-----------------------------
------------------------------
PSSAKATAKAFPRIYAFLLNKWYFDELYDFLFVRPAKKLGHFLWQK-GDVGVIDRFGPNGIA-
ALVVDVTNRVVKLQSGYVYHYAFAMLIGLAALVTWMMLGGAL-------
>WP_075697151_NuoL_Pseudovibrio_brasiliensis
MF-------SAIVFLPLLGFLIVGLFG----NRLGVKASEYITSGFMIVAALLSWVVFFQYGL--
GSESTQLITIF-RWITSGDLSVNWTIRVDTLTAVMLVVVNSVSCLVHIYSIGYMHHDPH-------
RPRFFAYLSLFTFAMLTLVTADDLLQMFFGWEGVGLASYLLIGFWYQKPSANAAAMKAFIVNRVGDFGFALGIF
GVFYLYGSIDF--
TSIFASGPSLLENKLNFLGYELMAEPALTVVCLLLFMGAMGKSAQFLLHTWLPDAMEGPTPVSALIHAATMVTA
GVFMVARLAPIFELAPTAMTVVTFFGATTAFFAATIGCVQNDIKRVIAYSTCSQLGYMFVALGVGAFGVGIFHL
FTHAFFKALLFLCAGSVIHAVSDEQDMRKMGGLRKHIPWTYWTMMIGTLALTGFGIPG-LAG-----
TAGFVSKDAIIEAAYVGHNAFASYAYWATVLAALLTSFYSWRLVFMTFHGKPR--ASVDVMKHI----------
---------------------------------------------------
HESPWVMLVPLILLSIGALFAGMAFKEAFMGH-DYAH---FW-
KDVFGGTNSELAVIEEMHHVHWLVAWSPTFAMITGFIIAYVFYIAK----------------------------
-------------------------------
PDMPKALAERHQPLYKFLLNKWYFDELYDFLFVKPAMALGRFLWKR-GDGTVIDGLGPDGIA-
SRVQQVTGWVVRLQSGYIYHYAFAMLIGVAFFVTLAMFSGGGVH-----
>WP_046478826_NuoL_Ca._Filomicrobium_marinum
MI-------LAIVFLPIVGALIAGLFG----RFIGARASEFITTGLLIIAAVLSWVAFVRVGF--
DAETARVPVM--RWITSGELDIAWSLRIDTLTAVMLVVVTTVSALVHVYSIGYMHEDPS-------
RSRFFAYLSLFTFAMLMLVTSDNLLQMFFGWEGVGLASYLLIGFWYTKPEANAAAIKAFVVNRVGDFGFMLGMF
GLFFVFNTLNF--DPLFEATAQAAGKS---
FNFLGYEVDILTTLCLLLFMGAMGKSAQFLLHTWLPDAMEGPTPVSALIHAATMVTAGVFMVARLSPIFEYAPT
ALEVVTLVGTVTAFFAATVGLVQNDIKRVIAYSTCSQLGYMFAALGVGAYSVGIFHLFTHAFFKALLFLGAGSV
IHAMHHEQDMRNMGGLAGKIPITWAMMLIGTLALTGVGIPH-VAG-----
FAGFHSKDAIIEATFAAHTPGNFVFWVLV-FTAFLTAFYSWRLVFMTFHGKTR--AAPDVFKHA----------
---------------------------------------------------

```
HESPWVMLVPLFVLAFGAVVAGFVFAPYFIGE-SYAE---FW--
GKALFTAPSNHILHDMHHVEEWVYWAPFLAMASGFALSWIFYVQA-----------------------------
------------------------------
PWLPEATARTFRPIYLFLLNKWYFDELYNWLFVRSAKGLGRFLWKA-GDGAVIDGTI-NGTA-
DSVQWTTGRVVKLQSGYIYHYAFAMLIGVALILTYFLRGLVMTGGAGQ-
>WP_111433419_NuoL_Rhodobium_orientis
MY-------SAIVFLPLLGFLIAGLFG----RIIGDKASEYITSGLLVVSCLLSWIAFFSVGY--
GDGPVEQVPIA-TWITSGALSIDWSLRIDTLTVVMLVVVTTISALVHIYSIGYMHHDPS-------
RARFFAYLSLFTFAMLMLVTADNLLQMFFGWEGVGLASYLLIGFWYHKPSANAAAMKAFIVNRVGDFGFSLGIF
ALFALTGSIGL--DAVFAAAGDLQETT---
FLFLSWDVNALTVICLLLFMGAMGKSAQFLLHTWLPDAMEGPTPVSALIHAATMVTAGVFMVARLSPLFELSHT
ALTVVVIIGATTAFFAATVGLVQNDIKRVIAYSTCSQLGYMFVALGVGAYGAGIFHLFTHAFFKALLFLSAGSV
IHAVSDEQDMRKMGGLRNHIPVTFWMMIIGTLALTG--FPL---------
TAGFFSKDAIIESSFVGHGAAAEYAFWMVVIAALFTSFYSWRLIFMTFFGKPR--ASAEVMGHV----------
---------------------------------------------------
HESPRVMTVPLMILALGALAAGVLFAGSFIGE-GYGE---FW--
KTALFVSEENHILHEMHNVPLWVKFAPFVMMVAGFVIAFYFYIMS-----------------------------
------------------------------
PETPKKLAKQHDGLYQFLLNKWYFDELYDVLFVRSAKWLGRFLWKQ-GDGRVIDGYGPDGVS-
ARVLQATGWVVRLQTGYLYHYAFAILIGVAALITWAMFTGGGAN-----
>WP_018698215_NuoL_Amorphus_coralli
MY-------AAIVFLPLIGFLVAGIFG----RLIGARASEIVTTSLLFVSAILSWVAFIHVGF--
SDVEPIKVPLM-PWITSGALEIDWSLRVDTLTVVMLIVVTSVSSLVHLYSIGYMSHDPD-------
RPRFFAYLSLFTFAMLMLVTADNLVQMFFGWEGVGLASYLLIGFWYTRPSAVAASIKAFVVNRVGDFGFMLGIF
ALFMLVGSVDL--DTIFGHGEELSTTT---
FSFLGYQVNAVETICFLLFIGAMGKSAQLGLHTWLPDAMEGPTPVSALIHAATMVTAGVFMVARMSPLFEYAPA
ALTFVTFIGATTAFFAATIGCTQNDIKRVIAYSTCSQLGYMFAALGVGAYSAGIFHLMTHGFFKALLFLCAGSV
IHAMHDEQDMRKMGGLVKKIPLTYAGMVIGTLAITG--FPY---------
TSGYFSKDAIIEATFAAHNPFSAYAFWLTTLAALLTAFYSWRLIFMTFHGKTR--ASADVFKHA----------
---------------------------------------------------
HESPPVMLIPLAVLSCGALFAGFILHGYFIGH-EYEH---FW--
RGSLYLGAENHILHDMHHLPAWVPLAPTVMMITGVAWAYWWYMLS-----------------------------
------------------------------
PKTPAAVAREHPALYQFLLNKWYFDELYDFLFVRPTKRIGHFLWKK-GDGWLIDGFGPDGVS-
HRVLDVTQRVVRLQTGYIYHYAFAMLIGVAALVTWAMMTGVGAP-----
>WP_132035152_NuoL_Aquabacter_spiritensis
MY-------HAIVFLPLIGFLIAGLFG----RSIGARASEVVTTSLLFIACLLAWIAFFNVGF--
GGHDTRVQVM--HWMYVGDLKIDWAFRIDTLTAVMLIVVTTVSSLVHLYSMGYMHEDPS-------
RPRFFAYLSLFTFAMLMLVTADNLLQMFFGWEGVGLASYLLIGFWYEKPSANAAAMKAFVVNRIGDFGFILGIF
GVFVMTGAVNF--ETVFAQAPGLAGKT---
IAFLGHHWDAPTVIAVLLFIGAMGKSAQFLLHTWLPDAMEGPTPVSALIHAATMVTAGVFMVARMSPIFELSPS
ALDLVMIVGATTSMFAATIALVQTDIKKVIAYSTCSQLGYMFVAMGAGAYSIGMFHLLTHACFKALLFLSAGSV
INAMHHEQDMRNMGGLYKKIPFTYAMMLVGTLAITG--FPF---------
LAGYYSKDAIIESAWGSTNSFALYGWVMTVGAAALTSFYSWRLVFLTFHGQPHD-HHHYEAA------------
---------------------------------------------------
HESPLVMTIPLGVLGFGSVVSGMILYPEFVGH-DVGE---FF--
RQSIFMGAENHVLHAMHDIPFWAKWMPTLMMALGLLVAYWFYVVD-----------------------------
------------------------------
```

```
RTLPVRLAHSQDALYKFLLNKWYFDEIYDFLFVRPALFIGRVLWKQ-GDVRIIDGMGPDGVS-
ARVLDVTGRVVRLQSGYLYHYAFVMLIGVAALITWFMFAGV--------
>WP_209483414_NuoL_Xanthobacter_flavus
MY-------QAIVFLPLLGFLIAGLFG----KKIGDRASEVVTTSFLFVACLLAWIAFFAVGF--
GDHDTRVQVM--KWIYVGDLKVDWAFRIDTMTVIMLIVVTTVSSLVHLYSIGYMHEDPS-------
RPRFFAYLSLFTFAMLMLVTADNLLQLFFGWEGVGLASYLLIGFWYEKPSANAAAMKAFVVNRVGDFGFMLGIF
AIFVMTGSIAF--EEVFAAAPGLAGKT---
ISFLGHNWDAPTVIALLLFVGACGKSAQFLLHTWLPDAMEGPTPVSALIHAATMVTAGVFMVARMSPIFELSQT
ALNVVMLVGATTAFFAATVGLVQNDIKKVIAYSTCSQLGYMFVAMGAGAYSIGMFHLLTHAFFKALLFLGAGSV
IHAMHHEQDMRHMGGLRKKIPFTYWTMVIGTLAITG--FPF---------
LAGYYSKDAIIEAAYASHNYFHVYGYWMTVIAAALTAFYSWRLIFLTFWGHPHD-HHHYEAA------------
---------------------------------------------------
HESPLVMTIPLGVLSLGALFAGIAFKNIFVGE-GVEH---FF--
RHSIFMGAENHILHAMHDIPGWAAFAPTVMMVVGSLFAIWFYIVN-----------------------------
------------------------------
PKVPAQLAKSQDALYQFLLNKWYFDEIYNFLFVRPTLWLGRTLWKR-GDGTIIDGLGPDGVS-
ARVLDITRGVVRLQTGYLYHYAFVMLIGVAALITWFMFAGV--------
>WP_213363409_NuoL_Ancylobacter_defluvii
MY-------QAIVFLPLAGFLIAGLFG----RAIGARASELITTGLLFISALFAWIVFFQTAF--
GDGGGTVEIF--TWIASGALDVKWALRVDTLTAVMLVLVTTVSSLVHLYSMGYMHEDPS-------
RPRFFAYLSLFTFAMLMLVTSDNLVQLFFGWEGVGLASYLLIGFWYDKPSACAAAMKAFIVNRVGDFGFALGIF
AVFMMTGSVGF--EQVFAAAPSLMDKT---
IHFIGIDWHAPTIICLLLFIGAMGKSAQLGLHTWLPDAMEGPTPVSALIHAATMVTAGVFMVARLSPLFELSPV
ALNVVMFVGASTAMFAATVGCVQNDIKRVIAYSTCSQLGYMFTALGAGAYSVGVFHLFTHGFFKALLFLAAGSV
IHAMHHEQDMRHMGGLWRKIPFTYATMLIGGLALTG--FPF---------
TAGYFSKDAIIETAFMSHSPFSTYAFVMTVAAAALTSFYTWRQMFMTFHGHPHD-HHHYEHA------------
---------------------------------------------------
HESPLTMLIPLGVLSLGALFAGMILYPYFVGH-DVEH---FF--
REAIFMGPDNKILEEIHHIPAWAKWSPTVMMVLGFVVAYWMYVAN-----------------------------
------------------------------
PAVPKALARRNDVLYRFLLNKWYFDELYDFLFVRPTMWLGRFLWKQ-GDGRVIDGWGPNGIS-
ARVIDVTNRVVRLQTGYLYHYAFVMLVGVAALITWFMFGGAH-------
>WP_086090017_NuoL_Pseudorhodoplanes_sinuspersici
MY-------QAIVFLPLIGCLIAGLFG----RIIGARPAELVTTLLLFASAALSWYAFARVGF--
GHIDERIVLF--PWIQSGDFKVDWVIRVDTLTAVMLVVVTTVSSLVHLYSIGYMHEDPH-------
RQRFFAYLSLFTFAMLALVTADNLAQLFFGWEGVGLASYLLIGFWYQKPEANAAAIKAFVVNRVGDFGFALGIF
ALFMMTGAIDF--DTIFAQAPALTGKT---
IDFFGWHADALTLICLMLFMGAMGKSAQFLLHTWLPDAMEGPTPVSALIHAATMVTAGVFMVARLSPLFELAPV
ALMVVTLIGATTAMFAATVGLVQNDIKRIIAYSTCSQLGYMFVAMGVGAWSVGMYHLFTHAFFKALLFLGSGSV
IHAMHHEQDIRHMGGLKNKIPLTYWLMIIGTLALTG--FPF---------
TAGYFSKDAIIEAAYAGKNPFAVYGFACTVLAAALTSFYSWRLIFKTFHGEPHD-RHHYEEA------------
---------------------------------------------------
HESPMVMLVPLMVLAAGSILAGYPFYEYFAGH-HVEE---FF--
RNSLAFSKTNTILEDMHHVPALVVWTPTVMMVLGFAIAYLFYIRD-----------------------------
------------------------------
PRIPVEMARNHQLLYRFLLNKWYFDELYDLIFVRPAKWLGTTLWKR-GDGWLIDGFGPDGVS-
ARVLDVTRNVVRLQTGYLYHYAFAMLIGAAALITWFMFAGAH-------
>WP_109919883_NuoL_Zavarzinia_compransoris
```

```
386 MI-------AAIVFLPLLGAVIAGFFG----RLIGARGAQIVTSTLLVVAAVLSWVTFFQVT---
GHEGGAYKMRLMTWVLSGAFEVDWAFRVDTLTAVMLVVVTSVSSLVHIYSVGYMAEDPH-------
IPRFHAYLSLFTFAMLMLVTADNFLQVFFGWEGVGLASYLLIGFWYKKPSANAAAIKAFVVNRVGDFGFALGIF
AIFLVFNSVSF--DTVFAAAAGKAGQT---
FNFLGYDVDIMTTICLLLFVGCMGKSAQLGLHTWLPDAMEGPTPVSALIHAATMVTAGVFLVARASPLFEFAPS
ALAVVTVVGATTAFFAATVGLVQNDIKRVIAYSTCSQLGYMFFAAGVGAYEAAVFHLFTHAFFKALLFLGAGSV
IHAMHHEQDMRNMGGTWKYIKFTYAMMWIGNLALA--GVPF---------
FAGFYSKDMVLEVAYAAHSGVGTYAFWLGIIAASFTAFYSWRLLFLTFHGEKRW-GGHGH-GHGHDDHGHG---
--------------------HGHAHTP------------------------
HESPLAMTIPLALLAVGAVFSGFWFYDSFVGE-HREA---FW--
GGAIFVAEANKVIEHAHHVPGWVKLLPLIVTAGGIFVAWVFYILK-----------------------------
------------------------------
PKWPDQLAKTHREAYAFLLNKWYFDELYDLLFVKGARLFGRVFWKQ-GDGAVIDGLGPNGIS-
ARVLDIARRAAKLQSGFVYHYAFAMLIGVALLVTWYVYRMGVTP-----
387 >WP_119778097_NuoL_Oleomonas_sp._K1W22B-8
388 MI-------AAIVFLPLLGAIIVGFFG----RRLGNRGAQVITSGFLVVAAVLSWITFFQAT---
GHATDEPLKVHLTWMLSGSLEVEWAFRIDALTAVMLVVVNTVSSLVHIYSAGYMEEDPH-------
IPRFQAYLSLFTFAMLMLVTADNFVQVFFGWEGVGLASYLLIGFWYKKPSANAAAIKAFVVNRVGDFGFALGIF
AIFFVFNSVSF--DTVFAAAAGETGKT---
FNFLGYDVDILTTICLLLFVGCMGKSAQLGLHTWLPDAMEGPTPVSALIHAATMVTAGVFLVARCSPLFEHAPV
ALEVVTIVGASTAFFAATVGLVQNDIKRVIAYSTCSQLGYMFFAAGVGAYEASIFHLFTHAFFKALLFLGAGSV
IHAMHHEQDMRNMGGLWRHIKLTYALMWIGNLALA--GIPL---------
FAGYYSKDTILEVAFAAHTGVGTYAYILGTLAAMMTAFYSWRLLFMTFHGEARW--GHGH-GHG-DHGHGD---
--------------------HGHAHTP------------------------
HESPLVMTIPLIVLALGAAAAGIIFFDDFVGH-HREA---FW--
GGAIFVAETNKVLEEAHHIPEWAALLPLVLSVFGIFLAWVFYIAR-----------------------------
------------------------------
PKWPAALAQTHKELYAFLLNKWYFDELYDLLFVRPAKAIGRLLWKG-GDGRIIDGLGPDGIS-
ARVLDIARRAVKLQSGFVYHYAFAMLIGVAALVSWYVYRIGVTP-----
389 >WP_091859036_NuoL_Bosea_robiniae
390 MY-------HAIVFLPLIGFLIAGLFG----RLIGARGSEIVTTSLLMVSAALSWVALTQVGF--
GSGTTRVQVA--SWISSGELQVDWAFRVDTLTAVMLVVVNTVSSLVHLYSIGYMHEDEH-------
RSRFFAYLSLFTFAMLMLVTADNLVQMFFGWEGVGLASYLLIGFWYQKPSANAAAMKAFIVNRVGDFGFLLGIF
LIFVLFGAVTF--DAIFPRAGEMVSQS---
FRFLGYDWNALTLTCLLLFMGAMGKSAQFLLHTWLPDAMEGPTPVSALIHAATMVTAGVFMVARLSPVFEYAPV
ALTVVVVIGATTAFFAATVGLVQNDIKRVIAYSTCSQLGYMFVALGVGNYGAGIFHLFTHAFFKALLFLGAGSV
IHAMHHEQDMRHMGGLRKHIPLTAAAMTIGTLALTG--FPG---------
FAGYFSKDAVIESAYASVAHGGLASSYAFVLAAGMTSFYSWRLYFMTFEGAPRWGHGHDAHGHGHDDHAHAAD-
--------------GHDDHGHGHDHTP------------------------
HESPWTMLVPLAVLSLGAIVAGYAFKEAFIGH-DFEH---FW--
KSALFMGKDNHILHAMHEVPKWVVWSPFVAMVIGFALAWYMYVRR-----------------------------
------------------------------
PEIPGKLAAANPALYQFLLNKWYFDELYDFLFVRPAKWLGRFLWKK-GDGLVIDGFGPDGVS-
ARVVDVTNRVVRLQTGYLYHYAFAMLIGVAGLVTWYLVARG--------
391 >WP_079591299_NuoL_Bosea_thiooxidans
392 MY-------HAIVFLPLIGFLIAGLLG----RLIGARGSEIVTTSLLFVSAVLSWIALAQVGF--
GSGTTRIQVA--NWISSGQLQADWAFRIDTLTAVMLVVVNTVSSLVHLYSIGYMHEDPH-------
RPRFFAYLSLFTFAMLMLVTADNLVQMFFGWEGVGLASYLLIGFWYQKPSANAAAIKAFVVNRVGDFGFLLGIF
```

```
LIFVLFGAVTF--DAIFPRAGEFVNTS---
FRFLGYDWNALTLTCLLLFMGAMGKSAQFLLHTWLPDAMEGPTPVSALIHAATMVTAGVFMVARLSPVFEYAPV
ALTVVVVVGATTAFFAATVGLVQNDIKRVIAYSTCSQLGYMFVALGVGNYGAGIFHLFTHAFFKALLFLGAGSV
IHAMHHEQDMRHMGGLRKHIPLTAAAMTIGTLALTGVGIPGTI-----
FGFAGFFSKDAIIESAYASVATGGFASSVLLVVAACMTSFYSWRLYFMTFEGTPRWGHGHAHQGHDDHAHAAHA
H--DDH------------GHDDHGHDHTP------------------------
HESPWTMLLPLVLLSIGAIAAGFAFKEAFIGH-DFEH---FW--
KSALFMGKDNHILHAMHEVPKWVILSPFVAMVIGFLLAYWMYVRR-----------------------------
------------------------------
PDIPGRLAAANPALYRFLLNKWYFDELYDFLFVRPAKWLGRFLWKK-GDGLIIDGFGPDGVS-
ARVVDVTNRVVRLQTGYLYHYAFAMLIGVAGLVTWYLVTRG--------
>WP_066611767_NuoL_Bosea_sp._PAMC26642
MY-------HAIVFLPLIGFLIAGLFG----RLIGARGSEIVTTSLLFVSAVLSWVAFFSVGF--
GSGTTRIQIA--TWMASGDLRVDWAFRIDTLTAVMLVVVNTVSCLVHLYSIGYMHEDEH-------
RPRFFAYLSLFTFAMLMLITSDNLVQMFFGWEGVGLASYLLIGFWYKKPSANAAAMKAFIVNRVGDFGFALGIF
LVFVLFGSVGF--DAIFPRVADLTNAT---
FHFLGRDWHALTLASLLLFMGAMGKSAQFLLHTWLPDAMEGPTPVSALIHAATMVTAGVFMVARLSPIFEYAPA
ALTVVIVIGATTAFFAATVGLVQNDIKRVIAYSTCSQLGYMFVALGVGAYSAGIFHLFTHAFFKALLFLGAGSV
IHAMHHEQDMRNMGGLRKHIPLTAAAMTIGTLALTG--FPL---------
FAGYFSKDAIIESAYASVAHTGFPASYAFVVAACMTSFYSWRLYFMTFEGKPRWGADA-
HAGHGHDAHDDHAHAAHAHDDNA-------HGDHGHGHAHTP------------------------
HESPLVMLVPLAVLSLGAVAAGFAFKEAFIGH-NYEH---FW--
KAALFTGKDNHILHEMHEVPGWVVASPFIAMVIGFLLSLYMYVLR-----------------------------
------------------------------
PDVPGKLAAANPALYKFLLNKWYFDEIYDFLFVKPAMWLGKFLWKK-GDGFVIDGMGPDGIS-
ARVVDVTNRVVRLQTGYLYHYAFTMLIGVAGLVTWYLLARG--------
>WP_103871693_NuoL_Bosea_lathyri
MY-------HAIVFLPLIGFLIAGLFG----RLIGPRGSEILTTSLLVVSAVLSWVALFNVGF--
GSGTTRIQVA--TWLASGDLRVDWAFRIDTLTVVMLVVVNTVSSLVHVYSIGYMAEDPH-------
RPRFFAYLSLFTFAMLMLVTADNLVQMFFGWEGVGLASYLLIGFWYQKPSANAAAMKAFIVNRVGDFGFLLGIF
LVFVLFGTVGF--ESIFPRVGELTTQT---
FRFLGYEWNALTLTCLLLFMGAMGKSAQFLLHTWLPDAMEGPTPVSALIHAATMVTAGVFMVARLSPIFEYAPV
ALTVVVVVGATTAFFAATVGLVQNDIKRVIAYSTCSQLGYMFVALGVGAYAPAIFHLFTHAFFKALLFLGAGSV
IHAMHHEQDMRHMGGLRKHIPLTAAAMTIGTLALTG--FPF---------
FAGYFSKDAIIESAYAAVAHTGFAASYLLVVAACFTSFYSWRLYFMTFEGRPRWA-
GHGHEAHGHDDHAHAAHGHDDH---------------GHGHAHTP------------------------
HESPLSMLIPLALLSLGAVAAGFAFKEAFIGH-DYEH---FW--
KGALFTGKDNHILHAMHEVPGWVVASPFIAMVLGFVFALYFYVLR-----------------------------
------------------------------
PDIPGKLAAANPVLYRFLLNKWYFDEIYDFLFVKPSMWLGRFLWKK-GDGLVIDGMGPDGIS-
ARVVDVTNRVVRLQTGYVYHYAFAMLIGVAGFVTWYLLARG--------
>WP_114829996_NuoL_Bosea_caraganae
MY-------HLIVFLPLIGFLIAGLLG----RLIGPRGSEIVTTSLLMVSAALSWVAFFQVGF--
GSGTTRIQIA--TWLASADLRVDWAFRIDTLTVVMLVVVNTVSSLVHLYSIGYMSEDPH-------
RPRFFAYLSLFTFAMLMLVTADNLVQMFFGWEGVGLASYLLIGFWYQKPSANAAAMKAFIVNRVGDFGFALGIF
LVFVLFGTVAF--DGIFPKAGELVTQT---
FRFVGYEWNALTLTCLLLFMGAMGKSAQFLLHTWLPDAMEGPTPVSALIHAATMVTAGVFMVARLSPIFEYAPV
ALTVVVVIGATTAFFAATVGLVQNDIKRVIAYSTCSQLGYMFVAMGVGAYSAGIFHLFTHAFFKALLFLGAGSV
```

```
IHAMHHEQDMRKMGGLRKYIPLTAAAMTIGNLALTG--FPF---------
FAGYFSKDAIIESAYASVAHGGFASSVLLVVACFTSFYSWRLYFMTFEGQPRWA-
GQGHDAHGHDAHGHDDHAAHGH------------DDHGHGHDHTP------------------------
HESPWVMMIPLAVLALGAVAAGFVFKEAFIGH-DYEH---FW--
KAALFTGKDNHILHAMHEVPGWVVWSPFVAMVLGFLVALYMYVLR-----------------------------
------------------------------
PDVPGKLAAANPALYQFLLNKWYFDEIYDFLFVKPAMWLGKFLWKK-GDGFVIDGMGPDGIS-
ARVVDVTNRVVRLQTGFVYHYAFAMLIGVAGLVTWYLLARG--------
>WP_114771645_Microvirga_subterranea
MY-------HAIVFLPLVGFLIAGLFG----RILGARPSELITTGLLMVAAVLSWLAFFSVAY--
GDGATRIQVA--QWMVSGDLVVDWAFRIDTLTAVMLIVVNTVSALVHLYSIGYMHEDPH-------
RPRFFAYLSLFTFAMLMLVTADNLVQMFFGWEGVGLASYLLIGFWYQKDSANAAAMKAFIVNRVGDFGFLLGIF
TLFVLFNSVTF--DQIFPRVAELANNR---
FHFLGIEWHALTIACLLLFMGAMGKSAQFLLHTWLPDAMEGPTPVSALIHAATMVTAGVFMVARLSPVFEYAPA
ALTVVTIIGGITAFFAATVGLVQNDIKRVIAYSTCSQLGYMFVALGVGAYSVGVFHLFTHAFFKALLFLGAGSV
IHAMHHEQDMRHMGALRRYIPFTTAMMAIGTLALTG--FPF---------
TAGYYSKDAVIEAAYASHSTAGSFAFLATVIAAFMTSFYSWRLFFMTFEGSARWG-
HHGHDAHAPHDHEGVEHDARSRDVEPAHAQHDHSHDHGHGHAHTP------------------------
HESPLVMLVPLIVLATGALLAGIIFHGAFIGE-GYEE---FW--
KGAVFTRPDNHILEEMHHLPGWVPLLPTLMMILGFLLAVYMYIID-----------------------------
------------------------------
SKQPAKLAADHPILYRFLLNKWYFDELYDAIFVRPAMAIGRFFWRT-GDQRIIDGLGPDGIS-
ARVLDVTRGVVRVQTGYLYHYAFAMLIGVAALVTFYLFRGAH-------
>WP_183451382_NuoL_Microvirga_lupini
MY-------HAIVFLPLVGFLIAGLFG----RVLGPRPSEIITTALLFVAAVLSWVSFIQVGF--
GDGATRIQIA--QWMSVGDLQVDWAFRIDTLTAMMLVVVNTVSALVHLYSIGYMHEDPH-------
RSRFFAYLSLFTFTMLMLVTADNLVQMFFGWEGVGLASYLLIGFWYQKDSANAAAMKAFIVNRVGDFGFLLGIF
TVFVLFNGVTF--DAIFPRVAEFADAK---
FHFLGIEWHALTIACLLLFMGAMGKSAQFLLHTWLPDAMEGPTPVSALIHAATMVTAGVFMVARLSPIFEYAPA
ALTVVTVIGGITAFFAATVGLVQNDIKRVIAYSTCSQLGYMFVGLGVGAYGAGVFHLFTHAFFKALLFLGAGSV
IHAMHHEQDIRHMGALRRYIPFTTAMMAVGTLALTG--FPF---------
SAGYYSKDAIIEAAYAAHSSAGSFAFLATVVAAFMTSFYSWRLFFLTFEGSARWG-
HHDHHAHAPHSHEGVEHDDHNRDVEP---ASHAQHEHGHHGAHTP------------------------
HESPLVMLIPLLVLAIGAVLAGIIFHGAFIGE-GYQE---FW--
KGALFTRPENHILEEMHHLPGWVPLLPTIMMVLGFGLAWYMYIID-----------------------------
------------------------------
NKQPAKLAADHPILYRFLLNKWYFDELYDAIFVRPAMAIGRFFWRT-GDQRIIDGLGPDGIS-
ARVLDVTRGVVRVQTGYVYHYAFAMLIGVAALVTFYLFRGAH-------
>WP_04718718_NuoL_Microvirga_vignae
MY-------HAIVFLPLVGFLIAGLLG----RVIGARPSEIITTALLFVSAVLSWVAFIHVGF--
GEEPMIRVQVA-QWMSVGDLQVDWAFRIDTLTAMMLVVVNTVSALVHLYSIGYMHEDPH-------
RPRFFAYLSLFTFAMLMLVTADNLVQMFFGWEGVGLASYLLIGFWYKKDSANAAAMKAFIVNRVGDFGFLLGIF
TIFVLFNGVTF--DAIFPRVGEFANAR---
FTFLGIEWHALTITCLLLFMGAMGKSAQFLLHTWLPDAMEGPTPVSALIHAATMVTAGVFMVARLSPVFEYAPA
ALTVVTVIGGITAFFAATVGLVQNDIKRVIAYSTCSQLGYMFVALGVGAYSAGMFHLFTHAFFKALLFLGAGSV
IHAMHHEQDMRNMGNLRRYIPLTTAMMAIGTLALTG--FPF---------
TAGYYSKDAIIEAAYASHSAAGSFAFLATVIAAFMTSFYSWRLFFMTFEGNARWGASHHHAPSDHEGVEHDERG
HDVEPAGH----ADHEHGHHGHGAHTP------------------------
```

404… HESPLVMVLPLIVLAIGAVVAGIVFHDAFLGE-GHQE---FW--
… KGALFTGPNNHILEEMHHLPGWVPLLPTVMMILGFVLAYYMYIVD----------------------------
… ------------------------------
… AKQPARLAADHPILYRFLLNKWYFDELYDAIFVRPAMAIGRFFWRT-GDQKIIDGLGPDGIS-
… ARVLDVTRGAVRLQSGYVYHYAFAMLIGVAALVTFYLFRGAH-------
405 >WP_203271756_NuoL_Microvirga_arabica
406 MY-------HAIVFLPLVGFLIAGLFG----RVLGPRPSEIITTALLFVAAVLSWVSFIQVGF--
… GDGATRVQVA--QWMSVGDLQVDWAFRIDTLTAMMLVVVNTVSALVHLYSIGYMHEDPH-------
… RSRFFAYLSLFTFTMLMLVTADNLVQMFFGWEGVGLASYLLIGFWYQKDSANAAAMKAFIVNRVGDFGLLGIF
… TVFVLFNGVTF--DAIFPRVAELANAN---
… FTFLGIEWHALTIACLLLFMGAMGKSAQFLLHTWLPDAMEGPTPVSALIHAATMVTAGVFMVARLSPIFEYAPA
… ALTVVTVIGGITAFFAATVGLVQNDIKRVIAYSTCSQLGYMFVALGVGAYSAGVFHLFTHAFFKALLFLGAGSV
… IHAMHHEQDMRHMGALRRYIPLTTAMMAIGTLALTG--FPF---------
… TAGYYSKDAIIEAAYAAHSTAGSFAFLATVVAAFMTSFYSWRLFFLTFEGTARWG--
… HGHHGHAAHAPHSHEGVEHDDHNRDVEPASHAQHEHGHHGDHTP------------------------
… HESPLVMLLPLIVLAIGAVVAGIVFHNAFIGE-GYQE---FW--
… KGALFTRPENHILEEMHHLPAWVPLLPTVMMALGFVLAYYMYIVD----------------------------
… ------------------------------
… AKQPAKLAADHPILYRFLLNKWYFDELYDAIFVRPAMAIGRFFWRT-GDQRIIDGLGPDGIS-
… ARVMDVTRGVVRVQTGYVYHYAFAMLIGVAALVTFYLFRGAR-------
407 >WP_201834068_NuoL_Microvirga_zambiensis
408 MY-------HAIVFLPLVGFLIAGIFG----RLLGPRPSEIVTTALLFVAAVLSWVSFIQVGF--
… GDGATRIQIA--QWMSVGDLQVDWAFRIDTLTAMMLVVVNTVSALVHLYSIGYMHEDPH-------
… RSRFFAYLSLFTFTMLMLVTADNLVQMFFGWEGVGLASYLLIGFWYQKDSANAAAMKAFIVNRVGDFGLLGIF
… TVFVLFNGVTF--DAIFPRVSEFANAR---
… FMFLGIEWHALTIACLLLFMGAMGKSAQFLLHTWLPDAMEGPTPVSALIHAATMVTAGVFMVARLSPLFEYAPS
… ALAVVTVIGGITAFFAATVGLVQNDIKRVIAYSTCSQLGYMFVGLGVGAYGAGVFHLFTHAFFKALLFLGAGSV
… IHAMHHEQDIRHMGALRRYIPLTTAMMAIGTLALTG--FPF---------
… TAGYYSKDAIIEAAYAAHSSAGSFAFLATVVAAFMTSFYSWRLFFLTFEGTARWG-
… HHDHHAHAPHSHEGVEHDDHNRDVEPAS---HAQHDHGHHGDHTP------------------------
… HESPLVMLIPLIVLATGAVIAGIVFHGAFIGE-GYQE---FW--
… KGALYTRPDNHILEEMHHLPGWVPLLPTIMMVLGFVLAWYMYIID----------------------------
… ------------------------------
… NKQPAKLAADHPILYRFLLNKWYFDELYDAIFVRPAMAIGRFFWRT-GDQKIIDGLGPDGIS-
… ARVLDVTRGVVRVQTGYVYHYAFAMLIGVAALVTFYLFRGAH-------
409 >WP_183332122_NuoL_Chelatococcus_composti
410 MY-------HAIVFLPLAGFLIAGLAG----RFIGARASEVITTSGLFVSCALSWIAFFDVAF--
… GDGATRVTVA--NWIISGDLAVDWAFRIDTLTAVMLIVVTTVSSLVHLYSIGYMHEDPS-------
… RPRFFAYLSLFTFAMLMLVTADNLVQMFFGWEGVGLASYLLIGFWYEKPSANAAAIKAFVVNRVGDFGLLGIF
… LLFVLFGAVTF--DGIFPRVAEFNEAT---
… FRFLGHEWHALTLACLLLFMGAMGKSAQFLLHTWLPDAMEGPTPVSALIHAATMVTAGVFMVARMSPVFEYAPT
… ALSVVTVIGGITAFFAATVGLVQNDIKRVIAYSTCSQLGYMFVALGVGAYGAGVFHLFTHAFFKALLFLGAGSV
… IHAMHHEQDMRHMGGLRKHIPFTAAMMTIGTLALTG--FPF---------
… TAGYFSKDAIIEAAYASHTPGANFAFIATVIAALFTSFYSWRLYFLTFEGKPRW-AGH-GHGHDD--------
… -----------------HGHGHGDHTP------------------------
… HESPLVMLIPLAVLAIGVVVAGFAFKEAFIGH-DYDN---FW--
… KGALFTGANNHILHEMHEVPGWVVASPFVMMVLGFLIALWFYVLN----------------------------
… ------------------------------

```
PRMPAALAAQQPLLYRFLLNKWYFDELYDFLFVRPAMWLGRVFWKK-GDGWLIDGFGPDGVS-
ARVVDVTNRVVRLQTGYVFHYAFAMLVGVAALVTWYLIGGTH-------
>WP_069691719_NuoL_Bosea_vaviloviae
MY-------HAIVFLPLIGFLIAGLFG----RLIGARGSEIVTTSLLVVSAVLSWIAFFQVGF--
GHGTTRVQIA--TWMASGDLRVDWAFRVDTLTAVMLVVVNTVSCLVHLYSIGYMHEDPH-------
RPRFFAYLSLFTFAMLMLVTADNLVQMFFGWEGVGLASYLLIGFWYQKPSANAAAMKAFIVNRVGDFGFALGIF
LVFVLFGSVGF--DAIFPRVGDLTSQS---
FHFLGRDWHALTLASLLLFMGAMGKSAQFLLHTWLPDAMEGPTPVSALIHAATMVTAGVFMVARLSPIFEYAPV
ALTVVVVIGATTAFFAATVGLVQNDIKRVIAYSTCSQLGYMFVALGVGNYGAGIFHLFTHAFFKALLFLGAGSV
IHAMHHEQDMRNMGGLRKHIPLTAIAMTIGTLALTG--FPG---------
FAGYFSKDAIIESAYASVAHGGFASSYAFVLLVCMTSFYSWRLYFMTFEGKPRWGADA-HAAHAHAAHAHDD--
-----------------HHGHGHDHTP------------------------
HESPWVMLIPLAVLSLGAVAAGFVFKDAFIGH-DYEH---FW--
KAALFTGKDNHILHAMHEVPGWVVASPFVAMLIGLALAYYMYVRR-----------------------------
------------------------------
PDVPGKLAAANPALYKFLLNKWYFDELYDFLFVKPAMWLGKFLWKK-GDGFVIDGMGPDGIS-
ARVVDVTNRVVRLQTGYVYHYAFAMLIGVAGFVTWYLVARG--------
>WP_181052769_NuoL_Microvirga_mediterraneensis
MY-------HAIVFLPLVGFLIAGIFG----RVLGARPSELITTALLFVAAVLSWVSFIHVGF--
GDEPMVRVQVA-QWMSVGDLQVDWAFRIDTLTAMMLVVVNTVSALVHLYSIGYMHEDPH-------
RPRFFAYLSLFTFAMLMLVTADNLVQMFFGWEGVGLASYLLIGFWYQKDSANAAAMKAFIVNRVGDFGFLLGIF
TIFVLFNGVTF--DQIFPRVAEFAEAK---
FHFLGIEWHALTIACLLLFMGAMGKSAQFLLHTWLPDAMEGPTPVSALIHAATMVTAGVFMVARLSPVFEYAPA
ALTVVTVIGGITAFFAATVGLVQNDIKRVIAYSTCSQLGYMFVALGVGAYGAGVFHLFTHAFFKALLFLGAGSV
IHAMHHEQDIRHMGALRRYIPFTTAMMAIGTLALTG--FPF---------
TAGYYSKDAIIEAAYAAHSSAGSFAFLATVVAAFMTSFYSWRLFFLTFEGSARWGHGHHSPSAHEGVEHDESGH
DVEPASHAQHEHGHDDHAHGHHGSHMP------------------------
LESPLVMLLPLLVLAIGAVVAGFMFHGAFIGE-GYQE---FW--
KGALFTRPENKILETMHHLPGWVPLLPTIMMILGFLVAVYMYIID-----------------------------
------------------------------
NKQPAKLAADHPILYRFLLNKWYFDELYDAIFVRPAMAIGRFFWRT-GDQRIIDGLGPDGIS-
ARVLDVTRGVVRVQTGYVYHYAFAMLIGVAALVTFYLFRGAH-------
>WP_027316570_NuoL_Microvirga_flocculans
MY-------HAIVFLPLVGFLIAGLFG----RVLGARPSEIVTTALLFVAAVLSWVSFIQVGF--
GDGATRVQIA--QWMSVGDLRVDWAFRIDTLTAMMLVVVNTVSALVHLYSIGYMHEDPH-------
RPRFFAYLSLFTFAMLMLVTADNLVQMFFGWEGVGLASYLLIGFWYQKDSANAAAMKAFIVNRVGDFGFLLGIF
TVFVLFNGVTF--DAIFSRVGEFANAK---
FHFLGIEWHALTIACLLLFMGAMGKSAQFLLHTWLPDAMEGPTPVSALIHAATMVTAGVFMVARLSPVFEYAPA
ALTVVTVIGGITAFFAATVGLVQNDIKRVIAYSTCSQLGYMFVALGVGAYGAGVFHLFTHAFFKALLFLGAGSV
IHAMHHEQDMRHMGALRRYIPITTAMMAIGTLALTG--FPF---------
TAGYYSKDAIIEAAYASHSSAGSFAFLATVVAAFMTSFYSWRLFFMTFEGSARWG--
HGDHAHAHAPHDHEGVEHDAHGRDVEAASH-AQHEHGHHGGHTP------------------------
HESPLVMLLPLFVLAAGAVVAGAVFHGAFLGE-GYEE---FW--
KGALFTRPENKILEAMHHLPVWVPLLPTLMMALGFALAYYMYIVD-----------------------------
------------------------------
AKQPAKLAADHPILYRFLLNKWYFDELYDAIFVRPAMAIGRFFWRT-GDQRIIDGLGPDGIS-
ARVLDVTRGVVRVQTGYVYHYAFAMLIGVAALVTFYLFRGAH-------
>WP_112664104_NuoL_Microvirga_flavescens
```

```
MY-------YAIVFLPLVGFLIAGLFG----RVIGARPSELITTAFLFIAAALSWVAFFQVGF--
GHGAVRIQIA--QWFSSGDLMVDWALRIDTLTAIMLIVVNTVSALVHLYSIGYMHEDPH-------
RPRFFAYLSLFTFAMLMLVTADNLVQMFFGWEGVGLASYLLIGFWYQKESANAAAMKAFIVNRVGDFGFLLGIF
LIFVLFKTVNF--DQIFPRVAEFEKAQ---
FHFLGFDWNALTLACLLLFMGAMGKSAQFLLHTWLPDAMEGPTPVSALIHAATMVTAGVFMVARLSPVFEYAPD
ALTVVTVIGGITAFFAATVGLVQNDIKRVIAYSTCSQLGYMFVALGVGAYSAGVFHLFTHAFFKALLFLGAGSV
IHAMHHEQDIRAMGGLRKYIPFTTGMMAIGTLALTG--FPF---------
SAGYYSKDAIIEAAYAAHSPAGSFAFLATVVAAFMTSFYSWRLFFLTFEGKARWN--
GEAHTHEGAEHNASGHDVSHAHAD-------HSHDDHGHGAHKP------------------------
HESPLVMLIPLLVLAIGAVVAGFAFSGAFIGH-GYEH---FW--
KGAIFTRPDNKILEDMHHLPGWVPLLPTVMMVLGFLLAVWMYLID-----------------------------
------------------------------
TKKPAQIAADHPILYRFLLNKWYFDELYDAIFVRPAMALGRFLWRT-GDGKIIDGLGPDGIS-
ARVLDVTRGVVRVQTGYLYHYAFAMLIGVAALVTYYLFRGAH-------
>WP_202055072_NuoL_Microvirga_aerilata
MY-------HAIVFLPLVGFLIAGLFG----RLLGPRPSEIVTTALLFVAAVLSWVSFIQVGF--
GDGATRVQVA--QWMSVGDLVVDWAFRIDTLTAMMLVVVNTVSALVHLYSIGYMHEDPH-------
RPRFFAYLSLFTFAMLMLVTADNLVQMFFGWEGVGLASYLLIGFWYQKDSANAAAMKAFIVNRVGDFGFLLGIF
TLFVLFNGVTF--DQIFPRAAELANAN---
FHFLGIEWHALTIACLLLFMGAMGKSAQFLLHTWLPDAMEGPTPVSALIHAATMVTAGVFMVARLSPVFEYAPA
ALTVVTVIGGITAFFAATIGLVQNDIKRVIAYSTCSQLGYMFVGLGVGAYSAGVFHLFTHAFFKALLFLGAGSV
IHAMHHEQDMRNMGALRRYIPVTTAMMAIGTLALTG--FPF---------
TAGYYSKDAIIEAAYASHSTAGTFAFLATVIAAFMTSFYSWRLFFLTFEGTARWS-
NADHHAHGHHEGVEHDGHSH--DVEPAT---HAQHEHGHHGDHTP------------------------
HESPLVMLLPLVVLAIGAVVAGIVFHGAFIGE-GYAD---FW--
KGALFTRPENHILEEMHHLPGWVPLLPTVMMILGFVLAVYMYMID-----------------------------
------------------------------
AKKPAQLAAEHPVLYRFLLNKWYFDELYDLLFVRSAMWIGRFFWRT-GDQRIIDGLGPDGIS-
ARVMDVTRGVVRVQTGYVYHYAFAMLIGVAALVTFYLFRGAH-------
>WP_009491127_NuoL_Microvirga_lotononidis
MY-------HAIVFLPLVGFLIAGIFG----RVLGARPSEIVTTALLFVAAVLSWVSFIHVGF--
GDEPMIRVQIA-QWMSVGDLQVDWAFRIDTLTAMMLVVVNTVSALVHLYSIGYMHEDPH-------
RPRFFAYLSLFTFAMLMLVTADNLVQMFFGWEGVGLASYLLIGFWYQKDSANAAAMKAFIVNRVGDFGFLLGIF
TIFVLFNGVTF--DQIFPRVAEFADAK---
FHFLGIEWHALTIACLLLFMGAMGKSAQFLLHTWLPDAMEGPTPVSALIHAATMVTAGVFMVARLSPVFEYAPA
ALTVVTVIGGITAFFAATVGLVQNDIKRVIAYSTCSQLGYMFVGLGVGAYGAGVFHLFTHAFFKALLFLGAGSV
IHAMHHEQDIRHMGALRRYIPFTTAMMAIGTLALTG--FPF---------
TAGYYSKDAIIEAAYAAHSSAGSFAFLATVVAAFMTSFYSWRLFFLTFEGSARWG-
HHDHHAHAPHDHEGVEHDDHNRDVEPAT---HAQHEHGHHGSHMP------------------------
HESPLVMLLPLLVLAIGAVVAGFVFHGAFIGE-GYQE---FW--
KGALFTRPENEILETMHHLPGWVPLLPTIMMILGFLVAVYMYIID-----------------------------
------------------------------
NKQPAKLAADHPILYRFLLNKWYFDELYDAIFVRPAMAIGRFFWRT-GDQRIIDGLGPDGIS-
ARVLDVTRGVVRVQTGYVYHYAFAMLIGVAALVTFYLFRGAH-------
>WP_106335304_NuoL_Alsobacter_soli
MY-------QAIVFLPLVGFLIAGLFG----RIIGARASEVVTTAFLMAAALLSWIAFFQVGL--
GHHEALRVQVA-QWMVAGDLKVDWAFRIDTLTVVMLVVVNTVSSLVHLYSIGYMHEDPH-------
RPRFFAYLSLFTFAMLMLVTSDNLVQMFFGWEGVGLASYLLIGFWYQKPSAVAAAMKAFIVNRVGDFGFSLGIF
```

```
LVFVLSGSVAF--PEIFPKAEEIAKGT---
FHFIGYDWNAMTIACLLLFMGAMGKSAQFLLHTWLPDAMEGPTPVSALIHAATMVTAGVFMVARLSPLFEQSHV
ALSVVMIVGATTAFFAGTIGLVQNDIKRVIAYSTCSQLGYMFVALGAGAYGAGIFHLFTHAFFKALLFLGAGSV
IHAMHHEQDIRHMGGLARKIPFTTVMFAIGTLALTG--FPL---------
TAGYYSKDAIIESSYAAHTFGNAYAFILLSITAGLTSFYSWRLFFVTFMGAPRWD---
TPHGHDLAVSHHVESEDDSHGS-------HAHAGDHGHGHHEP------------------------
HESPLTMLIPLGVLGLGALFAGLVFEGYFFGH-AYDE---FW--
KNALHTGPENHILHAVHEIPAWAKYSPTVEMIIGFVVAYIFYIQK-----------------------------
------------------------------
PYLPRQLAESQPLLYRFLLNKWYFDELYDVIFVRPAKWLGRFLWKK-GDGAVIDGFGPDGVS-
ARVVDVTQAVVRLQTGYVYHYAFAMLIGVAALVTWYMFGGAH-------
>WP_020175212_NuoL_Methyloferula_stellata
MY-------FAIVFLPLAGFLIAGLFG----RQIGARASEIITTSFLFGAAFLSWIAFSQVAL--
GSTPASVPVIG-TWFNVGALQVDWALRIDSLTVVMLIVVNTVSALVHLYSIGYMHEDPA-------
RPRFFAYLSLFTFAMLMLVTADNLVQMFFGWEGVGLASYLLIGFWYQKPSANAAAIKAFVVNRVGDFGFAIGIF
LVFYITNSVAF--DPIFAAAPGLAHKT---
IHVFSHDVDAMTITCLFLFMGAMGKSAQFLLHTWLPDAMEGPTPVSALIHAATMVTAGVFMVARLSPLFEQAPV
ALSFVTIIGGTTAFFAATVGLVQNDIKRVIAYSTCSQLGYMFVGLGVGGYDLGIFHLFTHAFFKALLFLSAGSV
INAMHHEQDMRKMGGLAKKIPFTFWMMVIGTLALTG--FPL---------
TAGFFSKDAIIEAAFASGRFGSFYAFLLIDFAAGLTSFYSWRLIFMTFYGPAHWD-HHGVDHHDEHAAHDE---
-----------------AHGHAHTFVP------------------------
HESPAVMLIPLAVLAFGSIFAGLAFGGFFIGE-GQAD---FW--
KGALFYASDNHILHEMHTIPAFVSYSPFWMMCGGFLVALYVYILR-----------------------------
------------------------------
PGTAANWAAANPALYKFLLNKWYFDELYDLIFVRPAFWLGRLFWKG-GDGAIIDGLGPDGVS-
ARVIDITHRVVKLQTGYIYHYAFAMLIGVAALMTYYLFGGIR-------
>WP_133769151_NuoL_Enterovirga_rhinocerotis
MY-------HAIVFLPLVGFLIAGLFG----RFIGARPSELITTGLMMVAALLSWVAFISVGF--
GSGATRVQVA--NWFTSGELVIDWAFRIDTLTAVMLVVVNSVSALVHLYSMGYMHEDPH-------
RPRFFAYLSLFTFAMLMLVTSDNLVQMFFGWEGVGLASYLLIGFWYEKPSANAAAMKAFIVNRVGDFGFALGIF
LVFVLFGTTTY--SEIFAKIGEVKDAH---
FHFIGFEWHALTLACLLLFMGAMGKSAQFLLHTWLPDAMEGPTPVSALIHAATMVTAGVFMVARLSPLFDAAPA
ALTVVTIIGAITAFFAATVGLVQNDIKRVIAYSTCSQLGYMFVALGVGAYGAAVFHLFTHAFFKALLFLGAGSV
IHAMHHEQDMRNMGGLRKYIPFTWAMMLVGTLALTG--FPF---------
TAGYFSKDAIIEAAFMSDRPGHAIAFVSTLVAAFMTSFYSWRLAFMTFEQTARWGAGHGHDAHGDAAHGDAGHG
HAAHADHAHDEHAAHGHDDHHGHGHDHKP----------------------
HESPLVMLVPLGVLAFGAIFAGMIFAGSFVGH-HAAD---FW--KGALAESQYAVFEAMHH-
APKWVVWSPFVAMVLGWALAWVMYIRK-----------------------------------------------
------------PYLPGELAASQPIAYRFLLNKWYFDELYDFLFVRPAKRLGYFLWKR-GDGTVIDGMGPDGIS
-ARVVDVTRGVVRLQTGYVYHYAFVMLIGVAALVSWYLIVGAGAH-----
>WP_088519014_NuoL_Rhodoblastus_acidophilus
MY-------SAIVFLPLLGFLIAGLLG----KKIGDRASEIVTTALLFVSCALSWITFFDVGL--
HDGHFSAPVIA-NWLTSGDLKIDWALRVDTLTSVMLIVVTTVSALVHLYSIGYMHEDKS-------
RPRFFAYLSLFTFAMLMLVTSDNLAQMFFGWEGVGLASYLLIGFWFEKPSANAAAIKAFVVNRVGDFGFSLGMF
LVFMLTGSISF--DVIFAAAPDLASKS---
VHAFGLDWNAMTLACLLLFMGAMGKSAQFLLHTWLPDAMEGPTPVSALIHAATMVTAGVFMVARLSPLFEQSAT
ALTVVTLVGGITAFFAATVGLVQNDIKRVIAYSTCSQLGYMFVGLGCGGYALGIFHLFTHAFFKALLFLGAGSV
IHAMHHEQDMRQMGGLWSKIPVTFACMTIGTLALTG--FPL---------
```

```
TAGYFSKDAIIEGAFAAGAHSFAASFALLVVAAGMTSFYSWRLVFMTFFGERRWS-
EQYKLLHPAPQAAHGHGHGDDGHG--------AHDDPHGHHIGEP------------------------
HESPATMMVPLFVLSAGALAAGYAFKDQFIGH-EAGE---FW--
RTSLFTGANNHILHEMHEIPAWVGYAPFAMMVGGFVVALAMYVIA-----------------------------
------------------------------
PKVPAALAEKFPRLYLFLLNKWYFDELYDKIFVKPAFALGRLFWKG-GDGAIIDRFGPDGVA-
ARVLDGAGVASRLQTGFIYNYAFAMLLGVAALITYGLFALGGGQ-----
>WP_188518029_NuoL_Alsobacter_metallidurans
MY-------QAIVFLPLVGFLIAGLFG----RIIGARASEIVTTAFLMAAALLSWVAFFQVGL--
GHGETRVQVA--QWMVSGDLKVDWAFRVDTLTVVMLVVVNSVSSLVHLYSIGYMHEDPH-------
RPRFFAYLSLFTFAMLMLVTADNLVQMFFGWEGVGLASYLLIGFWYNKPSAVAAAMKAFIVNRVGDFGFALGIF
LVFVLTGSVAF--SEIFPKVEEIAKGT---
FHFIGYDWNAMTIACLLLFMGAMGKSAQFLLHTWLPDAMEGPTPVSALIHAATMVTAGVFMVARLSPLFETSPV
AMSAVLFVGATTAFFAGTIGLVQNDIKRVIAYSTCSQLGYMFVALGAGAYGAGIFHLFTHAFFKALLFLGAGSV
IHAMHHEQDIRHMGGLARKIPFTTAMFAVGTLALTG--FPL---------
TAGYFSKDAIIESSYATHTYGNAYAFILLSITAGLTSFYSWRLFFLTFMGQPRWDTAHGHDAHASGHVESEDDS
HGSH----------AHAGDHGHGHHEP------------------------
HESPLSMLIPLGVLGVGALFAGLAFESFFFGH-AYEE---FW--
KGAVHTGPDNHILHAIHEVPGWVKYSPTVEMIIGFVVAYIFYVQK-----------------------------
------------------------------
PYLPRQLAESQPTLYRFLLNKWYFDELYDVIFVRPAKWLGRFLWKK-GDGLVIDGMGPDGVS-
ARVVDVTNAVVRLQTGYVYHYAFAMLIGVAALITWYMFGGAH-------
>WP_104509173_NuoL_Rhodoblastus_sphagnicola
MY-------SAIVFLPLLGFLIAGLAG----RKIGDRASEIVTTTLLFVSCALSWKVFLDVGL--
NDGHFSAPVLA-NWLTSGDLKIDWALRVDTLTSVMLVVVTTVSALVHLYSVGYMSEDPS-------
RPRFFAYLSLFTFAMLALVTADNLAQMFFGWEGVGLASYLLIGFWYEKPSANAAAIKAFVVNRVGDFGFSLGMF
LIFLLTGSISF--DTIFAAAPGLADKS---
IHAFGLDWNAMTLTCLLLFMGAMGKSAQFLLHTWLPDAMEGPTPVSALIHAATMVTAGVFMVARLSPLFEQSPT
ALTVVTLVGATTAFFAATVGLVQNDIKRVIAYSTCSQLGYMFVGLGCGGYALGIFHLFTHAFFKALLFLGAGSV
IHAMHHEQDMRQMGGLWTKIPITFACMTIGTLALTG--FPF---------
TAGYFSKDAIIEGAFAASGHSAAAFYAALVVAAGMTSFYSWRLVFMTFFGPRRWS-DEYKALHGHAAAQDDH--
--------------GHDDAHAHHIGEP------------------------
HESPLTMMIPLFVLSAGALGAGYVFKDHFINE-GAQE---FW--
RTSLFTGANNHILHEMHQTPYWVEPLPLLMMGGGFLVALVLYVIA-----------------------------
------------------------------
PKIPAALAARYPKLYLFLLNKWYFDELYDRIFVKPAFWLGRLFWKG-GDGAIIDRLGPDGIA-
ARVIDGAGLASRLQTGFIYNYAFAMLLGLAALITYGLFAVGGGQ-----
>WP_108659702_NuoL_Acuticoccus_kandeliae
MV-------QLIVFLPLFGFLFAFLFG----RQVGALAAQVVTTGLLFVCAVLSWIVFFNWGF--
HGAPTERILIA-QWIVSGDLEIAWRLRVDTLTAVMLIVVNTVSALVHLYSFGYMSHDPA-------
KAKFFSYLSLFTFAMLMLVTSDNLVQMFFGWEGVGLASYLLIGFWSDRPSANAAAIKAFVVNRVGDFGFLLGIF
TLFAMTGVVQL--DAVFASAPALSTHT---
FTFLGFEVPAIELICLLLFIGAMGKSAQIFLHTWLPDAMEGPTPVSALIHAATMVTAGVFMVARMSPVFEFAPS
ALAFVTFIGATTAMFAATIGCAQNDIKRVIAYSTCSQLGYMFAALGVGAFGIAIFHLFTHAFFKALLFLSAGSV
IHAMHDEQDMRRMGGLWKKIPYTYIGMMAGTLALTG--FPL---------
TAGYFSKDAIIEATFVGENPFAIYAFIMTVGAALLTSFYSWRLVFMTFHGTTRAPADTFKHA------------
---------------------------------------------------
HESPWTMLVPLGVLTLGALAAGYVFATVFLSE-PTGD---FW--
```

```
NGAIFASAANHVLHAMHEVPTWVILSPAIAMVIGFCIAVLTYILV-----------------------------
------------------------------
PTLPKRFIEAFPRVHQFFLNKWYFDELYDYIFVRPARWLGRMFWR--GDGRVIDGVGPDGIA-
ARVKDLSVIAHRFQSGYLYHYAFVMLTGIAVFVTVMIALNRGYL-----
>WP_012170207_NuoL_Azorhizobium_caulinodans
MY-------QAIVFLPLIGFLIAGLFG----RVLGPRPSELITTALLFIACLFAWITFIKVGF--
AGFDTRVQVM--HWIYVGDLKIDWALRIDTMTAIMLIVVTSVSSLVHLYSIGYMHEDPS-------
RPRFFAYLSLFTFAMLMLVTADNLVQLFFGWEGVGLASYLLIGFWFEKPSANAAAMKAFVVNRVGDFGFMLGIF
AVFVMTGSVSF--DTIFAQAPSLAGKT---
ITFLGHHWDAPTVIALLLFIGAMGKSAQFLLHTWLPDAMEGPTPVSALIHAATMVTAGVFMVARMSPIFELSPS
ALDVVTIVGATTAMFAATVALVQNDIKKVIAYSTCSQLGYMFVAMGAGAYSVGVFHLFTHAFFKALLFLGAGSV
IHAMHHEQDMRHMGGLYKKIPFTYGAMMIGTLAITG--FPF---------
LAGYYSKDAIIEAAYASHSHFRVYAYWMTVLAAALTSFYSWRLVFLTFHGHPHDH-HHYDHA------------
---------------------------------------------------
HESPLVMTIPLAVLSVGAVIAGFVCYNLFVGH-DVEH---FF--
RSSIFMGPDNHILHAMHEVPGWAAYMPTVMMVLGFVVAYWFYMVD-----------------------------
------------------------------
RRVPAALAKSQDALYQFLLNKWYIDELYNFLFVRPALCLGRILWKK-GDGAIIDGLGPNGIS-
ARVLDVTGQLVRLQTGYLYHYAFVMLVGVAAIITWFMFAGV--------
>WP_05503695_NuoL_Blastochloris_viridis
MY-------QLIVFLPLLGCVIAGLFG----RVIGSRPSEIVTTALVGVSALLSWIAFFNVGY--
DGESGRILVS--TWIEAGSFKADWAFRIDTLTVVMLVVVNTVSALVHFYSIGYMADDPH-------
RPRFFSYLSLFTFAMLMLVTSDNLVQMFFGWEGVGLASYLLIGFWYLKPSANAAAIKAFLVNRVGDFGFALGIF
AIFWMVGSTDF--TTIFAQVPTLQDKV---
IAVAGVPFDALTLICLLLFMGAMGKSAQFLLHTWLPDAMEGPTPVSALIHAATMVTAGVFMVARLSPLFELAPT
AQEVVILVGATTAFFAATVGLVQNDIKRIVAYSTCSQLGYMFVAMGVGAYSVGMFHLFTHAFFKALLFLGSGAV
IIAMHHEQDVRKMGGLRKAIPFTYAMMVIGTLALTG--FPF---------
TAGYYSKDAIVESAFVSHAALSGYGFWMLVIAAALTSFYSWRLVFLTFHGAPRFVAADGHHGHGDHGHHGVSRI
EDV------------------------------------------------
HEAPKTMLLPLLVLAVGALGAGFAFYPVFVDQ-HGVAE--FF--RDSISLSKL-
GLLEEMHHAPALVVFAPTIAMAAGLAVAVLFYLVS---------------------------------------
--------------------PGLPVALARSMRPLYLFLLNKWYFDELYDVIFVRAAKRLGRFLWKQ-
GDGAVIDGLGPDGIA-ARVLDTTDRVVRLQTGYLYHYAFAMLIGLAALVTWFMFAGGVR------
>WP_183750461_NuoL_Pseudochelatococcus_contaminans
MY-------QAIVFLPLVGFLVAGLFG----RFIGARASEIITTSLLFVTAALSWVAFVTVGF--
GTETFYVPIA--RWITSGTLIVDWALRIDTLTVVMLVVVNSVSALVHLYSIGYMHEDPD-------
RPRFFAYLSLFTFAMLTLVTADNLVQMFFGWEGVGLASYLLIGFWYEKPSACAAAMKAFIVNRVGDFGFMLGLL
LTFIVFGSVSF--GDIFSRTAEFQDAT---
FNFIGFEWHTLTLISLLLFIGAMGKSAQFILHTWLPDAMEGPTPVSALIHAATMVTAGVFMVTRMSPVFEYAPS
ASTVIIVIGAVTAFFAATIGLVQNDIKRVIAYSTCSQLGYMFVALGVGAYSAGVFHLFTHAFFKALLFLGAGSV
IHAMHHEQDMRKMGGLRKYIPFTALAMTIGTLALTG--FPF---------
TSGFYSKDAIIESAFASHSSVAGFAFTSTIVAALMTSFYSWRLYFMTFEGKPRWE-
HHDDHAHGHDHAHDDGHDHKP------------------------------------------------
HESPLVMLLPLVVLSIGALFAGAVFYNAFIGS-GYDA---FW--
KGAIFTNPDNHILHDMHEVPLWVKLAPVVTMVIGFLLALWFYVLD-----------------------------
------------------------------
TSKPRQLAAAFPGVYRFLLNKWYIDELYNFLFVRPAKELGRFFWKQ-GDGWLIDGHGPDGIS-
ARVLDVTGRVVKLQTGYVYHYAFAMLIGVAAFVTWYIFGGAV-------
```

```
>WP_015821652_NuoL_Methylorubrum-extorquens
MY-------HAIVFFPLIGALIAGLFG----RFIGARMSELVTTGCLAFAALLSWGAFFLVTG--
DGRAETVPVA--QWFTAGDLVVDWAFKVDTLTAVMLVVVTSVSTLVHLYSIGYMHEDPH-------
RPRFFAYLSLFTFAMLMLVTADNLVQMFFGWEGVGLASYLLIGFWYEKPSANAAAMKAFIVNRVGDFGFSLGIF
LVFVLTGSVGF--DAIFAKAPELKDAT---
FHFLGHDWHALTLACLLLFMGAMGKSAQFLLHTWLPDAMEGPTPVSALIHAATMVTAGVFMVARLSPLFELAPT
ALTVVTVIGGITAFFAATVGLVQNDIKRVIAYSTCSQLGYMFVGLGVGAYATGVFHLFTHAFFKALLFLGAGSV
IHAMHHEQDMRNMGGLRRYIPFTTAMMTIGTIALIG--FPF---------
TSGYYSKDAIIEAAYMSDRPGHVLAFLATVIAALMTSFYSWRLFFLTFEGPQRWVAH-
GAHGHDDHAHAEHDHVHAASAHEADGAPGHHEGVAHDDHGHDVEPASHSAV--EHHDHAPLTP----
HESPLVMTIPLAILAFGALFAGLIFKERFIGH-DMDK---FW--
GNALPHGSGNDIMHKIHDAPGWVAASPFVMLVLGFLLAFWMYLRR-----------------------------
------------------------------
PDLPHRLAESQPILYRFLLNKWYFDEIYDRIFVRPAKNFGLFLWKE-GDGRVIDGLGPDGIS-
ARVVDITRGVVRLQTGYVVYHYAFVMLVGVAGLITWYLVSGVP--GGTH-
>KIU34726_NuoL_Methylobacterium_radiotolerans
MY-------HAIVFFPLIGFLIAGLFG----RYIGARACEYITTAFLAFTALIAWGVFLDG----GHAE-
RVQVA--SWFSAGDLQVDWAFKVDTLTRVMLVVVTTVSTLVHLYSVGYMEEDPH-------
RPRFFAYLSLFTFAMLMLVTADNLVQMFFGWEGVGLASYLLIGFWYEKPSANAAAMKAFIVNRVGDFGFSLGIF
LTFVLTGSVAF--DAIFGKVDAIKTLT---
FHFLGYDWNALTLACLLLFMGAMGKSAQFLLHTWLPDAMEGPTPVSALIHAATMVTAGVFMVARLSPLFEEAPN
ALIVVTVIGGVTAFFAATIGLVQNDIKRVIAYSTCSQLGYMFVALGVGAYSAGVFHLFTHAFFKALLFLGAGSV
IHAMHHEQDMRNMGGLRRYIPYTSAMMAIGSLALIG--FPY---------
TAGYYSKDAIIEAAYASTRPGHTLAFLCVVVAAFFTSFYSWRLFFMTFEGPARWGAHA-
AHSHHDAPAVAHSTMAHEADGAPGHTEGVAHDDKGHDVEPAHRSDLVADDHHGHGHHGDHTP----
HESPLVMTIPLAILAFGALFAGIIFHNRFIGE-GMDA---FW--
GHALAHGPNNHIMHEIHEVPALVSYSPLVMLILGFVLAYWMYIRR-----------------------------
------------------------------
PELPGQLAAQQPSLYRFLLNKWYFDEIYDRIFVRPAKDFGLFLWKE-GDGRIIDGLGPNGIA-
ARVVDVTRGVVRLQTGYLYHYAFVMLIGVAGLISWYLMSGLP--KGAH-
>WP_18891554_NuoL_Salinarimonas_ramus
MY-------QAIVFLPLLGFLVAGIFG----RFIGARPSEIVTTALLVVSAALSWVAFFDVAL--
GEGPEVFRVQVATWMVSGGLSVDWAFRIDTLTVVMLVVVNTVSALVHLYSIGYMHEDPS-------
RPRFFAYLSLFTFAMLMLVTSDNLVQMFFGWEGVGLASYLLIGFWYQKPSANAAAMKAFVVNRVGDFGFALGIF
AVFFLFGSVQF--EEIFAAIPAYAETGGP-
VLLGLSSEGAITLAALLLFMGAMGKSAQFLLHTWLPDAMEGPTPVSALIHAATMVTAGVFLVARMSPLFEFAPT
ALTVVTYVGALTAFFAATVGLVQNDIKRVIAYSTCSQLGYMFVALGIGAYGAGIFHLFTHAFFKALLFLGAGSV
IHALHHEQDLRNMGGLRRHIPFTAAMMAIGTLALTG--FPF---------
TAGYYSKDAIIEAAYASPTTAGDVAFVATVIAALFTSFYSWRLFFLAFEGAPRWA-
GHGDATHAAHGHDDHEGVAHGAHDQDLEPAHAAAHEHA-DHGHG----------HGHGHGHGHAP----
HESPLVMTIPLGVLAVGALLAGMLFKESFVGH-DWHH---FW--
EGAIYTREGNTIMEDFHHVPAWVVWSPFVMMVLGFLGAFWMYIRD-----------------------------
------------------------------
KAAPARLAGRHPVIYRFLLNKWYIDELYDMLFVQPSKRLGRFLWRT-GDGRVIDGLGPDGVS-
ARVLDATSWVVRLQTGYVYHYAFVMMIGVAALVTWYLVTGAR-------
>SEC00671_NuoL_Rhizobiales_GAS188
MY-------SAIVFLPLFGFIIAGGFG----RIIGARGAEVVTTGFLLLGMVLSWIAFFQVGL--
GHQSFHVDIA--PWIISGGLRTDWALRIDTLTAVMLVVVNTVSGLVHLYSIGYMAEDTS-------
```

```
RPRFFAFLSLFTFAMLMLVTADNLVQMFFGWEGVGLASYLLIGFWYDRPSANAAAIKAFIVNRVGDFGFALGIF
LVFTLFGTVSL--SDIFARAPEMAKGS---
YHFIGFDWPALTLACLLLFMGAMGKSAQFLLHTWLPDAMEGPTPVSALIHAATMVTAGVFMVARLSPLFEQAPA
AQTVVLAIGGMTAFFAATVGLVQNDIKRVVAYSTCSQLGYMFVGLGAGAYSVGIFHLFTHAFFKALLFLGAGSV
ITAMHHEQDMRKMGGLWKSIPFTGWMMVIGTLALTG--FPL---------
TAGFYSKDAIIEAAYATDRPGHVFAFLCTTIAALMTSFYSWRLIFMTFFGTPHWAAADGHGAADAHAHADAHGD
DDAHEHHGPIVP------------------------------------
HESPLVMLVPLAVLALGSLFAGLAFKGVFFGG-AAEH---FW--NNSLIFDED--
IVKKIEEVPFWVEHSALVMMLIGGGVAYWFYVAN------------------------------------
------------------PSLPVRLAARNPILYQFLLNKWYFDEIYEVIFVRPAKAIGRFLWKQ-
GDGRMIDGLGPDGVS-ARVIDVARGAVRLQSGYLYHYAFAMLIGVAAFITYFMIAGAR-------
>WP_106748567_NuoL_Phreatobacter_cathodiphilus
MY-------QAIVFLPLVGFLIAGLFG----RAIGARASEIVTTSLLGIAALLSWVAFYQVGY-
AGQGTTVELMR---WISAGSLNVSWSLRIDTLTAVMLVVVNSVSFLVHLYSIGYMHEDPD-------
RPRFFAYLSFFTFAMLMLVTANDLLQMFFGWEGVGVASYLLIGFWYKKPSANAAAMKAFIVNRVGDFGFLLGIF
GLYTLTGSIGF--EAVFSAAQGLSGQK---
MAFLGYQLDALTVVCLLLFMGAMGKSAQFLLHTWLPDAMEGPTPVSALIHAATMVTAGVFMVARLSPVFEYAPG
ALQVVMFFGATTAFFAATVGLVQNDIKRVIAYSTCSQLGYMFVALGAGAYSVGIFHLFTHAFFKALLFLGAGSV
IHAMHHEQDIRRMGGLAPHIKFTYAMMVIGTLALTGFGIPFLH-----
IGFAGYHSKDAIIEVAYAAHGAMGPYAFFMTVIAALMTSFYSWRLIFMTFHGQPRW-AGAH-
GHHDDHAHADAAAHGHADPHAHAHGAK-DDHHHDDHGHGHAHQP--------------------
HESPLVMLIPLGALAVGAVFSGVVFMKKFTDE-AGIKG--FF--
RDSIFMRPENHIIHEFHSVPGWVPWTPFVMMAIGFALAWYMYIKR----------------------------
----------------------------
PDVPVQLAREHQGLYQFLLNKWYFDELYDRIFVRPARWLGRFFWKKGGDGWLIDGFGPDGIA-R-
VVDITRGVVRLQTGYVYHYAFAMLIGVAALATWFMFSGGVG------
>WP_136964045_NuoL_Phreatobacter_stygius
MY-------QAIVFLPLVGFLIAGLFG----RAIGARGSEIVTTSLLGISALLSWVAFVQVGF-
GGQTASIELFP---WISAGALEVSWTLRIDTLTAVMLVVVNSVSFLVHLYSIGYMNEDPH-------
RPRFFAYLSLFTFAMLMLVTADDLLQMFFGWEGVGLASYLLIGFWYQKPSANAAAMKAFIVNRVGDFGFALGIF
GLYKLTGSIEL--DTIFGAAQGLVGKK---
MEFLGYQLDALTVVCLLLFMGAMGKSAQFLLHTWLPDAMEGPTPVSALIHAATMVTAGVFMVARLSPVFEYSPT
ALSVVMFFGATTAFFAATVGLVQNDIKRVIAYSTCSQLGYMFVALGAGAYSVGIFHLFTHAFFKALLFLGAGSV
IHAMHHEQDMRNMGGLAPHIKVTYAMMLIGTLALTGFGIPFLH-----
VGFAGFHSKDAIIEVAYASHNVMGPYAFALTVVAAAMTSFYSWRLIFMTFHGKPRW-ANGH-
GAHGHDAHDEEEHGIGH----------------EPGNGHDHQP--------------------
HESPAVMLIPLVLLAIGAVFAGVVFAYQFADH-HGVEK--FF--
RESLFFRPDNHIMDEFHHVPGWVPWTPFVMMAFGFVLAWYMYIRH----------------------------
----------------------------
PEMPGQLARRHPALYQFLLHKWYFDELYDRIFVRPARALGRFFWKQGGDGWLIDGFGPDGVA-R-
VVDVTRGVVRIQTGYVYHYAFAMLIGVAALATWFMFSGGLH------
>KKB10330_NuoL_Devosia_chinhatensis
MI-------QAIVFLPLIGALIAGLLG----
RQIGHRPAEFITTGLLSVAAVLSWVVFLPVAFVGDGHAAVLKVEIMRWIQVGDMDLRWTLRVDTLTAIMLVVVN
TVSALVHVYSIGYMNEDPH-------
RSRFFAYLSLFTFAMLMLVTADNFLQMFFGWEGVGLASYLLIGFWYTRPSATAAAMKAFVVNRVGDFGFALGIF
GAFMVLGHIDF--
DGAFQAADEFATTGLPVIQFLGWQLDAMTVICLLLFMGAMGKSAQFLLHTWLPDAMEGPTPVSALIHAATMVTA
```

456 GVFMVARLSPLFETSPVALTVVIVVGAITAFFAATVGLVQNDIKRVIAYSTCSQLGYMFVALGVGAYSAGVFHL
FTHAFFKALLFLGAGSVIHAMHHEQDMRNMGGLRKKIPITYAMMMIGTLALTGVGIPGTN-----
FGFAGFFSKDAIIESAYAFGGNAGTMAFWLLVIAALFTSFYSWRLVHLTFHGSPRDAQHHGEGPHPDPAHSALE
SHDEPIDDSNADDHGHHGHHHGSAYDNA-------------------------
HESPNVMLVPLYVLSVGAVLAGVVFYGMFFHD-VEHIHE-FF--AGSIFVDHQ--
IIEDAHHVPTWVKWSATIAMIAGFVAAWFMYIRR----------------------------------------
-------------------PETPAKLAASNPGLYKFLLNKWYFDELYNTIFVRPALWIGNAIWKG-
FDDWLVDGKITEGLG-RRVQNVTSWVVKLQSGYLYHYAFAMLIGIAALLTWAITAGGLL------
457 >WP_108397312_NuoL_Devosia_submarina
458 MI-------QAIVFLPLIGALVAGLLG----
RTIGHRPSEYITTGLLIISAVLSWVVFLPVAFAGVGQAAAVKVEVMRWIQVGDMDVRWILRVDTLTAIMLVVVN
TVSSLVHVYSIGYMAEDPH-------
RSRFFAYLSLFTFAMLMLVTADNFVQMFFGWEGVGLASYLLIGFWYTRPSANAAAMKAFVVNRVGDFGFALGIF
GSFMLLGTVDF--DTAFSNLPAVSNFT---
MRFLGWDVHALTVICLLLFMGAMGKSAQFLLHTWLPDAMEGPTPVSALIHAATMVTAGVFMVARLSPMFETSPT
ALTVVIVIGAITAFFAATIGLVQNDIKRVIAYSTCSQLGYMFVALGTGAYSAGVFHLFTHAFFKALLFLGAGSV
IHAMHHEQDMRNMGGLGKKIPITYAMMLIGTLALTGVGIPGTF-----
LGFAGFFSKDSIIEAAYAYGGNVGSLAFWLLVIAAVFTSFYSWRLIHLTFHGSPRDAQHA-
HTAHPDVAHAAAETHHEPIDDSNAHDHGHHDHAHHGSAHDNA------------------------
HESPNVMLVPLYVLAVGAVLSGAVFYSMFFES-LEHVHE-FF--AGAIYVDHE--
IIEGAHHVPLWVKWSATIAMIIGFVTAWFMYIKE----------------------------------------
-------------------PAVPGRIAAQNPGLYRFLLNKWYFDELYDRIFVRPAVWVGRAFWRG-
FDDWLINDKLTEGLG-RRVQNVTSWVVRLQSGYLYHYAFAMLIGIAALLTWAIAAGGLI------
459 >MBO0732892_NuoL_Methylocapsa_sp._Bacteria.bin_47
460 MY-------AAIVFLPLLGFLIAGPLA----WRKGDRAAELVTTGLLFISLGLSWAVFFQVAL--
GTAPASVSVLG-NWFTSGTLQIGWSLKIDSLTAVMLVVVTTVSAFVHLYSIGYMAGDPC-------
RPRFFSYLSLFTFAMLMLVTADNLVQLFFGWEGVGLASYLLIGFWYEKPSANAAAIKAFVVNRVGDFGFELGIF
LVFALTQSVGL--DQIFAAAPALAHKS---
IHVFGSDLDALTVACLLLFMGAMGKSAQFLLHTWLPDAMEGPTPVSALIHAATMVTAGVFMVARLSPLFEQAPA
ALSFVIFIGATTAFFAATVGLVQNDIKKVIAYSTCSQLGYMFVGLGVGGYSLGIFHLFTHAFFKALLFLGAGSV
IIALHHEQDMRMMGGLWRKIPFTFAMMVIGTLALTG--FPF---------
TAGYYSKDAIIEAAFESGRVGAFYGFILTTLAAGLTSFYSWRLVFLTFFGAPRWAALA-
ELPKSGLAHEQSANAGTGQAHEDHAFD-HP------------------------------------
HESALTMLIPLAVLAAGALFAGIIFSSDFIG-HGATG--FW--KALILASTN--
EVTEGHTLPSYVSIGPTALMVIGFVVALAFYILW----------------------------------------
--------------------PTIPSLLAKSLRPLYEFLLNKWYFDELYDKIFVRPAFALGRLFWK-
GGDGAIIDRFGPDGVAARVIEITGRVVKLQSGYIYHYAFAMLVGLAAVITWYTAWGIR--------
461 >WP_127143657_NuoL_Pelagibacterium_montanilacus
462 MI-------QAIVFLPLIGALVAGLFG----RTIGHKAAEVLTTSLLVIAAILSWIVFIPFFL--
GNAEAYKVEIM-RWIVTGDLDLRWILRVDTLTAIMLVVVNTVSCLVHVYSIGYMHDDPH-------
RARFFAYLSLFTFAMLMLVTADNFLQMFFGWEGVGLASYLLIGFWYKKPSANAAAMKAFVVNRVGDFGFALGIF
GTFYLVGSLGF--EDTFAALPGLADAT---
IPILGGEYHALTVVCLLLFMGAMGKSAQFLLHTWLPDAMEGPTPVSALIHAATMVTAGVFLVARLSPMFEMSET
ATLVIIYVGAITAFFAATVGLVQNDIKRVIAYSTCSQLGYMFVALGVGAYSAGVYHLFTHAFFKALLFLGAGSV
IHAMHHEQDMRNMGGIARKIPITYAMMIIGTLALTGVGIPGTM-----
FGFAGFFSKDAIIEAAYAFGGPAGQLAFWMLVIAALFTSFYSWRLIHLTFHGKTRADQHT--FDHA--------
-----------------------------------------------------
HESPRIMLVPLYILAAGAVLAGVVFYDVFFGH-AEHVEH-FF--HGAVVVDAE--

```
IIDAAHYVPFWVKASAAMAMLAGFVLAWYFYIRR----------------------------------------
-------------------PEAPRELAAEHPGLYKFLLNKWYFDEIYDFLFVRPTRWVGQALWKG-
FDDWFVDQMLVEGLG-RRVKQVTTQVVRLQSGYLYHYAFAMLIGVAALITWAIAAGGLLG-----
>WP_127071105_NuoL_Pelagibacterium_lentulum
MI-------QAIVFLPLIGALVAGFFG----RVIGHKPAEILTTALLMISALLSWAVFIPFFL--
GNAEAYQVEVM-RWINSGDLDLRWVLRVDTLTAIMLVVVNTVSSLVHLYSIGYMHEDPH-------
RARFFAYLSLFTFAMLMLVTADNFLQMFFGWEGVGLASYLLIGFWYKKPSANAAAMKAFVVNRVGDFGFALGIF
GTFFLFGTLGF--DETFAVLSGAEGMT---
IPFLGAEFDAMTVICLLLFMGAMGKSAQFLLHTWLPDAMEGPTPVSALIHAATMVTAGVFMVARLSPMFEMSET
ATLVVIYIGAITAFFAATVGLVQNDIKRVIAYSTCSQLGYMFVALGVGAYSAGIFHLFTHAFFKALLFLGAGSV
IHAMHHEQDMRNMGGIAKKVPLTYWMMIIGTLALTGVGIPGTM-----
FGFAGFFSKDAIIEAAYAFGGPAGTLSFWLLVVAALFTSFYSWRLIHLTFHGKTRADHHT--FDHA--------
-----------------------------------------------------
HESPNVMLIPLYVLAVGAVLSGVIFYDVFFGH-AEHVEH-FF--HGSIVVDAA--
IIDAAHYVPTWVKWSATVAMLAGFVGAWYFYIRS----------------------------------------
-------------------PETPKQLAAEHPHLYKFLLNKWYFDELYHHIFVRPARWLGNALWKG-
FDDWFIDQTVVEGLG-RRVKQVTAQVVKLQSGYLYHYAFAMLIGVAALITWAIAAGGLLG-----
>WP_090596279_NuoL_Pelagibacterium_luteolum
MI-------QAIVFLPLIGALVAGLFG----RVIGHKPAEILTTSMLIAAAVLSWIVFIPFFL--
GDGEAYKVTVM-EWIHSGDLQLDWVLRVDTLTAIMLVVVNTVSSLVHLYSIGYMHEDPH-------
RARFFAYLSLFTFAMLMLVTADNFVQMFFGWEGVGLASYLLIGFWYKKESARAAAMKAFVVNRVGDFGFALGIF
GAFFLLGTLDF--DATFAALPGFADAT---
ISFLWWEAHAMTVICLLLFMGAMGKSAQFLLHTWLPDAMEGPTPVSALIHAATMVTAGVFMVARLSPMFELSEA
ATLFIIYIGAITAFCAATIGLVQNDIKRVIAYSTMSQLGYMFVALGVGAYSAGVFHLFTHAFFKALLFLGAGSV
IHAMHHEQDMRNMGGIRKKVPLTYAMMLIGTLALTGVGIPGTS-----
FGFAGFFSKDAIVEAAYAFGGTAGSFSFWMLVTAALFTSFYSWRLIHLTFHGPTRADHHT--FDHA--------
-----------------------------------------------------
HESPNVMMIPLYVLAAGAVLAGVVFYDVFFGH-AEHVEH-FF--HGSIVVDAA--
IIDEAHHVPTLVKWSATIAMILGFIAAWYMYIRN----------------------------------------
-------------------PSAPKQLAEEHRGLYQFLLNKWYFDELYDRIFVRPARWVGTALWKG-
FDDWLIDQTLVEGLG-RRVRQVTGYVTRLQSGYLYHYAFAMLIGVAALITWAIASGGLLG-----
>WP_013421165_NuoL_Rhodomicrobium_vannielii
MY-------QAIVLLPLIGALCAGFFG----RSLGDKNSGFLTSALVLVSAALSWLALYQVGI--
EGQEARITLF--PWIGVGDLQTSWSLRIDTLTAVMLVVVNTVSALVHIYSVGYMSDDDR-------
QPLFFSYLSLFTFAMLSLVTADNLVQLFFGWEGVGLASYLLIGFWYQRESANAAAIKAFVVNRVGDFGFAIGIF
AIWFIFRDVNF--DTIFAAAKEHQATT---
IPFFGYQVPALDFICLFLFMGAMGKSAQFLLHTWLPDAMEGPTPVSALIHAATMVTAGVFMVARLSPLFELSPI
ALLVVTCVGAITAFFAATVGLAQNDIKRVIAYSTCSQLGYMFVALGLSAYGAAVFHLFTHAFFKALLFLGAGSV
IHALHGEQDLRKMGGMRKAIPFTFGMMVIGTLSLTG--FPF---------
TAGYFSKDMIIEVAEVSHSAVGQTVYWMLIFAAFLTSFYSWRLIFMAFYGEPKDH-HA--FEHA----------
---------------------------------------------------
HESPPVMTGPLLILAAGALFAGLIFAPYFVGA-DYAQ---FW--RSSLLFLHG---
WHEPHVADVFMKWLPTIVMAGGFGVAYWAYISS-----------------------------------------
------------------PGIPAWTVKTFKPIHAFLYNKWYFDELYDWIFVRPTRWIARVLWKY-
GDGAIIDGLGPDGVA-ASVIATTKRVVRIQTGFVYTYAFVMLIGVTVLITWFAYLYRGALFTL--
>WP_119060945_NuoL_Dichotomicrobium_thermohalophilum
MY-------TAIVLLPAISALFVGLFG----RYIGNAASGWTASTLLVISAILSWIAFVSVGF--
DGETARVTLF--EWIVSGDMVAEWSLRIDTLTAVMLVVVTTVSSLVHIYSIGYMEHDEG-------
```

```
QPRFFAYLSLFTFAMLALVTADNLLQLFFGWEGVGLASYLLIGFWYHKPSANAAAIKAFVVNRVGDFGFLLGIF
AIFASFGTISL--DAIFAAAPGKVDET---
IHFLGQDFDLLTVACLLLFMGAMGKSAQFLLHTWLPDAMEGPTPVSALIHAATMVTAGVFLVARMSPLFELAPV
ALTVVTAFGAITAFFAATVGIAQNDIKRVIAYSTCSQLGYMFVALGVGAYSVGIFHLFTHAFFKALLFLGAGSV
IHALHEEQDLRKMGGIARKIPLTWAMMLIGTLALTG--FPL---------
TAGYYSKDAVVEAAYAGANPVAMYAFWATVIAAFLTSAYSWRLMFMAFHGQPRMGDEA--LGKV----------
---------------------------------------------------
HESPPVMTGPLIALAIGALGAGFAFKTFFIGD-GYEA---FW--
GASLFTAEGNTILYDLHHVPGWVVASPSVAMLIGLAVAWLSYIAF-----------------------------
------------------------------
PAVPGVLTRAFKPIHLFLLNKWYFDELYDWIFVRPYKWLSRQLWKV-GDGAIIDGLGPDGIS-
ARVTDITARVVRLQTGFVYHYAFIMLVGVALIISYFLYLILSRGGF---
>WP_161140169_NuoL_Propylenella_binzhouense
MY-------SAIVFLPLFGAIIAGLIGDAAHGGHGPGAAEIVTSAFLVICAVLSWIAFFRVGA--
NTGEAFTVPVL-RWIESGALAAGWALRVDTLTAVMLVVVTTVSALVHVYSIGYMSHDPH-------
RPRFFAYLSLFTFAMLMLVTADNLVQMFFGWEGVGLASYLLIGFWYHRPSANAAAMKAFIVNRVGDFGALGVF
GVFVLFGSVEF--
SSIFANTQAVAEGGLRIFGSVFTGEGALTVVCLLLFMGAMGKSAQVPLHTWLPDAMEGPTPVSALIHAATMVTA
GVFMVARLSPLFEHAPDALIVVTVVGAITAFFAATVGLVQNDIKRVIAYSTCSQLGYMFVALGLGAYSVAIFHL
FTHAFFKALLFLGAGSVIHAVSDEQDMRRMGGLRRVIPVTYAMMLVGTLALTG--FPF---------
TAGYYSKDAIIEAAYAGHTTAAGLAFVLTVVAAAFTSFYSWRLVFMTFHGKPRASAEV--MEHA----------
---------------------------------------------------
HESASVMLIPLYVLAAGALLAGIAFSGLFIGE-AQEG---FW--
KGALFFGPENEILEEMHHAPFWVGISPFVAMVLGFLVALWFYILS-----------------------------
------------------------------
PDTPRRIGEQHEGLYRFLLNKWYFDELYDFLFVRPVMALGRFLWKR-GDGWLIDGFGPDGVS-
ARVLDVTRNVVKLQTGYVYHYAFAMLIGVAILVTWMMFAG---------
>WP_026380115_NuoL_Afifella_pfennigii
MY-------TAIVFLPLLGAIIAGLIGEHHPAAAGSRAAEMITSSLLVISAVLSWIAFFDVAL--
GERQGFSVSVL-TWVQSGDLAADWTLRIDALTAVMLVVVTTVSALVHIYSIGYMSHDPH-------
RPRFFAYLSLFTFAMLALVTADNLLQMFFGWEGVGLASYLLIGFWYQRPSANAAAIKAFIVNRIGDFGFLLGIF
GLFVLFGSISF--
DTIFANAAAVARGEVEVFGTVWSGEAALTVVCLLLFLGAMGKSAQLPLHTWLPDAMEGPTPVSALIHAATMVTA
GVFMVARMSPVFEFSVDALTVVTFIGAATAFFAATVGLVQNDIKRVIAYSTCSQLGYMFVALGIGAYSVAIFHL
FTHAFFKALLFLGAGSVIHAVSDEQDMRKMGGLWRKIPLTYALMLVGTLALTG--FPM---------
TAGFFSKDAIVEAAFVGHNPMASIAFVLLVVAAAFTSFYSWRLIFMTFHGKPRASSEV--MSHV----------
---------------------------------------------------
HESPQVMLMPLYILAAGALFGGLLFAAAFIDP-ERSVE--FW--
KGSIFTAPGNDILEELHHVPAWVKASPIIAMALGFLVSLFFYVLR-----------------------------
------------------------------
PETPKALAQRMGSLYRFLLNKWYFDELFDAMFVRPAFALGRFLWRR-GDEGTIDRLGPDGIS-
ARVVDITGRVVKLQSGYLYHYAFVMLIGVAVLVTWMMFAG---------
>WP_099555500_NuoL_Hartmannibacter_diazotrophicus
MY-------SLIVFLPLVGFLIAGAIGHHHDAAPGSRPAELITTCLMGVVAILSWVAFFQVAL--
GESETLTVPVL-SWITSGTLSVDWAFRIDTLTAVMLVVVNTVSFLVHLYSMGYMSHDPA-------
RPRFFAYLSLFTFAMLMLVTADNLLQMFFGWEGVGLASYLLIGFWYEKPSACAAAMKAFIVNRVGDFGFLLGIF
GIYLFFHTISF--DAIFANAHSLEGAS---
VTFLGSEWDAATIICLLLFMGAMGKSAQFLLHTWLPDAMEGPTPVSALIHAATMVTAGVFMVARMSPVFEMSET
```

```
ALTVVTFIGATTAFFAATVGLVQNDIKRVIAYSTCSQLGYMFAALGVGAYPIAVFHLFTHAFFKALLFLGAGSV
IHAVDGEQDMRKMGGLRTKVPVTFWMMTIGTLALTG--FPF---------
TAGYYSKDAIIEATYVGTNAFSGYAYGMLVIAAAFTSFYSWRLAFMTFFGKPRASAEV--MKHA----------
---------------------------------------------------
HESPQVMLVPLYVLAAGALLAGILFHGSFAGE-AMSELTEFW--
KGAVALPEGEHVLEAMHHVPFLAAFAPTIAMVLGFLVALHFYILN-----------------------------
------------------------------
PEAPKQLAKRHDALYQFLLNKWYFDELYEILFVRTAKWIGTKLWKI-GDGTIIDGFGPNGVA-
ARVMDVTDRVVRLQTGYLYHYAFAMLIGVAALITWAMFYA---------
>WP_132806041_NuoL_Tepidamorphus_gemmatus
MY-------SAIVFLPLIGAILAGLIGDHHEAAEGSRAAELITTGCLIVSAILSWVSFVDVGI--
NGNDARVQVL--PWVVSGSLTAEWAFRIDTLTVVMLVVVTTVSALVHMYSIGYMSHDPH-------
RPRFFAYLSLFTFAMLMLVTADNFLQMFFGWEGVGLASYLLIGFWYQRPSANAAAIKAFVVNRVGDFGFALGIF
GVFMIYGSISF--DAVFANAADATGRT---
IHIFGADIDALTVMCLLLFMGAMGKSAQFLLHTWLPDAMEGPTPVSALIHAATMVTAGVFMVARLSPMFELSET
ALTVVTFVGATTAFFAATVGLVQNDIKRVIAYSTCSQLGYMFAALGVGAYSIGIFHLFTHAFFKALLFLGSGSV
IHAMSDEQDMRRMGGLRKLIPFTYWMMIIGTLALTG--FPL---------
TAGWVSKDAIIEATYVGHNPMAGYAFGLTTFAALLTAFYSWRLIFMTFHGKSRAPSDV--LHHV----------
---------------------------------------------------
HESPPVMLVPLAVLAVGALGAGLAFKSLFMGE-GYEA---FW--
RGALYLSEENHILHDMHEVPALIVWTPTVAMLLGLGLAWYFYVVS-----------------------------
------------------------------
PATPGALARQFDVLYRFLLNKWYFDEIYDFLFVRPTKWVGRFLWKK-GDGWLIDGFGPDGIS-
ARVLDVTRKVVRLQTGYLYHYAFAMLIGVTALITWYMFAGGVR------
>WP_115689547_NuoL_Pseudolabrys_taiwanensis
MY-------QAIVFLPLLGAILAGLIGPVEPAAVGSRTAELITTTLLMISMVLSWIALVQVGF--
GGQDIREPLM--PWIVAGDLKVDWVLRIDTLTAVMLVVVNSVSAFVHLYSMGYMHEEPY-------
RPRFFAYLSLFTFAMLMLVTSDNLVQMFFGWEGVGLASYLLIGFWYQKPEACAAAIKAFVVNRVGDFGFALGIF
SIFYLTGAIDF--DTAFAQAPSLVGKT---
ILFFGWHVDALTLTCLLLFMGAMGKSAQFLLHTWLPDAMEGPTPVSALIHAATMVTAGVFMVARLSPLFELSPN
AQTFVTFIGATTAIFAATIGLVQNDIKRIVAYSTCSQLGYMFVAMGVGGYSIGMFHLFTHAFFKALLFLGSGSV
IHAMHHEQDIRNMGGLKDRIPFTYAVMIVGTLALTG--FPL---------
TAGYFSKDAIIEAAYVGKNPMALYAFACVVVAALLTSFYSWRLVFKTFHGEPHDR-QH--WKDA----------
---------------------------------------------------
HESPLVMLIPLAFLAIGSFAAGFPFKEIFAGH-GVEH---FF--
RGSLTFGKDNHILEEMHHIPLGITTLPTVMMAVGFVIAWVFYIKR-----------------------------
------------------------------
PDIPVELARQHRPLYLFLLNKWYFDELYDVIFVRPVKWLGRTLWKR-GDGWLIDGFGPDGVS-
ARVLDITRGVVRLQTGYLYHYAFAMLIGAALLITWFMFSVH--------
>WP_029040010_NuoL_Cucumibacter_marinus
MQ-------AFLFQVIVLAPLLGALVAGLLGRSIGHRPAEIITSSLLVLAGLCSWIVFLGILF--
AGGGAEHGAEAASWVHVGDLSLDWMIRVDTLTGVMLVVVTTVSALVHIYSIGYMHEDPH-------
RARFFAYLSLFTFAMLMLVTADNFLQLFFGWEGVGLASYLLIGFWYKKPSANAAAMKAFIVNRVGDFGFALGIF
GAFMMLGSINF--DEAFAQVSHMEGVT---
IDFLWGQWDAMTVICLLLFLGAMGKSAQFLLHTWLPDAMEGPTPVSALIHAATMVTAGVFMVARLSPMFEASEV
ALTVVIYVGAITAFFAATVGLVQNDIKRVIAYSTCSQLGYMFVALGLGAYSAGIFHLFTHAFFKALLFLGAGSV
IHAMHHEQDMRKMGGLRTKIPLTYAMMMIGTLALTGVGIPGTP-----
IGFAGFFSKDAIIESAAGAHTAAGDFAFWMLVIAAVFTSFYSWRLVHLTFHGKTRASRET--YSHA--------
```

```
-----------------------------------------------------
HESPNVMMIPLYVLAAGAVLSGVLFHEVFFAN-AEHVTA-FF--RNAIVVDEH--
VIEEAHHVALWVKWSPAIAMIIGFLVAWQFYIRR----------------------------------------
-------------------PDLPKALAAEHSGLYQFLLNKWYFDELYDFIFVRPAKWLGRALWRG-
FDDWLIDGIIVGGLA-ARVKDVTDRVVRLQSGYLYHYAFAMLIGVAALITWAIATGGLL------
>WP_026790169_NuoL_Pleomorphomonas_oryzae
MY-------SLIVFLPLIGFLIAGAIGAVEPAAPGSRIAELVTTGLLFVSAVFSWVAFISVGF--
SEHETQSIQVL-SWIHSGAMQVDWAFRVDTLTVVMLVVVTTVSSLVHLYSIGYMAEDPD-------
RPRFFAYLSLFTFAMLMLVTADNLVQMFFGWEGVGLASYLLIGFWYKKPSASAAAIKAFVVNRVGDFGFMLGIF
GIFMMFGTISL--TDIFANVATVEGKT---
IHFLWGNWDAITVICLLLFMGACGKSAQFLLHTWLPDAMEGPTPVSALIHAATMVTAGVFMVARLSPLFDLSPT
ALTVVTLVGAVTAFFAATVGLVQNDIKRVIAYSTCSQLGYMFVALGVGAYGAGIFHLFTHAFFKALLFLGAGSV
IMASHHEQDMRSMGGLRNKIPLTFWAMTLGTLAITGVGIPGTE-----
IGFAGFLSKDAIIESAFAGHNPVSGFAAVLLVIAAGFTSFYSWRLVFLTFFGASRAPKDV--YDHA--------
-----------------------------------------------------
HESPLVVLIPLGVLSLGAVLAGMVFYGHYVGE-EQAE---FW--
KGSLFYGPDNHILHAMHDVSFLVKASPFLAMLVGLAIAYWFYILN-----------------------------
------------------------------
PALPKRLAAALPGLYKFLLNKWYFDELYDLIFVRPAFAIGRLFWKT-GDGAIIDGLGPNGIA-
ARVQDVTARVVRLQTGYVYHYAFAMVIGVAALITWAMFAGGQ-------
>WP_181701862_NuoL_Chthonobacter_albigriseus
MY-------SLIVFLPLIGFLIAGAIGAAEPAAPGSRISELVTTGLMFVSAFLSWISFISVGF--
GEAETLRVPVL-EWMTSGALVVDWSFRIDTLTAMMLVVVTTVSSLVHLYSIGYMHEDPD-------
RPRFFGYLSLFTFAMLMLVTSDNLVQMFFGWEGVGLASYLLIGFWYKKPSAVAAAIKAFVVNRVGDFGFALGIF
GVFVMFGSISF--DAIFANTEAVAGKT---
MVFLGHEFDAMTVICLLLFMGAMGKSAQFLLHTWLPDAMEGPTPVSALIHAATMVTAGVFMVARLSPLFELSHT
ALTVVTLVGATTAFFAATVGLVQNDIKRVIAYSTCSQLGYMFVALGVGAYGAGVFHLFTHAFFKALLFLGAGSV
IHAMHHEQDMRNMGGLRKKIPVTFWVMVIGTLAITGVGIPLTH-----
IGFAGFLSKDAIIEAAYAGHNAFSGYAFGLTVLAAAFTSFYSWRLIFLTFFGETRADHHT--FDHA--------
-----------------------------------------------------
HESPRVMLVPLAVLSVGAVLAGMVFYDAFVGE-AQAE---FW--
KGSVFYGAENHILHAMHEVPAWVKASPFFAMLAGLGVAYWFYIAN-----------------------------
------------------------------
PEMPKQLAARHDALYAFLLNKWYFDELYDILFVRSAKAIGRFLWKT-GDGTVIDGLGPNGIA-
ARVQDVSARVVRLQTGYVYHYAFAMLIGVAALITWAMFGGTH-------
>WP_131310004_NuoL_Siculibacillus_lacustris
MF-------TAIVFLPLLGFLIAAAMPADHREAPFARPAELVTSGFLLIAALLSWVAFVQIGL--
AEHAATVKIAVLPWIHTAGLNVDWSLRIDTLTAVMLVVVTTVSALVHVYSIGYMSHDPD-------
RPRFFAYLSLFTFAMLMLVTADNLVQMFFGWEGVGLASYLLIGFWFQKPSANAAAIKAFVVNRVGDFGFILGIF
GVFWLFGSIDL--DTIFASVGTMKGKE---
IVFLGGHVDAMTTLCLLLFLGAMGKSAQFLLHTWLPDAMEGPTPVSALIHAATMVTAGVFMVARLSPLFEQAPA
ALAVVTLVGAVTAFFAATVGLVQNDIKRVIAYSTCSQLGYMFVALGVGAYGPGIFHLFTHAFFKALLFLGAGSV
IHAVSGEQDMRKMGGLGPHIKLTYALMIIGTLALTGVGLPMTE-----
IGFAGYNSKDAIIEAAYVGDNIFASLAFGLLVIAALFTSFYSWRLIFLTFWGAPRAPKDV--MDHV--------
-----------------------------------------------------
HESPSVMTVPLMVLALGAVIAGMVFSSVFIGH-GYGE---FW--
KGSIFTSGGNHVLEHMHHVPVAVKFAPFVAMLLGLGTAYVFYVAK-----------------------------
------------------------------
```

488 PELPGQLAASQPGLYRFLLNKWYFDELYDFLFVKPAKAFGRFLWKE-GDGRVIDGFGPDGVS-
ARVVQITGRVVALQSGYLYHYAFAMLIGVALLATMVMFGGVR-------
489 >WP_075214650_NuoL_Mongoliimonas_terrestris
490 MY-------SLIVFLPLVGFLIAGAIGAAEPAAPGSRIAELVTTGFLIVSCLLSWVAFLTVGF--
GDGETFRVPVA-AWMTSGALVVDWSFRIDTLTVMMLIVVTTVSSLVHAYSIGYMSEDPG-------
RPRFFAYLSLFTFAMLMLVTSDNLVQMFFGWEGVGLASYLLIGFWYKKPSASAAAIKAFVVNRVGDFGFALGIF
GVFAVFGSVGF--DAIFANTEAVAEQS---
MIFLGAEWHALEIITFLLFIGAMGKSAQFLLHTWLPDAMEGPTPVSALIHAATMVTAGVFMVARLSPLFEHAHG
TLTFVTFIGATTAFFAATVGLVQNDIKRVIAYSTCSQLGYMFVALGVGAYGAGVFHLFTHAFFKALLFLGAGSV
IHAMHHEQDMRKMGGLRSKIPVTYWTMLIGTLAITGVGIPLTH-----
IGFAGFLSKDAIIESAYAGHNSFAGYAFMLTLIAAAMTSFYSWRLIFLTFFGQTRADHHT--FEHA--------
-----------------------------------------------------
HESPRVMLIPLIVLSVGAVLAGMVFYEAFVGE-AQGE---FW--
KGALLYGPENHILHDMHEVPAWVKASPFFAMLFGLGIAYWFYIAD-----------------------------
------------------------------
PEQPKRLAARHDALYQFLLNKWYFDELYDLLFVRSAKALGRFFWKA-GDGMVIDGLGPNGVA-
ARVQDVSARVVRLQTGYVYHYAFAMLIGVAALITWAMFAGGIN------
491 >WP_083255387_NuoL_Methylobrevis_pamukkalensis
492 MY-------SLIVFLPLVGFLIAGAIGDHHEEAPGSRPAELVTTGLMAVTALLSWVAFFSVAF--
GEGEAFTVPVL-SWIHSGALATEWAFRIDTLTVVMLVVVNTVSFLVHLYSMGYMSHDPA-------
RPRFFAYLSLFTFAMLTLVTSDNLLQMFFGWEGVGLASYLLIGFWYQKPSANAAAMKAFVVNRVGDFGFLLGLF
GVYVLFNSISF--DAIFANVGSLEGAQ---
LTFLGSSWDAATIVCLLLFMGAMGKSAQFLLHTWLPDAMEGPTPVSALIHAATMVTAGVFMVARMSPVFEMSHT
ALTVVTFIGATTAFFAATVGLVQNDIKRVIAYSTCSQLGYMFVALGVGAYPIAVFHLFTHAFFKALLFLGAGSV
IHAVSGEQDMRKMGGLRTVIPVTFWMMTIGTLALTG--FPF---------
TAGFYSKDGIIEAAFVGENAFAGYAFGMLVIAAVFTSFYSWRLAFLTFFGKPRASAEV--MKHA----------
---------------------------------------------------
HESPQVMLIPLYVLAVGALFAGVAFSGAFMGE-GVEE---FW-
KQSLYLPHGEHSILEAIHHVSFWVKISPFIAMLIGFAVAVYFYIVN----------------------------
-------------------------------
PEAPKQLAARHDALYQFLLNKWYFDELYDILFVRSAKWIGTKLWKV-GDGRIIDGLGPDGIA-
ARVQDVTTRVVRLQTGYLYHYAFAMLIGVALLVTWAMFYAGGM------
493 >WP_023432137_NuoL_Lutibaculum_baratangense
494 MY-------SAIVFLPLIGAIIAGAIGDGHHDHHGPGAAEYITTGALILSAILSWVAFFTVGL--
GEEPVHVQVM--PWVVSGDLAFDWAFRIDTLTVVMLVVVTTVSSLVHVYSIGYMSHDAH-------
RPRFFAYLSLFTFAMLMLVTADNLVQLFFGWEGVGLASYLLIGFWYTRPSANAAAIKAFVVNRVGDFGFSLGIF
AVFLVYGSVNF--EVIFANAADAEGMT---
LSFLGGEFDALTVICLLLFMGAMGKSAQFLLHTWLPDAMEGPTPVSALIHAATMVTAGVFMVTRLSPLFELAPT
ALAFVTFIGATTALFAATIGLVQNDIKRVIAYSTCSQLGYMFVAIGVGAYGAAIFHLFTHAFFKALLFLGSGSV
IHAMSDEQDMRRMGGLRKHIPLTYWMMVIGTLALTG--FPF---------
TAGFISKDAIIESAFAAHGTFAVYAFALTLFAAALTSFYSWRLIFMTFHGKPRASADV--MHHV----------
---------------------------------------------------
HESPLVMTVPLMILAVGALFSGLAFQGLFIGE-GYED---FW--
KQAIFLGSDNDILHEMHEVSALVVWSPTVAMIIGFGLAYLFYIRR-----------------------------
------------------------------
PSIPVALAKQFDLVYLFLLNKWYFDELYDLIFVKPSFWLGRVFWKK-GDGWLIDGFGPDGIS-
ARVLDVTRQFVRLQTGYLYHYAFAMLVGVTALITWYMFAGGAH------
495 >WP_096356271_NuoL_Variibacter_gotjawalensis

```
MY-------QAIVFLPLLGAILAGAIGPAEPAAVGSRAAEVITSALLVTSAVLSWITLVQVGY--
GHQEARIVLF--PWMVSGDLNVSWALRIDTLTAVMLVVVTTVSSLVHIYSIGYMHEDPH-------
RPRFMAYLSLFTFAMLMLVTSDNLAQLFFGWEGVGLASYLLIGFWFKKPTANAAAIKAFVVNRVGDFGFALGIF
ALFAVTGSINF--DQIFAEGPNLANKT---
IHFLSWDVNALTVICLLLFMGAMGKSAQFLLHTWLPDAMEGPTPVSALIHAATMVTAGVFMVARLSPLFEYSPT
ALTVVTFFGATTAFFAATIGLVQNDIKRVIAYSTCSQLGYMFVAMGVGAYSVGMFHLFTHAFFKALLFLGSGSV
IMAMHHEQDIRNMGGLRKKIPFTYWMMVIGTLALTG--FPF---------
TAGYFSKDAIIEAAYAGKNPMAIYGFMMTVIAALLTSFYSWRLIFKTFHGKPHDQHHY--DEA-----------
----------------------------------------------------
KESPLVVLVPLGILAVGSIAAGFVFKEVFIGH-HVAE---FF--
RESIKIIPENHILHDMHEVPKWVVASPFVMMVIGFLVAYQFYIRR-----------------------------
------------------------------
PDLPVGLAKQQPLLYNFLLNKWYFDELYNYIFVKPALWLGRLFWKR-GDGTIIDGFGPDGVS-
ARVADVTRGVVRLQTGYLYHYAFAMLIGVAAFITWFLFSGTTGGAH---
>WP_073629719_NuoL_Pseudoxanthobacter_soli
MY-------HLIVFLPLLGFLIAGAIGDHHAAAPGSRAAEIITTSLLLVSAILSWVAFFSVGV--
GETQVPTVDLL-TWVTSGSLSFNWALRVDTLTVVMLVVVNTVSMLVHLYSMGYMAHDEH-------
RPRFFAYLSLFTFAMLMLVTSDNLMQMFFGWEGVGLASYLLIGFWFQKPSACAAAIKAFVVNRVGDFGFLLGIF
ALVYMTGVLQF--DQVFAAAPSLAEKS---
IHFLGGDYNAVTVICLLLFMGAMGKSAQFLLHTWLPDAMEGPTPVSALIHAATMVTAGVFMVARMSPLFSLSAD
ALTFVTLIGATTAFFAATIGCVQNDIKRVIAYSTCSQLGYMFAALGVGAYNVAVFHLFTHAFFKALLFLSAGSV
IHAVGGEQDMRKMGGLRKTVPFTFWAMLFGGLALTG--FPL---------
TAGYFSKDAIIEATFVGTNSMHVYAFWMTTIAALLTAFYTWRQMFMTFFGKPRASAEV--MAHA----------
----------------------------------------------------
HESPLVMLIPLGVLSLGALFGGIVFHNLFIGE-SFEE---FW-
KHALDMESVEHVLHAMHTEVPGWARWAPTVAMLIGLFSAWYAYIRV----------------------------
-------------------------------
PTLPAALAARNKGLYEFLLNKWYFDELYDFLFVRPAKRLGSFLWKR-GDGWLIDGFGPNGVS-
ARVVDITNRVVRLQTGYIYHYAFVMLIGIALLVTWSMFHGGGNP-----
>WP_01147286_NuoL_Rhodopseudomonas_palustris
MI-------QAIVFLPLIGALLAAAIGPVETPAVGSRAAELITTGLLLVAAGLSWLALVDVGF--
MHHDARIPLF--PWINSGDLQVAWSLRVDTLTAVMLVVVNTVSSLVHLYSIGYMNEDPH-------
RPRFFAYLSLFTFAMLMLVTADNLVQMFFGWEGVGLASYLLIGFWYQKPEANAAAIKAFVVNRIGDFGFLLGIF
AIFMLVGSIDF--ETIFAAAPGLTGKT---
INFFGWHADALTLTCLFLFMGAMGKSAQFLLHTWLPDAMEGPTPVSALIHAATMVTAGVFMVARLSPLFELAPD
AQNVVIFFGATTAFFAATIGLVQNDIKRIVAYSTCSQLGYMFVAMGTGAYSVGMFHLFTHAFFKALLFLGSGSV
IHAMHHEQDIRKMGGLKDKIPFTYIVMVIGTLALTG--FPL---------
TAGYFSKDAIIEAAFASHKSVAFYGFLMTVIAAGLTSFYSWRLIFKTFHGTPHDQ-KN--YDAA----------
----------------------------------------------------
HESPYWMLIPLVVLAAGSILAGFPFKEVFAGH-GVGE---FF--RESLKMNPHII--
DEMHHIPVAVAYLPTAMMALGFVVSWMFYIRK------------------------------------------
-----------------PYLPVELANQHQLLYKFLLNKWYFDELYDLIFVRPAKWIGRFLWKA-
GDGTIIDGLGPDGIS-ARVLDITRGAVRLQTGYLYHYAFAMLVGVAGLITWFMLGGGGR------
>WP_143577931_NuoL_Tardiphaga_sp._vice278
MI-------QALVFLPLIGAILAGLIGVAEPAAAGSRAAELITSTLLVVSAALSWMTLVDVGF--
LHHDVRVVLM--PWITSGDLQLSWTLRVDTLTAVMLVVVTTVSALVHIYSLGYMEDDPN-------
RPRFMGYLSLFTFMMLMLVTADNLLQLFFGWEGVGLASYLLIGFWYQKPSANAAAIKAFVVNRVGDFGFLLGIF
AIFMMVGSTDF--DTIFAALPGLTGKT---
```

```
INFFGWQADALTLIGLLLFMGAMGKSAQFLLHTWLPDAMEGPTPVSALIHAATMVTAGVFLVARMSPLFELAPN
AQAVIMLVGATTAFFAATIGLVQNDIKRIVAYSTCSQLGYMFVALGAGGYSIAMFHLFTHAFFKALLFLGSGSV
IYAMHHEQDIRKMGGLWRKIPFTYAMMVIGTLALTG--FPL---------
TAGYFSKDAIIEAAYVSTNPFALYGFLMTIVAAGLTSFYSWRLIFKTFHGAPHDQ-KH--YEAA----------
---------------------------------------------------
HEAPLVMLVPLGVLAAGAILAGFPFKELFAGH-GVEE---FF--RDSLKNHPHLL--
DEMHHIPAMIANLPTVMMAAGFLVSYLFYIRR------------------------------------------
-----------------PYLPVELAGQHPLLYKFLLNKWYFDELYDLIFVRPAKWLGRFLWKV-
GDGKIIDGLGPDGIS-ARVLDITRGAVRIQTGYLYHYAFAMLIGVAGLITWFIFGLGGH------
>WP_068730882_NuoL_Tardiphaga_robiniae
MI-------QALVFLPLIGALIAGFIGVAEPAALGSRAAELITAGLLVVSAVLSWMTFVDVAF-MHHDARIVLL
---NWINSGDLQLSWTLRVDTLTAVMLVVVTTVSSLVHIYSLGYMEDDPN-------
RPRFMAYLSLFTFMMLMLVTADNLLQLFFGWEGVGLASYLLIGFWFKKPSANAAAIKAFVVNRVGDFGFLLGIF
ALFMLVGSTDL--DTIFASAPSLTGKT---
INFFGWHADALTLTCLLLFVGAMGKSAQFLLHTWLPDAMEGPTPVSALIHAATMVTAGVFMVARMSPLFELAPN
AQAVVMLVGATTAFFAATIGLVQNDIKRIVAYSTCSQLGYMFVALGTGAYSVAMFHLFTHAFFKALLFLGSGSV
IYAMHHEQDIRKMGGLWNKIPFTYAMMTIGTLALTG--FPF---------
TAGYFSKDAIIEAAYASHNPFAFYGFLMTVVAALLTSFYSWRLVFKTFHGEPHDK-KH--YDAA----------
---------------------------------------------------
HEAPLWMLVPLAVLGIGSFAAGFPFKELFAGH-HVGE---FF--RESLKMNPHII--
EEMHHIPGWVPYVPTVMMVLGFLISYLFYVRK------------------------------------------
-----------------PYLPVELANQHPLLYKFLLNKWYFDELYEIIFVRPAKWIGHFLWKK-
GDGMIIDGLGPDGVS-ARVLNITRGVVKLQTGYLYHYAFAMLIGVAGLITWFIFGLGGQ------
>OJY10283_NuoL_Rhizobiales_bacterium_62-47
MI-------QAIVFLPLLGAILAGLIGPAEPPAAGTGAAELITTALLFVSAALSWITFVNVGF--
LKQDLLVHVL--PWISSGDLQIAWSLRVDTLTAVMLVVVTTVSSLVHLYSVGYMDEDPH-------
RPRFFAYLSLFTFAMLMLVTSDNLVQLFFGWEGVGLASYLLIGFWYQKPSANAAAIKAFVVNRVGDFGFSLGIF
AVFMLIGSTDF--ETIFHAAPGLTGKT---
INFLGWHADALTLTCLLLFMGAMGKSAQFLLHTWLPDAMEGPTPVSALIHAATMVTAGVFMVARLSPLFELAPN
AQAFVMLIGGTTAFFAATIGLVQNDIKRIVAYSTCSQLGYMFVAMGAGAYSVGMFHLFTHAFFKALLFLGSGSV
IYAMHHEQDIRNMGGLKDKIPFTYVAMVIGTLALTG--FPL---------
TAGYFSKDAIIESAFVAQNPFAFYGFLMTVIAACLTSFYSWRLIFKTFHGRPHDQ-HH--YEAA----------
---------------------------------------------------
HEAPLWMLVPLGVLAAGSILAGFPFKELFAGH-GIEE---FF--RESVKMHPHII--
EEMHHIPATIAYLPTVMMVVGFVVAWLFYIQR------------------------------------------
-----------------PYLPVELASQHPLLYKFLLNKWYFDELYDVIFVRPAKWLGRFLWKK-
GDGFVIDGFGPDGVS-ARVLDVTRNVVRLQTGYLYHYAFVMLIGVAGLITWFMFGLGGQ------
>WP_184082102_NuoL_Afipia_massiliensis
MI-------QAIVFLPLIGALIAGAVGPVEPPAAGSRTAEVVTTALLLISAALSWMALVDVGF--
LHHDARVQLF--TWIGSGDLVVNWALRVDTLTAVMLVVVTSVSSLVHLYSIGYMHEDPN-------
RPRFMAYLSLFTFAMLMLVTADNLVQLFFGWEGVGLASYLLIGFWYQKPSANAAAIKAFVVNRVGDFGFALGIF
AVFMLVGSTDF--DTIFAAAPGLTGKT---
IHFFGWDVDALTLTCLLLFMGAMGKSAQFLLHTWLPDAMEGPTPVSALIHAATMVTAGVFMVARLSPLFELAPN
AQAAVMLIGATTALFAATIGLVQNDIKRIVAYSTCSQLGYMFVAMGAGAYSVGMFHLFTHAFFKALLFLGAGSV
IHAMHHEQDIRAMGGLKDKIPFTYITMVVGTLALTG--FPL---------
TAGYFSKDAIIESAFVAQNPFALYGFLCTVIAAGLTSFYSWRLIFKTFHGEPHDK-EH--YEAA----------
---------------------------------------------------
HESPYTMLIPLGVLAAGSILAGFPFKELFAGH-GIGE---FF--RESLKNHPKII--
```

```
DEMHHIPAMIAFLPTVMMVVGFIVSWLFYIRK------------------------------------------
-----------------PYLPVELANQHPGMYRFLLNKWYFDELYDFIFVRPTKWLGRFLWKK-
GDGAIIDGFGPDGVS-ARVLDVTRNVVRLQTGYLYHYAFAMLIGAAGLITWFIFGLGGQ------
>WP_100382102_NuoL_Afipia_broomeae
MV-------QAIVFLPLIGAVLAGLIGAVEPPAQGSRAAELITTGLLIVSAVLSWITFFDVGF-MHHDVRIALF
---QWINSGDLQVAWSLRVDTLTAVMLVVVTTVSSLVHLYSIGYMDEDPN-------
RPRFFGYLSLFTFAMLMLVTSDNLVQLFFGWEGVGLASYLLIGFWYQKPSANAAAIKAFVVNRVGDFGFALGIF
AIFALIGSTDF--ETIFAGAPGLTGKT---
IDFFGWHADALTLTCVLLFMGAMGKSAQFLLHTWLPDAMEGPTPVSALIHAATMVTAGVFMVARLSPLFELSPN
AQAVVMFFGATTAFFAATIGLVQNDIKRIVAYSTCSQLGYMFVAMGAGAYSVGMFHLFTHAFFKALLFLGSGSV
IYAMHHEQDIRKMGGLWRKIPYTFTVMCIGTLALTG--FPL---------
FAGYFSKDAIIESAYASHNPFSTYAYLLTVGAAGLTSFYSWRLIFKTFFGEPHDR-EH--YEAA----------
---------------------------------------------------
HESPIWMLIPIGVLAAGSILAGFPFKELFVGH-GAEE---FF--RDSLKMNPQIF--
EDMEHMPRLLGFMPFIMMVLGLAVAYLFYIRR------------------------------------------
-----------------PYLPEELASQQPMLYQFLLNKWYFDELYDLIFVRPAKRIGRFLWKF-
GDGYIIDGFGPDGVS-AWVLDVTRNVVKLQTGYLYHYAFAMLIGVAGLITWFMFGVGGQ------
>WP_111383984_NuoL_Rhodoplanes_piscinae
MY-------QAIVFLPLLGFLIAAAIGAAEPAAWGSRPAEVVTTTLLFVSMALSWIAFARVGF--
GGLEVREAIA--PFIFSGELKVDWALRVDTLTAVMLVVVTTVSALVHLYSIGYMVDDPG-------
RPRFFAYLSLFTFAMLMLVTADNLVQLFFGWEGVGLASYLLIGFWYKKPSANAAAIKAFVVNRVGDFGFLLGIF
AVFVVTAAVDF--DTIFAQAPALAGKP---
IPFLSWEVDALTVICLLLFMGAMGKSAQFLLHTWLPDAMEGPTPVSALIHAATMVTAGVFMVARLSPLFELAPT
ALAVVTFVGATTALFAATVGLVQNDIKRIVAYSTCSQLGYMFVAMGVGAYSVGMYHLFTHAFFKALLFLGSGSV
IVAMHHEQDIRMMGGLSRKIPFTYAMMLVGTLALTG--FPL---------
TAGYFSKDAIIEAAFASDHAMAFYGFVCTVAAAALTSFYSWRLVFKTFHGQPHDR-HH--YDHA----------
---------------------------------------------------
HESPLPMLIPLGVLAAGSILAGLPFYDAFAGN-SVAA---FF--RGSLSVTG-IL--
EEMHHVPAMIAYLPTVMMVAGFAVAWVFYIKS------------------------------------------
-----------------PGIPRALARTFKPVYLFLLNKWYFDELYDRIFVRPAMWLGRALWKG-
GDGRLIDGLGPDGVS-ARVLDVTRNVVRLQTGYLYHYAFAMLIGVAALATWFILGGLR-------
>WP_002711593_NuoL_Afipia_clevelandensis
MI-------QAIVFLPLIGALIAGAIGPVEPPAAGSRAAEVITTALLLISAALSWVTLVDVGL-LHHDARVQLF
---NWIGSGNLVVNWAFRVDTLTAVMLVVVTSVSSLVHLYSIGYMHEDPN-------
RPRFMAYLSLFTFMMLMLVTADNLVQMFFGWEGVGLASYLLIGFWFQKPSANAAAIKAFVVNRVGDFGFLIGIF
SVFLMVGSVDF--DTIFAAAPGLTGKT---
IHFFHWNIDALTLICLFLFMGAMGKSAQFLLHTWLPDAMEGPTPVSALIHAATMVTAGVFMVARLSPLFELAPN
AQAFVMLIGATTAMFAATVGLVQNDIKRIIAYSTCSQLGYMFVAMGAGAYSVGMFHLFTHAFFKALLFLGAGSV
IHAMHHEQDIRAMGGLKDRIPFTYITMVIGTLALTG--FPL---------
TAGYFSKDAIIESAFVAHNPFALYGFLCTVVAAGLTSFYSWRLIFKTFHGEPHDK-HH--YDAA----------
---------------------------------------------------
HESPYTMLIPLGVLAMGSILAGFPFKELFAGH-GVEE---FF--RESLKNHPHIL--
EEMHHIPAMIAFLPTVMMVIGFLVSWLFYIRK------------------------------------------
-----------------PYLPVELANQQPLLYNFFLNKWYFDELYEVIFVRPAKWLGRFLWKK-
GDGFIIDGFGPDGVS-ARVLDVTRNVVRLQTGYLYHYAFAMLIGAAGLITWFIFGLGGQ------
>WP_073051127_NuoL_Kaistia_soli
MY-------SAIVFLPLIGAIIAGVIADHAPAAPGSRPAEIITTGFLFVSAILSWIAFLTITA--
PTGEPVFVNVL-PWVTSGTLSFNWSFRIDTLTSVMLILVTTVSSLVHLYSWGYMAHDEH-------
```

```
RPRFFGYLSLFTFAMLMLVTSDNLVQMFFGWEGVGLASYLLIGFWYQRPSANAAAIKAFVVNRVGDFGFAVGIF
GVFVLFNSVDF--QTIFANAGGAVGTK---
MHFLSWDFDALTTVCLLLFVGAMGKSAQFLLHTWLPDAMEGPTPVSALIHAATMVTAGVFMVARMSPLFGLSET
ALTMVTFFGATTCFFAATIGLVQNDIKRVIAYSTCSQLGYMFVGLGTGAFATGIFHLLTHGFFKALLFLCAGSV
IHAVSGEQDMRKMGGLRKHLPITYWAMIAGTLAITG--FPF---------
TAGYFSKDAIIEAAYVAENPLAGYAWFMTVAAALLTSFYSWRLIFMTFHGKPR--ASAEVMAHV-----------
---------------------------------------------------
HESPKPMTIPLIVLSLGALFAGMIFENRFIGE-AYAE---FW--
KSALFTNPDNHVLHNMHEMSELVKLAPTVMMVIGFLLTYWFYIRS-----------------------------
------------------------------
PEVPKRLAEEQPLLYRFLLNKWYFDELYDFLFVRPVKWLGRFLWKK-GDGWLIDGFGPDGIS-
ARVVDVTKRVVRLETGYVYHYAFAMLIGVSALVTWMMF--AR-------
>WP_111199216_NuoL_Aestuariivirga_litoralis
MN----TVLSAIVFLPLLGALIAGLFGGHHGAYDGPKWPMYLTTALLCISAVLSWYVFA--GF--
LHEPRAEKIELLHWINSGALSANWMIRVDTLTAVMLVVVNTVSALVHIYSLGYMSHDES-------
QPRFFAYLSLFTFAMLMLVTADNLVQMFFGWEGVGLASYLLIGFWYQRDSACAAAMKAFIVNRVGDFGFALGIF
GCFLVFQTVDF--DTIFAAAPGMAGKP---
MQFLAWTPDTLTVLCLLLFMGAMGKSAQFLLHTWLPDAMEGPTPVSALIHAATMVTAGVFMVARLSPMFELAPH
AKDFVVVIGAITAFFAASVGLVQNDIKRVIAYSTCSQLGYMFVALGVGAYGAGVFHLFTHAFFKALLFLSAGSV
IHAVSGEQDMRKMGGLGPFIKITFAMMVIGNLALTG--FPY---------TSGYFSKDAIIEAAYVS---
GWGFAFWMLIVAAVFTSFYSWRLTFMTFNGQPR--ASREVMEHV------------------------------
-------------------------------HESPPVMLVPLFVLAIGALFAGFFFKDYFIGH-DYKE---FW-
-GASLANGEVMHEMHEEGMVPALVKWSPFIAMVLGFVVAWIFYIRD----------------------------
-------------------------------
TSIPKRLAEQHQPLYRFLLNKWYFDELYDVIFVRPTMWLGRFLWKK-GDGFLIDGFGPDGVS-
AVVNDVTKGVVRLQTGYLYHYAFAMMIGVAALISWFMF-GGAH------
>WP_080917694_NuoL_Pseudaminobacter_manganicus
MY-------QAIVFLPLIGFLIAGLFG----NSLGAKASEAITSGLLVIAAVLSWVAFFTVGF--
GTGEVFTVPMM-TWFHAGGVDVSWALRIDTLTVVMLVVVNTVSALVHIYSIGYMHHDPS-------
RPRFFAYLSLFTFAMLMLVTADNLLQMFFGWEGVGLASYLLIGFWFKKPSANAAAIKAFVVNRIGDFGFLLGIF
GVFVLFGTLDL--GTIFANAATYLPAE---
GAPAGDTVLNFLGYALLLFMGAMGKSAQMPLHTWLPDAMEGPTPVSALIHAATMVTAGVFMLARLSPLFELSHT
ALTIVTFIGAFTAFFAASVGLVQNDIKRVIAYSTCSQLGYMFVALGVGAYGVAIFHLFTHAFFKALLFLGSGSV
IHAVSDEQDMRNMGGLRKLIPTTYWMMVIGSLALTGVGIPLTV-----
IGTAGFFSKDAIIEASFVSHNPVAAFAFVMLVIGA--TSFYSWRLIFMTFHGKPR--ASEEVMHHV--------
-----------------------------------------------------
HESPSVMLVPLYVLAAGALFAGVLFSTYFIGE-GYEA---FW--
KAALFTLPDNHILHEMHDVPLWVSLAPIVVTIIGFLVAYKFYITS-----------------------------
------------------------------
PEMPEQVAGRHRMLYAFLLNKWYFDEIYEFLFVQPAKNIGHFLWKT-GDGKVIDGIGPDGIS-
ARVVDVTNRVVKLQTGYLYHYAFAMLIGVAALVTWMML-----------
>WP_018633546_NuoL_Neomegalonema_perideroedes
ML-------TAIVFLPLIGSLISGFFG----KRLGETLAQWIPTGLLFVSALFSWIVFAQTL---
GGEDRLVRLF--TWISSGELQTDWALRADPLTAVMLVVVTTVSALVHLYSVGYMAEDPS-------
KARFFSYLSFFTFAMLMLVTADNFLQMFFGWEGVGVASYLLIGFWHQKDSANAASMKAFIVNRVGDFGFLLGMF
GVFAVFRSLDF--DTVFAAAPQYAEST---
FHFLWMEVKVLEAICFLLFIGAMGKSAQFFLHTWLPDAMEGPTPVSALIHAATMVTAGVVMICRLSPLFEWAPG
ALAFVTFIGATTALFAATVGLVQNDIKRIIAYSTCSQLGYMFFAAGLGAYPVAMFHLFTHAFFKALLFLGAGSV
```

```
IHAMHHEQDIRKMGGLWRYIPLTFVMMVVGTLAITG--LPL---------
LSGFYSKDAIIEVAFVGKGWTAQYAFFVSLLVAAMTSFYSWRLIFKVFFGPRVWKDDPDPYPT--
PVEEAHDDHAHGDDHGHGHHELHP------------------------------------
HESPKTMLIPIGVLTVGAILAGAIFAPGFIGDGYEEFW------
RGSIFFGPNNHVMHDFHYVPLWVKFAPFMMMVTGFFIAWTLYERR-----------------------------
------------------------------
RDLPAKLARSQEALYTFFLNRWWFDQAYDFLFVKPAQALGRFLWKK-GDGATIDG-
AIDGASMKIAPAIAGLLRRAQSGYVFHYAFAMILGVVVLVGLAVV-----------
>WP_138015039_NuoL_Pelagicola_litoralis
ME-------KIILFAPLVGSLIGGFGW----RFIGEKGAQYVTTGLLFLACALSWIVFLGHD---GQTQYIHIL
---
DWIQSGTLDTAWAIRLDRLTAIMLIVVTSVSAVVHLYSFGYMDHDPQWKEGESYKPRFFAYLSFFTFAMLMLVT
SDNLVQMFFGWEGVGVASYLLIGFYYRKPSANAAAIKAFVVNRVGDFGFGLGIFGLFMLTDSVKF--
DDIFAAAPALADTQ---
LHFLWADWNAANLIAFLLFIGAMGKSAQLFLHTWLPDAMEGPTPVSALIHAATMVTAGVFLICRMSPLMEYAPE
ATAFITFLGATTAFVAATIGLVQNDIKRVIAYSTMSQLGYMFVAAGVGVYSVAMFHLFTHAFFKAMLFLGAGSV
IHAMHHEQDMRNYGALRKKIPYTFMAMMIGTLAITGVGIPLTH-----IGFAGFLSKDAVIESAWA--
GTAGGYAFWMLVIAALFTSFYSWRLMFMTFYGKAR--GDKHTHDHA----------------------------
---------------------------------HESPMVMLVPLGLLALGAVFSGMIWYGSFFGDHDKVNK--
FFG-GHGAFMAPDNHVMDEAHHAPTWVKVSPFVAMIIGFVVAMWFYIWN-------------------------
----------------------------------
PSMPKKLADAQRPLYLFLLNKWYFDELYDVIFVRPAKAIGRFLWKR-GDGNVIDG-
ALNGVAMGIVPFFTKLAGRAQSGYIFTYAFAMVIGIVVLITWMTLSGGAH------
>EAU50537_NuoL_Rhodobacterales_HTCC2255
ME-------IVLLFAPLVGSIICGFGH----RILGEKISMMVSTGALFISAILSWVVFINFHGD-GHTI-SLFR
----
FIESGSLSADWAIRIDRLTAIMLVVINTVSALVHLYSWGYMDKDPQWKQGESYKPRFFAYLSFFTFAMLMLVTS
DNLLQMFFGWEGVGVASYLLIGFYYRKPSAGAAAMKAFIVNRVGDFGFALGIFGLFYLSGSINF--
SEVLDHNAKELSKTS--
FQFLSWQLNALEVITMLLFIGAMGKSAQLMLHTWLPDAMEGPTPVSALIHAATMVTAGVFMVCRLSPLMEYATF
TPNFIVLIGATTAFFAATVGLVQNDIKRVIAYSTCSQLGYMFVAAGVGAYGAAMFHLFTHAFFKAMLFLGAGSV
IHAMHHEQDMRNYGALRKKIPYTFAAMMIGTLAITGVGIPLTH-----
YGFAGFLSKDAIIESAYGAGTGVGQYAFVMLVVAAAMTSFYSWRLIFLTFYGKAR--GNSHTHDHA--------
-----------------------------------------------------
HESPLVMILPLAVLALGAVFSGMIWYEDFFGD-HWEE---FF-
GSSLYANHDTNHVVHDAHYVANWVKASPFFAMLFGFIMALWFYIWA----------------------------
-------------------------------
PENPAKLAKQQNAVYLFFLNKWYFDELYEIIFVRPSKWLGNFLWKR-GDGSLIDG-
GINGIALGIIPLLTRKYQGMQSGYLFHYAFTMVIGVAVIVTWFALTGSN-------
>WP_011748520_NuoL_Paracoccus_denitrificans
ME-------KFVLFAPLIASLIAGLGW----RAIGEKAAQYLTTGVLFLSCLISWYLFLSFD---GVP--
RHIPVL-
DWVVTGDFHAEWAIRLDRLTAIMLIVVTTVSALVHMYSLGYMAHDDNWTHDEHYKARFFAYLSFFTFAMLMLVT
ADNLLQMFFGWEGVGVASYLLIGFYYKKASANAAAMKAFIVNRVGDFGFLLGIFGIYWLTGSVQF--
DEIFRQVPQLAQTE---
MHFLWRDWNAANLLGFLLFVGAMGKSAQLLLHTWLPDAMEGPTPVSALIHAATMVTAGVFLVCRMSPLYEFAPD
AKNFIVIIGATTAFFAATVGLVQNDIKRVIAYSTCSQLGYMFVAAGVGVYSAAMFHLLTHAFFKAMLFLGAGSV
IHAMHHEQDMRNYGGLRKKIPLTFWAMMIGTFAITGVGIPLTH-----LGFAGFLSKDAIIESAYA----
```

```
GSGYAFWLLVIAACFTSFYSWRLIFLTFYGKPR--GDHHAHDHA------------------------------
-------------------------------HESPPVMTIPLGVLAIGAVFAGMVWYGPFFGD-
HHKVTEYFHIVGGAIYMHPDNHIMDEAHHAPAWVKVSPFVAMVLGLITAWTFYIAN------------------
-----------------------------------------
PSLPRRLAAQQPALYRFLLNKWYFDEIYEFIFVRPAKWLGRVLWKG-GDGAVIDG-
TINGVAMGLIPRLTRAAVRVQSGYLFHYAFAMVLGIVGLLIWVMM-RGAH------
>WP_038147201_NuoL_Thioclava_atlantica
ME-------TIILFSPLVGAIIGGFGW----RVIGEKGAMVLTTGLLFLAAALSWIVFLTL----GDTTQ-
HIHIL-
NWISSGDLDVNWSIRMDRLTAIMLIVVTTVSALVHLYSWGYMAHDENWTENENYKARFFAYLSFFTFTMLMLVT
SDNLLQMFFGWEGVGVASYLLIGFYYKKPSANAAAMKAFIVNRVGDFGFLLGMFGLFYMTGSLKM--
DDVFAAAPALADTQ---
LHFLWTDWNAIELIGVLLFVGAMGKSAQLFLHTWLPDAMEGPTPVSALIHAATMVTAGVFLVCRMSPVYEYAPH
AKEFILIIGATTAFFAATVGLVQNDIKRVIAYSTCSQLGYMFAAAGVGVYSASMFHLLTHAFFKALLFLSAGSV
ITAMHHEQDMRNYGGLRKKIPLTFWAMIMGTLAITGVGIPFTH-----LGFAGFLSKDAIIESVWA--
GDASGYGFWMLVIAAVFTSFYSWRLVFMTFFGKPR--GDHHAHEHA----------------------------
---------------------------------HESPWVMTVPLVVLSLGAIFAGMIWYNAFFGDHAKLVK--
FFNLPAAIFMGAENDVIDAAHHAPAWVKMSPFFAMLAGLIVAWLFYIRD-------------------------
----------------------------------
VSLPRRLAEAQPSLYRFLLNKWYFDELYDRIFVRPALWLGYQFWKK-GDGQ-IDR-
AIDGMAVGIVPSFTRFLNRMQSGYLFHYAFAMVIGIVGLMFWVVLNGGAN------
>WP_078546626_NuoL_Thioclava_sp._DLFJ4-1
ME-------TIILFAPLVGAILGGFGW----RVIGEKGAMVLTTGLLFLAALLSWIVFLTL----GQTQHIHVL
---
NWITSGELDVSWAIRMDRLTAIMLIVVTTVSSLVHLYSWGYMAHDENFGKGENYKARFFAYLSFFTFTMLMLVT
SDNLLQMFFGWEGVGVASYLLIGFYYKKPSANAAAMKAFIVNRVGDFGFLLGMFGLFYMTGSLQL--
DDVFSAAPALSETN---
LDFLWGSWNAVELCAFLLFVGAMGKSAQLFLHTWLPDAMEGPTPVSALIHAATMVTAGVFLVCRMSPVYEYAPH
AQTFIVVIGASTAFFAATVGLVQNDIKRVIAYSTCSQLGYMFAAAGVGVYSVAMFHLLTHAFFKALLFLSAGSV
ITAMHHEQDMRNYGGLRKKVPLTFWAMIFGTLAITGVGIPFTP-----IGFAGFLSKDAIIESVWG--
GSGAGYAFWLLVIAAAFTSFYSWRLIFMTFFGKPR--GDHHAHDHA----------------------------
---------------------------------KESPWVMTVPLVVLSFGAIFAGMIWYNVFFGDHAKLVK--
FFGLPAAIYMGQENEVIDKAHHAPAWVKVSPFVAMLAGLLVAWLFYIKD-------------------------
----------------------------------
TSLPKRFAEAQPGLYRFLLNKWYFDELYERIFVRPALWFGYQFWKK-GDEGSIDR-
VIDGMAVGIVPTFTRFLNRMQSGYLFHYAFAMVIGIVGLMLWVVLNGGAN------
>WP_095595910_NuoL_Actibacterium_pelagium
MA-------SIILFAPLVGAIICGFGW----RIIGEKAAMWVATGLLFFACLLSWIVFLGFD---GQMQQIHIL
---
DFIQSGSLDTAWAIRLDRLTAIMLIVVTSVSALVHLYSFGYMDHDPQWGEGETYKPRFFAYLSFFTFAMLSLVT
ADNLVQMFFGWEGVGVASYLLIGFYWKKPTASAAAIKAFVVNRVGDFGFALGIFALFMLTDSIAF--
DDVFAAGPALADTT---
LHFLWRDWNAANLIGFLLFVGAMGKSAQFILHTWLPDAMEGPTPVSALIHAATMVTAGVFLVCRMSPIYEYAPQ
ATSFIVFLGALTAFFAATVGLVQNDIKRVIAYSTCSQLGYMFVAAGVGVYSVAMFHLFTHAFFKAMLFLGAGSV
IHAMHHEQDMRNYGNLRKKIPYTFWAMMIGTLAITGVGIPLTH-----FGFAGFLSKDAVIESAYA---
GGTGFAFWALVVAACFTSFYSWRLMFMTFFGEER--GDKHTHEHA-----------------------------
--------------------------------HESPKVMLIPLGALALGAVFSGMVWYGSFFGDNVNK----
FFGIPAAYAMGPDNHVLHDAHYVPKWVKISPFIAMLIGLGVAYLFYIVN-------------------------
```

```
----------------------------------
PALPKKLAEQQRPLYLFLLNKWYVDEIYDFLFVRPARALGNFLWKK-GDGTVIDG-
FLNGVAMGIIPFFTRLAGRAQSGYLFHYAFAMVIGIAVLVTWMTISGGAE------
>WP_105513845_NuoL_Defluviimonas_denitrificans
ME-------TIIVLAPLVAAIIAGFGW----RIIGETAAMALTTGVLFLACLMSWIVFLGFD---GTTQHIHVL
---
DWIQSGSLDTAWSIRVDRLTTIMLIVVTTVSSLVHLYSWGYMAHDENWGHHEHYKARFFAYLSFFTFAMLALVT
ADNLAQMFFGWEGVGVASYLLIGFYYKKPSANAAAIKAFVVNRVGDFGFALGIFGLFYLTDSILM--
DDVFAAAPTLAETR---
LGFLWTDWNAANLLAFLLFIGAMGKSAQLFLHTWLPDAMEGPTPVSALIHAATMVTAGVFLVCRMSPLFEYAPE
AKTFVIYIGAATAFFAATVGLVQNDIKRVIAYSTCSQLGYMFVAAGAGIYSAAMFHLLTHAFFKAMLFLGAGSV
IHAMHHEQDMRNYGNLRKKIPLTFWAMMIGTLAITGVGIPLTS-----IGFAGFLSKDAIIESAW---
GTGNGFAFWALVVAALFTSFYSWRLMFLTFYGKER--GDHHAHEHA----------------------------
---------------------------------HESPAVMTVPLGVLALGAIFAGMLWYKPFFGDHERMLS--
FFAMPVEAASEATNTVFDNAHHAPAWVKVSPFIAMLIGFFTAWVFYIRD-------------------------
----------------------------------
PSIPGRLAAQQPILYRFLLNKWYFDEIYEAVFIRPARWLGNFLWKK-GDGDVIDG-
TINGVAIGFVPMLTRIAARLQSGYIFTYAFAMVIGIAVLLLWMTLSGGAH------
>HDZ53908_NuoL_Sulfitobacter_litoralis
ME-------TIILFAPLVGALICGFGW----KLIGEKAGQYVATGLLFLAALLSWIVFLSFD---GETQHIQIL
---
RWIESGSLSTDWAIRLDRLTAIMLIVITTVSSLVHLYSFGYMAHDENFSDAEPYRARFFAYLSFFTFAMLMLVT
SDNLVQMFFGWEGVGVASYLLIGFYYKKPSANAAAIKAFVVNRVGDFGFLLGIMALFYLTDSIKF--
DDIFAATPDLAETT---
LGFLWSEWNAANLIAILLFIGAMGKSAQLFLHTWLPDAMEGPTPVSALIHAATMVTAGVFLICRMSPLMEVAPE
AMMFVTFLGATTAFVAATIGLVQNDIKRVIAYSTMSQLGYMFVAAGVGMYSAAMFHLFTHAFFKAMLFLGAGSV
IHAMHHEQDMRNYGALRKKIPYTFWAMMIGTLAITGVGIPLTH-----FGFAGFLSKDAIIESAW---
GGGSMYGFWMLVIAAAMTSFYSWRLMFLTFFGKAR--GDKHTHDHA----------------------------
---------------------------------HESPMVMLVPLGVLSLGAIFAGMVWYNSFFGHAEDVGE--
FYGIPMALYIAPDNHVLDDAHAAPAWVKVSPFIAMVFGLVMALWFYIWN-------------------------
----------------------------------
PSLPARLAANQRPLYLFFLNKWYFDELYNVIFVKPAMAIGRFFWKR-GDGNVIDG-
SLNGVAMGIIPFLTRLAGRAQSGYIFTYAFAMVIGIAVFVTWMTMSGGAH------
>WP_118942038_NuoL_Profundibacter_amoris
ML-------QTILFAPLIGALIAGFGW----RIIGEKGAQMLTTGLLFLSALLSWVVFLTFD---GETQHIQIL
---
RFIESGTLSTDWGIRLDRLTAIMLIVVTTVSSLVHLYSFGYMEDDPQWGEGQNYKARFFAYLSFFTFAMLALVT
ADNLVQMFFGWEGVGVASYLLIGFYYKKPSANAAAIKAFVVNRVGDFGFALGIFGLFYLTDSIKM--
DDVFAAAPQLAETN---
LQFLWTEWNAANLLAVLLFIGAMGKSAQLFLHTWLPDAMEGPTPVSALIHAATMVTAGVFLVCRMSPLMEYAPQ
ATTFIVVIGAATAFVAATIGLVQNDIKRVIAYSTMSQLGYMFVAAGVGAYGVAMFHLLTHAFFKAMLFLGAGSV
ITAMHHEQDMRNYGNLRKKIPMTFYAMLIGTLAITGVGIPLTS-----IGFAGFLSKDAIIESAF-
GGTGASYYAFWALVIAAAFTSFYSWRLMFMTFWGKAR--GDAHTHDHA--------------------------
-----------------------------------HESPLTMLVPLGVLGLGAVFSGMVWYNSFFGDHEKMLT-
-FFGMEEAIFMGPENEVIEKAHESPAWVKVSPFIAMVLGFVLAWYFYIVN-----------------------
-----------------------------------
TAAPRRLAEIQPMLYRFLLNKWYFDEIYDALIVRPAFAIGRFLWKK-GDMGVIDG-
SINGVAMGIIPFFTRLAGRAQSGYVFTYALAMVIGIVVLITWMAISGGAE------
```

```
>MBT6020484_NuoL_Planktomarina_temperata
ME-------TILLFAPLVGALIAGFGH----GVIGDKAAQVLTTALLFLSALLSWVLFLSFD---GQTESIHIL
---
RWIESGTLSTDWAIRMDRLTTIMLIVITTVSALVHLYSMGYMAHDDHFRDGESYRPRFFAYLSFFTFAMLMLVT
SDNLLQLFFGWEGVGVASYLLIGFYIRKQSAGAAAIKAFVVNRVGDFGFLLGIFGLYLVADSIKF--
EDIFLIGPQLATME---
LHFLWRDWNAANLIAFLLFVGAMGKSAQLILHTWLPDAMEGPTPVSALIHAATMVTAGVFLVCRMSPIMEFAPD
TMNFIVFLGASTAFFAATVGLVQNDIKRVIAYSTCSQLGYMFVAAGLGVYSVAMFHLLTHAFFKAMLFLGAGSV
IHGMHHEQDMRNYGGLRKKLPYTFWAMMIGTLAITGVGIPLTH-----YGFAGFLSKDAVIESAYA--
GTMGGYAFWMLVIAAMFTSFYSWRLMFLTFFGAAR--GDKHTHEHA----------------------------
---------------------------------HESPKVMLVPLAVLAVGAIFSGMVFYKPFFGDHHKVEV--
FFGIVEAFYMAPSNTVLDDAHHVHWLVKLSPFIAMVLGFAGAWAFYMTK-------------------------
----------------------------------
LDLPRRVAESNPVLYRFLLNKWYFDEIYDVIFLRPAQWLGRTLWKV-GDGKIIDG-
GINGLAMGIIPFFTKWAGRMQSGYIFTYAFGMVIGIAALVTWVTIAGGK-------
>WP_133489491_NuoL_Aliiroseovarius_marinus
MV-------QFILFAPLFGALVAGFGH----RFIGDKGAQILTTALLFWSAFLSWITFFGLG---SETQIIHLM
---DWVQSGSLSVDWSIRLDRLTAIMLVVVTSVSSLVHLYSMGYMAHDPQF-
EGESYRPRFFAYLSFFTFTMLMLVTSDNLLQMFFGWEGVGVASYLLIGFYYKKPSANAAAIKAFVVNRVGDFGF
ALGIFGLFYMTDSIRF--DDVFAAAPALAETN---
IHFLWTDWNAANLLAFLLFVGAMGKSAQLILHTWLPDAMEGPTPVSALIHAATMVTAGVFLVCRMSPLMEYAPE
ATNFIVFLGATTAFFAATVGLVQNDIKRVIAYSTCSQLGYMFVAAGVGVYSVAMFHLFTHAFFKAMLFLGAGSV
IHGMHHEQDMRNYGDLRKKLPITFWAMMIGTLAITGVGIPLTT-----IGFAGFLSKDAVIESAFA--
GTNGGYAFWMLVIAALFTSFYSWRLMFMTFYGKAR--GDKHTHEHA----------------------------
---------------------------------HESPLVMTIPLGVLAIGAIFAGMIWYGPFFGKEDKVNQ--
FFGIEGEAEMHPDNHVLHDAHYVPKWVKLSPFVAMIIGFVTAFWFYIKN-------------------------
----------------------------------
PSLPKRLAETFPHVYNFLLNKWYFDEIYDILFVKPAFMVGRFLWKR-GDGNVIDG-
FLNGVAMGIVPWFTKQAGRAQTGYLFSYAFAMVLGIVALVTIMIFSGGAH------
>WP_125403843_NuoL_Rhodovulum_robiginosum
ME-------KIILFAPLLGALLCGFTW----RLIGEKAAMWTATVLVFVSMVLSWVVFFGFD---GEMRQVEVF
---
RFIESGTFTADWAIRIDRLTAIMLIVVTTVSAFVHLYSFGYMDHDPQWQEEEHYKARFFAYLSIFTFAMLALVT
ADNLIQMFFGWEGVGLASYLLIGFYFKKPSANAAAMKAFIVNRVGDFGFALGIFALYFLVDSVNF--
TDVFAAAPEIAETE---
LSFLNGSWNAANLIGILLFIGAMGKSAQLLLHTWLPDAMEGPTPVSALIHAATMVTAGVFLICRMSPLIEFAPQ
AQTMIVVVGAATAFFAATVGLVQNDIKRVIAYSTCSQLGYMFVAAGVGVYSAAMFHLLTHAFFKALLFLGAGSV
IHAMHHEQDMRNYGGLKDKIPYTFYAMLIGTLAITGVGIPLTS-----IGFAGFLSKDAIIESAFG---
SGDTWAFWTLVIAAAMTSFYSWRLMFMTFWGTPR--GDKHTHEHA-----------------------------
--------------------------------HESPRSMLIPLGVLAFGAVFAGMAWYGSFFGDHDRVNR--
FYGIPSTAEAGADNHVMDEAHHAPQWVKAAPFLAMLLGLGLAYQFYLVS-------------------------
----------------------------------
PSVPGQLARQFRPVYLFLLNKWYFDELYDAIFVNPARNIGRALWKG-GDGSLIDG-
AINGLSMGFVPFLTRLSGRMQSGYLFHYAFAMVLGIALLVTWMSLGGGAN------
>WP_187428657_NuoL_Roseobacter_litoralis
ME-------TIILFAPLLGALLCGFGW----KVFGEAAAMWIATALLFVSAILSWVVFLSFD---GTTEQIQIL
---
RFIESGTLSTDWAIRMDRLTAIMLVVITTVSSLVHLYSFGYMDKDPQWKEGETYKPRFFAYLSFFTFAMLMLVT
```

```
ADNLVQMFFGWEGVGVASYLLIGFYYRKPSANAAAIKAFVVNRVGDFGFALGIFALFFLTDSINF--
DDVFAAAPDLAETQ---
LTFLWGEWNAANLIAMLLFIGAMGKSAQLLLHTWLPDAMEGPTPVSALIHAATMVTAGVFLVCRMSPIMEFAPE
AMAFVTVIGATTAFFAATVGLVQTDIKRVIAYSTCSQLGYMFVAAGVGMYSAAMFHLFTHAFFKAMLFLGAGSV
IHAMHHEQDMMNYGGLRKKIPYTFAAMMIGTLAITGLGIPVWGDIP--IGFAGFQSKDAIVESAYA---
AGTMYGFWALVIAALFTSFYSWRLMFLTFYGKPR--GDKHTHDHA------------------------------
--------------------------------HESPMTMILPLGVLALGAIFAGMVWYGSFFGHTDKVAK--
FYGIPYA-EAGADNTLLDDAHKVPYWVKTSPFAAMLIGFLTAYWFYIKN-------------------------
----------------------------------
PSLPGRLAQNQRPLYLFLKNKWYFDELYDVIFVGPAKWIGRMLWKR-GDGDVIDG-
GLNGVAMGIIPFFTRLAGRAQSGYIFTYAFAMVIGIAVLVTWMTMSGGAN------
>WP_102223097_NuoL_Acidimangrovimonas_sediminis
MV-------EFILFAPLVGAIVAGFGW----RFITETGAEWLTTALLFAAAIAAWFVFFSFD---GNTQVVTIF
---
RWIDSGSLQANWGVRLDRLTTIMLIVVTTVSALVHLYSIGYMAHDEDESKGENYRARFFAYISLFTFTMLALVT
ANNLLQLFFGWEGVGLVSYLLVGFYYKKPSANAAAIKAFIVNRVGDFGFLLGIFAIFVATDSINF--
DIIFPKSAELAQMH---
IGFLWTTWNAADVIGVLLFIGAAGKSAQLFLHTWLPDAMEGPTPVSALIHAATMVTAGVFLVCRMSPLYEYAPG
AAALVLVIGATTAFFAATVGLVQTDIKRVIAYSTCSQLGYMFAAAGAGVYSAAMFHLMTHAFFKALLFLCAGSV
ITAMHHEQDMRSYGGLRKKIPFTFWMMVIGTLAITGTGIPFTGDGP--IGFAGYDSKDAIIEGVWG--
ANGAGYAFWMLVIAAAFTSFYSWRLIFMTFFGAPK--GDKHAHEHA-----------------------------
---------------------------------HESPMVMLIPLGVLALGATFAGMIWYNSFFGNEKSMES--
FFHLP--THAERDNQAIEHAHHAPAWVKISPFIAMALGFLLAWVMYIRS-------------------------
----------------------------------
PSMPGKLAESQRPLYLFLLNKWYFDELYDAIFVRPMLAVGRFFWRW-GDGVVIDG-
SINALATRIIPAMTRAAGRLQSGFVYTYAFTMVIGITLLVTWMTLTGGAQ------
>WP_050686206_NuoL_Phaeobacter_italicus
ME-------TILLFTPLVGALVCGFGH----KILGEKVATVFATALLFLTAALSWIVFLTFDPS-
AHENGYYTVEILRWIQSGSLDTSWQFRVDRLTAIMLIVITSVSSLVHLYSFGYMDHDPQWKDGESYKPRFFAYL
SFFTFAMLMLVTADNLVQMFFGWEGVGVASYLLIGFYYRKPSAGAAAMKAFIVNRVGDFGFALGIFALFFLTGS
INF--DDIFAATPMLAETQ---
LGFLWTEWNAANLIAFLLFVGAMGKSAQLILHTWLPDAMEGPTPVSALIHAATMVTAGVFLVCRMSPLMEVAPE
ATAFITVLGATTAFFAATVGLVQTDIKRVIAYSTCSQLGYMFVAAGVGMYSAAMFHLFTHAFFKAMLFLGAGSV
IHAMHHEQDMMNYGGLRKKIPYTFWAMMIGTLAITGVGIPLTH-----IGFAGFLSKDAIIESAYA---
GGSMYGFWMLVIAAAMTSFYSWRLIFLTFYGKPR--GDKHTHDHA------------------------------
--------------------------------HESPMVMLVPLGVLALGAVFSGMVWYNSFFGHTDTVGK--
FYGIPYAEAAEADNTLLDDAHAVNKWVKVSPFIAMVLGLSLAIWFYIVN-------------------------
----------------------------------
PSLPGRLAQTHQPLYQFLKNKWYFDELYNYVFVKPALALGRFFWKR-GDGSTIDG-
ALNGLAMGIIPFFTRLAGRAQSGYIFTYAFWMVLGIAALVTWMSIGGGAH------
>WP_16085423_NuoL_Oceanomicrobium_pacificus
MT-------DIILFAPLVGALICGFGH----RFIGEKAAMWTATGLLFLACALSWIVFLFGD---
YHHHAVSHAV-
LRWIESGTLSSEWGIRLDTLTAIMLIVVTSVSALVHLYSFGYMDHDPQWKEGESYKPRFFAYLSFFTFAMLMLV
TSDNLLQMFFGWEGVGVASYLLIGFYYRKPSANAAAMKAFIVNRVGDFGFLLGIFATYMLVDSIRF--
DDIFAAAPALAEES---
FHFLSWEVPAVETIAVLLFIGAMGKSAQLFLHTWLPDAMEGPTPVSALIHAATMVTAGVFLVCRMSPVMEYAPG
ALMFVTVIGASTAFFAATIGLVQNDIKRVIAYSTCSQLGYMFAAAGVGVYQAAMFHLFTHAFFKAMLFLGAGSV
```

```
IHAMHHEQDMRNYGGLRSKIPYTFWAMMLGTLAITGVGIPLTH-----
IGFAGFLSKDAIIESAFAAHSGVGTYAFTLLVLAAAMTSFYSWRLMFLTFYGKPR--GDAHTHEHA--------
-----------------------------------------------------
HESPMVMVVPLGILALGAIFAGMIWYEDFFGHHAKE----FFGA--
SVFMHPDNHVMHDAHEVSKLVKVSPFLAMLVGLALAYWFYIVN-------------------------------
----------------------------PSIPGRIAKTNHGLYQFLLNKWYFDELYDAVFVRPAKALGRFLWKR
-GDGSVIDG-GINGLAMGVVPFFTRLVGRAQSGYIFHYAFAMILGVVVIVTWFAVSGGAQ------
>WP_012177999_NuoL_Dinoroseobacter_shibae
ME-------SIILFAPLVGALICGFGW----KFIGEKAALWVSTGFVFLAAILSWFVFLTFD---GTTEQIQVL
---
RWIESGSLAADWAIRMDRLTAIMLIVVNTVSALVHLYSFGYMDHDPQWREGETYKPRFFAYLSFFTFAMLMLVT
SDNLVQMFFGWEGVGVASYLLIGFYWRKPSAGAAAMKAFIVNRVGDFGFLLGIFALFYLTDSVNL--
TDIFAAAPELAETQ---
ISFLWTDWNAANLIAFLLFIGAMGKSAQLFLHTWLPDAMEGPTPVSALIHAATMVTAGVFLVCRMSPVMEFAPQ
AMTFVTFVGATTAFFAATVGLVQNDIKRVIAYSTCSQLGYMFVAAGVGAYPVAMFHLFTHAFFKAMLFLGAGSV
IHAMHHEQDMRNYGGLRKKIPYTFWAMMIGTLAITGVGIPLTGW----
IGFAGFASKDAVIESAFAGTNAAHMYAFWMLVIAALMTSFYSWRLMFMTFYGTPR--GDKHTHEHA--------
-----------------------------------------------------
HESPKVMLIPLGVLALGSVFAGAIWYGSFFGHTDEVAK--
FYGIPYAEYMAPENTVMADAHDVPKWVKVSPFVAMLIGLGLSYLFYIRK-------------------------
----------------------------------
PSLPGTFAETLWPVYNFIYNKWYFDEIYDAVFVKPSKAIGRFLWTK-GDGATIDG-
GINGLAMGIIPFFTRLAGRAQSGYLFHYAFAMVLGITILVTWMMIGGGAE------
>WP_150968212_NuoL_Aureimonas_leprariae
MY-------ALIVLLPLAGFLIVGFLG----NSIGAKASEYVTSGLMIVAAVLSWIAFLFLAP--EAPQSHAIL
---QWMQAGSLDVSWSLRIDRLTLVMLVVVNTVSALVHVYSIGYMHHDPH-------
RPRFFCYLSLFTFAMLMLVTSDNLVQMFFGWEGVGLASYLLIGFWYKKPSASAAAMKAFIVNRVGDFGFALGIF
GIFVLFGSVNF--DTIFAGAADLATKL---
GEGGGEAAAAVTAICLLLFMGAMGKSAQLGLHTWLPDAMEGPTPVSALIHAATMVTAGVFMLARMSPIFELSHP
ALTFVTFIGATTAFFAATIGLVQNDIKRVIAYSTCSQLGYMFAALGVGAYGPAIFHLFTHAFFKALLFLGAGSV
IHAVSDQQDMRNMGGLRKHIPKTYWMMVIGTLALTG--FPF---------
TAGYYSKDAIIESTWAGDNAFAVYACGLTIIAAALTSFYSWRLIFMTFHGKPR--ASHEVMHHI----------
---------------------------------------------------
HESPPVMLVPLYVLGAGALLAGILFEHFFIGE-GYGE---FW--
RAALVTGAENHVLHEMHEAPFLIKIAPFLAMAGGFIVAWIFYIRS-----------------------------
------------------------------
QSLPARTAATNPGLYRFLLNKWYFDELYDVVFVRSAKALGRFFWKTGGDQKVIDGLGPDGIS-
ARVLDVTDRVVRIQSGYLYHYAFAMLIGIAALVTLMMV--GGR------
>WP_160587554_Pyruvatibacter_mobilis
MY-------SAIVFLPLLGFLIAGLFG----RVIGHRGSEIVTTSLLMLAALLSWIAFFDVAF--GGYT-
GKVHIL-RWIDSGALEVDWMIRVDTLTAVMLVVVNTVSSLVHLYSIGYMSHDPH-------
RSRFFAYLSLFTFAMLMLVTADNFVQMFFGWEGVGLASYLLIGFWYKKPSANAAAMKAFVVNRVGDFGFLLGVA
GTFLVFGSLDF--DTVFAAVPEVAGQT---
FAFAGMQVDIMTTLCLLLFMGAMGKSAQFLLHTWLPDAMEGPTPVSALIHAATMVTAGVVLVARTSPMFEFAPD
ALWFVTLIGATTAFFAATVGLVQNDIKRVIAYSTCSQLGYMFVALGLGGYQMAVFHLFTHAFFKALLFLGAGSV
IHAMSDEQDMRKMGGIFKMIPGTWIMMIIGTLGLT--GVPY---------
LAGYYSKDAIIEAAYMANSGLAGYAFAMTVVAALMTSFYSWRLIFMTFHGESR--ASNEVLSHV----------
---------------------------------------------------
```

```
HESPWVMLIPLIVLSAGTVLAGFPMREFFVGH-DQAT---FW--
NGSLFTLPSNNLIEEFHHAPALVVWSPFIMMVIGLATAWLFYIVR-----------------------------
------------------------------
PGIPKALAREQQPLYQFLLNKWYFDELYNVIFVKPAMWLGRTLWKGGGDGFIIDGFGPNGIA-
ARVMDVTRNVVKLQTGYVVHYAFAMLIGVAAFVTYYSL--GAH------
>WP_213162180_NuoL_Kaustia_mangrovi
MY-------SLIVFLPLIGALVAGLAG----RWIGAQASRVLTSGLVVVSAVLSWVAFYDVGL--
GHETYKVQVL--SWIHSGAFEADWAFRVDTLTAVMLVVVNTVSALVHIYSIGYMSHDPH-------
QPRFFAYLSLFTFAMLMLVTADNLVQMFFGWEGVGLASYLLIGFWYQRPSANAAAIKAFVVNRVGDFGFSLGIF
ACFTVFGAVDL--DTIFASAPDVAGQT---
MVFLGYEVDILTTICLLLFMGAMGKSAQFLLHTWLPDAMEGPTPVSALIHAATMVTAGVFLVARMSPIYEFAPD
ALAFVAFIGATTAFFAATIGLVQNDIKRVIAYSTCSQLGYMFAALGVGAYGVAIFHLFTHAFFKALLFLGSGSV
IHAMSDEQDMRKMGGLFPHLKKTWAMMLVGTLALTG--FPL---------
TAGYFSKDAIIEAAFAAHSGVGQYAFWLTVIAAFLTSFYSWRLMFMTFHGRQR--ASRDVMAHV----------
---------------------------------------------------
HESPNVMLVPLYLLAIGALVAGGLFKGLFIGH-EEAE---FW--
GAALLRGEGNEIVEAMHHVPGWVPFAPTVMMVAGFALAWLFYIAR-----------------------------
------------------------------
PEMPKALAEMHRPLYLFLLNKWYFDELYDLVFVRGAKALGRFLWKR-GDGWAIDGFGPDGVS-
ARVIDITNRIVRLQTGYLYHYAFAMLVGIAALITWLMF--GSM------
>WP_167095161_NuoL_Parvibaculum_indicum
MF-------SAIVFLPLLGFLTAGIFG----RWLGPRGAQIATCVPMVIAAILAAFAFVDVGLQ-GETY-
RIQVL--TWINSGAFEADWRLRVDTLTAVMLVVVTWVSALVHIYSIGYMSHDPQ-------
QPRFFAYLSLFTFAMLMLVTADNFVQLFFGWEGVGLASYLLIGFWYKKPSANAAAIKAFVVNRIGDFGLILGIA
TLFFTIGSVDF--DTVFKAIPELQDKT---
FLFLGYDVPVVTTACLLLFMGAMGKSAQFLLHTWLPDAMEGPTPVSALIHAATMVTAGVFLVARMSPVFEFSPF
ALTVVTIIGAITAFFAATVGLVQNDIKRVIAYSTCSQLGYMFAALGVGAYEPAVFHLFTHAFFKALLFLGSGSV
IHAMSDEQDMRKMGGLFRMIPATWLMMIIGTLALTG--FPF---------
TAGYFSKDAIIEATFAGENPAHMFAFVMLVVAALFTSFYSWRLIFLTFHGQSR--ASNETLSHV----------
---------------------------------------------------
HESPMVMMLPLVVLAVGAVFAGMAFSDLFIGK-AHEA---FW-LSS-
IFKGPDNHVLHDMHHIPAWAIWAPTAMMVIGLAVAVLFYLVK--------------------------------
---------------------------PELPKQLAREQEPLYKFLLNKWYVDEIYDFLFVRPAFWVGRLLWKQ-
GDGRIIDGLGPDGVS-ARVLDVTRGVVRLQSGYLYHYAFAMLLGVAALATYFMF-GGAH------
>ATY40894_ND5_Picozoan_Picobiliphyte_sp.MS584-11
MY-------LTIVFLPLVGSMIAGLGG----RWVGPIGACLVTTTCLLISFFFSCIVFYEVGLC-GSP--
CHIHLL-DWFDSAAFVGAWGFQFDTLTATMLIVVTGVSSCVHIFSVDYMSGDPH-------
RARFMSYLSFFTFFMLTLITADNFIQLFFGWEGVGLCSYLLINFWYSRITANRAALKAFIMNRVGDFGFGLGIF
GCYMVFQSIEF--ETIFACAPLYANQT---
FEFFNFEVDQLTCIVCLLFVGAVGKSSQIGLHTWLPDAMEGPTPVSALIHAATMVTAGIYMLLRCSPLLEYAPD
ALSVIAFFGASTAFFAATTGLLQNDLKKVIAYSTCSQLGYMCFAVGLSKYSVSFFHLANHAFFKALLFLSAGCV
IHAMQDEQEMRRMGGLLSSLPFTYSTMTLGSLALMG--TPF---------
LSGFYSKDAILEYASIHYFIAGSFTFWLGVLAAFCTAFYSFRLGYMTFLTNTN--AYRYCVEHI----------
---------------------------------------------------
HDAPGIVLVALCPLAVGSIWSGYFGADMFLGV-GTP----FW--
GNALFFGWTTTLTLEPHFLPMATKFLPLVFTILGGFLAFLAFNYLAN---------------------------
------------------------------
ITFTLTFHPYIKPIFAFLNKKWYFDKIYDEVFLVQSMLFGRNVTYRLIDGFVFEQLGPMGIQ-
```

```
KTIGVLSGSHKRFHNGFIPNYTSIFFFGILLFLVGFVALPKCLVLYPGL
>BBU60045_NAD5_Cyanidioschyzon_merolae
MY-------LTTVFAPLLGFLTCMLFG----RFLGKIGACFIVCFCTAISLLFSIIIFYEVCIA-DYT--
NNIFFT-LWLSTGLLEIPWGFLFDPLTAVILLTINFVSLLVNVYSVEYIGEDPH-------
QVRFIAYLNIFAFFITILVTANNFLQIFLGWEGVGLASYLLINFWFTRLLASKSAIKAIIINRVGDFFLSLAIF
LIFLKYKSLNY--DIVFSITPFMSNYI----
ECLGIKFSYLDMISLFLFLGAIGKSAQIGMHIWLPDAIEGPTPVSALIHAATMVTAGVFLIVRCSPILEYSSNV
LIFISVIGASTSLFAGSVAIFQNDLKKVIAYSTCSQLGYMIFACGISNYIPSIFHLVNHAFFKALLFLSAGYII
HSLFNEQDIRRIGSLLRILPIAYLMIMIGSLSLIG--IPF---------
LTGFYSKDLILELSFTSYFIINDFVYYLAIVSACLTAIYSTRLILLTFITKPN--FSKISLIKL---------
-------------------------------------------------
HYSSNWITLPLLILAFASICFGFILKDIFVGL-GTN----FW--GSSILINYNHIYFIEAEN-
PIIIKWIPFFASIIGFFIIIIMQNNYAI-------------------------------------------
-----------FSFSFVRY-MHSFFY-FFNKKWYLDKIIIIIA-NSVLNFGYYVSLKIFDKGVIEYFGPFGLI-
KIIPYMSKQSRNLQSGQIAHYVFIILIGLLVLIIVIEFNNLYFIINYIK
>AIA61058_ND5_Cyanidiaceae_sp._MX-AZ01
MH-------LTVILAPLIGFLTCGLFG----RFIGKKGANFISCFCICVSLLFSIIIFYEACFS-NYK--
NHILLS-SWIDSAFLEVSWGFIFDSLTAVILLIVNFISLLVIIYSIEYIQEDPH-------
QIRFISYLTIFVFFITILVTANNFMQLFLGWEGVGLASYLLINFWFTRLQANKSAIKAIIINRVGDFFLSLGIL
FIFIKFQSLNY--NVVFSIVPFITDNF---
LECYGIKFSYLDIITLLLFLGAMGKSAQLGIHIWLPDAIEGPTPVSALIHAATIVTAGVFLVVRCSPIFEFSNF
TLVFVSVVGALTSFFAGSVAIFQNDLKKVIAYSTCSQLGYMIFVCGISNYTSSIFHMVNHAFFKALLFLSAGYI
IHSLINEQDIRRIGALMRILPIAYIIMLIGSLSLIG--IPF---------
ITGFYSKDLILELIFESYFVIHDFVYWLAIVSVFLTAMYSIRVILLVFVKNSN--TARINLINL---------
-------------------------------------------------
HYSSFFITSPLFILAFASICFGFLLKDIFVGL-GTN----FW--
GSSVIIYYSQVNFLEVENLSTVIKWLPFIISIIGVFLILLSINYDLF--------------------------
-----------------------------RLFPLEK--IHTIFY-FFNKKWYFDKIITTIA-
NFILSFGYDISLRIFDKGIIEYFGPFGLI-
KIVPYFSIRSKNLQTGQIAHYIFVILIGFLSLVLFTEFLNIINLIHFEK
>YP_010007659_NAD5_Cyanidium_caldarium
MY-------LIILFLPFCGSLFSILFG----RWVGGFGSSVITCLCLILAIFLSCIAFYEVGIL-NYP--
CYIKLT-SWIDSGIFHVSWGFLFDSLTVTMIVIVSVISLLVHIYAIEYIRLDPH-------
LPRFISYLSVFTFFILILVTANNFIQIFLGWEGVGFASYLLINFWFTRLSANKAALKAIVINRIGDVGLSLGIL
VIFLKFHSVDY--LTVFNLVPNVFFES---
FNFFNIKVNFCTIICLLLFIGVIGKSAQIGLHTWLPDAMEGPTPVSALIHAATIVTAGVFLIIRCSPLFEFSDT
ALFIVAIIGGITAFFSASIGLFQYDLKKIIAYSTCSQLGYIVFSCGLSNYSVGFFHLFNHAFFKALLFLSAGSV
IHAFLNEQDIRFIGGLKNLLPFTYAIMIVGSFSLLG--FPF---------
ITGFYSKDLILESAYNNFNIGSDFIYWLGLITVLLTAFYSFRLLFLVFLNYPN--FSTLIFNKV---------
-------------------------------------------------
NDCSFYIKTSLTFLAIGSIFLGYLTKEMLVGM-GTD----FW--
DISLFFKLNHIDFVENEYLYFLIKLIPIFFSFFGILF-CYLFNYFIN--------------------------
-----------------------------LKF-LEVQLRFRYFLFFLNKKWGWDYIYTYIA-
YQFMNFGYYISFKLIDKGILEFFGPQSLM-
KLITNMLNKLKFIQTGQITHYVFIIILGALVLFTLVELLDVLVYFFNIK
>YP_009968183_ND5_Cyanidiococcus_yangmingshanensis
MY-------LITIFAPLLGFIICGLFG----RFLGKKGACFITYFCSSISLVFSIIIFYEVSVA-AYD--
NHILLA-KWVESSIFFVSWGFNFDSLTALILVMVSFISLLVNIYSIEYIEEDPH-------
```

```
QVRFISYLNIFVFFMFMLVTANNFIQLFLGWEGVGLSSYLLINFWFTRLIANKAAIKAIIINRVGDFFLSLGIF
FIFFKFKSLDY--NVVFSIAPSIIDNY---
LECLGIKFLYLDVVTLFLFLGAIGKSAQIGIHMWLPDAIEGPTPVSALIHAATMVTAGVFLIVRCSPIFEFSHF
TLVIISVVGALTSLFAGSVAIFQNDLKRVIAYSTCSQLGYIMFACGISNYMASIFHLLNHAFFKALLFLSAGYI
IHSLANEQDIRRIGSLIRIIPIAYIAIIIGSLSLIG--IPF---------
LTGFYSKDLILELTFGSYFIIHDFTYWLAIISASMTAIYSMRLVLLTFIKSTN--VIKINLIHI----------
---------------------------------------------------
HYSSSWMIVPLVILIIGSIYFGFILKDIIIGL-GTD----FW--
GSSILTHYSRANFVEVENLHVIIKWLPFFVSILGVVIILFIYNYDFK---------------------------
------------------------------IFSTIFLKYTHSFFY-FFNKKWYFDKIANIIA-
IYLLSFGYNVCLKVFDKGIIEYFGPFGLM-
KIVPYLSTESKSLQTGQIAHYIFVILMGLLFLILSTEFINIFYIINYIK
>NP_062497_ND5_Chondrus_crispus
MY-------LLILFLPLLGSLISGFGG----RWLGCRGTNTFSTLCVVVSSLFSLLAFFEIGLT-NTT--
CTIFLV-SWIKSGAFYVSWGFLFDSLTVTMLVVITLVSSLVHIYSIKYMENDPH-------
QPRFMSYLEIFTFFMLILVTADNLIQMFLGWEGVGLASYLLINFWYTRLAANQSAIKALIVNRVGDFGLSLGIF
LIFWVFNSVDY--SVIFSLVPLFDNQF---
LTFLGFKLHVLTLISLFLFIGAIGKSAQLGLHTWLPDAMEGPTPVSALIHAATMVTAGVFLMIRFSPLLEFSPT
ILFILITFGSLTAFFAAVTGVFQHDLKRVIAYSTCSQLGYMIFSCGMSCYDVSLFHLANHAFFKALLFLSAGSV
IHAVSNEQDMRRMGSLLKFMPLTYSVMLIGTLALIG--FPF---------
LTGFYSKDFILELTSSLQMSYISFACWLGTMSVFFTSFYSFRLIYLTFLNNTN--LAKSSLNLV----------
---------------------------------------------------
HESSLLMIFPLIILSIGSIFAGYLIRDLFVGS-GSD----FW--
GAAIFILPKHSTFIEAEELPIVVKWLPFILSLLGIFFASFVQIFLKT---------------------------
------------------------------
FYFKSNLQNLLSFFTFLINKKWYWDVLYNRLIVLPILNFGYSISFKILDRGFIELSGPYGFT-
KFVSFWSQILIKLQTGQITHYLFFMIFTFCSF---SIILVYSYINLTFN
>QKZ95183_NAD5_Pyropia_pulchra
MY-------LLIVVLPLVGTLVTGLGG----RWIGRKGANLFSTTCVILCCCLSIVAFFEVGLC-GVP--
CYLSLS-PWISSGALKISWGFLFDSLTTTMLVVITSISSLVHLYSIQYMEYDPH-------
CPRFCPSWKFFTFFMIVLVTADNFVQMFLGWEGVGLASYLLINFWYTRLCANQAAIKALVVNRVGDFGLSLGIF
TIFFLFGSVDY--ELVFASASLYTNYS---
IYFLGCSVNFLTIIGIFLLIGAIGKSAQLGLHTWLPDAMEGPTPVSALIHAATMVTAGVFLIVRCSALINLSSN
VLFLITILGSSTAFFASIVGVFQNDIKRVIAYSTCSQLGYMLFVCGLSYYNVGMFHLVNHAFFKALLFLSAGSV
IHALSNEQDMRRMGSLANSLPITYAAMLIGSLSLAG--FPF---------
LTGFYSKDLIIEITSSLQITFGIFACWLANISVFFTAFYTFRLLFLTFVKNSN--SYEKNIENI----------
---------------------------------------------------
HESPILILIPLILLSLASIFVGFLTKDLFVGV-GTS----FWGNAINILP-TSCNLLEVEF-
MSYLIKWLPFVLSINGAIFAYTLNIGYYK---------------------------------------------
------------NNIQFAYNHIFRKLAFSLSKKLYWDKLYNFLVVSPLMQFGYNVSFKNIDRGFIELLGPYGIS
-RTIKKWSTQILKIQTGQVTHYTFFVVSGLCLFFLFAPVWSSLEFFIDIR
>YP_006665880_NAD5_Porphyra_umbilicalis
MY-------LLIIALPLIGTLFTGLGG----RWLGRKGSNIFSTTCVILCFFLSLLAFFEVGLC-GVP--
CYVSIS-PWISSGVLNITWGFLFDSLTTTMLVVITSISSLVHLYSIQYMEYDPH-------
CPRFMSFLEIFTFFMLLLVTADNFVQMFLGWEGVGLVSYLLINFWYTRLCANQAAVKALIVNRVGDFGLSLGIL
TIFSLFGSVDY--EVVFSLVHTYSNHN---
IYLFGAHLNTLTLVGIFLLIGAVGKSAQLGLHTWLPDAMEGPTPVSALIHAATMVTAGVFLIVRCSPLIDLAPN
VLFLITILGSSTAFFASIVGVFQNDIKRVIAYSTCSQLGYMVFVCGLSYYNVGMFHLVNHAFFKALLFLSAGSV
```

```
IHALSNEQDMRRMGSLMHNLPITYAAMLIGSLSLAG--FPF---------
LTGFYSKDLIIEITSSLYITYGIFACWLANISVFFTSFYTFRLIFLTFIKNNN--SYRTYMDGI----------
---------------------------------------------------
HESPKLILIPLILLAIASIFVGFISKDLFVGV-GNS----FWGNSISTMP-IACNLLEVEF-
MTTSIKWLPFVLSTLGAFFAYSVNAGIFK---------------------------------------------
------------NNITFAYSNRFRKLAFSLSKKLYWDKLYNGLIVSFLLNFGYNISFKNIDRGFIELLGPYGIS
-KIISNLSFKIAKIQTGQITHYTFFVVTGLCLLLLLVPFFTFLENLIDIR
>AIU44693_NAD5_Cyanophora_paradoxa
MY-------LTLIILPLLSALVG-FLV----YFIGNRFAAFICTTLLGFTLALSIKSFYYIALL-QNP--
CYIQTL-PWFLGENLVIFWGFLFDSVTATMLVVVTSISFLVHFYSIEYMGADPH-------
LGRFMSYLSFFTFFMLILVTADNFVQMFVGWEGVGLCSYLLINFFFNRIQANKAAIKAMIMNRIGDFGLSLAIM
VIFYTCKSVDY--HTVFACVPFFIDST---
FMFFNFEVNLITCICILLFIGAVGKSAQIGLHTWLPDAMEGPTPVSALIHAATMVTAGVFLIIRCSFLFEFSDT
ALDILTIIGALTAFFAATTGLLQNDAKRVIAYSTCSQLGYMVFVCGFSGYSVSLFHLANHAFFKALLFLTAGSL
IHGMQDEQDFRKMGGLSYLLPYSYSMLLIGSLSLAG--FPF---------
LTGFYSKDMILELTFSSYLFKGTFAYTLGVVSAFFTAFYSFRLIYWAFLAKPN--GYKYSYANA----------
---------------------------------------------------
SESRFFITIPLAILAFASIFIGYILRDMFIGV-GTS----FWNNSIFILP-ENNFILESEF-
IPIHNKLTPVIFSLCGLFTAFFLYHIL-----------------------------------------------
------------FKKIFILNSPYIKFYTFLNKRWYFDKIYNEFIALPVISSGYKITFKIIDRGILELFGPAGIV
-ANLLNIARNTSNLQSGFMYHYLFLILLSITGAIVLIIGLLYNYFTLPIG
>YP_009092497_NAD5_Cyanoptyche_gloeocystis
MH-------LTVIILPLLSFVINTLFG----RSLGNSKIAIISTFNMAVTFIISVILFYETSII-GQV--
LNIDVS-QWIRSEYLIVNWGFLFDQLTTTMLIIVNSISFLVHLYSTEYMRNDPH-------
LGRFMSYLSFFTFFMLILVTANNYLQMFVGWEGVGLCSYLLINFFSARIQANKAAIKAMVLNRIGDFGLTIGML
MLLFVFKSLDY--SIIFSLAPYYTEEY---
IIIMNKSFSVLDLTTFFLLIGAIGKSAQLGLHTWLPDAMEGPTPVSALIHAATMVTAGVFMIIRSSPLFEFAPT
TLFTISIVGCLTAFFAASTGLLQNDLKRVIAYSTCSQLGYMVFACGVSAYNVAIAHLFNHAFFKALLFLTAGSI
IHSLADEQDLRKMGGLTQLLPLAYVMMLIGSLSLMG--FPF---------
LTGFYSKDFILEVTLSKYTINSAFCYWLGTLSAFFTAYYSFRLLYYTFLSKPN--GYKSTYNNI----------
---------------------------------------------------
HESNITMVLPLCALGFLSIFVGFAFKDMMVGI-GTN----FWDSSIFILPSIMNHINDAEN-
IQLFLKWLPVIFSILGAMLALLVYNYL-----------------------------------------------
------------KNILLSNLIMYKALYSFFNKKWYFDKIYNEIIAQYISMMGYHYTFKTLDRGIIEIIGPFGIA
-DTFIKISKSTSFLQSGYLHHYIYTIFFGAITLVILTSNYIKLQFWTDLR
>YP_009092462_NAD5_Gloeochaete_wittrockiana
MY-------LVIVFLPLVSAIMAGLFG----RFLGNKNSAYLATFCLGFTFFLCLIAFYEVGLC-HSP--
CYIVSF-PWIQSELFKVNWSFLFDSITVVMLIVVTSVSFLVHMYSIEYMGQDPH-------
LSRFMSYLSFFTFFMLILVTADNFLQMFVGWEGVGLCSYLLINFYFTRVQANKAALKAMIVNRIGDFGLSLGMM
ALFFTFKTLDY--NVVFNLAPLVVNEN---
FCFFSFELNKITVICLLLFVGAVGKSAQLGLHTWLPDAMEGPTPVSALIHAATMVTAGVFLIIRCSPLFELSAI
ALDVLMVFGSLTAFFAATTGLLQNDVKRVIAYSTCSQLGYMVFACGASSYSVAMFHLANHAFFKALLFLTAGAI
IHALRDEQDMRKMGGLAMLLPFAYSMIVIGSLSLMG--FPF---------
LTGFYSKDSILELVYSSYSSPAGFAYVLGTLAAFCTAFYSFRLIYLVFLSVPN--GYKSYFNQV----------
---------------------------------------------------
HETPYLMALPLVLLAFGSIFIGYLTKDLMLGM-GTN----FWGGSIYFLP-ERAGLFEAEF-
IPTNIKILPVCLSLLGAFTACLFYSSF-----------------------------------------------
------------FKFLVYLSVYVKDFYTFLNKKWYFDKIYNEVVGKFFIWFGYNMSFKLIDRGFVELFGPYGMS
```

```
-KLSFNLAKQFSLLQTGFLYHYIFMLLVGVTFFVGILSFGYYLNFWVDTR
>YP_004222736_NAD5_Glaucocystis_nostochinearum
MY-------LSIVFLPLLSAFVAGLLG----KFLGNKISSYFTTICLGITFILSIIAFYEVALC-ASP--
CYLTTI-SWIKSELFHVNWGFLFDTLTVVMLIVVTSVSFLVHMYSIEYMSHDPH-------
LSRFMCYLSFFTFFMLILVTADNFLQMFVGWEGVGLCSYLLINFYFTRIQANKSAIKAMIMNRIGDFGLSLGMM
AIFLTFKSLNY--DIIFTSVNLYSHEF---
FLFLNFEVNKITLICILLFVGAVGKSAQVGLHTWLPDAMEGPTPVSALIHAATMVTAGVFLIARCSPLFEYSTL
ALSILTIFGATTAFFAATTGLLQNDIKRVIAYSTCSQLGYMVFSCGISSYSAAVFHLANHAFFKALLFLTAGSI
IHSFQDEQDMRKMGGLGLILPYSYSMILIGSLSLMG--FPF---------
LTGFYSKDIILELAYSTYSIDSTFAYWLGTISAFFTAFYSFRLIYLVFLNKPN--GYKNAYQNA----------
---------------------------------------------------
DDSHFWIALPLSLLAFGSIFIGYLSKDMMLGL-GTN----FWGNSLFLLS-NHTTILDSEY-
IDYYLKLIPVIFSLIGSVCSYLLYRFS-----------------------------------------------
------------KNFLFTLNLRFKTIYIFFNKRWLFDKIYNEFIGLPALSFGYNISFKLLDRGFFEIFGPYGLA
-FVLLNFAKNTSRLQTGLLYHYIFIFILGLTFFVGLIKFGEQFFIYAFIF
>YP_009317212_NAD5_Palpitomonas_bilix
MY-------LLIVALPLISALTSGLFG----RFLGSKGAGFISVCCLIATSFLSFIAFYEVALN-GSP--
CYIKLA-TWVDSEMLHADWGFVFDSLTVIMLVVVTFVSALVHLYSTGYMEGDPH-------
VPRFMSYLSLFTFFMVMLVTADNFIQMFLGWEGVGLCSYLLITFWFTRVQANKAALKAFIVNRVGDFGLSLGVF
AIFYLFSSLDF--STVFALAPYMVGSQ---
LIFCGFEVDSLTLICILLFVGAVGKSAQLGLHTWLPDAMEGPTPVSALIHAATMVTAGVFMIARCSPLFEFAPT
ALFVVAIVGAMTAFFAATTGMLQNDLKKVIAYSTCSQLGYMVFVCGLSNYSVGVFHLANHAFFKALLFLSAGSV
IHAMSDEQDMRRMGGLAQIIPFTYVMMVIGSLSLMG--FPF---------
LTGFYSKDVILEIAYAKYSFAGTFSHWLGCISAFFTAFYSFRLLYLTFIQNTN--AYKRVIEHA----------
---------------------------------------------------
HESSFVMLLPLILLSFGSIFVGYLTKDMIIGL-GTS----FWGNSIFTLP-TNITMVDAEF-
LPYYIKLIPVILSLSGAVISLLVYHFMSQ---------------------------------------------
------------ILFNVQTSFIGRHLYIFFNKKWFFDKIYNYFFVQHVLRFGYSITFKSLDKGLIEFLGPYGIA
-NVVSRVSSLTSALHTGYIYHYAFVMFVGVVFILRI------------IF
>YP_316588_NAD5_Thalassiosira_pseudonana
MY-------LLLIFLPLIGSICAGLFG----RILGPFGSSFVTVICLMVTFCLSLFAFYEVALL-DCC--
VYIKLA-PWINSEMLNVDWGFLFDSLTVVMCCVVTFVSSIVHLYSTEYMAYDPH-------
LPRFMSYLSLFTFFMLILVTADNFIQMFVGWEGVGLCSYLLINFWFTRIQANKAAIKAMILNRIGDFGLVIGIL
IIFVEYKAVDY--ATVFALTPIFTNKV---
FHFLNFDFDLISLICFFLFIGAVGKSAQLGLHTWLPDAMEGPTPVSALIHAATMVTAGVFLIARTSPLFEYTSS
ILSLVTIVGACTAFFAATVGLLQNDLKRVIAYSTCSQLGYMVFACGLSNYSVGVFHLVNHAFFKALLFLGAGSI
IHAVADEQDMRKMGGLKKLVPFTYSMMVIGSLALIG--FPF---------
LTGFYSKDVILEVAYGKYTLEGHFSYILGTLGAFLTAFYSTRLIYLTFLSKPN--GYKSVICAA----------
---------------------------------------------------
YDSSYQISASLFFLVIPSVLIGFYAKDMIIGF-GSD----FWGNAIFTSI-ENMNRIDSEF-
ITHFYKILPVALSILGVTSSFLLYLFGSK----------------------------------------------
------------ILVKLKLSTIGKKIYNFFNKKWFFDKVYNEYVSQFFFTISYTVTYKTIDRGIIEIFGPMGLS
-SAITKKALYISKLQTGYLYHYTFLILTGLTLILGVRQFWVFLGDFIDFK
>YP_009050520_NAD5_Sargassum_fusiforme
MY-------LNIVFLPLLGSIIAGFFG----RFVGVYGSCFITVFCVGLTFCLSCLSFYEVGLA-GSG--
TYLSLM-TWLESDSFYVEWAFCFDSLTVVMLIVVTFISTLVHIYSTEYMGGDPH-------
LPRFVSYLSLFTFFMLILVTADNFVQMFVGWEGVGLCSYLLINFWFTRIQANKAAIKAMIVNRIGDFGLALGIF
AIFICFGALDY--ATVFAFSPQLTSCT---
```

```
LSFLNIKFKALDLIGVLLFIGAVGKSAQLGLHTWLPDAMEGPTPVSALIHAATMVTAGVFLLARCSPLLEYCSG
ALLLISVVGGMTAFFAATTALVQNDLKRVIAYSTCSQLGYMVFACGLSNYSVAVFHLANHAFFKALLFLSAGSV
IHAVGDEQDMRKMGGLRRLLPFTYAMMVIGSLALMG--MPF---------
LTGFYSKDVILEIAYATYSIPAHFSYWLGSLAAFCTAFYSIRLIALCFLVEPN--GSRKLLLSA----------
---------------------------------------------------
SEGGKAIGFVLGLLAIPSIFVGYFSRDLFIGL-GTH----FWGSSLFCLP-SNLSLIDAEF-
MSTFSKMLPLCLSIVGGALSFTLYRYYNY---------------------------------------------
------------NMYFWKTSLVGRKFYTFLSRKWFFDKIYNGFISQNILDFGYHFTYKSIDRGLIESLGPFGLS
-NVLTSQVQQARVWHSGYLYHYLVIFVWGNLLF----GLWFLFGFPSLRI
>YP_009254724_NAD5_Ectocarpus_siliculosus
MY-------LNIVFLPLLGSLVAGFFG----RFLGGRGSGLITISCVGLSFFLSGLAFYEVGLG-GSS--
TYLYLM-PWLESDSFHVDWAFCFDSLTVVMLIVVTFISTLVHLYSTEYMGGDPH-------
LPRFMSYLSLFTFFMLILVTADNFVQMFVGWEGVGLCSYLLINFWFTRIQANKAAIKAMLVNRVGDFGLALGIF
GVFICFGAVDY--ATVFSLAPQLSNFT---
LLFFNFEFNALNLIGILLFVGAVGKSAQLGLHTWLPDAMEGPTPVSALIHAATMVTAGVFLLARCSPLLEYCPD
ALFIISIIGGMTAFFAATTALVQNDLKRVIAYSTCSQLGYMIFACGLSNYSVAVFHLANHAFFKALLFLGAGSV
IHAVGDEQDMRKMGGLRRLLPFTYAMMVIGSLALMG--MPF---------
LTGFYSKDVILEVGYATYSSASHFAYWLGSFAAFCTAFYSIRLLALCFLSEPN--GSRTFLLSA----------
---------------------------------------------------
SEGSWPMGIVLGILAVPSMFIGYFSRDLFIGL-GTD----FWGNALFFLP-SNISIIDAEF-
MSTGIKILPLCLSILGGFSSFILYRNYSR---------------------------------------------
------------NLFLLKTSILGRKLYTFLNRKWLFDKIYNDFITQNTLDLGYHFTYKSIDRGLIESLGPFGFS
-NVLLDQVNKSRLWHSGYLYHYLVIL--GWSNLLF--GVWFIFGFKFLPI
>YP_005090357_NAD5_Phaeodactylum_tricornutum
MY-------LLVVFLSAIGSCFAGLFG----KHLGPLGSIFVTTSCLFLSLFISFFIFYEVALS-NSV--
VYIKLT-TWISSEVLHVDWGFMFDSLTATMCIVVISISFLVHLYSVEYMSHDPH-------
LPRFMSYLSLFTFFMLILVTADNFIQMFVGWEGVGLCSYLLINFWFTRIQANKAAIKAMIINRIGDFSLLIGII
LIFANYKSVDY--ATVAILSPFFKNSS---
ASFLNFNLELLTTIGIFLFLGAVGKSAQLGLHTWLPDAMEGPTPVSALIHAATMVTAGVFLIVRSSFIYEHTFS
VLEFIAILGATTSLFASTTGLLQNDLKRVIAYSTCSQLGYMVFACGLSNYSVSVFHLSNHAFFKALLFLSAGSV
IHAVNDEQDMRKMGGLKNLVPFTYSTMIIGSLTLMG--FPF---------
LAGFYSKDLILEVAYGKYNSTGHFCYFLGTTGAFLTAFYSIRLIYLTFLSKPT--GFKKTICCA----------
---------------------------------------------------
LDSGQSICIALGCLAILSLFIGYLTKDLLVGL-GSD----FFGNAIYVSV-KNLNMFDAEF-
IPSFYKTLPVSLSLCGLLLGFVLYNFKSK---------------------------------------------
------------ELFLLKISDFGKKYYSFLNRKWFFDKIYNNCLGQVFFRFGYSTSYKFIDRGIFETLGPTGLS
-LVSLRIASNLHKTQTGYLYHYTLAILIGLTLFISIRQMLFFMEHLLDYR
>YP_008999866_NAD5_Prasinoderma_coloniale
MY-------LVLLLLPTLGTLFGCFFG----RAMGGRGTAILTTTAVALSALLSFVGLYEVGYA-GSP--
CALSLG-RWVHSEAFDAEWGFLFDTLTMVMLCMITGVSSLVHLYSIGYMSADPH-------
LPRFMTYLSAFTIFMMLLVTGNNFLILFLGWEGVGLASYLLINFWFTRAQANKASIKAMIMNRVGDVGLALGIF
GIYALFKSLDY--ATVFSTAHLHAEDV---
LIFFGYEVHALTAIGCLLFIGAIGKSAQVGLHTWLPDAMEGPTPVSALIHAATMVTAGVFLLARVSPLLAYAPG
ALQVVTVVGATTALLGAVTGLTQNDMKRVIAYSTCSQLGYMVFAAGVGAYAVALYHMIIHAFFKALLFLCAGSV
IHAIGGEQDMRKMGGLARLLPLTYACMVVGSLALVG--FPY---------
LGAFYSKDLVLEAAYSRGMATGLLAYLLGIFAAFCTSYYSFRLLFLVFWGESR--TPKAGVRLA----------
---------------------------------------------------
HEGSWTMLIPLLVLTTCSIFGAEGLKEAFTGL-GTP----FWGGSLVAAG-GLTQSFEAEL-
```

VPLSAKLAPLVATGLGATLAYLVHLGPLRKL-------------------------------------------
------------ALKALTESQLGRTFYIFTNKRWFIDKIYAEVFGWSALFFGYHVTFKTIDKGLLEVFGPTGAK
-FSAASLIPALRLRSWTSLPHYASVLAFCLILLLSILGPASGMLSLGIGP
>ADU04611_NAD5_Mesostigma_viride
MY-------GTIIALPLIGSFCAGALG----RWLGFRGAAFITTLFVTISTIFSWIAFYEVALS-GSF--
CYIPVG-AWILSEMFDVSWLFLFDPLTVVMLIVVTSVSTLVHLYSISYMWSDPH-------
LPRFMSNLSIFTFFMLILVTAGNLVQMFLGWEGVGLASYLLINFWFTRLQANKSAIKAMLMNRVGDVGLALAIM
GIFYTFQSVDF--SVLLAL-SASEHSS--------
SSENFFTFVSLLLFVGSIGKSAQLGLHTWLPDAMEGPTPVSALIHAATMVTAGVFLIARCSALFEQSVHASFLV
ALVGAMTAFFAATTGAAQNDLKRVIAYSTCSQLGYMVFASGLSHYDLSVFHLMNHAFFKALLFLGAGSVIHALS
DEQDLRRMGGLVQLLPFTYTMMLIGSLSLVG--IPF---------
LTGFYSKDAILEVAYAKYTVSAHFAYWLGSLSVLFTSTYSFQLLQSAFLTSTN--AKRFALAKI----------
---------------------------------------------------
HESDLLLSIPLVLLALGSIFVGYLAKDMMIGL-GTD----FWSQSLFVQP-
YNVRLLEAEFATPISIKMVPLIFSFLGALIAHLLQNNKNVQTFIFS----------------------------
-----------------------
LLLFNPATDFLRSIYSFWLKRWLFDKVYNDFIVRPCMKFGYYVTFKTIDKGFLEWLGPLGIS-
YTIQKFGSTLSRFQSGYIYHYAFVMLIAFTTFISMQTPFAESTFQLQST
>NP_054400_NAD5_Marchantia_paleacea
MY-------LLIVILPLIGSFAAGFFG----RFLGSRGVAVVTTTCVSLSSIFSCIAFYEVALC-ASA--
CYIKIA-PWIFSELFDAAWGFLFDSLTVILLLVVTIVSSLVHIYSISYMSEDPH-------
SPRFFCYLSIFTFFMLMLVTGDNFIQLFLGWEGVGLASYLLINFWFTRIQANKAAIKAMLINRVGDFGLALGIM
GCFTIFQTVDF--STIFACASAFSEPHHY-
FLFCNMGFHAITVICILVFIGAVGKSAQIGLHTWLPDAMEGPTPVSALIHAATMVTAGVFMIARCSPLFEYSPN
ALIVITFVGAMTSFFAATTGILQNDLKRVIAYSTCSQLGYMIFACGISNYSVSVFHLMNHACFKALLFLSAGSV
IHAMSDEQDMRKMGGLASLLPFTYAMMLIGSLSLIG--FPF---------
LTGFYSKDVILELAYTKYTISGNFAFWLGSVSVFFTSYYSFRLLFLTFLAPTN--SFKRDLSRC----------
---------------------------------------------------
HDAPILMAIPLILLAFGSIFVGYLAKDMMIGL-GTN----FWANSLFILP-
KNEILAESEFATPTIIKLIPILFSTLGSFVAYSVNFVVNP----------------------------------
-----------------------
LIFALKTSTFGNRLYCFFNKRWFFDKVFNDFLARSFLRFGYEVSFKALDKGAIEILGPYGIS-
YTIRKMAQQISKIQSGFVYHYAFVMLLGLTIFISVIGLWDFISFWVDNR
>YP_009047585_NAD5_Sphagnum_palustre
MY-------LLIVTLPLLGSCVAGAFG----RFLSSRGTAIVTSTCVSLSFILSLIAFYEVALE-ASA--
CYIKMA-PWIFSEMFDASWGFFFDSLTVIMLIVVTFVSSLVHIYSISYMFEDPH-------
SPRFMCYLSIFTFSMLMLVTGDNSIQLFLGWEGVGLASYLLINFWFTRLQANKAAIKAMLVNRVGDFGLALGIM
GRFTIFQTVDF--STLFACVSAFSEPYHY-
LIFCNMKFHAITVICILLFIGAVGKSAQIGLHTWLPDAMEGPTPVSALIHAATMVTAGVFMIARCSPLFEYPPN
ALIVITFVGAMTSFFAATTGILQNDLKRVIAYSTCSQLGYMIFACGISNYSVSVFHLMNHAFFKALLFLSAGSV
IHAMSDEQDMRKMGGLASLLPFTYAMMFIGSLSLIG--FPF---------
CTGFYSKDVILELAYTKYTISGNFAFWLGSVSVFFTSYHSFRLLFLTFLAPTN--SFKQDILQC----------
---------------------------------------------------
HDAPILMAIPLIFLAFGSIFAGYLAKDMMIGL-GTN----FWANSLFVLP-
KNEIIAESEFATPTIIKLIPILFSTLGAFVAYNINFVANQ----------------------------------
-----------------------
FIFALKTSTLGNRLYCFLNKRWFFDKVFNDFIVRSFLRFGYEVSFKVLDKGAIEILGPYGIS-
YTFRKLAKQISKLQSGFVYHYAFVMLIGLTIFITIIGLWDFISFWVDNR

```
>AJF36693_NAD5_Klebsormidium_flaccidum
MY-------LLIVTLPLLGCATSALFG----RFLGSRGAAIVTTSCVGTSFLFSLVAFYEVALV-GSA--
CTLQFA-PWFDSEMFDASWGFLFDSLTVVMLIVVTLVSTLVHTYSISYMSEDPF-------
LPRFMCYLSIFTFFMLMLVSADNLIQMFLGWEGVGLASYLLINFWFTRLQANKAAIKAMLVNRVGDFGLALGIM
SCFLVFQSVDY--
HVLFAVANEWAMKESESFVFCNMHVDALSAICLLLFVGAVGKSAQIGLHTWLPDAMEGPTPVSALIHAATMVTA
GVFLIARCSPLFEYAPNALVVVTCMGAMTAFFAATTGILQNDLKRVIAYSTCSQLGYMVFACGISNYAVSIFHL
MNHAFFKALLFLSAGSVIHAMSDEQDMRKMGGLNQVLPFTYAMMLIGSLALIG--FPF---------
LTGFYSKDVILELAYTKYTISGHFAFWLGSLSVLFTSYYSFRLLFLTFLENTN--AFKQDIKNA----------
---------------------------------------------------
HDAPPLMAAPLIVLAFGSLFVGYLCKDMMIGL-GTD----FWGQSIFVLP-
DNSNLSESEFSTPQIVKLIPLLLSTLGAGIAYWINNHKNS----------------------------------
-----------------------
FVYDFKTSSLGRRFYTFLNKRWLFDKVLNEFLAQSVLRFGYSISFVTLDKGVIELLGPYGIA-
TTMRTLSKRTSGLQSGLIYHYAFIMLIGLTVLITFIGLWDIVSFWVDSR
>NP_943684_NAD5_Chara_vulgaris
MY-------LLIVVLPLLGSLVAGVFG----RFLGSRGAALVTTTCVSISSGLCFIAFYEVALG-ASA--
CYIKFA-PWILSEMFDASWGFLFDTLTVVMLIVVTFVSSLVHIYSISYMSEDPF-------
LPRFMCYLSIFTFFMLMLVTGDNLIQMFLGWEGVGLASYLLINFWFTRLQANKAAIKAMLVNRVGDFGLALGIM
GCFAIFQTVDF--STIFACASAFVW-EPS-
FIFLNMKIHALTAICVLLFIGAVGKSAQIGLHTWLPDAMEGPTPVSALIHAATMVTAGVFMMARCSPLFEYAPK
ALIVITFVGAMTSFFAATTGILQNDLKRVIAYSTCSQLGYMVFACGISNYSVSVFHLMNHALFKALLFLSAGSV
IHAMSDEQDMRKMGGLASLLPFTYAMMLIGSMSLIG--FPF---------
LTGFYSKDVILELAYTKYTISGNLAFWLGSLSVFLTSYYSFRLLFLTFLAPTN--AFKRDIERC----------
---------------------------------------------------
HDAPTLMAIPLILLAFGSLFVGYLAKDMMIGL-GTH----FWANSIFILP-
KNEIMFSSEFATPRMMKLIPILFSAIGAFLAYRVNFCANK----------------------------------
-----------------------
FIYVLKTSTLGTKCYCFLNKRWLFDKVFNDFIAKSFLRFGYEVSLKTLDKGAISILGPYGIS-
TTFRKLAKQISTLQSGFVYHYAFVMLIGLTIFITIIGLWDFISFWVDNR
>YP_009472097_NAD5_Arabidopsis_thaliana
MY-------LLIVFLPLLGSSVAGFFG----RFLGSEGSAIMTTTCVSFSSILSLIAFYEVALG-ASA--
CYLRIA-PWISSEMFDASWGFLFDSLTVVMLIVVTFISSLVHLYSISYMSEDPH-------
SPRFMCYLSIFTFFMLMLVTGDNFLQLFLGWEGVGLASYLLIHFWFTRLQADKAAIKAMLVNRVGDFGLALGIL
GCFTLFQTVDF--STIFACASVPRNS----
WIFCNMRLNAISLICILLFIGAVGKSAQIGLHTWLPDAMEGPTPVSALIHAATMVTAGVFMIARCSPLFEYSPT
ALIVITFAGAMTSFLAATTGILQNDLKRVIAYSTCSQLGYMIFACGISNYSVSVFHLMNHAFFKALLFLSAGSV
IHAMSDEQDMRKMGGLASSFPLTYAMMLIGSLSLIG--FPF---------
LTGFYSKDVILELAYTKYTISGNFAFWLGSISVLFTSYYSFRLLFLTFLVPTN--SFGRDISRC----------
---------------------------------------------------
HDAPIPMAIPSILLALGSLFVGYLAKDMMIGL-GWN----FWANSLLVLP-
KNEILAESEFAAPTIIKLIPILFSTLGAFVAYNVNLVADQ----------------------------------
-----------------------
FQRAFQTSTFCNRLYSFFNKRWFFDQVLNDFLVRSFLRFGYEVSFEALDKGAIEILGPYGIS-
YTFRRLAERISQLQSGFVYHYAFAMLLGLTLFVTFFCMWDSLSSWVDNR
>YP_588334_NAD5_Zea_mays
MY-------LLIVFLPLLGSSVAGFFG----RFLGSEGTAIMTTTCVSFSSILSLIAFYEVALG-ASA--
CYLRIA-PWISSEMFDASWGFFFDSLTVVMLIVVTFISSLVHLYSISYMSEDPH-------
```

```
SPRFMCYLSIFTFFMLMLVTGDNFLQLFLGWEGVGLASYLLIHFWFTRLQADKAAIKAMLVNRVGDFGLALGIF
GCFTLFQTVDF--STIFACASAPRNE----
WIFCNMRFNAITLICILLFIGAVGKSAQIGLHTWLPDAMEGPTPVSALIHAATMVTAGVFMIARCSPLFEYSPT
ALIVITFAGAMTSFLAATTGILQNDLKRVIAYSTCSQLGYMIFACGISNYSVSVFHLMNHAFFKALLFLSAGSV
IHAMSDEQDMRKMGGLASSFPLTYAMMLMGSLSLIG--FPF---------
LTGFYSKDVILELAYTKYTISGNFAFWLGSVSVLFTSYYSFRLLFLTFLVPTN--SFGRDRLRC----------
---------------------------------------------------
HDAPIPMAIPLILLALGSLFVGYLAKDMMIGL-GTN----FWANSPFVLP-
KNEILAESEFAAPTITKLIPILFSTSGASLAYNVNLVADQ----------------------------------
-----------------------
FQRAFQTSTFCNRLYSFFNKRWFFDQVLNDFLVRSFLRFGYSVSFEALDKGAIEILGPYGIS-
YTFRRLAERISQLQSGSVYHYAFAMLLGSTPFVTFSRMWDSLSSWVDSR
>ANA57055_NAD5_Pyramimonas_parkeae
MY-------LLLVVLPLMGSLFAGFFG----RYLGYRGAGLFSSTCVGLSAIGSWVAFYEVGLS-GSP--
CYLKFA-PWFHSEFFDASWGFLFDSLSVVMLVTVTTASTCIHLYSISYMGEDPH-------
LPRFMSYLSIFTFFMLVLVTADNFIQMFLGWEGIGLASYLLINFWVTRLQANKAALKAMLVNKIGDFGLALGIL
GVFQLVKTVDY--ASLFACVPYINESS---
ICFFQWELDGLTLIACLLFLGVVGKSSQIGLHTWLPDAMEGPTPVSALIHAATLVTAGVFLMARISPLLEYAPQ
ALSLIAFTGAMTCFLAASTGIMQNDLKRVIAYSTCSQLGYMVFACGLSSYGVSIFHLMNHAFFKALLFLSAGSL
IHALADEQDMRKMGGLGRLLPYTYGMILIGSLSLAG--FPF---------
LTGFYSKDVILEMAYAKYSLSGNFAYWLGALSVFFTSYYSFRLISLAFFAPTN--CIKGSIRTV----------
---------------------------------------------------
HDAPFLLGAPLIPLALGSLFLGFLGKDMMIGA-GST----FWGNALFILP-QNSLSIESEW-
IPQNVKLLPLVLTIGGAITAYLINLVWIE---------------------------------------------
------------SAFKIKTHWLGREIYLFLNKRWLFDKVYNEFIAKTFMQFGYNVSFKSLDKGAFEIIGPYGLI
-QSVPILLKETSRFQSGFLYHYALIMLIGITFLIGLIGLGPILNYWIDPR
>QIQ59668.1_Nad5_Trebouxia_sp._A1-2
MY-------LSLIFLPLLGSLCAGFFG----RFLGFRGACLISTTSVFSSFIMSTVAFYEVALS-GSS--
CYIKYS-SWFVSEMFDSSWGFYFDTLTVVMLVVVTSVSTLVHLYSISYMSGDPH-------
LPRFMSYLSIFTFFMLVLVTADNFLQMFFGWEGVGLASYLLINFWFTRLQASKASIKAMLVNRVGDFGLSLGIM
AIFSVFKSVDF--ATVFSCAPHFADTE---
FLFCNIECTLLNVVCILLFIGAVGKSAQLGLHTWLPDAMEGPTPVSALIHAATMVTAGVFLIARCSPLFEYAPN
ALMIVTLVGGMTTFFAATTGIVQNDLKRVIAYSTCSQLGYMVFACGISNYSVGVFHLMNHAFFKALLFLSAGSV
IHALSDEQDMRKMGGVVQLLPFTYGMMLIGSLSLTG--FPF---------
LTGFYSKDTILELAFASYTVNGNFAYWLGAICVLFTSYYSFRLLFLTFLSPTN--LYKSSLKNT----------
---------------------------------------------------
HDANFIMALPLILLAFGSIFVGYLGKDMMIGL-GTN----FWGNALFVLP-KNINLLESEY-
IPQVQKMVPLFFTLLGAFLAYLVNYSLFR---------------------------------------------
------------ETYFIKTSWIGRKFYYMLSKRWLFDKVYNDFIGQKNLDFGYRISFKTLDKGCFEILGPYGIS
-FLFQNLTRYLSKLQSGMIYHYAVVMLLGVILLISIVGLWEFLEVFVDNR
>YP_717300.1_nad5_Ostreococcus_tauri
MY-------LLLVFLPFFGFLLVTVFG----RFFGYRGAPLLSTFAVMSSACLSFVALYEVGIC-GSP--
CYVQFL-PWFQTDMFVASWGFLFDSLTVIMCVVVTFVSSLVHIYSISYMGEDPH-------
LPRFMSYLSVFTFFMLMLVTSDNFLQMFFGWEGVGVASYLLINFWYTRLQANKSAIKAMLVNRVGDFGLALGIM
GIFHIFKAVDF--ETVFACASEFACHH---
FLFFHMEVHTLTCISLLLFIGAIGKSAQLGLHTWLPDAMEGPTPVSALIHAATMVTAGVFLLARCSPLVEYSSQ
ALVVITLVGASTAFFAATTGVVQNDLKRVIAYSTCSQLGYMMFACGISQYALGVFHLMNHAFFKALLFLSAGSV
IHALGDEQDMRKMGGLVRLLPFTYSMMMLGSLALIG--FPY---------
```

```
LTGFYSKDVILEVAYAKYTLSGNFAYWLGSVSALFTSYYSFRLLYLTFLASPQ--MLKSSVQNV----------
---------------------------------------------------
HDAPFLMGLSLCCLALGSIFGGFIAKDMMIGL-GTN----FWGNAVFTLP-TNTLVLESEY-
IPQAVKFVPIIFSILGAVFAVNLNGFGSS---------------------------------------------
------------FSYSLKTSVLGKQLYTFLNKRWLFDKVYNDFVARPSLHFGYTISFKLVDKGFLEMFGPTGIM
-QQSAFMRHFVKTVQTGSIYHYAFMMFVSLTSLAFFLVATTSVGAFLDTR
>YP_008994799_NAD5_Bathycoccus_prasinos
MY-------LTILFLPLLGFLLAASGG----RFFGWRGTPLITTFCVFLSSLFSFIAFYEVGIA-GSP--
CYIQLA-PWFTSEFFDATWGFMFDSLTVVMLVVVTFVSTLVHIYSISYMSEDPH-------
LPRFMSYLSIFTFFMLMLVTADNFIQLFFGWEGVGLASYLLINFWYTRLQANKSAIKAMLVNRVGDFGLALGII
ATFSLFKSVDF--ATVFACSAHFAENS---
FIFFHLEWHALSLICALLFVGAVGKSAQLGLHTWLPDAMEGPTPVSALIHAATMVTAGVFMIARCSPLFEQAPQ
TLILVTVTGALTAFFAATTGVVQNDLKRVIAYSTCSQLGYMVFACGISQYAVGVFHLMNHAFFKALLFLSAGAV
IHALADEQDMRKMGGTIKVLPFTYSMMFIGSLALIG--FPF---------
LTGFYSKDVILEVAYAKYTVAGTFTYWLGAVSALFTSYYSFRLLFLTFIGPVN--SLKDSLKHV----------
---------------------------------------------------
HDAPFLMGFPLFLLAFGSIFVGYLAKDMMIGM-GTQ----FWGNALFMGP-DHGLLVESEY-
IPQAIKYIPILFSFLGFLFALNLNLLGAS---------------------------------------------
------------FSYSLKTSTFGRTLYIFLNKRWLFDKMYTDFVALPALFFGYNVSFKTLDKGLLEIVGPAGVI
-QTVSQGGRQLRKMQSGLIYHYAFIMLCSLTFLIAFVGLWDFFSFFVETH
>YP_009121385_NAD5_Thecamonas_trahens
MY-------SLIIFLPLIGSFAAGFFG----RLIGEKGAAIITTSAMVFSTLFSYVALYNLVVT-GNP--
VYIDLF-TWIDSGLFEARWGFTFDSLTVIMLIVVTTVSTLVHLYSTEYMAGDPH-------
VPRFMSYLSLFTFFMLMLVTSNNLIQLFFGWEGVGLASYLLINFWFTRIPANKAAIKAMIVNRVGDFGLSLGIM
AIFFIFKSVDF--ATIFALIPHFDGYT---
FVFFNDTYNIFNVICLLLFIGAVGKSAQIGLHTWLPDAMEGPTPVSALIHAATMVTAGVFLLIRCSALFEYAPD
VLLIVTFVGAVTALFAATAGMLQNDLKKVIAYSTCSQLGYMVFACGISNYSVSMFHLMNHAFFKALLFLSAGSV
IHAMADEQDMRKMGGIINLLPFTYTMILIGSLSLMG--FPF---------
LTGFYSKDVILELAFAKYTITSSFSYWFGTLAAFFTAFYSFRLVYLTFISNTN--AYKKVMEQA----------
---------------------------------------------------
HDAPIRMAIPLIILLFGSIFVGYLTRDMIIGL-GTD----FWGNAIFILP-SNLNMIDSEF-
IPWNIKLIPVIFSLIGASLAFILNAFYDR---------------------------------------------
------------FLVDFKLSKLGLRLYGFINKKWYVDNIYNEFIVKNFLAFGYHVSFKTIDRGLIEYIGPYGIV
-KLAQKFSNTISKLQTGYVVYNYTAMTLIGSILFITLIGLWDILNVFVDHR
>BAA78087_NAD5_Dictyostelium_discoideum_3D
MY-------IVNLILPLIGSIITGIFG----HKLGNRISIKIAVGCMMLTAISSLYIGYEILLC-NSV--
VHFKLG-TWMQVGSLNVEYGLLYDSLTSIMIIVITCISSMVHLYSMDYMKEDPH-------
KTRFFSYLSLFTFFMMLLVTADNFVQLFFGWEGVGIMSYLLINFWYTRLQANKSALKAVILNRFGDFGLFFGIL
LVFLVFKSVDF--SVIFTIAPFITEYT---
INLLGYEVNAITLIGSFIVIGVVGKSAQLGLHMWLPDAMEGPTPVSALLHAATMVTAGVFLVLRTSPLLSYSIT
ILNILTIIGALTTLFATTIGIVQNDIKRVIAYSTCSQLGYMIFACGLLNYNASIYHLTTHAFFKALLFLSAGSV
IHGLNDEQDMRKMGGLVNLMPLTYQCMLIGTLALTG--FPF---------
LSGYYSKDIILETSYATYYWEGTFAAIIGYVAAFGTTFYSFRLLILTFFNKPR-MQYKTIAGV-----------
---------------------------------------------------
HEASTNMVIPLVILALCSIFIGYVTKDFFVGL-GTP----VWNNSFFAYP-YNNLILESEV-
LQRELKLLPLFAFIYGVITPVLFYFNIKEID-------------------------------------------
------------
RMINVKQNLMVKESYFFFVKKWYFDFLSRVLIVVPFFHLSYDVMNKNLDKGLWEKIGVTGVAKNEITLSTYISY
```

```
IVQTIILIIVVGIFSFMTGFIYMELCIIIGILYICLPS
>YP_009476712_NAD5_Proteomonas_sulcata
MY-------LTIIFLPLVGCFFSGFFG----RFLSPKGSAFITILGIFIAFIFSLVAFFEVGLS-LCP--
CYIQLL-PWIDCELFDASWGFMFDTISVGMCVTVTFISLLVHIYSTSYMSHDPH-------
QPRFMAYLSLFTFFMLCLVTADNFIQLFLGWEGVGLCSYLLINFWFTRLQANKAAIKAMIINRIGDVGLALGVF
AAFIVFKTTNF--SVIFSLVPYYVNHE---
IFFFTYKCNAITLIGLLLFIGSVGKSAQLGLHTWLPDAMEGPTPVSALIHAATMVTAGVFLIIRCSPLYEYSES
ALFIIVIFGALTAFFAATIGLLQNDLKKIIAYSTCSQLGYMVFACGLSSYSVSMFHLVNHAYFKALLFLSAGSV
IHALNNEQDMRRMGGLLQVLPFTYCMFVVGSLALMG--FPF---------
LTGFYSKDAILEIAYAKYNFIGTFAHWLGIISAFFTAFYSFRLIYFTFVISPN--GYKTNMQNA----------
---------------------------------------------------
HDAPLLMGLPLFLLSVASIFIGYLTRDLYIGL-GTP----FWNNAIYTAP-QNLSYIDAEF-
IPTSIKWLPVIFSLLGAILSIFLYHNCKQ---------------------------------------------
------------ILYNLTSISLFRYIYAFLNRRWLFDRLQNDFIGDSFLKIGYSITYKKLDKGIIEVIGPKGII
-NLTTNIGKQVISLQSGLIHQYASLIFVGVIFIVIKITNFNFFHPFIDIH
>BBD14148_NAD5_Ophirina_amphinema
MY-------LLIVFIPLVTSLVSGLGG----RFIGKLGSQVLTSLGVLMSATLSSVAFYEVGIC-GSP--
CYVEFL-PWISSDMLRVYWGFQFDSLTVVMLVVVTYVSTLVHLYSIGYMSEDPH-------
IPRFMSYLSLFTFFMLMLVTGDNLVQLFFGWEGVGVCSYLLISFWYTRIQANKAALKAMVMNRVGDFGLSLGIF
ASFYVFKSLDY--ATIFSLAPYYVDFQ---
IPFLYWSFGSLNLIGILLFVGAVGKSAQLGLHTWLPDAMEGPTPVSALIHAATMVTAGVFLISRCSPLYEYAPK
ALLVVTIIGGTTAFFAATTGLVQNDLKRVIAYSTCSQLGYMIFACGLSGYSIGMFHLMNHAFFKALLFLSAGSV
IHALADEQDMRKMGGMLNIVPLTYTMMLIGSMSLMG--FPF---------
LTGFYSKDVILEFSFAKYTIDSSFVFWLGVISAFFTAFYSFRLIYLTFFGEPN--SIRSVVSKS----------
---------------------------------------------------
HDAPVIMALPLMLLCLGSVFVGYIFRDLIIGL-GTD----FWGSALYVSP-HNSAHIDSEF-
IPHMVKLMPVCLSVLGALASLYVYEKLAL---------------------------------------------
------------EIGSLYKTAIWLSVYKFLNGRWLFDSVYNYFVAEKLMSFGYHISFKTLDKGIIEFMGPSGLI
-SLISTWSIRVSRLNTGYLYHSAFLMMLGAVIVL---GVTRVCQMESDIA
>YP_007890522_NAD5_Andalucia_godoyi
MY-------LLIVFLPLLGSILAGLFG----RFLGRHGASLVSTTSVGLTCLFSWIAFYEVGIC-ESP--
VYLTLL-PWIDSGMLTANWGFLFDSLTVVMLIVITTVSFLVHIYSTSYMSEDPH-------
LPRFMSYLSFFTFFMLMLVTADNFVQMFLGWEGVGLCSYLLINFWFTRLQANKAAIKAMIMNRIGDFGLSLGMM
ALFAVFQALDF--STLFAIAPLFANQD---
FLFLNFQVDLLTTICILLFVGSVGKSAQLGLHTWLPDAMEGPTPVSALIHAATMVTAGVFLIVRCSPLFEYAPS
ALLVVTFVGAMTAFFAATTGMLQNDLKRVIAYSTCSQLGYMVFACGLSSYSVSMFHLMNHAFFKALLFLSAGAV
IHALADEQDMRRMGGLVQLLPFTYSMMLIGSLALMG--FPF---------
LTGFYSKDVILELAYAKYSLDGTFAHWLGTVSAFFTSFYSFRLLYLTFLTKTN--AYKSSVSHA----------
---------------------------------------------------
HDAPFVMAFPLMVLALGSLFVGYLFRDMMIGL-GTS----FWGNSLFVLP-EHLILIDSEF-
IPYSIKSIPVLLSFVGASLAWLLNHSYGA---------------------------------------------
------------FLYELKTSSFGRDLYTFLNKRWLFDKVYNDYVGKVLLQFGYNVSYKTLDKGILEILGPYGII
-RLVRHLSDRVSSFHTGYLYHYAFIMLIGVTLLISVIRFWDVLSQVVDPR
>AGH24310_NAD5_Reclinomonas_americana
MY-------LLIVFLPLLGSITAGFFG----RSLGKQGAAIITTSCVALSSLFSMVAFYEVGLC-GSP--
CYIRLF-NWIDSEMLHASWGFLFDSLTVVMLIVVTIVSSLVHLYSVGYMSHDPH-------
LPRFMSYLSLFTFFMLMLVTGDNFVQMFLGWEGVGLCSYLLINFWFTRLQANKSAIKAMIMNRIGDFGLSLGMM
AIFFIFKSVDF--ITVFALSPYMTDAT---
```

```
IVFLNYEVHALTLICILLFVGAVGKSSQLGLHTWLPDAMEGPTPVSALIHAATMVTAGVFLIARCSPIFEYAPT
ALLVVTIVGAMTAFFAATTGLLQNDIKRVIAYSTCSQLGYMVFACGISGYSVGMFHLMNHAFFKALLFLSAGCV
IHALADEQDMRRMGGIVKIVPFTYGMMLIGSMSLMG--FPF---------
LTGFYSKDVILELAFAKYTIDGTFAHWLGTVAAFFTAFYSFRLIYLTFLGETN--APRTIINHA----------
---------------------------------------------------
HDAPFIMALPLMILAIGSIFVGFIMKDMMIGL-GTD----FWGNSLFTHP-KNLTLIESEF-
IPTPIKLLPVILSIVGASLAIILNNFYAT---------------------------------------------
------------FLVSLKTSLLGREIYSFLNKRWYFDIVYNEYVGKTLLWFGYNISFKSVDKGLIEILGPYGLE
-RLTRRLTSKVSALQTGYIYHYAFIMLLGVTLIITIIGLWDYISMWTDYR
>YP_009245579_NAD5_Diphylleia_rotans
MY-------LSIIVLPGIAALLAGFFG----RFLGPKGSSLITTTLVGITCFLSLTSLYEVGIL-GSV--
CEIKLA-KWIESDLLDADWGFLFDSLTVSMLVVITSISTLVHLYSIDYMSEDPH-------
LPRFMSYLSFFTFLMIILVTADNYLQMFVGWEGVGLCSYLLINFWFTRIAANKSAMKAIIVNRIGDLGLILGII
CLMTTFKSVDY--ETIFSIVSGSDLIT-------
GVPSSILTLTTLLLFIGAVGKSAQIGLHTWLPDAMEGPTPVSALIHAATMVTAGVFLIIRSSPLFIVAEDILVL
MTIMGSLTALFAATSGLLQNDLKRVIAYSTCSQLGYMVFACGITSFATSFFHLANHAFFKALLFLSAGAVIHAL
GDEQDMRKMGGLIRLLPFTYVMILIGSLSLMG--FPF---------
LTGYYSKDVILEFAYSFVSFDSSFAYWLGTISAFFTAFYSIRLIYLTFITETN--AYHSIIKNV----------
---------------------------------------------------
HEVPLKMAIPLMILAFGSIFIGYLTKDLIIGF-GTP----FWNNAIVLLP-N-----DAEF-
LPFFIKLIPVIFSLFGAFLGYTLYSSNI----------------------------------------------
------------LWNMPEQGSVLRFVYSFFNQKWYFDQLYNRFCVQPLLKIGYHTTFKLLDKGIIEILGPYGIA
-SIIRYTVDRARYLQSGLIYHYTFIMVSGLILLLLVISQKEFSIITPFPI
>YP_009446429_NAD5_Ancoracysta_twista
MY-------LLIVFLPLLSALTVLSFG----RYLGREGSVLLTVSSLFITAAMSTTAFYEVALC-GSV--
CHIKLA-PWITSELLEISWGFLFDSLTAVMLVVVTYVSALVHLYSVSYMEEDPH-------
LPRFMSYLSLFTFFMLMLVTGDNFLQLFLGWEGVGLASYLLINFWYSRIQANKSAIKAMVMNRIGDFGLALGLM
AIFAVFKSIDY--STVFATAPFFSNEN---
FLFCNLELDQLTVICLLLFVGAVGKSAQLGLHTWLPDAMEGPTPVSALIHAATMVTAGVFMICRCSPLFEYAPD
ALFVITLVGASTAFFAATVGVVQNDLKRVIAYSTCSQLGYMVFSCGLSSYSVSMFHLMNHAFFKALLFLSAGSV
IHAMGDEQDMRRMGGLLNLLPFTYTMMLIGSLSLVG--FPF---------
LTGFYSKDVILELAYGKYTVSGTFAHWLGSLSVFFTSFYSFRLIYLTFINKTN--SHKSLIEGA----------
---------------------------------------------------
HDAPIIMAIPLMILAVGSIFVGYVTKDLMIGL-GTT----FWSHSLFVLP-VNMTSVESEF-
LDASIKCIPLFFSGMGATVAFILCYANRR---------------------------------------------
------------LNFRLTSNPVGRGIYTFLNKRWLVDAVYNVYGAAPLMRFGYSTSFRLLDTGFIQMFGPRGLS
-TLLLELSSLASRFQSGYVYHYAFIMVCGATILVGFINLWPFTSLYLSNK
```
